# AI-Enhanced Adaptive Virtual Screening Platform Enabling Exploration of 69 Billion Molecules Discovers Structurally Validated FSP1 Inhibitors

**DOI:** 10.1101/2023.04.25.537981

**Authors:** Domiziana Cecchini, AkshatKumar Nigam, Ming Tang, Joana Reis, Matt Koop, Andrea Gottinger, Callum Robert Nicoll, Yao Wang, Abhilash Jayaraj, Süleyman Selim Çınaroğlu, Ricarda Törner, Yehor Malets, Minko Gehev, Krishna M. Padmanabha Das, Kelly Churion, Jongwan Kim, Nidhin Thomas, Yong Li, Hyuk-Soo Seo, Sirano Dhe-Paganon, Christopher Secker, Mohammad Haddadnia, Alexander Hasson, Minkai Li, Abhishek Kumar, Roni Levin-Konigsberg, Eun-Bee Choi, Geoffrey I. Shapiro, Huel Cox, Luke Sebastian, Chelsea Braithwaite, Puspalata Bashyal, Dmytro S. Radchenko, Aditya Kumar, Lei Yang, Pierre-Yves Aquilanti, Henry Gabb, Amr Alhossary, Gerhard Wagner, Alán Aspuru-Guzik, Yurii S. Moroz, Charalampos G. Kalodimos, Konstantin Fackeldey, John D. Schuetz, Andrea Mattevi, Haribabu Arthanari, Christoph Gorgulla

## Abstract

Identifying potent lead molecules for specific targets remains a major bottleneck in drug discovery. As structural information about proteins becomes increasingly available, ultra-large virtual screenings (ULVSs) which computationally evaluate billions of molecules offer a powerful way to accelerate early-stage drug discovery. Here, we introduce AdaptiveFlow, an open-source platform designed to make ULVSs more accessible, scalable, and efficient. AdaptiveFlow provides free access to a screening-ready version of the Enamine REAL Space (1), the largest library of ready-to-dock, drug-like molecules, containing 69 billion compounds that we prepared using the ligand preparation module of the platform. A key innovation of the platform is its use of a multi-dimensional grid of molecular properties, which helps researchers explore and prioritize chemical space more effectively and reduce the computational costs by a factor of approximately 1000. This grid forms the basis of a new method for identifying promising regions of chemical space, enabling systematic exploration and prioritization of compound libraries. An optional active learning component can further accelerate this process by adaptively steering the search toward molecules most likely to bind a given target. To support a broad range of applications, AdaptiveFlow is compatible with over 1,500 docking protocols. The platform achieves near-linear scaling on up to 5.6 million CPUs in the AWS Cloud, setting a new benchmark for large-scale cloud computing in drug discovery. Using this approach, we identified nanomolar inhibitors of two disease-relevant targets: ferroptosis suppressor protein 1 (FSP1) and poly(ADP-ribose) polymerase 1 (PARP-1) (2, 3). By leveraging newly solved crystal structures of FSP1 in complex with NAD^+^, FAD, and coenzyme Q_1_, we validated these hits experimentally and determined the co-crystal structures of FSP1 bound to small-molecule inhibitors, enabling insights into inhibitor binding mechanisms previously unknown. With its high scalability, flexibility, and open accessibility, AdaptiveFlow offers a powerful new resource for discovering and optimizing drug candidates at an unprecedented scale and speed.

## Introduction

A central challenge in early-stage drug discovery is the identification and optimization of initial hit and lead compounds that bind specifically and potently to biological macromolecules. Historically, experimental high-throughput screening (HTS) has served as the primary method for discovering such hits (4). While these approaches have yielded many successful leads, they are inherently constrained by high costs, long timelines, and the need for robust, scalable assays. These limitations are further compounded by the vastness of chemical space: the number of synthetically accessible, drug-like small molecules is estimated to exceed 1060 (5), yet typical HTS campaigns screen only hundreds of thousands of compounds. This minuscule sampling poses a major bottleneck - particularly for challenging targets such as protein–protein interaction interfaces or allosteric sites, where binding pockets are often shallow, dynamic, or poorly defined. Virtual screenings have emerged as a compelling alternative, offering the ability to computationally evaluate extremely large numbers of molecules. Virtual screening surpassed millions of compounds over a decade ago (6) and, in recent years, has expanded to hundreds of millions to billions of molecules in silico, giving rise to ultra-large virtual screens (ULVSs). Compared to traditional HTS, ULVSs dramatically reduce cost and time, while expanding the scope of chemical space that can be explored. In addition to identifying potent candidate compounds, virtual screening can provide insights into binding mechanisms and structure–activity relationships, making it especially valuable for targets that are difficult to modulate using conventional experimental approaches.

In recent years, multiple studies have demonstrated the remarkable effectiveness of ULVSs in identifying potent hits across a wide range of protein targets, including enzymatic active sites, orthosteric sites of GPCRs, and protein–protein interaction interfaces. In 2019, Lyu and colleagues (7) targeted the D4 dopamine receptor and AmpC in a pioneering work using ULVSs and obtained picomolar binders directly from the screen. In 2020, the melatonin receptor MT1 was successfully targeted with ULVS, resulting in compounds with sub-nanomolar potency (8). In the same year, ULVSs were used for the first time to target protein–protein interactions. That study performed the first screen of one billion compounds to target the KEAP1-NRF2 interaction, leading to the identification of inhibitors in the sub-micromolar range (9). In 2021, the σ2 receptor was targeted by ULVSs, leading to low-nanomolar ligands (10). Notably, in 2020, the first open-source, fully automated platform dedicated to ULVSs called VirtualFlow was published (9, 11). This platform allows users to routinely perform ULVS with a library of approximately 1.4 billion compounds in a “ready-to-dock format”. In 2020, there were approximately 2 billion on-demand molecules available. Driven by the success of billion-compound screens and advancements in synthesis and computational technologies, the number of available on-demand molecules, with an established synthetic route, has expanded rapidly, with current estimates reaching up to 3 trillion compounds across multiple vendors (1).

While ultra-large virtual screenings (ULVSs) enable the exploration of vast chemical libraries, significant challenges remain in making these searches efficient, affordable, and intelligent. Navigating such immense chemical space to rapidly identify promising hits requires new strategies that go beyond brute-force docking. The financial cost alone is a major bottleneck - docking one billion compounds using standard cloud-based CPU providers typically costs $20,000–30,000, (9) rendering it impractical as chemical databases continue to grow exponentially, with newer libraries exceeding one trillion compounds, and growing (7). Moreover, effective virtual screening requires the ability to flexibly integrate a wide range of docking algorithms, each built on different theoretical foundations and requiring distinct input formats, to better match the characteristics of diverse target classes. Compounding these challenges is the need to fully leverage modern computational infrastructure: while many existing platforms are limited to CPUs, there is increasing demand to exploit both CPU clusters and GPU architectures to scale performance and reduce turnaround time. These limitations - high cost, limited flexibility, and insufficient scalability - highlight the urgent need for a new solution. To this end, we developed AdaptiveFlow, an open-source platform designed to intelligently navigate ultra-large chemical spaces, seamlessly incorporate diverse docking protocols, and operate efficiently across heterogeneous computing environments, including cloud-scale CPU and GPU resources.

AdaptiveFlow is a next-generation, open-source platform purpose-built for navigating ultra-large chemical libraries comprising 69 billion drug-like molecules with scale, speed, and intelligence. A key innovation of the platform is the implementation of Adaptive Target-Guided Virtual Screens (ATG-VSs), which use an 18-dimensional grid of molecular properties to identify and hone in on the most promising regions of chemical space, favorable for a chosen target site, substantially reducing computational cost relative to exhaustive docking. An optional active learning layer can be added to further refine and accelerate this process by iteratively selecting the most promising compounds of each tranche. To demonstrate the utility of AdaptiveFlow, we applied it to two therapeutically relevant targets: ferroptosis suppressor protein 1 (FSP1), a key regulator of lipid peroxidation and ferroptotic cell death with emerging roles in cancer resistance (2), and poly(ADP-ribose) polymerase 1 (PARP-1), a well-established DNA repair enzyme and clinical target in oncology (3). In both cases, the platform identified nanomolar inhibitors, which were experimentally validated through high-resolution co-crystal structures confirming target engagement.

### AdaptiveFlow

Deploying ultra-large virtual screenings (ULVSs) at the scale of billions of compounds requires not only massive computational resources but also software systems that can efficiently coordinate the preparation, management, and docking of libraries across heterogeneous high-performance computing (HPC) environments. To meet these requirements, we developed AdaptiveFlow, an open-source platform built to streamline, scale, and intelligently guide ULVS workflows from end to end. AdaptiveFlow combines deep flexibility with performance, offering support for diverse ligand and receptor types, GPU acceleration, and modular integration of classical and machine learning-based docking protocols.

AdaptiveFlow introduces major enhancements across three core components: the AdaptiveFlow Ligand Preparation (AFLP) module for efficient preprocessing of massive libraries; the AdaptiveFlow for Virtual Screening (AFVS) engine, which supports over 1,500 docking protocols; and the AdaptiveFlow Unity (AFU) module, which integrates these components into a unified, modular workflow adaptable to both CPU and GPU-based infrastructures (overview in Fig. 1 and in Extended Data Table 1). These modules have been engineered to support both traditional and emerging use cases, while optimizing for large-scale parallel execution on cloud and on-premise infrastructures. AdaptiveFlow is natively compatible with cloud services such as AWS (Extended Data Fig. 1), where it demonstrates near-linear scaling up to 5.6 million virtual CPUs (Extended Data Fig. 2), enabling efficient exploration of ultra-large libraries. The use of spot (preemptible) instances further reduces the costs.

**Fig. 1.**
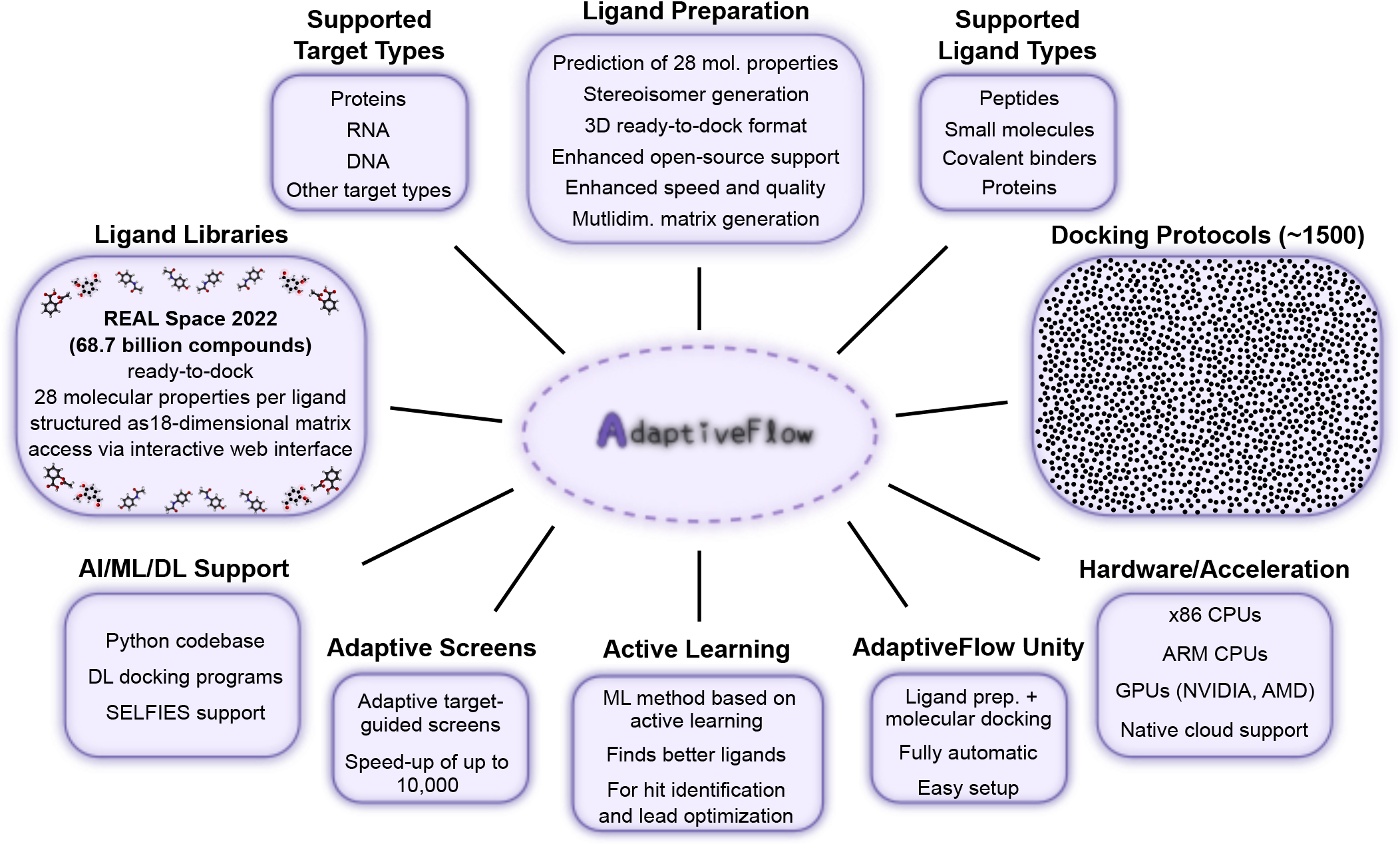
Overview of features of AdaptiveFlow. AdaptiveFlow introduces several key advances that significantly expand the capabilities of ULVS, including, i) curation of the Enamine REAL Space library, comprising 69 billion molecules in a ready-to-dock format compatible in multiple chemical file formats, ii) integration of an extensive suite of approximately 1,500 docking protocols (defined as specific combinations of sampling algorithms and scoring functions), enabling the targeting of a broad range of receptor types, including RNA and DNA, iii) support for emerging hardware architectures such as ARM CPUs, along with GPU acceleration to enhance computational performance, iv) advanced ligand preparation features, including stereoisomer enumeration and the calculation of 28 molecular properties, to enable the creation of more comprehensive screening libraries, and v) built-in support for machine learning and deep learning approaches, including compatibility with SELFIES and neural network–based docking protocols. The integrated module, AdaptiveFlow Unity, further streamlines workflows by combining ligand preparation and docking into a single environment. Importantly, AdaptiveFlow also introduces a novel screening paradigm known as Adaptive Target-Guided Virtual Screens (ATG-VSs), which enables efficient, target-focused exploration of ultra-large compound libraries at a fraction of the computational cost typically associated with conventional ULVSs.

The AFLP module provides a powerful pipeline for preparing massive compound libraries for docking. In addition to generating 3D conformers, it enumerates stereoisomers and tautomers, validates geometries, and calculates 28 molecular properties (see Extended Data Table 2, 3; Supplementary Fig. 1, 2, 3). These properties are used to disperse the molecules within a multidimensional grid, up to 18 dimensions, based on physicochemical descriptors. This grid-based representation forms the foundation for Adaptive Target-Guided Virtual Screens (ATG-VSs), which allow users to intelligently select and prioritize chemically diverse subspaces relevant to a given target, thereby optimizing computational resources focusing on the most promising regions of chemical space. Furthermore, an optional active learning component can be layered on top of ATG-VSs to iteratively refine the search using data-driven feedback, dynamically selecting molecules predicted to yield the highest information gain or binding likelihood (Extended Data Fig. 3, 4, 5, 6, 7).

The AFVS module enables large-scale docking campaigns using over 1,500 distinct docking protocols. We define a docking protocol as a specific combination of a sampling algorithm (pose prediction) and a scoring function, allowing for a vast number of unique workflows assembled from more than 40 individual docking programs. AFVS supports a wide range of ligand and target types, including peptides, DNA, and RNA, and includes recent advances in machine learning-based docking such as DiffDock and TANKBind (12, 13), which have shown promise in speed and accuracy, especially for blind or flexible docking scenarios (Extended Data Table 4, 5, 6; Supplementary Table 1, 2; Supplementary Fig. 4).

To unify and simplify large-scale workflows, we developed AdaptiveFlow Unity (AFU), a new module that integrates the functionalities of AFLP and AFVS into a cohesive interface. AFU enables users to perform ligand preparation and docking within a single automated pipeline, dramatically reducing setup complexity and increasing accessibility for non-specialists. This streamlined design supports rapid iteration and facilitates integration with other tools in computational drug discovery and AI-driven design workflows (Supplementary Fig. 5, 6, 7, 8, 9, 10, and Methods: AdaptiveFlow Unity).

### Enamine REAL Space

Virtual screening performance scales significantly with library size, because larger screening sizes tend to yield both higher potencies and improved hit rates (i.e., the number of confirmed hits divided by the number of tested compounds) (7, 9–11). Until recently, the largest publicly available ready-to-dock libraries, including the 2018 version of the REAL Database (9) and the ZINC20 library (14), each contained approximately 1.5 billion molecules. To enable virtual screens of an order of magnitude larger scale, we prepared the 2022 version of the Enamine REAL Space into a ready-to-dock format, comprising 68.7 billion commercially available on-demand small molecules.

The current version of the Enamine REAL Space (2022q1–2, accessed November 15, 2022 (15)) contains 31,507,987,117 enumerated molecules (becoming 69 billion after ligand preparation, e.g. due to stereoisomer and tautomer enumeration). These compounds are synthetically accessible derivatives of 137,000 established building blocks, assembled using 167 internally validated one-pot synthetic protocols developed at Enamine (16–25). This combinatorial framework enables high-throughput compound generation with a reported synthetic success rate of 82%, based on 386,000 experimental reactions conducted in 2021. 99% of the compounds in the library conform to Lipinski’s Rule of Five (26). Structurally, it offers exceptional chemical diversity, comprising 1,098,811,629 unique Murcko scaffolds.

Following ligand preparation with AFLP, the number of ready-to-dock molecules expanded to 68.7 billion in the REAL Space. This transformation involved a comprehensive set of preprocessing steps, including stereoisomer and tautomer enumeration, desalting, neutralization, protonation state prediction, 3D conformer generation, and conversion into formats compatible with a broad range of docking programs. In addition, 28 physicochemical properties and cheminformatic descriptors were calculated for each compound (Extended Data Table 2). To organize this vast chemical space and enable efficient navigation, a subset of 18 key molecular properties was selected to construct an 18-dimensional matrix, in which each axis corresponds to a descriptor such as molecular weight, logP, hydrogen bond donor and acceptor counts, rotatable bond count, topological polar surface area (TPSA), aqueous solubility (logS), aromatic ring count, molecular refractivity (MR), formal charge, positive and negative charge counts, fraction of sp^3^-hybridized carbons (*fsp*^3^), number of chiral centers, halogen and sulfur atom counts, and stereoisomer count. Each dimension was discretized into multiple property intervals, resulting in a combinatorial matrix of 24.4 billion potential tranches (i.e., matrix elements). Molecules from the REAL Space occupy approximately 12 million of these tranches, allowing for highly granular, property-based selection of compound subsets. The 18 properties were chosen to balance chemical relevance for drug discovery with computational efficiency, as excessive dimensionality would lead to sparsity and reduce practical tractability. The binning schemes and occupancy statistics for each dimension are provided in Extended Data Fig. 3.

The REAL Space is not only vast, but also chemically rich and structurally diverse, particularly in its representation of three-dimensional and stereochemically complex molecules (14, 27). Notably, over 50.5 billion molecules (74%) contain at least one chiral center, contributing to what is often referred to as “nonflat” chemical space, a feature increasingly associated with favorable pharmacological properties (28, 29). Ligand–receptor binding is inherently three-dimensional, and the presence of multiple stereoisomers allows subtle shape complementarity to be exploited. In this context, differences in docking scores across stereoisomers can serve as a proxy for specific target engagement, aiding in the prioritization of high-affinity binders. The REAL Space also includes a substantial number of halogenated compounds, 20.2 billion in total, including 17.4 billion containing fluorine atoms, which are well suited for ^19^F-NMR–based binding assays that do not require isotopically labeled proteins. Despite its breadth, the 18-dimensional matrix remains sparsely populated, revealing large underexplored regions of chemical space. These unoccupied tranches present compelling opportunities for prospective synthesis campaigns to populate novel chemical spaces, enabling the development of molecules to selectively target previously undruggable regions of the proteome.

To facilitate exploration of this high-dimensional chemical library, we developed two interactive web interfaces (Extended Data Fig. 5 and 6), each built with different rendering frameworks to optimize performance across devices and browsers. These interfaces are a projection of the 18-dimensional matrix onto a two-dimensional matrix, allowing the users to dynamically visualize the properties. These interfaces present the 18-dimensional matrix as a dynamic two-dimensional projection, allowing users visualize the matrix. Each of these two axes of the two-dimensional projection is based on one of the 18 properties that define the full matrix, and the user can select the two properties. Although the visualization is limited to 2D projections, the range of the 18 properties (of the corresponding 18 dimensions) can be selected interactively independently of the visualization. Selected subsets can then be used directly for virtual screens via AFVS. The REAL Space is available in PDB, PDBQT, MOL2, and SDF formats, ensuring compatibility with most structure-based docking programs. Additionally, the library is distributed in SMILES, SELFIES, and Parquet formats to support cheminformatics, generative modeling, and large-scale analytics. Both the full library and user-defined subsets can be accessed via the AdaptiveFlow project homepage or through the AWS Open Data repository, enabling direct use in the cloud without the need for local storage.

### Adaptive Ultra-Large Virtual Screenings

Screening ultra-large ligand libraries containing billions of molecules is computationally intensive, especially when using classical physics-based docking programs. As the number of compounds in the library increases into the tens of billions, the cost of screening the library becomes prohibitively expensive.

Virtual screening is a process of searching a vast chemical space for the best candidate that matches the structural and biophysical properties of the target site in a given receptor. However, the fraction of compounds in an ultra-large screen that will optimally engage a specific site in the receptor is often very small and share specific molecular characteristics. To be cost- and time-effective, it is desirable to focus the screen on the optimal chemical space that is likely to contain hits for a given target. To rapidly narrow the search space for a given target, we introduce the Adaptive Target-Guided Virtual Screening (ATG-VS) technique. This approach consists of the following steps:

1. **Library Decomposition**. On average, a tranche in the current library has approximately 5600 molecules, and for each tranche up to ten representative ligands are selected for a pilot screen. The number of representatives per tranche (1–10) can be selected by the user. Since there are approximately 12 million tranches (that contain molecules from the REAL Space), if one representative per tranche is selected, the total number of molecules screened in the prescreen would be approximately 12 million for the entire library. If more than one representative molecule, per tranche, is chosen by the user then these additional molecules are automatically selected to be diverse from each other within the given tranche.
2. **Screening Representative Molecules (ATG Prescreen)**. Instead of screening the entire library, a prescreen of only the selected representative ligands is performed. In addition, instead of screening representative ligands from all the 12 million tranches, users can apply an optional dynamic filtering procedure that corresponds to the 18-dimensional properties of the ligands. Here, users can manually apply criteria that the molecules/tranches need to satisfy, e.g. based on the properties of the targeted site, which can significantly reduce the number of dockings required in the prescreen. After the prescreen, the sub-space (tranches) of the ligand library that have the best docking scores among the representative ligands are selected for the next screening stage.
3. **Selection of Active Tranches**. AdaptiveFlow automatically selects the most promising tranches based on docking scores of the ATG Prescreen. The total number of ligands to be screened in the next stage can be specified by the user.
4. **Active Cell Screening (ATG Primary Screen)**. In the next stage, the tranches that were selected after the prescreen are fully screened (e.g., in Fig. 3, the three tranches that contain the best ligands are marked purple). To further refine this selection, a machine learning classification model (see **Methods: Machine Learning Classifier for Tranche Prioritization**) is employed to identify the most promising molecules within each tranche. The model is trained on Morgan fingerprints (size 1024) of the prescreened compounds, using the docking scores obtained during the prescreen as labels. It applies a threshold-based classification approach, where the 75th percentile of docking scores serves as the decision boundary. Only molecules classified as high-confidence binders are prioritized for full docking, ensuring that computational resources are allocated efficiently to the most promising candidates.

**Fig. 2.**
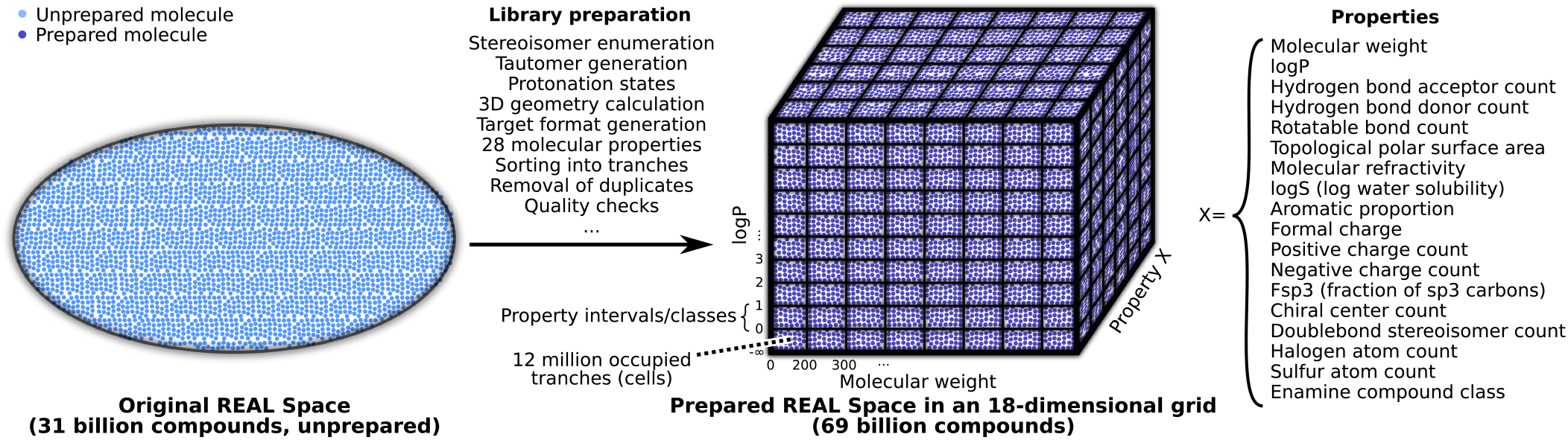
AdaptiveFlow version of the Enamine REAL Space. The initial version of the REAL Space (enumerated release, 2022q1–2) comprised approximately 31 billion molecules represented as SMILES strings in an unprocessed format, e.g. without full stereoisomer enumeration or structural standardization. A selected subset of 18 of these properties was used to partition the library into an 18-dimensional property matrix, where each dimension was discretized into multiple intervals. In this figure, the high-dimensional matrix is illustrated as a 3D schematic for clarity, with three representative properties each split into 6–10 intervals, resulting in 8 · 6 · 6 = 288 tranches. Each box represents a tranche that contains ligands sharing the corresponding property ranges. This structure enables flexible selection of compound subsets and serves as the foundation for adaptive, target-guided virtual screens (ATG-VSs) that are introduced in this work.

**Fig. 3.**
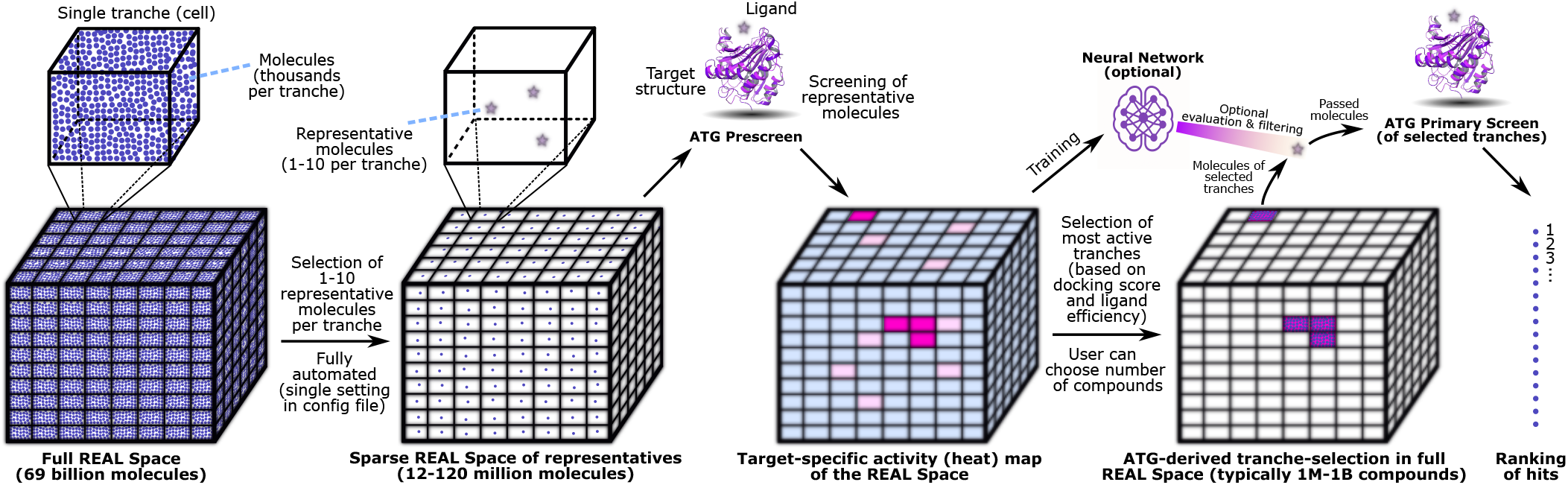
Conceptual overview of Adaptive Target-Guided Virtual Screens (ATG-VSs). The REAL Space, divided into an 18-dimensional grid, populates over 12 million individual tranches (cells) represented in the figure schematically as a 3-dimensional matrix and each of these tranches has on average 5600 molecules. During the first step of ATG-VSs, the AFVS module of AdaptiveFlow selects a few (1–10) representative molecules in each tranche and docks these molecules to the target receptor. This screening mode (and stage) is referred to as a prescreen and allows AdaptiveFlow to detect the tranches of the library that contain molecules of high potential binding affinity to the given target receptor/binding site. AdaptiveFlow then selects these tranches and screens them fully against the target receptor. In addition, an optional machine learning (ML) model can be trained on the ATG Prescreen results, which can evaluate the molecules of the selected tranches during the ATG Primary Screen.

ATG-VSs significantly reduce the dimension of the chemical space to be screened, thereby reducing the required computation time. AFVS can also prepare new and/or proprietary ligand libraries with ATG-VS support, which requires that the prepared ligand libraries are structured into a multidimensional tranche matrix with pre-assigned representative molecules for each tranche. An optional third-stage screen can be performed to rescore the best hits with higher accuracy, such as by including receptor flexibility. AdaptiveFlow can carry out all three stages in a fully automated workflow.

AdaptiveFlow is a highly flexible platform that supports funnel-type screening strategies, either as a standalone method or in combination with ATG-VS. Users can perform iterative filtering by screening a subset of the library (or the full library, if computationally feasible) to generate an initial ranking, then re-screening the top candidates with increased exhaustiveness or different scoring functions. This allows for an arbitrary number of screening stages to be deployed. Additionally, users can manually select library subsets based on molecular properties or generate smaller custom libraries via the AFLP module to facilitate multi-stage workflows.

To investigate the effectiveness of the various parameters including the number of representative molecules from each tranche, and the docking exhaustiveness (which is a measure of how extensively the ligands’ conformational space is sampled), we carried out multiple ATG prescreens on an array of targets. We examined the following scenarios, i) one representative molecule per tranche and exhaustiveness of 1, ii) 10 representative molecules and exhaustiveness of 1, and iii) 1 representative molecule with exhaustiveness of 10. The higher the exhaustiveness parameter the more thoroughly the conformational space is searched, but also the more computation time is needed. The results of these trials can be seen in Extended Data Fig. 8. As shown in the figure, the results (based on the distribution of the top docking scores, and the ranking of the tranches within each property category) are relatively similar for all three scenarios. Therefore, ATG prescreens with exhaustiveness 1 and one representative molecule per tranche are expected to be sufficient for many applications. In this case, an ATG prescreen involves docking 12 million molecules to explore the 69 billion compound library, which is over 5000 times less computational effort than screening all molecules of the library. If a second stage screen with higher accuracy is added in the context of multistaging, it can be beneficial to increase the exhaustiveness (9).

To systematically evaluate the performance of Adaptive Target-Guided Virtual Screens (ATG-VSs) relative to standard ultra-large virtual screens (ULVSs), we conducted benchmarking experiments across ten diverse protein targets, including kinases, phosphatases, GPCRs, and protein–protein interaction interfaces (Extended Data Table 7). These evaluations were performed in two settings: (1) test-based benchmarks using a 5-million-compound subset of chemical space (Fig. 4), and (2) large-scale production runs using the full 69-billion–compound Enamine REAL Space (Extended Data Figs. 9, 10). The production-scale benchmarks compared standard ULVSs to ATG-VS without the optional active learning component, while the subset-based studies also assessed ATG-VS with active learning enabled. Across these targets, we evaluated performance under the following scenarios:

i. Comparison to standard ULVSs (production-scale benchmarks): We evaluated performance at full production scale (Extended Data Figs. 9, 10) by comparing the top 50 docking scores obtained from two approaches: (1) a standard ULVS involving 100 million randomly selected compounds from the full Enamine REAL Space, and (2) the ATG-VS method without active learning. For ATG-VS, we conducted a prescreen of 12 million compounds, comprising 10 representative ligands per tranche across the entire library, followed by a primary screen of up to 1 million compounds. To assess the impact of screening depth, we systematically varied the number of compounds in the primary screen (10K, 100K, and 1M) across ten targets. Active learning was not applied in this setting to isolate and evaluate the impact of tranche prioritization alone; however, based on the test-based benchmarks we expect that incorporating active learning could yield further improvements in efficiency and enrichment. We found that ATG-VS achieves comparable or superior top docking scores to standard ULVSs while requiring dramatically fewer docking evaluations, highlighting its effectiveness and scalability in production-scale virtual screening campaigns.
ii. Impact of machine learning (test–based benchmarks): To further evaluate the benefit of incorporating an ML classifier into the ATG workflow, we conducted experiments using a sub-sampled version of the REAL Space comprising 5 million compounds. We compared three approaches: (1) a random screen of 1 million compounds (“1M Random” in Fig. 4); (2) the ATG-VS method, which involved decomposing the library based on 18 molecular properties, performing a prescreen, and selecting compounds for a primary screen, with a total of 100,000 docking evaluations across both stages (“100K ATG”); and (3) an enhanced ATG-VS approach in which a machine learning model, trained on docking scores and Morgan fingerprints from the prescreen, was used to prioritize compounds for the primary screen (“100K ATG+AL”). It should be noted that in the case of ATG and ATG+AL scenarios, the total number of docking evaluations, including the prescreen, was 100K, compared to a million in the random screen. As shown in Fig. 4, the ML-augmented ATG-VS consistently improved the enrichment of high-affinity virtual hits relative to the standard ATG-VS. Though both versions of the ATG screen sampled only 100K molecules, they outperformed the random screen sampling of 1 million molecules.

**Fig. 4.**
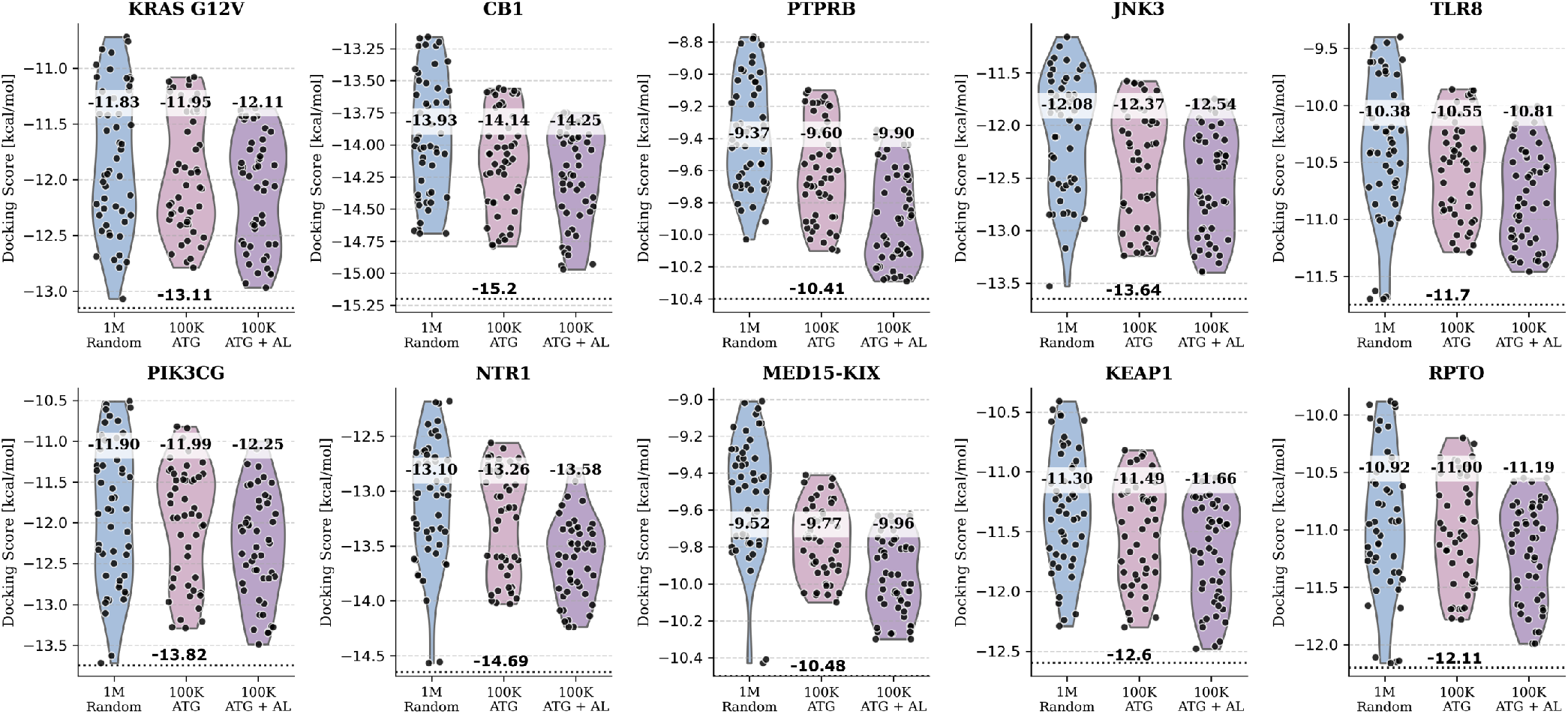
Benchmarks Involving Adaptive Target-Guided (ATG) Primary Virtual Screens and Standard ULVSs. Violin plots of the docking scores (the more negative, the better) of the top 50 virtual screening hits obtained using three approaches: (i) standard ultra-large virtual screening (ULVS) of 1 million random compounds (blue), (ii) adaptive target-guided (ATG) screening with 100,000 molecules (rose), and (iii) ATG screening augmented with an additional active learning step (purple). Black dots represent individual data points, and the numbers above the violin plots indicate the mean docking scores of the top 50 hits. The horizontal dashed lines indicate the best docking score obtained from a brute-force screen of the entire library, serving as a reference for the absolute best possible hit (5-million-compound subset). For most target proteins, ATG primary screens (100,000 molecules) with or without active learning performed comparably to standard ULVS (1 million random compounds), achieving a 10-fold reduction in the number of compounds screened while maintaining similar docking performance.

While ATG-based screens consistently outperformed random screening, the extent of improvement varied depending on the biophysical properties of the target. Proteins with complex, feature-rich binding pockets, particularly those combining extensive hydrophobic surfaces with well-defined polar subpockets, derived the greatest benefit from ATG-based prioritization and ML enhancement. Taken together, these findings underscore the scalability, efficiency, and adaptability of ATG-VS. By intelligently narrowing the search space and optionally integrating machine learning for compound prioritization, AdaptiveFlow enables high-performance virtual screening at the scale of tens of billions of molecules.

To evaluate the sensitivity of our binning strategy to different sampling algorithms and scoring functions, we repeated the benchmarks using Smina (employing the Vinardo scoring function) and GWOVina (incorporating flexible side-chain docking). The physicochemical property-based binning remains effective independent of the docking protocol used. As shown in Supplementary Fig. 11, the ATG-VS approach consistently identified high-scoring compounds with speedups and enrichment factors comparable to those observed with QuickVina 2. Specifically, the average predicted binding affinity of the top 50 molecules improved relative to standard random screens, while the computational costs remained dramatically lower. This demonstrates that the descriptor-based binning correlates well with binding potential across docking protocols of varying sophistication.

### Machine Learning and Cloud Support

AdaptiveFlow introduces expanded capabilities designed to support artificial intelligence (AI) and machine learning (ML) workflows across molecular biology and drug discovery. In addition to integrating state-of-the-art ML-based docking programs, including EquiBind (30), DiffDock (13), and TANKBind (12), the platform offers several features aimed at facilitating the development and application of ML models. The AFLP module computes a wide range of physicochemical properties commonly used as features or optimization targets in ML pipelines, including the Quantitative Estimate of Druglikeness (QED) and octanol–water partition coefficient (logP) (32). AFLP also supports the generation of SELFIES (Self-Referencing Embedded Strings) molecular representations (33, 34), which are widely adopted in generative modeling due to their syntactic robustness and ability to encode valid molecules across arbitrary token sequences (35) (Extended Data Table 2, 3, 3, 4, 5, 6, 7; Supplementary Table 1, 2). Additionally, AFVS introduces numerous deep learning-based docking approaches by modularly assembling scoring functions and pose prediction tools into comprehensive docking workflows. To further streamline ML-driven workflows the AFU module provides a fully integrated pipeline for ligand preparation and docking. This unified interface simplifies data generation for model training and enables seamless incorporation of docking scores as objective functions in generative or optimization-based ML models (36). With support for approximately 1,500 docking protocols accessible through a single command-line interface, AFU eliminates the need for users to individually configure and learn multiple docking programs, thus significantly lowering the barrier to entry for ML and drug discovery communities alike. Together, these features position AdaptiveFlow as a versatile platform not only for applying ML methods to virtual screening but also for developing new ML architectures and pipelines for structure-based drug discovery (Extended Data Fig. 11, 12).

In parallel with its support for machine learning, AdaptiveFlow is optimized for efficient deployment in cloud computing environments. The platform runs natively on the popular workload manager Slurm (37), widely supported across providers like Google Cloud and Microsoft Azure, and AWS Batch (38). Users provide a SLURM submission template (e.g., template1.slurm.sh) in the configuration. AdaptiveFlow populates this template with the necessary variables (job arrays, partitions, memory) at runtime. This decouples the workflow logic from the specific cluster configuration. Users can adapt the submission script to their local requirements (e.g., specific headers, modules, or partition names) without modifying the core Python codebase. Further, the size of computing jobs (both size in cores and length of time) can be configured based on site-specific cluster configurations.

In this study, we demonstrate near-linear scalability on AWS using over 5.6 million virtual CPUs, setting a new global benchmark for parallelization in cloud-based drug discovery. This level of scalability enables the completion of ultra-large virtual screens involving billions of compounds within hours, timelines that would otherwise require weeks or months on traditional on-premise compute clusters. By combining cloud-native performance with ML-native features, AdaptiveFlow serves as a high-performance and accessible foundation for next-generation AI-driven drug discovery.

### Experimental validation of AdaptiveFlow

To experimentally validate the capabilities and performance of AdaptiveFlow, we selected two therapeutically significant target proteins, ferroptosis suppressor protein 1 (FSP1) and poly(ADP-ribose) polymerase 1 (PARP1). FSP1 is a recently characterized oxidoreductase that plays a central role in protecting cells from ferroptosis, an iron-dependent form of regulated cell death triggered by membrane lipid peroxidation (39–42). FSP1 maintains pools of reduced coenzyme Q_10_ and vitamin K hydroquinone using NAD(P)H, operating independently of GPX4 to suppress oxidative membrane damage. Inhibiting FSP1 disrupts this defense, sensitizing cells, especially redox-imbalanced cancer cells (43), to ferroptosis. Despite its importance, only a few FSP1 inhibitors have been described and their binding to FSP1 has been structurally characterized (2, 44–46), making it an ideal target for exploring novel inhibitor space using AdaptiveFlow. We also targeted PARP1, a well-established target involved in DNA single-strand break repair, for benchmarking AdaptiveFlow on a mechanistically distinct, clinically validated enzyme class.

#### Structural Studies of FSP1

To streamline hit discovery and structural studies, we employed ancestral sequence reconstruction, a method known to yield highly stable proteins while preserving functional and structural fidelity to modern homologs (47, 48). Using this approach, we reconstructed the tetrapod ancestor of human FSP1, which shares 76% sequence identity and retains all active-site residues (Supplementary Fig. 12, 13). The ancestral FSP1 variant expressed efficiently in *E. coli* (~ 100 mg/L culture), bound FAD, exhibited high thermostability (*T*_m_ = 62 °C; Supplementary Fig. 13B), and retained full enzymatic activity (Supplementary Table 3; Supplementary Fig. 14 and 15). Crucially, it yielded high-quality, well-diffracting crystals (Supplementary Table 4), enabling detailed structural analyses.

We determined high-resolution crystal structures of FSP1 bound to FAD, to FAD and NAD^+^, and in complex with FAD, NAD^+^ and coenzyme Q_1_ (Fig. 5; Supplementary Fig. 16a; Supplementary Table 4). The overall structure of the ancestral protein is very similar to that of the human enzyme (Supplementary Fig. 16b). NAD^+^ triggers a 2 Å shift of the flavin ring and its binding mode is fully consistent with a hydride transfer mechanism for flavin reduction as the nicotinamide C4 atom is located right in front of the isoalloxazine N5 (Fig. 5B). Coenzyme Q_1_ binds between two aromatic side chains of Phe354 and Y290, with its quinone moiety at the edge of the pyrimidine ring of the flavin in a canyon leading from the protein surface (Fig. 5C-D). Specifically, quinone oxygen engages the N(3)H group of the flavin through a hydrogen bond. This geometry and the proximity between the two molecules can enable the facile electron transfer from the flavin to coenzyme Q. Comparison between the FSP1-FAD-NAD^+^ and FSP1-FAD-NAD^+^-coenzyme Q_1_ complexes shows that NAD^+^ binding is unaffected by the presence of coenzyme Q. Thus, it is demonstrated that coenzyme Q and NAD(P)^+^ can simultaneously bind to the protein and this structural attribute perfectly aligns with the ternary-complex mechanism evidenced by the kinetics data (Supplementary Fig. 15). The structure also suggests that there is ample space for the oxygen to access the flavin N5 locus where its reaction with the flavin is known to take place. This feature explains why FSP1 can also operate as a NAD(P)H oxidase (49). This detailed experimental visualization of the NAD(P)H and quinone binding modes and their interactions provided the foundation for inhibitor discovery through in silico screenings targeting the substrate binding sites.

**Fig. 5.**
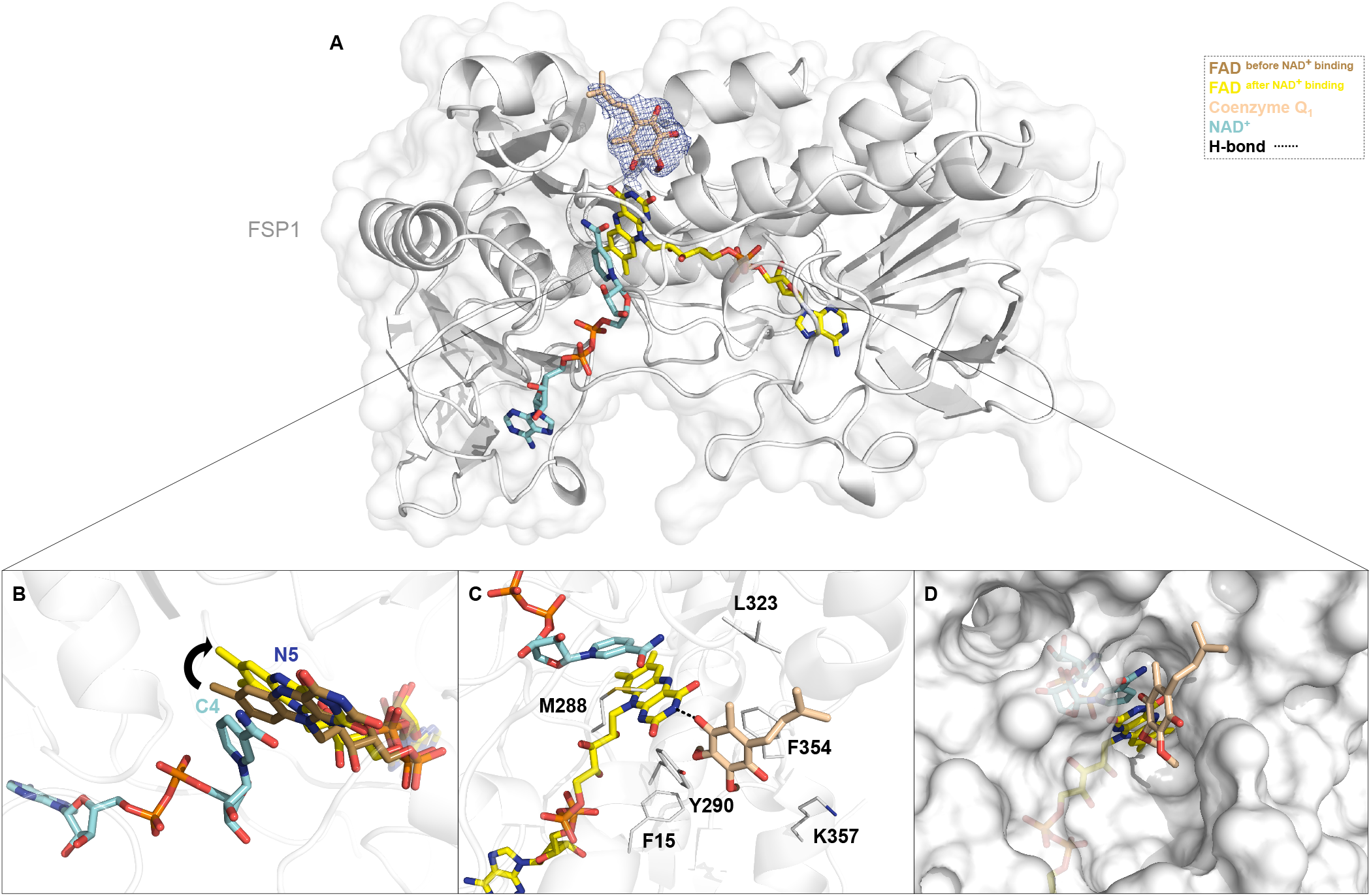
Co-Crystal Structures of FSP1 in Complex with its Cofactors and Substrate. We determined three crystal structures of FSP1: (i) bound to FAD alone, (ii) bound to FAD and NAD^+^, and (iii) bound to FAD, NAD^+^, and coenzyme Q_1_. (A) Overall structure of the fully assembled FSP1 complex with FAD (yellow carbons), NAD^+^ (cyan), and coenzyme Q_1_ (beige). The polder omit map for coenzyme Q_1_ is contoured at the 3.0 σ level. (B) Superposition of structures (i) and (ii) reveals that NAD^+^ binding induces a rotation of the FAD isoalloxazine ring. The FAD conformation prior to NAD^+^ binding is shown in brown, and the conformation after NAD^+^ binding is shown in yellow. No evidence for 6-hydroxyflavin formation was observed, in contrast to prior reports for human FSP1 (50, 51) (see also Supplementary Fig. 16A). (C) In the ternary complex, coenzyme Q_1_ interacts with Tyr290 and Phe354 via edge-to-edge aromatic stacking and forms a hydrogen bond (dashed black line) with the flavin’s pyrimidine ring. (D) NAD^+^ and coenzyme Q_1_ occupy distinct but converging channels that meet at the FAD active site, illustrating the architecture of the substrate-binding cavity.

#### Targeting FSP1 with AdaptiveFlow

To assess the effectiveness of AdaptiveFlow, we screened FSP1 inhibitors targeting its coenzyme Q site using AdaptiveFlow. Our high-resolution crystal structure of FSP1 in complex with FAD and NAD^+^ (PDB code: 9IFT) was used as receptor. Maestro from Schrödinger (52) was applied to prepare the receptor’s structure for screening, by removing solvent molecules, assigning missing hydrogen bonds, correcting protonation states at pH 7.4, and assigning correct bond orders. AutoDockTools (v.1.5.7) (53) was utilized to convert the prepared structure from the PDB format to the PDBQT format. The coenzyme Q site of FSP1 (with FAD and NAD^+^ bound to the corresponding sites) was targeted. In the ATG Prescreen, 1 representative compound was used per tranche (resulting in 12 million dockings), and based on these results 10 million compounds were chosen for the primary screen using the ATG-VS method (without ML feature). Basic filters were applied to the top hit candidates, such as logP less than 5, LogS greater than −5, Topological Polar Surface Area (TPSA) smaller than 140 Å^2^, number of stereo centers not greater than 3, and compounds with problematic functional groups and predicted toxicity (mutagenic compounds, compounds with reproductive effects, tumorigenic compounds) were excluded. In total, 42 compounds from 37 different clusters with zero alerts suggested by SwissADME (54) were ordered from Enamine. 33 compounds were synthesized and experimentally tested.

#### Experimental Validation: Inhibitors extend from the coenzyme Q site to the NAD site

We assessed the potential binding and inhibitory activities of the synthesized 33 virtual hits from AdaptiveFlow, using thermal shift and NADH-depletion assays. Two of the hits elicited stabilizing shifts of at least 1.5 °C in the protein melting temperature and inhibited FSP1 oxidoreductase activity by at least 50% at a 10 µM concentration. These two compounds were found to potently inhibit coenzyme Q-reduction activity (Table 1; Supplementary Fig. 17). Their *K*_*i*_ values are comparable to, or better than, those reported for previously described FSP1 inhibitors (0.283 µM for afi-FSP1-1 and 0.777 µM for afi-FSP1-2; Supplementary Table 5; Supplementary Fig. 18, 19) (2, 44, 45). Critically, a cellular thermal shift assay (CETSA; (55)) performed on FSP1-expressing HEK293 cells shows that afi-FSP1-1 and afi-FSP1-2 effectively engage human FSP1 in cells, as evidenced by a thermal shift of 2.8 °C and 3.1 °C, respectively (Supplementary Fig. 20A-J). The activity of afi-FSP1-1 was tested against a T-cell acute lymphoblastic leukemia cell line (Jurkat) that expresses FSP1 (Supplementary Fig. 20M) and is known to undergo ferroptosis (56). Cells proved to be sensitive to the combination of afi-FSP1-1 with 0.5 *µ*M erastin, a ferroptosis inducer. The activity of afi-FSP1-1 matched that shown by the widely used iFSP1(2). When used as a single agent, afi-FSP1-1 showed no cell toxicity at concentrations up to 10 *µ*M. The pro-ferroptotic effect induced by 25 µM inhibitor was completely rescued by liproxstatin-1, a ferroptosis inhibitor. (Supplementary Fig. 20N-P). We determined the crystal structure of the most potent compound (afi-FSP1-1) bound to FSP1-FAD-NAD^+^ which reveals that the inhibitor occupies the coenzyme Q binding site in a mode consistent with the predicted docking pose (Fig. 6A-B; Supplementary Fig. 16a). With its elongated conformation, afi-FSP1-1 extensively interacts with the protein by contacting several aromatic side chains, including Tyr290 and Phe354, residues that sandwich the quinone ring of coenzyme Q. Tyr290 is involved in a *π*-stacking interaction with the benzene ring of the isochroman moiety, whereas Phe354 and Phe15 contribute to the binding via hydrophobic interactions with the piperidine and a *π*-stacking interaction with the triazole group. The central carbonyl oxygen of afi-FSP1-1 forms an H-bond interaction with the N(3)H group atom of the flavin akin to the interaction established by the carboxyl oxygen of the coenzyme Q’s ubiquinone head (Fig. 6B and 5C). Furthermore, the inhibitor’s central carbonyl oxygen consistently interacts with the backbone amine group (NH) of Leu323 via a water bridge (Supplementary Fig. 21). Despite its proximity with the inhibitor, the binding conformation of NAD^+^ is identical to that observed both in the absence of a ligand and in the complex with coenzyme Q_1_. Similar features characterize the binding compound afi-FSP1-2 whose bulkier structure enables even more extended interaction with the protein (Supplementary Fig. 21, 22). Expanding on these results, we conducted a structure activity relationship study. The three-dimensional structure of the FSP1/afi-FSP1-1 complex was used as input for an additional round of in silico screening. This procedure returned 20 analogues of compound afi-FSP1-1 that were experimentally analyzed (Supplementary Fig. 23). Rewardingly, four additional molecules (afi-FSP1-3-6) displayed inhibitory activities with *K*_*i*_ values in two-digit nanomolar for achiral molecules to low micromolar range for molecules tested as either a racemic mixture or a diastereomeric mixture (Supplementary Fig. 17C-F; Supplementary Fig. 20). Notably, these compounds share an aromatic-amide-saturated heterocycle core as displayed also by afi-FSP1-1 and 2 (Table 1). Consistent with this common feature, the crystal structures of afi-FSP1-3 and afi-FSP1-4 bound to FSP1-FAD-NAD^+^ show that the ligands occupy the coenzyme Q site with their amide carbonyl oxygen engaged by the flavin N(3)H group via a hydrogen bond and further interacting through a water molecule with the backbone amine group of L323 in FSP1 as observed for afi-FSP1-1 (Fig. 6B-D; Supplementary Fig. 21, 22). Although compound afi-FSP1-4 replaces the dimethyl-tetrahydropyran portion of the isochroman group of afi-FSP1-1 with a methylthiomethyl-triazole, the two inhibitors bind in a very similar way with their common phenyl-triazole-piperidine moiety interacting with Phe15 and Phe354. Conversely, the binding of afi-FSP1-3 is more distinct as it adopts a less extended, U-shaped conformation laying on Tyr290. These binding modes are validated by mutagenesis experiments (Supplementary Table 6, 7; Supplementary Fig. 21, 22) and correlate with the measured *K*_*i*_ values. Compounds afi-FSP1-1 and afi-FSP1-4 exhibit practically identical *K*_*i*_s, whereas compound afi-FSP1-3 displays a twofold decrease in potency. Nonetheless, Absorption, Distribution, Metabolism, and Excretion (ADME) assays indicated that afi-FSP-3 is the most promising candidate for further medicinal chemistry optimization (see Supplementary Information Section H for details). Besides being the best inhibitors, compounds afi-FSP1-1, afi-FSP1-2, and afi-FSP1-3 exhibit a 10-12-fold reduced efficacy against the NADH oxidase activity of FSP1, while compound afi-FSP1-4 exhibited no efficacy at all (Supplementary Fig. 25; Supplementary Table 8). The weaker inhibition reflects the retention of NAD(P)H binding in the inhibited protein as shown by crystal structures and indicates that reactivity with oxygen is only mildly or not affected by inhibitor binding in the coenzyme Q site. We posit that preferentially inhibiting the quinone-reducing activity while sparing the reactivity with oxygen, might be pharmacologically beneficial. Inhibitors of this type may convert the enzyme from being anti-ferroptotic (coenzyme Q reducing activity) to being pro-ferroptotic (uninhibited lipid-damaging ROS production). Future work is needed to explore and validate this concept in FSP1 pharmacology.

**Table 1.** Competitive *K*_*i*_ and Binding *K*_*d*_ Values of the Most Potent FSP1 Inhibitors. *K*_*i*_ values were determined by monitoring NADH oxidation at 340 nm (pH 7.2) using variable NADH concentrations (5–200 *µ*M), 100 *µ*M coenzyme Q_1_, and 0.1–0.2 *µ*M FSP1 (Supplementary Fig 17). *K*_*d*_ values were derived from independent binding experiments based on surface plasmon resonance (SPR, Supplementary Fig. 19). The experiments were performed using the racemic mixture of compound afi-FSP1-5 and the diastereomeric mixture of compound afi-FSP1-6. Inhibitors’ concentrations ranged from micromolar to low nanomolar. Supplementary Table 5 compares the *K*_*i*_ values of the afi-FSP1 compounds with those of known FSP1 inhibitors whereas the IC_50_ values measured using the indirect resazurin assay, commonly used in previous literature (2, 44, 45), are listed in supplementary Table 9.

| Compound | Structure | $K_i$ ($\mu\text{M}$) | $K_d$ ($\mu\text{M}$) |
| --- | --- | --- | --- |
| afi-FSP1-1 | | $0.283^b \pm 0.05$ | 0.098 |
| afi-FSP1-2 | | $0.777 \pm 0.22$ | 0.266 |
| afi-FSP1-3 | | $0.520 \pm 0.11$ | 0.184 |
| afi-FSP1-4 | | $0.284 \pm 0.07$ | 0.198 |
| afi-FSP1-5 | | $1.00 \pm 0.32$ | n.d. |
| afi-FSP1-6 | | $3.94 \pm 0.71$ | n.d. |

**Fig. 6.**
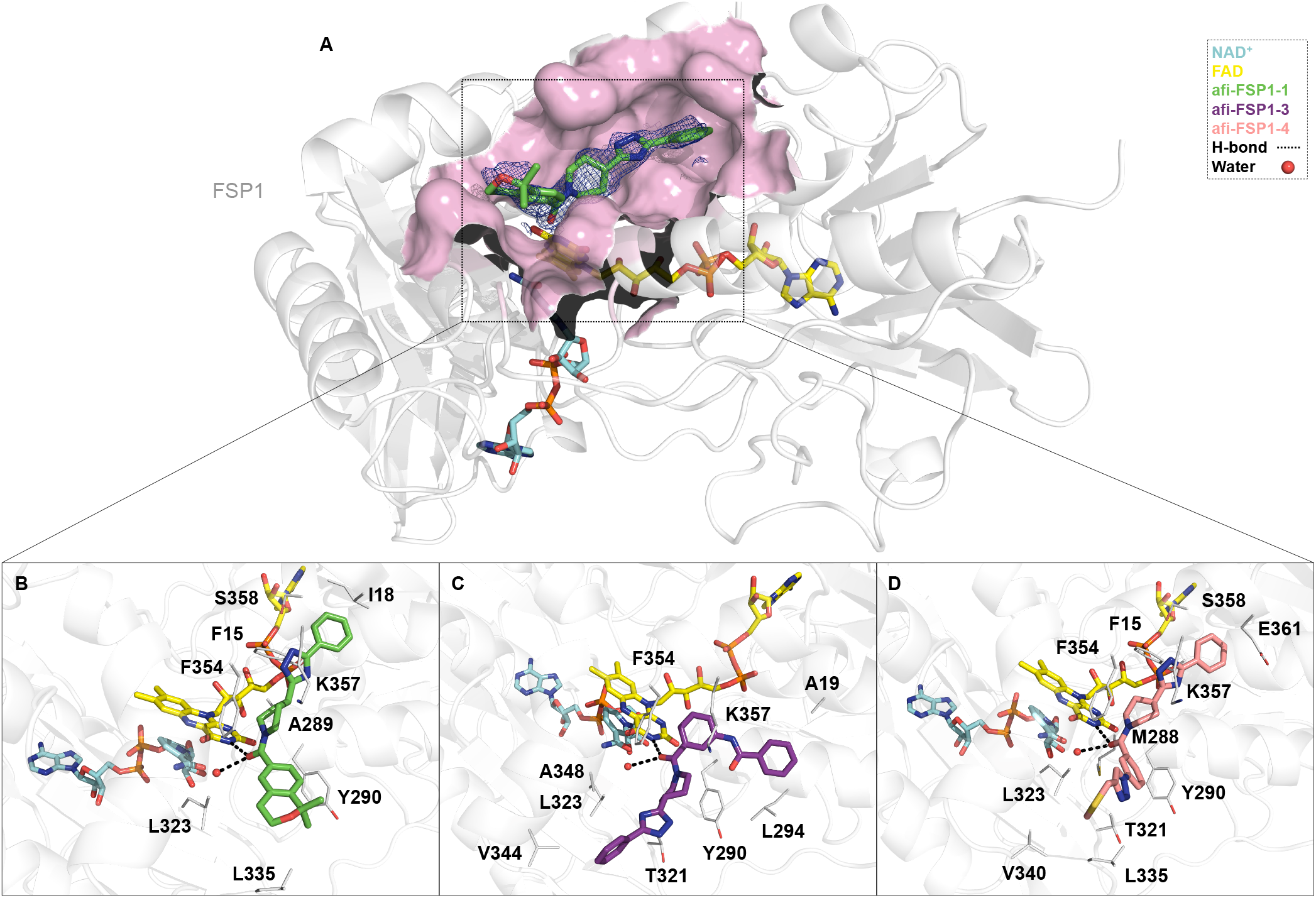
Co-Crystal Structures of Inhibitors binding to FSP1. (A) Overall structure of FSP1 in complex with inhibitor afi-FSP1-1. The polder omit map for the inhibitor is contoured at the 3.0 σ level. (B-D) Close-up views of the binding site with afi-FSP1-1 (B, green), afi-FSP1-3 (C, violet), and afi-FSP1-4 (D, pink) bound in the region normally occupied by coenzyme Q_1_, adjacent to FAD (yellow) and NAD^+^ (cyan). All inhibitors displace coenzyme Q_1_ and interact directly with the catalytic cavity.

#### Applying AdaptiveFlow to PARP1

To further validate the effectiveness of the AdaptiveFlow platform, we selected human Poly(ADP-ribose) polymerase 1 (PARP1) as a benchmark target due to its well-characterized structure, critical role in DNA repair, and relevance in cancer therapy. Following the ATG-prescreen and a primary screen of 100 million molecules, 160 promising candidates were synthesized for further experimental verification, as detailed in the Methods section. A two-concentration high-throughput enzymatic screen identified seven PARP1 inhibitors, four of which (iParp1, iParp2, iParp3, and iParp4) showed sub-250 nM IC50 values and were selected for further characterization. Protein NMR confirmed direct binding to the active site of PARP1, with chemical shift perturbations supporting the predicted docking poses. X-ray crystallography revealed the 2.05 Å structure of the PARP1 catalytic domain bound to a hydrolyzed form of iParp1, validating its binding mode and key interactions. Clonogenic assays in BRCA1-deficient TNBC cells demonstrated selective cytotoxicity for iParp1 and iParp2, though reduced cellular activity, especially for iParp1, was attributed to poor membrane permeability due to hydrolysis. Remarkably, AdaptiveFlow identified PARP1 inhibitors with submicromolar to low nanomolar potency, including iParp1, which exhibited an IC_50_ of 8.8 nM, comparable to the FDA-approved drug Olaparib. This result is particularly notable given that our hits were derived from a purely virtual screen without any prior knowledge of known inhibitors or medicinal chemistry optimization of the hit. Structural similarity analyses confirmed that several hits shared key scaffolds with known PARP1 inhibitors, while others showed different scaffold variations. Details can be found in the Supplementary Information Section I. Overall, this validation exemplifies AdaptiveFlow’s ability to efficiently and accurately discover drug-like molecules with clinically relevant potency, streamlining early-stage drug discovery. See Supplementary Information; Supplementary Fig. 27, 29, 30, 31, 33, 32; Supplementary Table 13.

## Discussion

The accessible chemical space for drug discovery is expanding at an unprecedented pace, with the number of purchasable, on-demand molecules growing from approximately 2 billion in 2020 to trillions of molecules. Yet, even this remarkable growth represents only a minuscule fraction of the estimated 1060 drug-like molecules that populate theoretical chemical space (5). Identifying high-quality hits within this vast and ever-expanding landscape remains a central challenge in early-stage drug discovery. To address this, we curated the largest ready-to-dock library to date, the Enamine REAL Space, comprising 69 billion commercially accessible small molecules, fully preprocessed and made available in multiple formats for structure-based virtual screening.

While ultra-large libraries increase the likelihood of identifying potent binders and enhance true-hit rates (7, 9–11, 57, 58), exhaustively screening billions of compound impose substantial computational costs, even with today’s cloud-based computing capabilities. AdaptiveFlow addresses this challenge through Adaptive Target-Guided Virtual Screening (ATG-VS), a highly efficient prescreening strategy that narrows docking efforts to target-specific, chemically diverse subspaces enriched for high-affinity candidates. When applied to the 69-billion-compound REAL Space, ATG-VS reduces screening costs by up to 1,000-fold compared to exhaustive searches, while preserving strong enrichment of top hits. Additionally, AdaptiveFlow incorporates deep learning–based docking and GPU acceleration, achieving a further 10-to 100-fold increase in throughput (59, 60). To experimentally validate the practical utility of AdaptiveFlow, we focused on two biologically and clinically relevant targets. First, we targeted ferroptosis suppressor protein 1 (FSP1), a redox enzyme involved in lipid peroxide detoxification and resistance to ferroptosis, and identified a series of nanomolar inhibitors. Structural characterization of FSP1 in complex with cofactors and ligands revealed that both NAD(P)H and coenzyme Q can simultaneously bind in catalytically competent orientations, providing mechanistic insights into its function. The identified hits were validated through crystallography and biochemical assays. We also applied AdaptiveFlow to poly(ADP-ribose) polymerase 1 (PARP1), a clinically validated oncology target central to DNA damage repair. Our ultra-large screen yielded seven inhibitors, including iParp1, which displayed an IC_50_ comparable to that of the FDA-approved drug olaparib (61). These results underscore the power of AdaptiveFlow in rapidly identifying potent hits across diverse protein classes and demonstrate the advantage of adaptive screening over brute-force enumeration. Notably, while our previous identification of nanomolar KEAP1–NRF2 inhibitors required an unbiased screen of 1.3 billion molecules, the ATG-guided screen for FSP1 achieved comparable success with just 22 million docking instances. Although the effectiveness of chemical space focusing is target-dependent, our benchmarking and experimental studies highlight the potential of ATG-VS to tailor the search space to specific binding pockets, significantly enhancing screening efficiency. As chemical space continues to expand, focusing on target-specific subspaces will no longer be optional, but essential.

With the infrastructure now in place, adding molecules to the library and directing them to their respective tranches is a computational process that scales linearly. As new tranches, beyond the 12 million currently populated are filled, only representatives from these newly populated tranches need to be incorporated into the prescreen set. The open-source nature of AdaptiveFlow enables seamless adaptation, extension, and development of the platform by both academic and industry users. With the increasing availability of high-resolution structural data, enabled by techniques such as cryo-EM and de novo structure prediction, combined with site-specific functional insights from gene editing technologies, in silico screening is poised to deliver chemical tools capable of modulating precise biomolecular functions. As modalities like molecular glues and targeted protein degraders mature, the ability to identify small molecules that bind specific protein surfaces with functional consequences is becoming increasingly valuable. AdaptiveFlow’s modular architecture supports over 1,500 docking protocols, encompassing combinations of scoring functions, sampling algorithms, and ML-based pose predictors such as DiffDock and TANKBind. The platform also natively supports SELFIES representations, physicochemical property calculations, and active learning–driven workflows, making it fully compatible with generative models and reinforcement learning pipelines (62). As generative models continue to advance, platforms like AdaptiveFlow will be essential for rapidly evaluating designed molecules against biologically relevant structural targets. In conclusion, AdaptiveFlow offers a unified, open-source, and massively scalable solution for ultra-large virtual screening. It provides seamless integration of conventional and AI-driven docking protocols, access to the largest ready-to-dock chemical library, and adaptive strategies for efficient navigation of trillion-scale chemical space. With demonstrated success across diverse therapeutic targets, AdaptiveFlow stands as a next-generation infrastructure poised to accelerate the design and discovery of future therapeutics.

## Methods

### AdaptiveFlow

AdaptiveFlow is an in silico platform consisting of three modules (AFLP, AFVS, and AFU) and ready-made ligand libraries for virtual screens. In this section, general platform-wide features have been described that span multiple modules.

#### Python Code-base

AdaptiveFlow has been entirely rewritten in Python, a programming language that enables the implementation of more sophisticated and advanced code while reducing maintenance work. This rewrite has also made it easier for scientists and programmers to join the open-source project and/or extend AdaptiveFlow independently, as Python is the most widely used programming language in scientific communities. In contrast, the previous version of AdaptiveFlow was written in Bash.

#### Parallelism

In addition to being rewritten in Python, a key goal for AdaptiveFlow was to improve parallelism in order to decrease the overall time required for screening. AdaptiveFlow has increased the levels of parallelism compared to the previous version. To help manage this increased parallelism, it maintains the use of *collections*, that group around 1,000 ligands together, reducing the overhead of scheduling individual ligands for processing.

AdaptiveFlow also introduces the concept of *subjobs*, which are atomic units of computational work that include one or more collections for processing. These subjobs are automatically generated by AdaptiveFlow to match the runtime requirements specified in the job configuration. Subjobs are then combined into larger groups called *workunits*, which are designed to allow schedulers to efficiently process the high levels of parallelism generated by AdaptiveFlow. These work units are sent as job arrays to the end schedulers for processing. By optimizing for parallelism, the team was able to achieve 5.6 million concurrent vCPUs running AdaptiveFlow.

#### Native Cloud Support

The AFLP and AFVS modules of AdaptiveFlow can run on computer clusters as well as on the cloud by using a batch system. Currently, two batch systems are supported, Slurm (37) and AWS Batch (38). Slurm is able to run on many cloud infrastructures, including Google Cloud, Microsoft Azure, as well as AWS. Within AWS, AdaptiveFlow uses AWS Batch and S3 object storage. CloudFormation is used to set up AdaptiveFlow and all necessary components. The AWS-native architecture allows AdaptiveFlow to utilize millions of virtual CPUs (vCPUs) in parallel, while the previous version was only demonstrated to scale up to 160,000 vCPUs.

AdaptiveFlow is able to utilize various features from AWS in addition to supporting traditional clusters, making it easier to use and increasing the scale of computations available to researchers. Some of these features include:

- Scheduling and Orchestration: AWS Batch is a cloud-native scheduler and orchestrator for containerized jobs. As part of running a job, AdaptiveFlow can make use of AWS Batch to automatically scale cloud infrastructure up and down as needed and run all of the individual subjobs. AdaptiveFlow can also automatically generate the Docker container used for processing, eliminating the need to understand containerization or other infrastructure.
- Object Storage (Amazon S3): In addition to the usage of standard cluster filesystems, AdaptiveFlow also supports the usage of object storage for both ingesting of input datasets and the output of the resulting data from the processing.
- Spot Capacity Awareness and Spot Resilience: Spot capacity is unused computing capacity that cloud providers make available at significantly discounted prices compared to normal on-demand access. In exchange for a lower price, a workload running on a spot instance may be reclaimed (stopped) with a short warning. AdaptiveFlow is designed to take advantage of this capacity model by using shorter running subjobs (approximately 30 minutes) and automatically re-running jobs that have prematurely exited through integration with AWS Batch. Additionally, through integration with AWS Batch, AdaptiveFlow is able to select instances that are less likely to be interrupted through the “spot capacity aware” scheduling method.

The architecture of AdaptiveFlow when deployed on AWS is shown in Supplementary Fig. 1. The architecture makes use of multiple *Compute Environments*, each of which may contain thousands of instances (compute nodes). The instances are automatically allocated and deallocated when the job starts and ends, respectively. Individual subjobs are allocated to the instances by AWS Batch until all subjobs have been completed. These subjobs run as Docker containers on the instances.

#### ARM

AdaptiveFlow is also able to run on the compute ARM architecture when using ARM-compatible choices in the settings. AFLP is fully ARM compatible, and AFVS is ARM compatible when docking programs are used that either make the source code available or provide binaries for ARM. ARM-based computing infrastructure can be more cost-efficient than traditional x86-based architectures.

### Enamine REAL Space

The REAL Space (enumerated, version 2022q1-2) was prepared using AFVS. JChem Suite was used for neutralizing the molecules, stereoisomer enumeration, tautomer generation, protonation state prediction, 3D coordinate calculation, and the calculation of multiple molecular properties, JChem Suite 18.20.0, ChemAxon (https://www.chemaxon.com). Open Babel (version 2.3.2) was used as backup program for protonation state prediction, 3D coordinate calculation (63), and target format conversion (all formats except SELFIES). The pip-installable SELFIES package (https://github.com/aspuru-guzik-group/selfies) was used for the conversion of molecules between SMILES and SELFIES. An overview of the processing steps and preparation methods used is shown in Supplementary Table 2.

#### Multiple Formats

AdaptiveFlow provides the REAL Space library in ready-to-dock 3D formats including PDB, PDBQT, MOL2, and SDF. These formats are compatible with a wide range of docking programs for small molecules. The total size of the ready-to-dock library is approximately 50 terabytes in compressed form for each of these formats, totaling around 200 terabytes in compressed form (uncompressed, the library is around 2 petabytes). In addition to these ready-to-dock formats, the REAL Space library is also provided in the SELFIES (33), SMILES (64), and Parquet formats. These formats are useful for postprocessing the results of virtual screening (e.g., for easy representation of each molecule), and they can also be used as standalone libraries for other purposes such as in the AI/ML community, where SMILES and SELFIES are the standard molecular input file formats. The Parquet format of the REAL Space only stores the properties of the molecules, which can be useful for fast library-wide queries.

### AdaptiveFlow for Ligand Preparation (AFLP)

AFLP is the module of AdaptiveFlow dedicated to the preparation of ligand libraries into a ready-to-dock format and can handle libraries of any size, including ultra-large libraries. AFLP requires a batch system and supports Slurm as well as AWS Batch as batch systems. AFLP is configured via a configuration file (an example configuration file is shown in Supplementary Listing K). Details on the workflow are available in Supplementary Information Section D. The new version of AFLP has the following features.

#### Preparation into Ready-To-Dock Format

The primary task of AFLP is to prepare ligands into a ready-to-dock format. To accomplish this, AFLP desalts the original ligands (in case they are salts), neutralizes them, generates stereoisomers and tautomers, predicts the protonation states, computes the minimum-potential energy conformation of the ligand (3D coordinates), optionally validates the generated 3D conformers, and converts them into the desired target formats. In total over 100 output formats are supported. A detailed overview of the processing steps can be seen in Supplementary Fig. 7. The support input formats are SMILES, SELFIES, as well as amino acid sequences. AFLP uses external tools such as ChemAxon’s JChem, Open Babel, and RDKit (65), to carry out the preparation steps. For many of the preparation steps, the user can choose which external tool should be used. Details on which tools are available for each processing step can be found in Supplementary Table 3.

#### Molecular Properties Calculation

AFLP enables researchers to calculate a variety of physicochemical properties and molecular descriptors for the molecules in the input ligand library, in addition to its primary task of preparing them for molecular docking. By calculating properties such as QED (Quantitative Estimate of Druglikeness), logP, molecular weight, and others (a full list of supported properties can be found in Supplementary Table 2), researchers can more effectively select and prioritize ligands, which can help reduce costs and the time required for screens using AFVS. AFLP uses mostly external tools to calculate the molecular properties, such as ChemAxon’s JChem (https://chemaxon.com/calculators-and-predictors, Open Babel (63), and RDKit (for details see Supplementary Table 3). By combining the results of AFLP with those of AFVS, researchers can gain valuable insights into the structural and chemical characteristics of their ligands relative to specific target proteins, which can help them design and optimize new molecules with improved affinity and specificity.

#### Automatic Tranche Assignments

In AdaptiveFlow, ligand libraries are organized into tranches. Each tranche corresponds to a property of the ligands and is divided into multiple intervals or categories. For example, the molecular weight of the ligands can be used as a tranche type, and can then be partitioned into several categories or intervals such as 0-200, 200-300, 300-400, 400-500, and 500-∞ daltons. Multiple tranches allow for the partitioning of a ligand library into a multidimensional grid or table, with one dimension corresponding to each tranche type (ligand property).

While the previous version of AdaptiveFlow required that the input ligand collection was already organized and structured in the tranche format, AdaptiveFlow can automatically reorganize the input ligand library into a new tranche output format based on the computation of specific molecular properties of the ligands (see the corresponding paragraph above). This feature is useful because most ligand libraries (e.g., from chemical compound vendors) are initially in the SMILES format and not in the tranche format. AFLP can then be used to prepare the entire library in the multidimensional grid/tranche format for AFVS.

To characterize how the Enamine REAL Space library populates the 18-dimensional property grid used by Adaptive-Flow, we quantified tranche occupancy (the number of molecules assigned to each grid cell) and summarized it as a histogram (Supplementary Fig. 34). We counted molecules per tranche across the full library. The resulting occupancy counts span several orders of magnitude, so we report the distribution using logarithmically spaced occupancy ranges (1, 2–10, 11–100, 101–1K, 1K–10K, 10K–100K, 100K–1M, 1M–10M, and >10M molecules per tranche). The histogram displays the number of tranches in each occupancy range (y-axis), highlighting that only a small fraction of the theoretical grid cells are populated and that populated tranches vary widely in size (from single-molecule tranches to highly populated tranches exceeding 10 million molecules), with an average of 5600 molecules per populated tranche.

#### Support of SELFIES

SELF-referencing Embedded Strings (SELFIES) (33, 34) has become one of the most popular line notation formats for small molecules within machine-learning communities. The primary reason is that SELFIES are fully robust, meaning that every SELFIES string corresponds to a valid molecule, which is not the case for SMILES. Thus generative ML models cannot create invalid molecules when using SELFIES. AFLP now supports SELFIES as an input format for the molecules in addition to SMILES. Additionally, the entire REAL Space is free to be downloaded in the SELFIES format, enabling the creation of powerful datasets.

#### Stereoisomer Enumeration

One of the main tasks of AFLP is to prepare ligand libraries from unprepared formats (e.g., SMILES) into ready-to-dock formats. The new version of AFLP enables the enumeration of stereoisomers for the ligands, which was not possible in the previous version. Accurate stereochemistry is essential for virtual screenings using structure-based molecular docking, but many ligand libraries in unprepared formats (e.g., from chemical compound vendors) do not contain stereochemical information. The ability to enumerate stereoisomers allows AFLP to include this information in the prepared ligand library, ensuring that the virtual screening results are accurate.

#### Extended Open Source Support

The previous version of AFLP required external tools (from ChemAxon) that are not open source for some of the steps when preparing ligands in a ready-to-dock format. The new version provides the option to choose open-source alternatives for each of the steps to prepare molecules into a ready-to-dock format. In particular, Open Babel and RDKit are supported by AdaptiveFlow as open-source alternatives.

#### Enhanced Speed

AFLP has been improved in terms of speed compared to the original version. Firstly, this has been achieved by AFLP using its own Java code to call the necessary functions of ChemAxon’s JChem package via Nailgun, which reduces the number of calls to Nailgun significantly. Additionally, timeout parameters that can be adjusted by users have been added to the control file, allowing AFLP to skip ligands that require unusually long processing times during preparation. These improvements make AFLP more efficient, robust and able to handle larger and more chemically complex ligand libraries.

#### Enhanced Quality Checks

AFLP checks the validity of each ligand that it prepares in the ready-to-dock format by calculating the potential energy of the ligand in 3D format using Open Babel. This allows AFLP to automatically remove ligands that have an excessively high potential energy, which can be an indication that the compound was damaged or corrupted during the preparation process. Additionally, AFLP supports optional plausibility checks on generated 3D conformers using PoseBusters (66). This helps to ensure the integrity and accuracy of the prepared ligand library.

#### Application

AFLP was used to prepare the REAL Space (version 2022q1-2). It was run on AWS using over 5.6 million Intel vCPUs in parallel. Spot instances were used, and the used capacity was sustained without significant preemption. In total less than 0.1 percent of the used CPU hours were interrupted due to preemption. The scaling behavior of AFLP was found to be perfectly linear (as shown in Supplementary Fig. 2). This demonstrates the efficiency and effectiveness of AFLP in preparing large ligand libraries in a ready-to-dock format.

### AdaptiveFlow for Virtual Screening (AFVS)

AFVS is the module of AdaptiveFlow dedicated to ultra-large virtual screens but can also screen libraries of smaller sizes. It requires a batch system and supports Slurm as well as AWS Batch. AFVS is configured via a configuration file (an example configuration file is shown in Supplementary Listing L). AFVS and AFU share many virtual screening features, which are described in a later section. AFVS and AFLP share the same parallelization mechanisms when using Slurm or AWS Batch, and therefore the scaling behavior of AFVS is expected to be the same as for AFLP, which was shown to be perfectly linear up to 5.6 million CPUs (see also Supplementary Fig. 2).

#### Parquet Output Format

AFVS has the ability to store output data in the Apache Parquet format, an open-source column-oriented data storage format. When using this format, AFVS can take advantage of any database and query system that can read this format, including Amazon Athena, a serverless service that allows users to efficiently query output score files stored in Amazon S3 (object storage) on AWS. Additionally, AFVS has a feature for automatically postprocessing ATG Prescreens using Amazon Athena, enabling researchers to efficiently prepare the input data for the ATG primary screens. Tools are provided with AFVS to facilitate common queries, but researchers can also create their own queries using the open-source format of the dataset.

#### ATG-VSs and Benchmarks

Following completion of the ATG Prescreen, tranches can be selected for the ATG Primary Screen using two main strategies. In the first, individual tranches from the 18-dimensional REAL Space matrix can be manually selected, offering fine-grained control over specific chemical subspaces. In the second, selection can be restricted to a hyperrectangular region of the grid. This is achieved by identifying the top-performing intervals (i.e., bins) within each of the 18 molecular property dimensions and computing their Cartesian product to form a structured, intersecting subset of tranches. This hyperrectangle approach enables chemically diverse yet targeted selection based on multiple favorable properties. In both selection modes, users define the total number of molecules to include in the ATG Primary Screen, and AdaptiveFlow automatically prioritizes the tranches containing the best-performing representative ligands.

To evaluate the effectiveness of ATG-VS, we performed two types of benchmark studies: (1) full-scale production benchmarks using the entire Enamine REAL Space, and (2) smaller-scale “test set-based” benchmarks designed for rapid prototyping and detailed comparison across different ATG configurations. In the test set-based benchmarks, we randomly selected synthetic reactions from the Enamine space to create a compound subset totaling approximately 5 million molecules. For each of the 10 protein targets tested (listed in Supplementary Table 7), a different random subset was used to ensure unbiased evaluation across targets and screening scenarios.

The test set-based benchmarks allowed us to systematically assess the benefits of ATG-VS in three configurations: standard ULVS (random selection), ATG without active learning, and ATG augmented with the optional ML-based active learning component. In contrast, the full-scale production benchmarks were conducted using the entire 69 billion–compound library, but without the active learning step, to minimize computational overhead while validating the performance of ATG-VS at scale. All docking in these benchmarks was performed using QuickVina 2 with default parameters, unless otherwise specified. For the “100K ATG” benchmark configuration presented in Fig. 8, tranches for the primary screen were selected using the hyperrectangle method, enabling a balanced, property-informed sampling of chemical space.

#### Machine Learning Classifier for Tranche Prioritization

To improve the efficiency of Adaptive Target-Guided Virtual Screens (ATG-VSs), the AdaptiveFlow platform incorporates an optional machine learning (ML) classification step into the primary screening stage. This step prioritizes compounds that are most likely to be high-affinity binders, thereby reducing the number of molecules subjected to full docking. The ML classifier is applied after the ATG prescreen and is used to filter undocked molecules within the selected tranches prior to docking.

The classifier is implemented as a fully connected feedforward neural network (FNN) designed to distinguish between promising and less likely binders. Compounds are represented using Morgan fingerprints of length 1,024 and radius 2, generated via RDKit (v2023.03.1). The model is trained on docking scores obtained from all representative molecules used in the prescreen. These scores are converted into binary classification labels using a percentile-based thresholding strategy: the top 25% of compounds with the most negative docking scores are labeled as positive examples (“high-confidence binders”), while the remaining compounds are labeled as negatives. To mitigate class imbalance, the majority class is undersampled during training.

The neural network architecture comprises four fully connected layers: an input layer of 1,024 nodes, followed by hidden layers of 512, 256, and 128 nodes respectively, each using ReLU activation. The output layer consists of a single sigmoid-activated node that outputs the predicted binding probability. Hyperparameters and layer configurations were selected through limited tuning using cross-validation on the prescreen data. The model is trained with the Adam optimizer (learning rate 0.001), binary cross-entropy loss, and early stopping based on validation loss with a patience of 5 epochs. A validation split of 10% is used, with training capped at 50 epochs and a batch size of 256, although convergence typically occurs earlier.

Once trained, the model is applied to all undocked compounds within the selected tranches. Compounds with predicted binding probabilities greater than 0.5 are retained for full docking, while the rest are filtered out. This filtering step typically reduces the number of compounds subjected to docking by 30–70%, depending on the target, while maintaining or improving hit enrichment.

The ML classification module is fully integrated into the AdaptiveFlow Unity (AFU) environment and can be activated through a configuration parameter. The training and inference pipeline is implemented in PyTorch (v2.0.1) and supports GPU acceleration for scalable and efficient deployment.

### AdaptiveFlow Unity

AdaptiveFlow Unity (AFU) is the all-in-one version of AdaptiveFlow, combining both ligand preparation and virtual screening into a single streamlined workflow. It is the third module besides AFLP and AFVS. This new module adds many conveniences for workflows where both ligand preparation and virtual screenings have to be carried out in tandem. Notably, AFU is designed to be run as a package, supporting approximately 1500 docking protocols that can be run independently of job schedulers (see Supplementary Data Table 1). A conceptual overview of the workflow of AFU can be found in Supplementary Fig. 11, and a detailed UML workflow diagram is shown in Supplementary Fig. 5. Specifically, as input, a user specifies the molecule to be docked (as a SMILES, SELFIES, or amino-acid sequence), the protein file, the docking site, and the docking program to be run. Based on the user’s choices, error checks in the form of missing docking-choice-specific files are performed, and calculations are run. As a result of a successful calculation, the docking score and pose are returned. AFU can be run in standalone mode, where it is run via the command line and specifies the options via a configuration file (an example configuration file is shown in Supplementary Data Fig. M). This mode is for example useful for scientists in the drug discovery communities who want to carry out molecular dockings in the most convenient and simple way, without requiring computational expertise, while having a large number of docking protocols available. In addition, AFU can be run in API mode to directly interface with other programs and code, which can for instance be useful for the machine learning communities that develop new methods for drug discovery.

AFU comes in two versions. The first is dedicated to single workstations or compute nodes without multinode parallelization. The second, AFUparr, is the parallelized version of VitualFlow Unity. Designed to operate seamlessly on SLURM systems, AFUparr allows users to easily parallelize the workflow on a larger number of CPUs and GPUs. Execution within SLURM environments is highly optimized, with computations distributed in parallel across multiple CPUs and nodes. This design ensures efficient linear scaling relative to the number of molecules provided.

### Joint Features of AFVS and AFU

The AFU and AFVS modules support the same docking protocols and therefore share all docking-specific features.

#### Supported docking protocols

AdaptiveFlow supports approximately 40 docking protocols, listed in Tables 4, 5, 6. These include software that performs both pose prediction and scoring (Table 4), solely pose prediction (Table 6) and only scoring (Table 5). Notably, within AdaptiveFlow, the scoring functions can be combined with pose prediction methods, giving rise to approximately 1500 methods (listed in Table 1). We note that many of the newly supported docking software possess special features such as protein-protein/RNA docking, explicit parameterization of solvents/metals, and covalent docking. In Tables 4,5,6, we describe these as “Special features”. Additionally, a key consideration for ULVS is the timing required for carrying out a single calculation. As such, in Fig. 12, we provide timing estimates for running a single calculation, averaging over 25 independent runs for all docking programs.

#### Special Types of Ligands

In addition to traditional small molecule ligands, AdaptiveFlow supports the use of peptides, protein ligands, and covalently-bonded ligands in AFVS. Peptides are of increasing interest as therapeutics, and several docking programs already support their use. For protein-protein/RNA docking, specialized programs require two protein/RNA targets in order to generate co-complexes. Covalent docking involves reduced flexibility for certain parts of the input molecule during pose sampling, and a few specialized docking programs support this ligand type (see Supplementary Table 4).

#### Enhanced Performance

The QuickVina 2 molecular docking tool is used to accelerate AutoDock Vina with no loss of accuracy (67). QuickVina 2 execution was profiled in an effort to improve performance even further. To establish baseline performance, the open-source GitHub repository was cloned and the qvina_1buffer branch was compiled using the GNU C++ compiler (version 9.4), the Boost C++ libraries (version 1.79, available from https://www.boost.org), and the default parameters (see https://qvina.github.io/compilingQvina2.html for the build instructions). A single docking experiment of Avanafil binding to unsolvated 5FNQ was run through the Intel VTune Profiler to look for performance bottlenecks. Unfortunately, the execution profile was flat. The most time-consuming function accounted for less than 20% of the total run-time. Tuning the source code would give rapidly diminishing returns at the risk of affecting accuracy, so compiler-level optimization was the best option to improve performance. QuickVina 2 was rebuilt using the Intel oneAPI DPC++/C++ Compiler (version 2022.2.0.20220730) and aggressive optimization: automatic vectorization, interprocedural optimization, and relaxed floating-point constraints. Profile-guided optimization was also tried, but it did not improve performance. The final executable was 18% faster than the default builds with no loss of accuracy. The new binaries are provided on the QuickVina and AF homepages.

#### GPU Support

Several of the docking programs support graphics processing units (GPUs), which can strongly reduce computation time and cost. For pose prediction, these include softwares: AutoDock-GPU (59), QVina2-GPU (60), QVina2-W-GPU (60), Vina-GPU (60). Additionally, deep-learning methods such as TANKBind (12), DiffDock (68), EquiBind (69) use GPU-compatible neural networks for direct pose prediction of several ligands in parallel. Additionally, Gnina (70), NNScore 2.0 (71), DeepAffinity (72) and DeepBindRG (73) use GPU-compatible deep learning models for scoring protein-ligand complexes. AdaptiveFlow supports GPU compatibility of all these software. The relative speedups of these methods for single calculations can be compared within Fig. 12.

#### Molecular Dynamics-Based Methods

Molecular docking is a computationally fast way to predict binding poses and binding-free energies, but it has sometimes trouble distinguishing between compounds with similar binding affinities. Molecular docking algorithms often prioritize speed over accuracy by using fast pose generation and approximate scoring functions to save time, especially when screening extensive collections of compounds against a protein target. This trade-off reduces the effectiveness of the algorithms. More precise algorithms based on molecular dynamics simulations are sometimes used downstream of docking in drug discovery processes to improve the number of accurate predictions. The combination of molecular docking and molecular dynamics can yield results near experimentally determined crystal structures (74, 75). AFVS includes methods for selecting and scoring poses based on molecular dynamics simulations, such as MM/PBSA and MM/GBSA, for evaluating docking poses and predicting binding affinities (76). MM/PBSA and MM/GBSA are popular for binding free energy prediction because they are more accurate than molecular docking and less computationally intensive than alchemical methods. AdaptiveFlow also supports Binding Pose Metadynamics (BPMD), which uses a stability score based on the resistance of the ligand to perturbation away from the initial pose to rerank predicted binding poses from docking (74, 77). The OpenBPMD algorithm can accurately predict binding poses within 2 Å RMSD of the crystallographic complex structure for many systems and efficiently rerank docked poses. The MM/PBSA, MM/GBSA, and BPMD methods can be used within AdaptiveFlow as rescoring and pose refinement methods.

#### Enhanced Quality Checks

The docking pose of each ligand screened by AFU is checked for corruption by evaluating the coordinates and computing the potential energy of the docked compound with Open Babel. Any corrupted compounds are automatically removed. AFVS supports optional plausibility checks on generated docking poses of Vina-based docking protocols using PoseBusters, an open-source tool, which checks for chemical as well as intramolecular ligand and intermolecular validity of ligand-protein complexes (66).

### Experimental Validation Studies Involving FSP1

Materials and methods used in the experimental validation of the molecules for FSP1 are described below.

#### Chemicals

All chemicals and proteins employed for binding and enzymatic assays were purchased from Sigma Aldrich, except for 2,3,5-trimethylcyclohexa-2,5-diene-1,4-dione (α-tocopherolquinone hydrophilic head) purchased from Synthonix and menaquinone (Vitamin K2) from Supelco Analytical. FSP1 inhibitor candidates were purchased from Enamine; iFSP1 from Medchemexpress; viFSP1 and FSEN1 from Cayman Chemical. Inhibitors were analyzed by NMR (supplementary data). Samples were prepared by dissolving the compounds in deuterated DMSO (d6-DMSO) to 1 mM final concentration. Spectra were acquired with 100 scans on a Bruker Avance III 400 MHz spectrometer and processed using TopSpin version 4.5.0.

#### Phylogenetic Inference and Ancestral Sequence Reconstruction

Phylogenetic analysis and ancestral sequence reconstruction were carried out following the methods described in (78). Briefly, human FSP1 sequence (UniprotID: Q9BRQ8) was employed as query for homology searches with BLASTP. Datasets were constructed vetting all chordate classes according to TimeTree (79) by mining at least two species with fully sequenced genomes. Multiple sequence alignments were constructed in MAFFT v7 (80) and manually trimmed for single sequence insertions/terminal extensions. Neighbour-joining guide trees were constructed in MEGA v10.2 to assess the quality of datasets under construction. Once the working multiple-sequence alignments were obtained, best-fit substitution models and gamma distribution values (α) were calculated using ProtTest (data for each protein phylogeny are shown in Supplementary Fig. 12B). Maximum likelihood phylogenies were inferred using RaxML v8.2.10 (HPC-PTHREADS module (81)), using rapid bootstrap analysis and searching for the best-scoring ML tree, with 500 bootstraps replicates and the given best-fit model under gamma distribution. When required, a species tree was used to constrain the phylogeny, this was constructed using TimeTree (timetree.org). Once the phylogenies were inferred, the bootstrap values were subjected to transfer bootstrap expectation values using BOOSTER online (82). Figtree v1.4.2 was employed for analysing and visualizing the trees. Ancestral sequence reconstruction was performed employing PAML v4.9a (CODEML module) as marginal reconstruction, using the phylogenies obtained previously, empirical amino acid substitution model (model=2) and JTT substitution matrix, four gamma categories and re-estimation of gamma shape parameter (83, 84). The distribution of the posterior probabilities for each of the ancestral states was analysed at the node corresponding to the tetrapod ancestor in the phylogeny. Sites that displayed posterior probabilities <0.8 were considered ambiguously reconstructed when alternative states displayed posterior probabilities >0.2 (85). The length of the targeted nodes was treated by Fitch’s parsimony. To visualize the degree of sequence conservation across tetrapod organisms, multiple sequence alignments were generated using ESPript 3 (86).

#### Construct design and expression of FSP1

The gene encoding for human full-length FSP1-twinSTREP-eGFP was cloned into pcDNA3 (Genscript). HEK293-EBNA1-6E cells were used for the transient transfection. Cells were maintained in suspension with FreeStyle medium (Invitrogen). The day before transfection, cells were seeded at a concentration of 0.5 × 10^5^ cells/mL. The day after (1 × 10^6^ cells/mL), cells were transiently transfected using polyethyleneimine with a ratio of 1:3 DNA:polyethyleneimine. After 48 h, cells were harvested by centrifugation at 1,200 g for 5 min at 20 °C. Trypan Blue exclusion assay (Trypan Blue solution w/v, Sigma) was used to evaluate cell viability. The genes encoding for His_6_-SUMO-ancestral FSP1, His_6_-SUMO-ancestral L323A FSP1, His_6_-SUMO-ancestral F15A FSP1, His_6_-SUMO-ancestral F354A FSP1, lacking the Gly-2 myristoylation site, were cloned in a pET-24a(+) vector with kanamycin resistance (Genscript). Plasmids were then transformed by heat shock into *E. coli* BL21(DE3) cells (25 s at 42 °C). Cells from a single colony were picked and pre-inoculated into 100 mL LB broth containing 100 µg mL^-1^ kanamycin and grown overnight at 37 °C. 10 mL of pre-cultures were inoculated into 1 liter of Terrific Broth and grown at 37 °C, 200 r.p.m. for 3 h until the optical density (OD_600_) reached 0.5-0.7. Protein expression was induced with isopropyl β-D-1-thiogalactopyranoside (0.1 mM final) and incubated overnight at 30 °C while shaking 200 r.p.m.. Cells were harvested by centrifugation (5,000 g, 15 min, 4 °C), flash frozen in liquid nitrogen, and stored at −20 °C.

#### Protein purification

Cells (ca. 29 g) were re-suspended 1:5 (grams:milliters) in Buffer A (50 mM HEPES pH 7.2, 250 mM NaCl, 10% (v/v) glycerol) supplemented with 1 mM FAD and protease inhibitors (1 mM phenylmethylsulphonyl fluoride, 10 µM leupeptin, 10 µM pepstatin) and 5 µg DNAse I /g cells. Cell resuspensions were stirred at 4 °C for 25 min before cell lysis, which was performed with a high-pressure homogeniser (Emulsiflex c-3, ATA Scientific) in three cycles. Cell lysates were centrifuged (4 °C, 56,000 g, 1 h) using an Avanti J-26 XP centrifuge equipped with a JA-25.15 rotor (Beckman Coulter). The resulting supernatant was filtered through a 0.45 µm filter and loaded on a gravity column containing Nickel Sepharose HP IMAC resin (Cytiva) pre-equilibrated with Buffer A. The crude extract was passed onto the column twice. The resin was then washed with at least five column volumes of Buffer A. The wash up of the resin was performed adding 5 column volumes of Buffer A supplemented with increasing concentrations of imidazole, 5 to 30 mM. The protein was then eluted with at least ten column volumes of Buffer B (Buffer A supplemented with 300 mM imidazole). The purified protein sample was concentrated using an Amicon Ultra (Merck) with a 30 kDa cut-off up to 2.5 mL. The imidazole was then removed with a PD-10 batch desalting column (Cytiva) pre-equilibrated with Buffer A. To cleave the His_6_-SUMO tag, the desalted sample was concentrated down to 800 µL and then added of a His_6_-tagged SUMO protease (1.2 mg mL^−1^) to a volume ratio of 100:1 and incubated overnight at 4 °C on a rotating wheel. The sample was then loaded onto an Äkta Pure system (Cytiva) equipped with a Ni HiTrap-HP 5 mL column (Cytiva) pre-equilibrated in Buffer A and proteins were eluted out using an imidazole concentration gradient of 2% Buffer B corresponding to 6 mM imidazole. The elution of the tag-less protein was monitored with a multiwavelength detector following the absorbance at 280 nm and 450 nm. The sample was loaded onto an Äkta Pure system (Cytiva) equipped with a Superdex 200 10/300 GL (Cytiva) pre-equilibrated in 100 mM NaCl, 50 mM HEPES pH 7.2. The purified protein was then concentrated down to about 1 mM, measuring the concentration using a NanoDrop ND-100 UV/Vis spectrophotometer (Thermo Scientific) based on the absorption at 280 nm and at 458 nm (ε280 nm predicted by ProtParam tool - Expasy). Purity of the sample was evaluated by SDS-PAGE. The purified protein was finally flash frozen in liquid nitrogen and stored at −80 °C.

#### Enzyme kinetics

The activity of FSP1 and mutants was monitored using 0.2-1 µM protein (estimated by 458 nm absorbance) in a 150 µL final volume of Buffer A. Reactions were followed in 10.00 mm quartz cuvettes (Hellma) and a Cary Eclipse Fluorescence Spectrophotometer (Agilent) or a Cary 100 UV-vis spectrophotometer (Varian) equipped with a thermo-stated cell holder (T = 25 °C). Reactions were started by adding NADH and rates were determined by following NADH consumption, caused by the oxidation to NAD^+^ (excitation 340 nm, emission 460 nm or monitoring the absorbance at 340 nm). A calibration line was built by measuring the fluorescence at known NADH concentrations. GraphPad Prism 9 was used to perform both linear and non-linear regression.

#### Superoxide assay

Superoxide radicals generated by oxidation of NADH were detected by the reduction of nitroblue tetrazolium to nitroblue diformazane using Tris-HCl buffer (16 mM, pH 8.0) 50 µM nitroblue tetrazolium, and 0.2 µM final FSP1 (final volume 100 µL). The reaction was started by adding various concentrations of NADH. The absorbance was measured at 560 nm using a ClarioStar plate reader.

#### Enzyme inhibition

K_i_ were measured following NADH oxidation by monitoring the absorbance 340 nm with Cary 100 UV-vis spectrophotometer (Varian). 0.2-1 µM purified FSP1 was added to a final volume of 150 µL of 50 mM HEPES buffer pH 7.2. Inhibitors were tested with concentrations varying from micromolar to low nanomolar.

#### Resazurin assay

The activity of FSP1 and mutants was monitored using 0.2-0.1 µM (final, absorbance at 458 nm) protein in a 100 µL 50 mM HEPES buffer pH 7.2 and 50 µM resazurin. The activity was monitored by the formation of resorufin (excitation. 572 nm, emission 583 nm) using varying concentrations of iFSP1 and viFSP1 (2, 44, 45), afi-FSP1-1, afi-FSP1-2, afi-FSP1-3, afi-FSP1-4, afi-FSP1-5 and afi-FSP1-6. The reaction was started by adding 200-100 µM NADH. The fluorescence was read using a ClarioStar plate reader, with a gain of 800.

#### Nano Differential Scanning Fluorimetry

Thermo-stability analyses were carried out using a TychoTMNT.6 system (NanoTemper GmbH). The protein was diluted to 1 mg mL^-1^ in Buffer A and pre-incubated with ligands at room temperature for 15 minutes before experiments. Melting curves were obtained following the intrinsic fluorescence of tryptophan and tyrosine residues (emission 350 nm and 330 nm, respectively) applying a temperature gradient from 35 °C to 95 °C for a time range of three minutes. Data were analysed as F_350_/F_330_ ratio and derived to determine the inflection temperature (T_m_). All measurements were performed in duplicate. The following condition were tested: FSP1 in presence of 1 mM FAD, NAD^+^ or 1 mM CoQ_1_ or 1 mM α-tocopherolquinone hydrophilic head or 1 mM vitamin K3 or 10 µM inhibitors or 5% DMSO v/v as a control.

#### ThermoFAD

Experiments were performed with 20 µM FSP1 after 10 min incubation with inhibitors (10 µM) or 5% v/v DMSO in HEPES buffer (50 mM, pH 7.2) (87). The temperature gradient was set to 30–90 °C with fluorescence detection at a constant rate 0.1 °C/s at 450 ± 30 nm excitation and 535 ± 30 nm emission (BioRad MiniOpticon Real-Time PCR System). Data were normalized and fitted with sigmoidal non-linear regression using GraphPad Prism 9.

#### Cellular thermal shift assays

Inhibitor afi-FSP1-1: HEK293-EBNA1-6E cells were seeded at a density of 3.0 × 10^7^ cells/mL and incubated with either 5% v/v DMSO or 10 µM of 1 for 1 h at 37 °C in 2 ml of culture medium. Cells were collected by centrifugation at 1,200 × g for 5 min at room temperature, washed twice with PBS, and resuspended at the initial cell concentration. Aliquots of 50 µL were distributed into PCR strip tubes. Samples were subjected to a heat treatment at different temperatures (ranging from 38.7 to 86.2 °C) using a Bio-Rad T100 Thermal Cycler. Following heat exposure, cells were lysed by adding 1% Triton X-100, protease inhibitors (see Protein purification section), and DNase I (5 µg per g of cells). Lysis was performed for 1 h at 4 °C on a rotation wheel. Lysates were centrifuged at 20,000 × g for 20 min at 4 °C, and 15 µL of each supernatant was loaded onto Bio-Rad Mini-PROTEAN TGX SDS-PAGE gels. Proteins were transferred to 0.2 µm PVDF membranes (Bio-Rad) using the Trans-Blot Turbo system. Membranes were exposed with an anti-AIFM2 antibody (clone 1I15 ZooMAb®, rabbit monoclonal, Sigma-Aldrich), followed by an HRP-conjugated anti-rabbit secondary antibody (Millipore). Chemiluminescent signals were detected with the Bio-Rad ECL detection kit on a ChemiDoc MP imaging system. For data analysis, band intensities at each temperature were quantified and normalized by setting the strongest and weakest signals to 100% and 0%, respectively. Melting curves were fitted using a Boltzmann sigmoidal model in GraphPad Prism 9. Aggregation temperatures were calculated from two independent biological replicates, and results are reported as single replicates.

Inhibitors afi-FSP1-2, 3, 4: 1.5 × 10^7^ HEK293-EBNA1-6E cells/mL expressing human FSP1-twinSTREP-eGFP were treated with 1 µM compound 2, or 10 µM compound 3, or 10 µM compound 4 or 5% v/v DMSO following the above-described protocol. The heat treatment ranged from 31.6 to 78.1 °C. PVDF membranes containing proteins were incubated with Precision Protein™ Strep-Tactin® HRP-conjugate (BioRad). The detection and data analysis were described before.

#### Immunoblot analysis

Jurkat T-lymphoid cell line was obtained from ATCC and cultured in RPMI 1640 with 10% FBS, 100 units/mL penicillin and streptomycin. The cells were lysed using sonication in M-PER buffer (cat:78501 from Thermo Scientific) and 50 µg of total protein per lane was loaded on the gel. FSP1 antibody is a Rabbit-derived cat. # HY-P80679 from MCE; GPX4 antibody is mouse-derived cat. # 67763-1-1g from Protein tech; GAPDH antibody (6C5) is mouse-derived, cat. # AM4300 from Invitrogen

#### Cell viability assays

Trypan blue dye exclusion was used to determine cell viability. Logarithmic phase cells were seeded at 1 × 10 ^5^ cells/mL in a 12-well plate. Cells were then treated with either DMSO (control) or the test compounds solubilized in DMSO, gently mixed and incubated for 24 hours under culture conditions as described. Viability was assessed by removing an aliquot (10 µL) of the cell suspension and mixing with an equal volume of 0.4% Trypan Blue solution. Live cells were then enumerated by light microscopy. Viability was expressed as a percentage of the DMSO control number.

#### Statistical Analysis and Reproducibility

Significance levels are indicated in legends of each figure as follows: *P< 0.05, **P< 0.01, ***P<0.001, ****P< 0.0001 (Student’s t test). Tests used to derive significance are indicated in legends. Individual data points are indicated in graphs. In the latter case all replicates are independent and not multiple measurements taken from a single sample. Analysis of significance was done using unpaired Student’s t-test, two-sided.

#### Crystallisation and structural determination of FSP1

Crystallisation was performed using 13 g/mL in 50 mM HEPES pH 7.2, 1 mM FAD, 1 mM NAD^+^ with or without 1 mM coenzyme Q_1_. Small crystals or crystalline precipitate appeared in a few days under several conditions of the Mopheus screening kit (Molecular Dimensions) using the vapour diffusion sitting drop methods. The best-looking crystals grew in HEPES/MOPS 0.1 M pH 7.5, 12.5% PEG 1000, 12.5% PEG 3350, 12.5% MPD, 0.06 M MgCl_2_, 0.06 M CaCl_2_ at 20 °C with a ratio of 1:1 in a sitting drop. After harvesting, they were smashed to generate small crystals, and used for microseeding. Specifically, the drops were composed of 0.15 µL protein, 0.05 µL seeds, and 0.2 µL reservoir from the following commercial screens.

- FSP1-FAD structure; 0.06 M MgCl_2_, 0.06 M CaCl_2_, 0.1 M HEPES/MOPS (acid) pH 7.5; 20% v/v PEG500*MME, 10% PEG20000.
- FSP1-FAD-NAD^+^ structure; 0.06 M MgCl_2_, 0.06 CaCl_2_, 0.1 M HEPES/MOPS (acid) pH 7.5, 12.5% PEG1000, 12.5% PEG3350, 12.5% MPD.
- FSP1-FAD-NAD^+^-coenzyme Q_1_ structure; 0.06 M MgCl_2_, 0.06 CaCl_2_, 0.1 M imidazole/MES monohydrate (acid) pH 6.5, 20.0% PEG500* MME, 10% PEG20000. Crystals were then soaked with 20 mM coenzyme Q_1_ added to the crystallisation drop.
- FSP1-FAD-NAD^+^-afi-FSP1-1; 0.06 M MgCl_2_, CaCl_2_, 0.1M imidazole/MES monohydrate (acid) pH 6.5, 12.5% MPD, 12.5% PEG1000, 12.5% PEG3350. Crystals were then soaked with 1 mM afi-FSP1-1 added to the crystallisation drop.
- FSP1-FAD-NAD^+^-afi-FSP1-3; 0.12 M diethylene glycol, 0.12 M triethylene glycol, 0.12 tetraethylene glycol, 0.12 pentaethylene glycol, 0.1 M tris (base)/bicine pH 8.5, 12.5% MPD, 12.5% PEG1000, 12.5% PEG3350. Crystals were then soaked with 10 mM afi-FSP1-3 added to the crystallisation drop.
- FSP1-FAD-NAD^+^-afi-FSP1-4; 0.06 M MgCl_2_, 0.06 M CaCl_2_, 0.1 M imidazole/MES monohydrate (acid) pH 6.5, 12.5% MPD, 12.5% PEG1000, 12.5% PEG3350. Crystals were then soaked with 10 mM afi-FSP1-4 added to the crystallisation drop.
- FSP1(L323A)-FAD-NAD^+^-afi-FSP1-2; 0.06 M MgCl_2_, 0.06 M CaCl_2_, 0.1 M HEPES/MOPS (acid) pH 7.5, 20% ethylene glycol, 20% PEG8000. Crystals were then soaked with 1 mM afi-FSP1-2 added to the crystallisation drop.

Electron densities from co-crystallisations with previously reported inhibitors (Supp. Table 5) did not show any ligand binding. Data were collected at the European Synchrotron Radiation Facility (Grenoble, France) and processed with the XDS and CCP4 packages (88, 89). The phase problem was solved by molecular replacement using an Alpha Fold model and CCP4 programs. Model building and refinement were conducted using COOT and Refmac5 (90). Figures were then generated using ChimeraX (91), PyMOL (DeLano Scientific; www.pymol.org).

#### SPR Binding Assays

SPR binding assays were carried out using the Biacore 8K SPR system (Cytiva) with the running buffer (50 mM HEPES, pH 7.2, 150 mM NaCl, 0.05% Tween-20, and 2% DMSO) under controlled conditions at 25 °C. The His-Sumo–FSP1 protein was immobilized on a preconditioned and activated Series S NTA sensor chip (Cytiva). Pre-conditioning involved treatment with 350 mM EDTA, followed by activation with 0.5 mM NiCl_2_. To enhance stability and minimize signal drift, the chip was further activated using EDC (1-ethyl-3-(3-dimethylaminopropyl) carbodiimide) and NHS (N-hydroxysuccinimide) before coupling the protein to the surface. The protein was immobilized to a response level of 7500 RU.

Compounds were applied in a three-fold serial dilution series (6, 18.9, 60, 189, 600, 1897, 6000, 18973, and 60000 nM) using a single-cycle kinetics (SCK) setup. Compound plating and DMSO normalization were performed using the Tecan D-300e digital dispenser (Tecan Life Sciences). Binding was monitored for 60 s during the association phase and 120 s during the dissociation phase. Reference channel signals and buffer blanks were used to correct the raw binding signals (RU).

Binding data were collected from three independent experiments; however, only a single representative dataset is shown for each compound in the manuscript. Data processing and binding analysis were performed using Biacore Insight software (Cytiva). Steady-state binding levels (*R*_eq_) were plotted against compound concentrations and fitted to a one-site binding model using the Levenberg–Marquardt algorithm, enabling accurate determination of binding affinities for each compound.

#### Virtual Screens

The AdaptiveFlow open-source drug-discovery platform was used to carry out the virtual screen. The size of the docking box was set to 14 × 18 × 16 Å. The FSP1-FAD-NAD^+^ complex structure was held rigid during the virtual screens (ATG prescreen and primary screen). QuickVina 2 (92) using a hybrid empirical and knowledge-based scoring function was used as the docking program. The exhaustiveness value was set to 1. The number of replicas was set to 1. The virtual screen was run in Amazon Web Services (AWS). Postprocessing of the results including clustering with the similarity limit set to 0.8 was done with DataWarrior (v6.0.0) (93). In detail: For each target, the virtual library was docked and the top ~2,000 compounds by docking score were retained for further processing. Compounds flagged by DataWarrior “Nasty Functions” and PAINS alerts were removed to exclude known assay-interfering or chemically problematic substructures (e.g., strong Michael acceptors, redox-active quinones, rhodanines, catechols/hydroquinones, azo dyes). The remaining compounds were filtered by simple physicochemical criteria, requiring predicted aqueous solubility ≥ 10 *µ*M, overall aromaticity ≤ 75%, and fraction of sp^3^ carbons ≥ 0.1. This filtered set was then clustered by 2D similarity, and from each cluster one or more representatives were chosen, prioritizing the best docking scores, to yield a final diverse set of ~80 compounds per target for experimental testing. Apart from these automated substructure and physicochemical filters and the docking scores, no additional interaction-based or other structural criteria were applied.

#### Discovery of Inhibitor Analogs

Analog Hunter and Scaffold Hopper modules in the infiniSee (94) module from BioSolveIT were used to screen analogs of compound afi-FSP1-1. Top-ranking 5000 compounds from each module were exported to SeeSAR (95) from BioSolveIT and docked with the FlexX algorithm in the general docking mode to the coenzyme Q binding site of the FSP1/FAD/NAD/afi-FSP1-1 complex structure (PDB code: 9IFU) with compound afi-FSP1-1 removed. The docked poses were scored with the HYDE scoring function. Top-ranking 168 compounds with a similar binding pose as compound afi-FSP1-1 were tested with 100-ns-long molecular dynamics (MD) simulations with the PMEMD module (CUDA version) of the AMBER 22 package and protein.ff19SB force field for protein (96). The Molecular Mechanics/Generalized Born Surface Area (MM/GBSA) binding free energies between protein and the ligands that remained in the binding pocket were calculated by the MMPBSA.py script in AmberTools 23 (97). The top-ranking 20 analogs with the lowest MM/GBSA binding free energies were ordered from Enamine for testing.

## Data Availability

The ready-to-dock Enamine REAL Space can be explored online via an interactive web interface available on the AdaptiveFlow homepage http://adaptive-flow.ai/. The data can be accessed directly via an AWS S3 bucket provided by AWS Open Data (https://aws.amazon.com/opendata). Additional information is available on the AWS Open Data website of this dataset https://aws.amazon.com/marketplace/pp/prodview-m32p7engdxk7k?sr=0-1&ref_=beagle&applicationId=AWSMPContessa, as well as on the AdaptiveFlow homepage https://adaptive-flow.ai/real-space-2022q12 (free registration required).

## Code Availability

The source code of AdaptiveFlow is released under the GNU GPL v2.0, a free and open-source license. The code can be accessed via GitHub. The AdaptiveFlow GitHub repos can be found at https://github.com/QuantumAI4Bio, which contains multiple repositories. The AFLP and AFVS modules of AdaptiveFlow can be found in AFLP (https://github.com/QuantumAI4Bio/AdaptiveFlow-LP) and the AFVS (https://github.com/QuantumAI4Bio/AdaptiveFlow-VS) GitHub repositories, respectively. The AFU module (both single machine and parallelized versions) can be found in the GitHub repo (https://github.com/QuantumAI4Bio/AdaptiveFlow-Unity).

## Acknowledgments

The authors would like to thank ChemAxon for a free academic license for JChem. AK.N. thanks Anshul Kundaje and Michael Bassik for their support. The authors thank AWS for research credits for the AWS Cloud and thank Nelson Gonzales, Stephen Litster, Mayank Thakkar, Robert Wang, Arun Subramaniyan, and Christine Tsien Silvers for technical and administrative support within AWS. The authors would like to thank Akanksha Bilani and Caroline Rhoades from Intel for their support, and Brigitte Klein for proofreading the manuscript. AK.N. acknowledges funding from the Bio-X Stanford Interdisciplinary Graduate Fellowship (SGIF). C.S. and K.F. acknowledge funding by the Deutsche Forschungsgemeinschaft (DFG, German Research Foundation) under Germany’s Excellence Strategy – The Berlin Mathematics Research Center MATH+ (EXC-2046/1, project ID: 390685689). H.A. gratefully acknowledges support from the NIH/NIGMS (GM136859 and GM158220), a grant from Pivotal Life Sciences, and a gift from J. Goldberg through Dana-Farber. A.M acknowledges funding from the European Research Council (ERC Advanced Grant, MetaQ no. 101094471), and by the Associazione Italiana per la Ricerca sul Cancro (AIRC) Investigator Grant (no. 28754). This work was supported by ALSAC, the fundraising and awareness organization of St. Jude Children’s Research Hospital, and the Blue Sky Kinase Project at St. Jude Children’s Research Hospital.

## Extended Data

**Extended Data Fig. 1.**
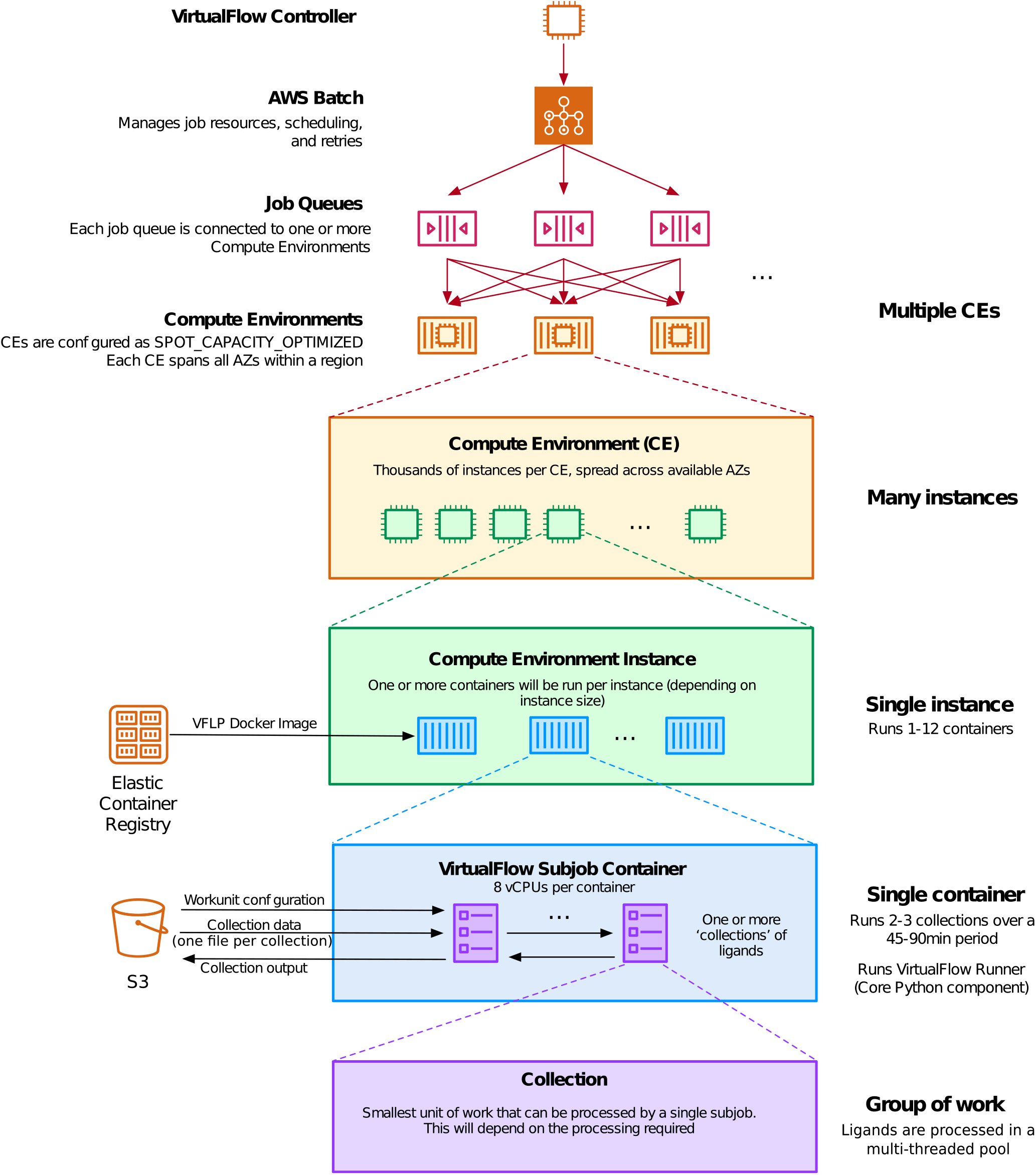
Compute Stack deployed by AdaptiveFlow in AWS. The login node is used by the user to control the workflow in AWS via the workload manager AWS Batch. AWS Batch has multiple Job Queues to which jobs are submitted by AdaptiveFlow, and each job queue deploys multiple Compute Environments that can spin up and use a large number of virtual machines (instances) of specific types. On each instance, one or more subjobs are run, and each subjobs deploys a docker container that carries out the computations of AdaptiveFlow by executing AdaptiveFlow Runner (a core python component of AdaptiveFlow). Each subjob typically processes a few ligand collections, where each ligand collection contains multiple ligands (e.g. 1000).

**Extended Data Fig. 2.**
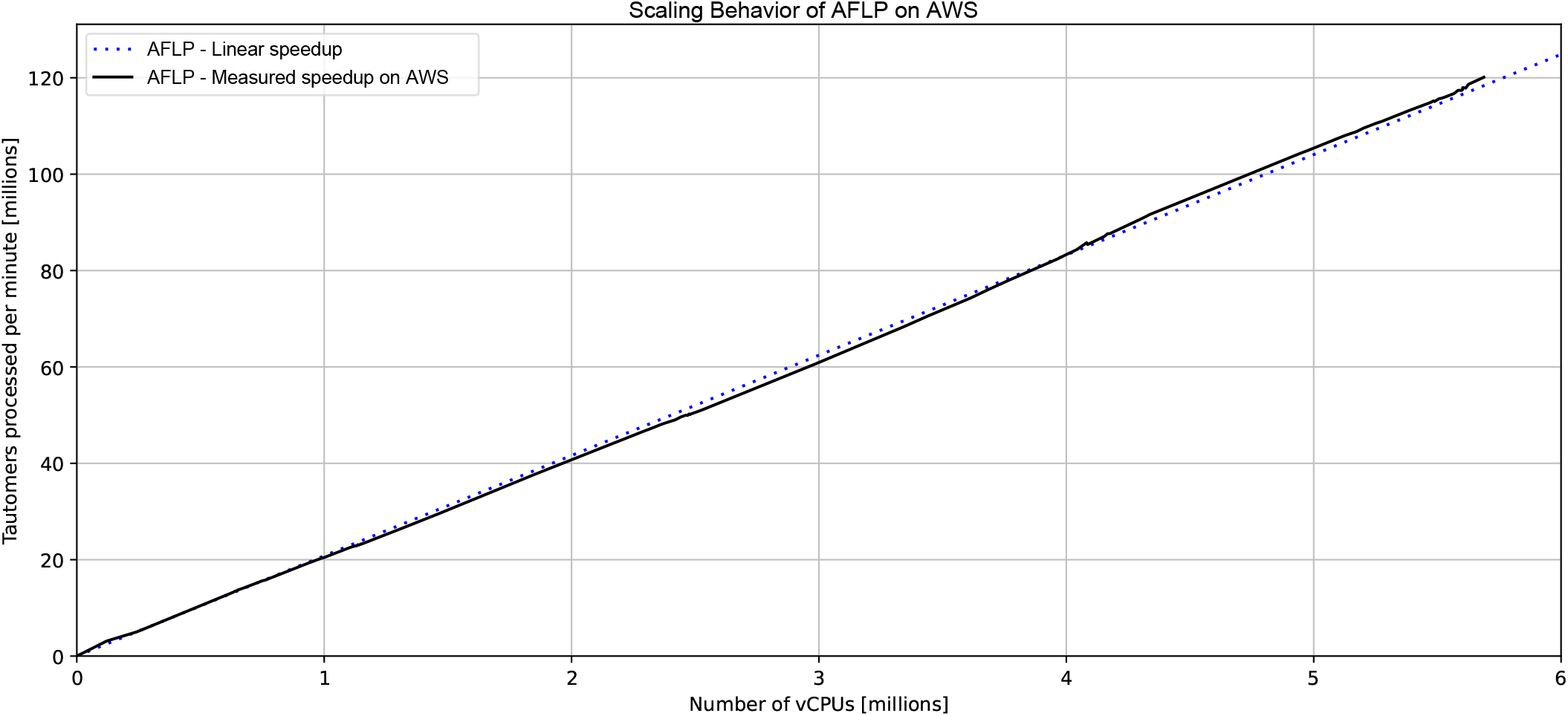
Scaling behavior of AFLP of on AWS. The scaling behavior is perfectly linear even when using extremely large numbers of CPUs. Up to 5.6 million vCPUs were used in parallel across multiple AWS regions. As can be seen in the plot, the scaling behavior is perfectly linear, implying there is no slowdown even when using extremely large numbers of vCPUs.

**Extended Data Fig. 3.**
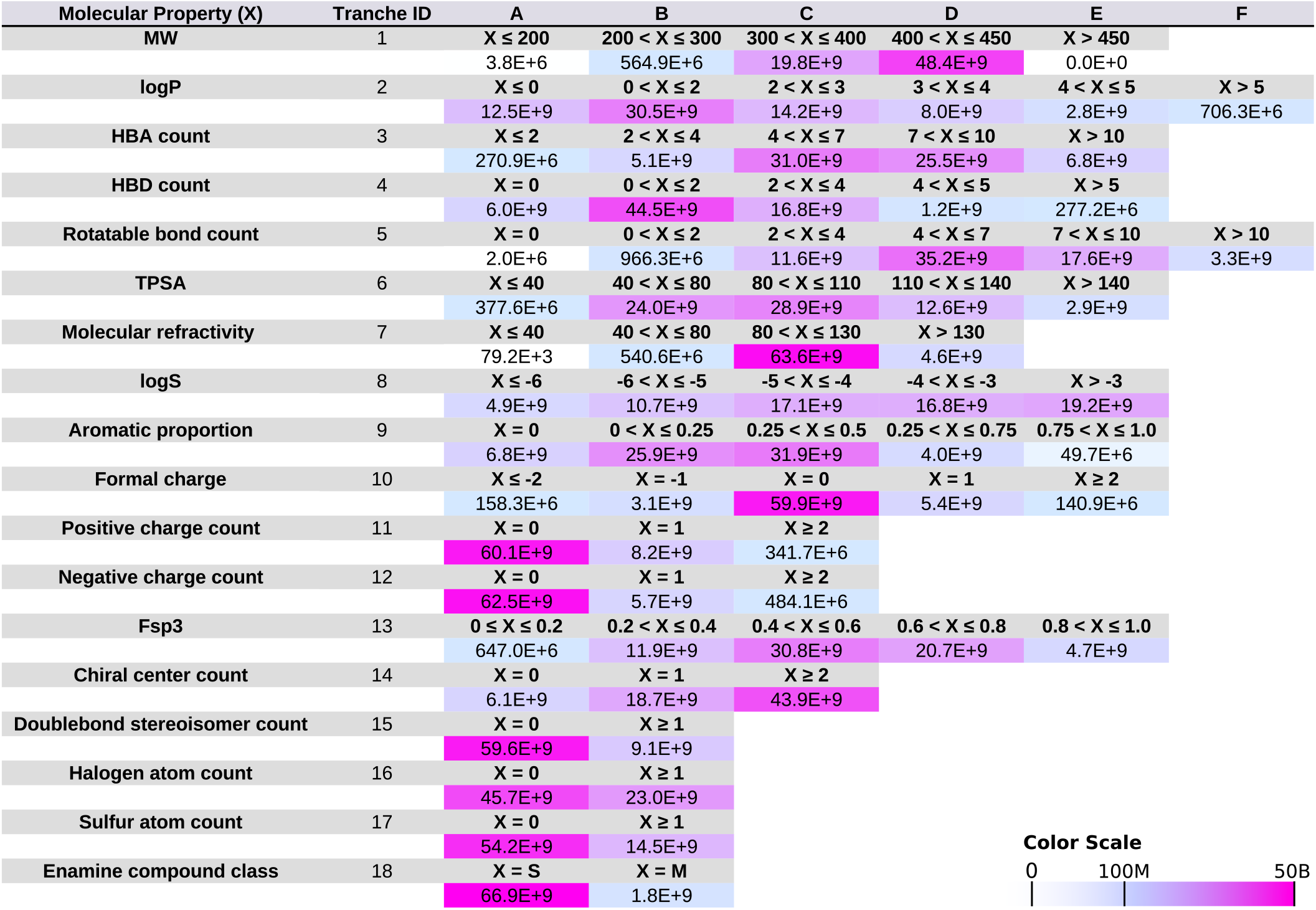
Overview of 18-dimensional tranche table. Molecular properties associated with each tranche, the partitions that are used for each tranche property, and the number of compounds per partition interval for each property (colored as a heat map).

**Extended Data Fig. 4.**
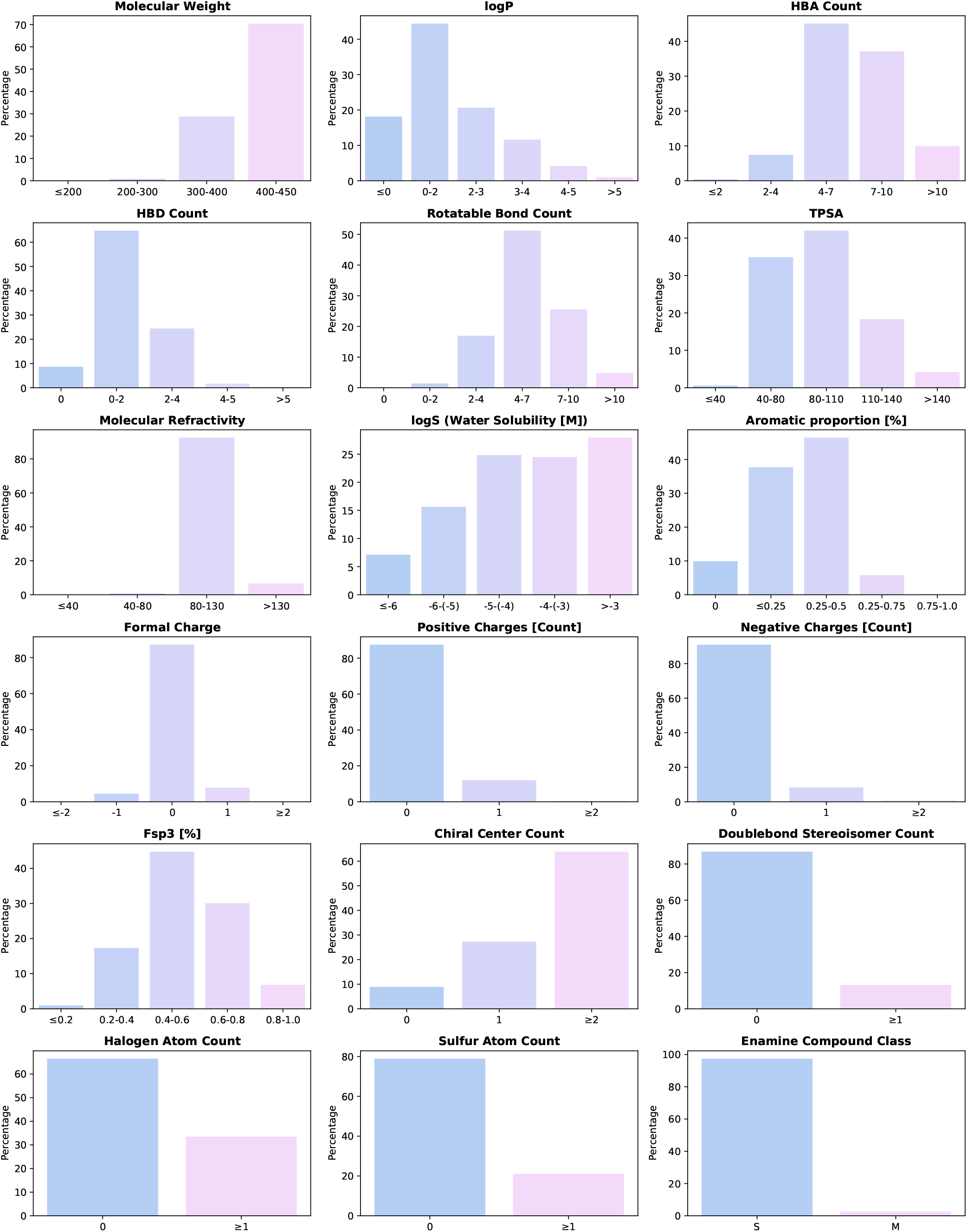
Overview of the distribution of the molecular properties of the Enamine REAL Space (version 2022q1-2). Each subplot shows the distribution of the molecular property associated with each dimension of the 18-dimensional grid into which the REAL Space was partitioned.

**Extended Data Fig. 5.**
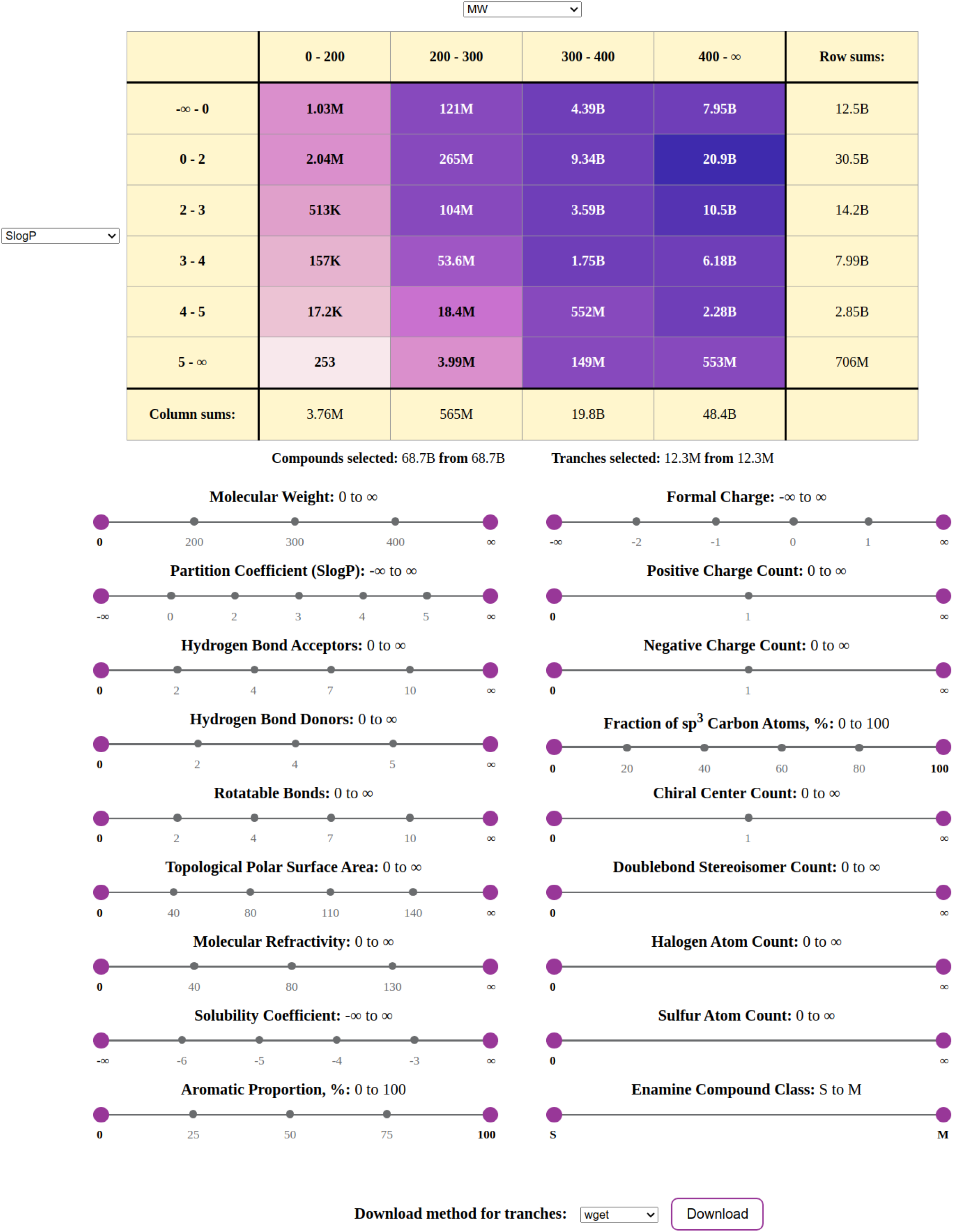
Webinterface (version 1) of the REAL Space. Shown is an interactive web interface that is made available on the AdaptiveFlow homepage (https://adaptive-flow.ai/real-space-2022q12-interface-2) that allows selecting subsets of the REAL Space partitioned into an 18-dimensional matrix. Each dimension corresponds to a molecular property and has a slider that can be adjusted to select the desired property ranges. The number of selected molecules is updated automatically in the web interface. Effectively, the sliders allow the selection of multidimensional rectangles within the 18-dimensional matrix.

**Extended Data Fig. 6.**
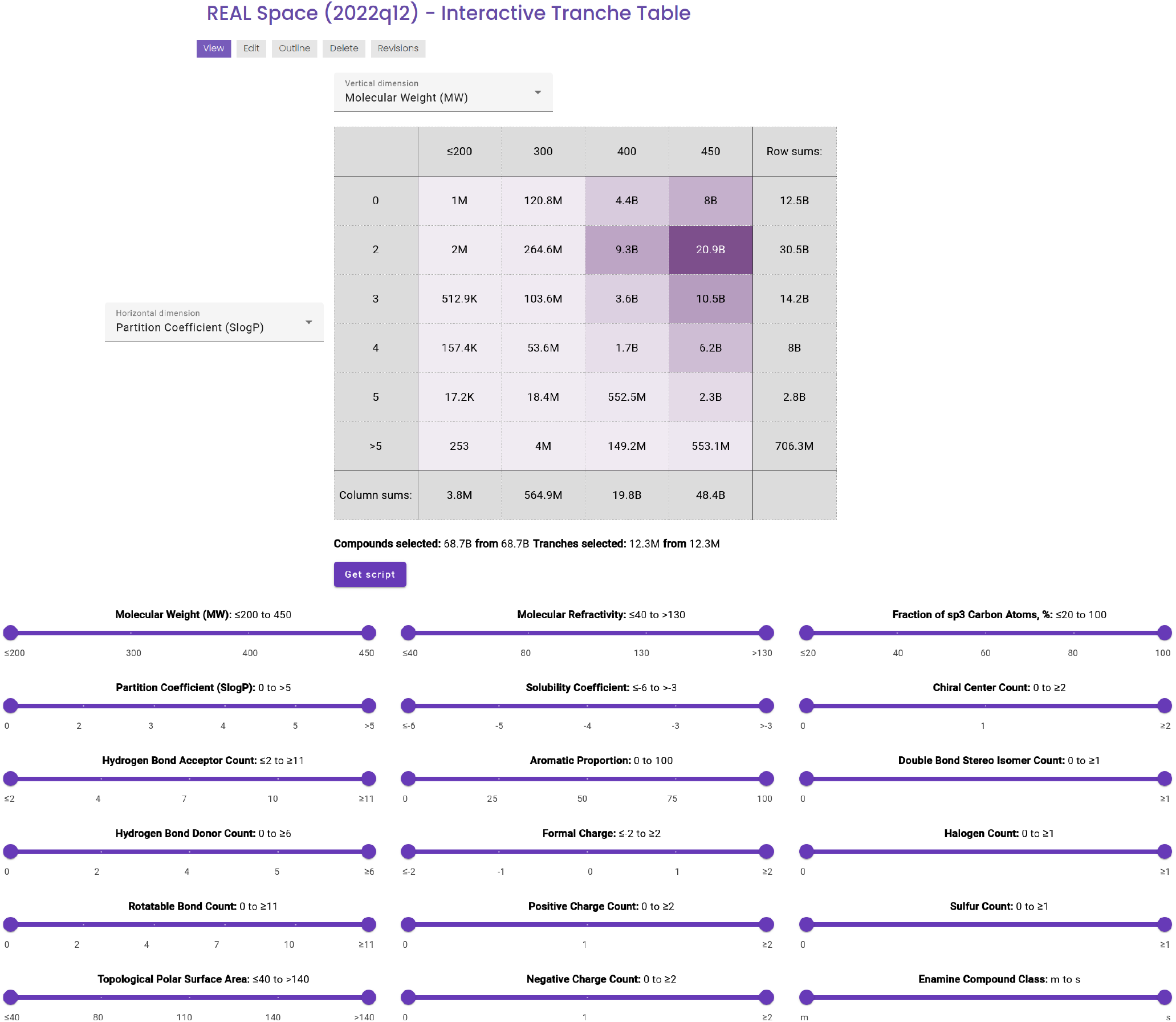
Webinterface (version 2) of the REAL Space. Shown is an interactive web interface that is made available on the AdaptiveFlow homepage (https://adaptive-flow.ai/real-space-2022q12-interactive-tranche-table) that allows selecting subsets of the REAL Space partitioned into an 18-dimensional matrix. Each dimension corresponds to a molecular property and has a slider that can be adjusted to select the desired property ranges. The number of selected molecules is updated automatically in the web interface. Effectively, the sliders allow the selection of multidimensional rectangles within the 18-dimensional matrix.

**Extended Data Fig. 7.**
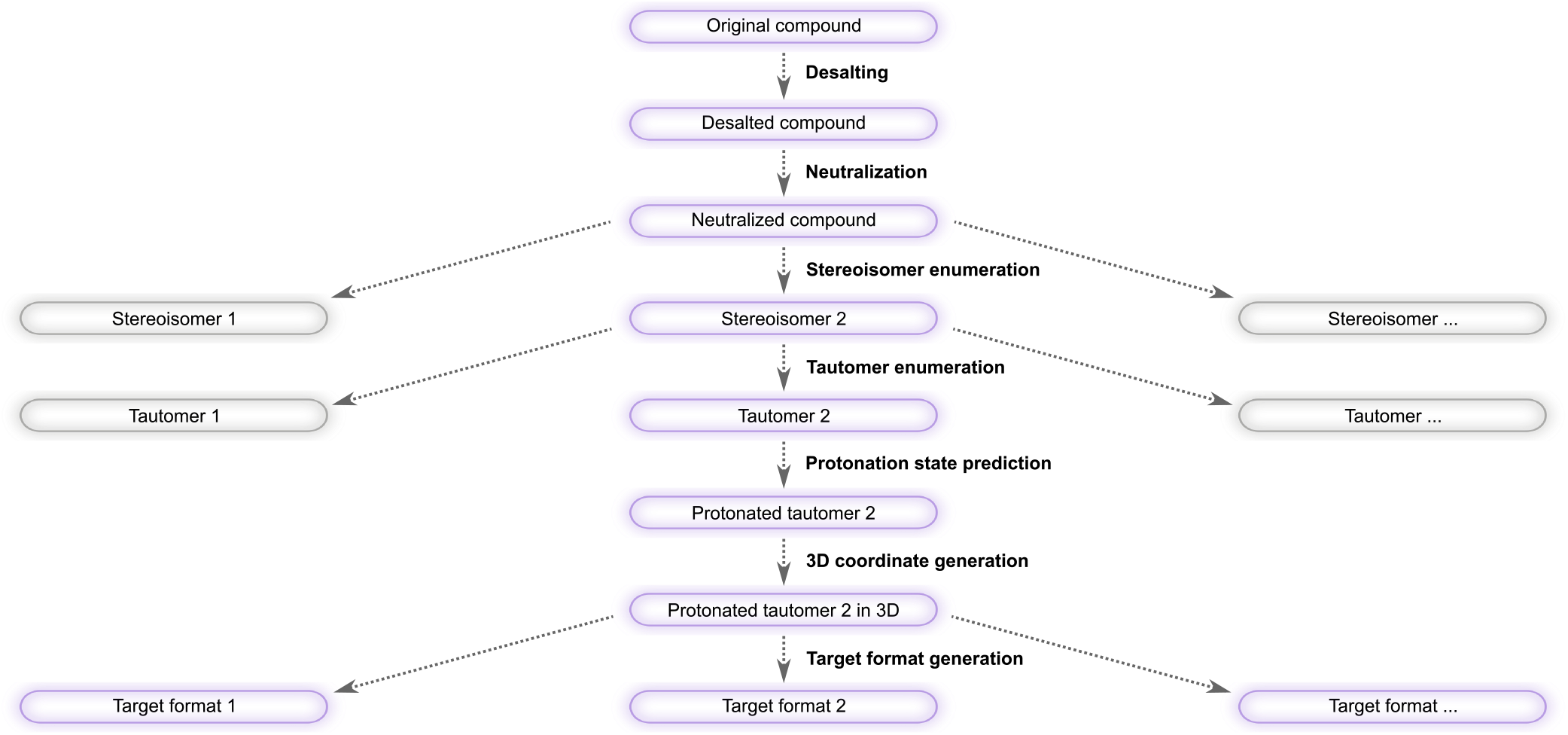
AFLP - Overview of the Ligand Preparation Procedure. Ligands are initially in a format in line notation (e.g. SMILES, SELFIES, or amino acid sequences). The preparation steps include desalting, neutralization, stereoisomer enumeration, tautomer enumeration, protonation state prediction, 3D coordinate generation, and target format generation. Stereoisomer and tautomer generation can result in multiple chemical species. Highlighted in purple are the steps of a molecule from the beginning to the end of the preparation procedure. Colored in white are the intermediate molecules/tautomers whose subsequent preparation steps are not shown in the diagram.

**Extended Data Fig. 8.**
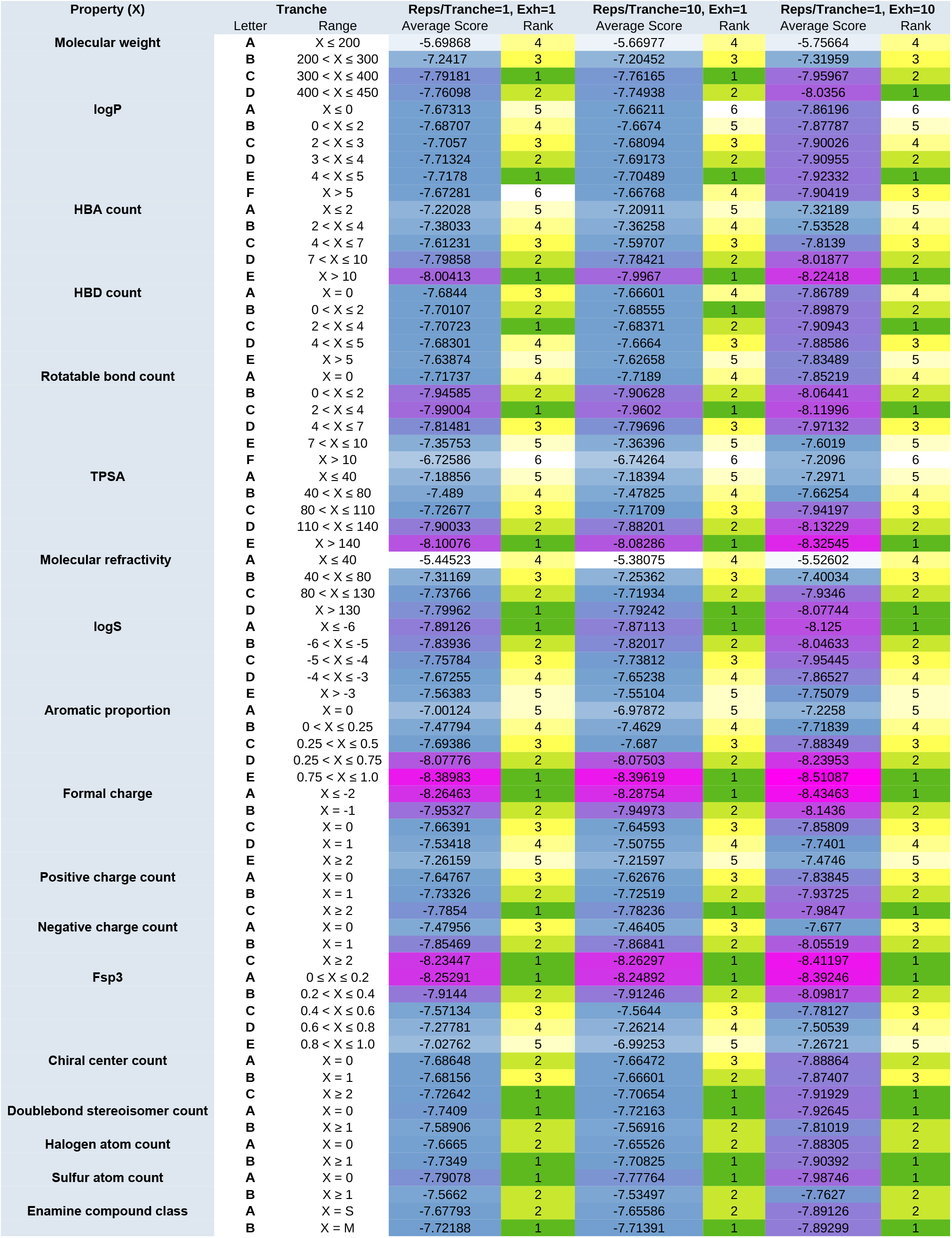
Adaptive Target-Guided Prescreening Benchmarks. Results of several prescreens for a test system within the Adaptive Target-Guided Virtual Screening (ATG-VS) approach, where different settings were used. In the first run (left), one representative per tranche was used with docking exhaustiveness 1. In the second run (middle), 10 representatives per tranche are used with exhaustiveness 1. In the third run (right), one representative per tranche was used with exhaustiveness 10. The average docking scores were computed for each tranche of each property (averaging over all other properties/dimensions). For each property, the tranches have been ranked within their property realm based on the average docking scores for the tranches. As can be seen, the patterns and rankings are very similar to each other, indicating that one representative with exhaustiveness 1 can be sufficient for the prescreenings.

**Extended Data Fig. 9.**
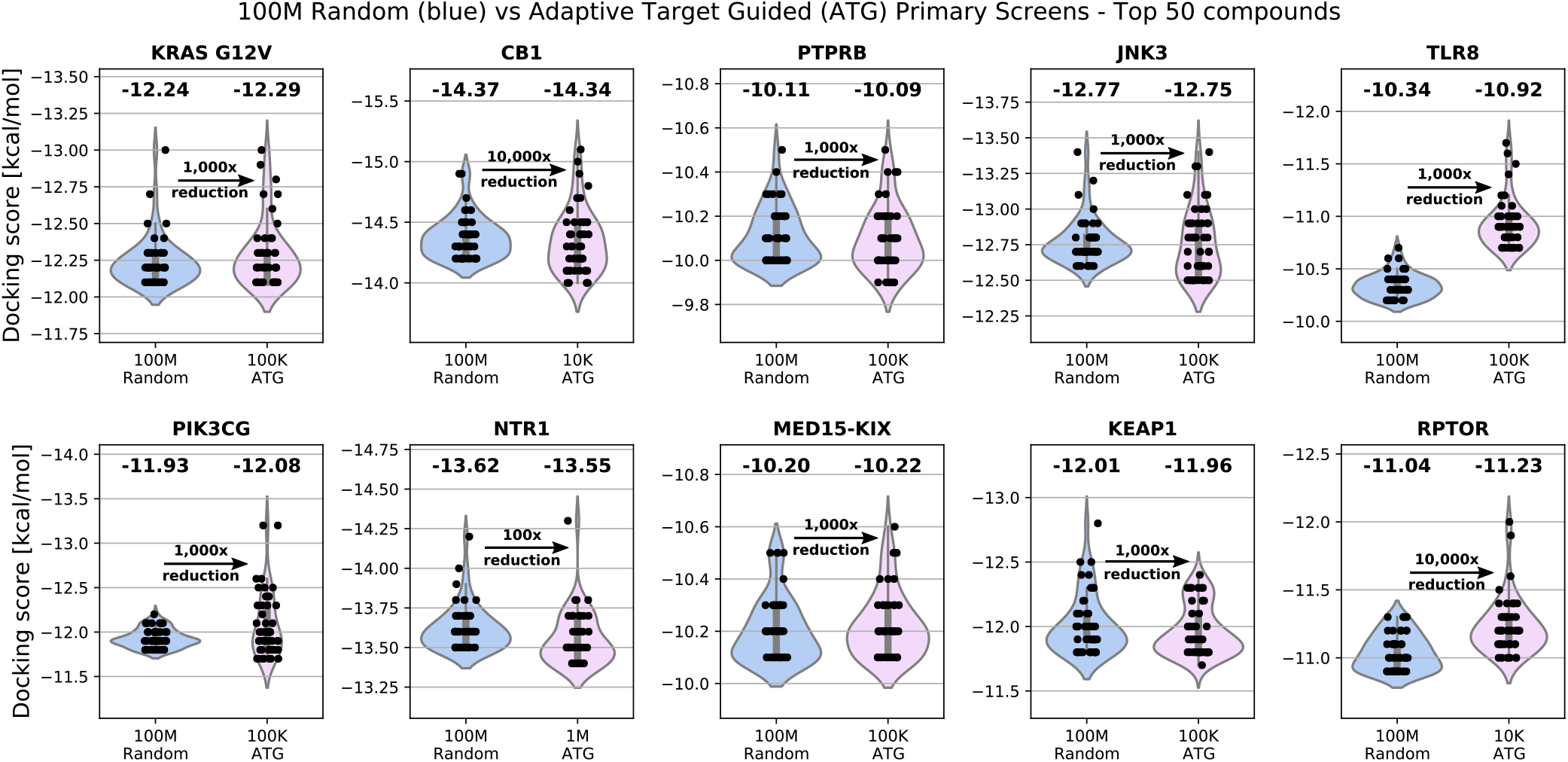
Benchmarking large-scale virtual screening strategies. Violin plots compare the distribution of docking scores for the top 50 molecules identified by two approaches: (1) a standard ultra-large virtual screen of 100 million randomly selected compounds, and (2) the Adaptive Target-Guided (ATG) screening strategy developed in this work. For each ATG run, a prescreen of 11 million representative compounds was used to identify the most promising regions of chemical space, followed by a focused primary screen whose size is indicated for each target. ATG consistently achieves comparable or superior docking scores using a fraction of the computational budget.

**Extended Data Fig. 10.**
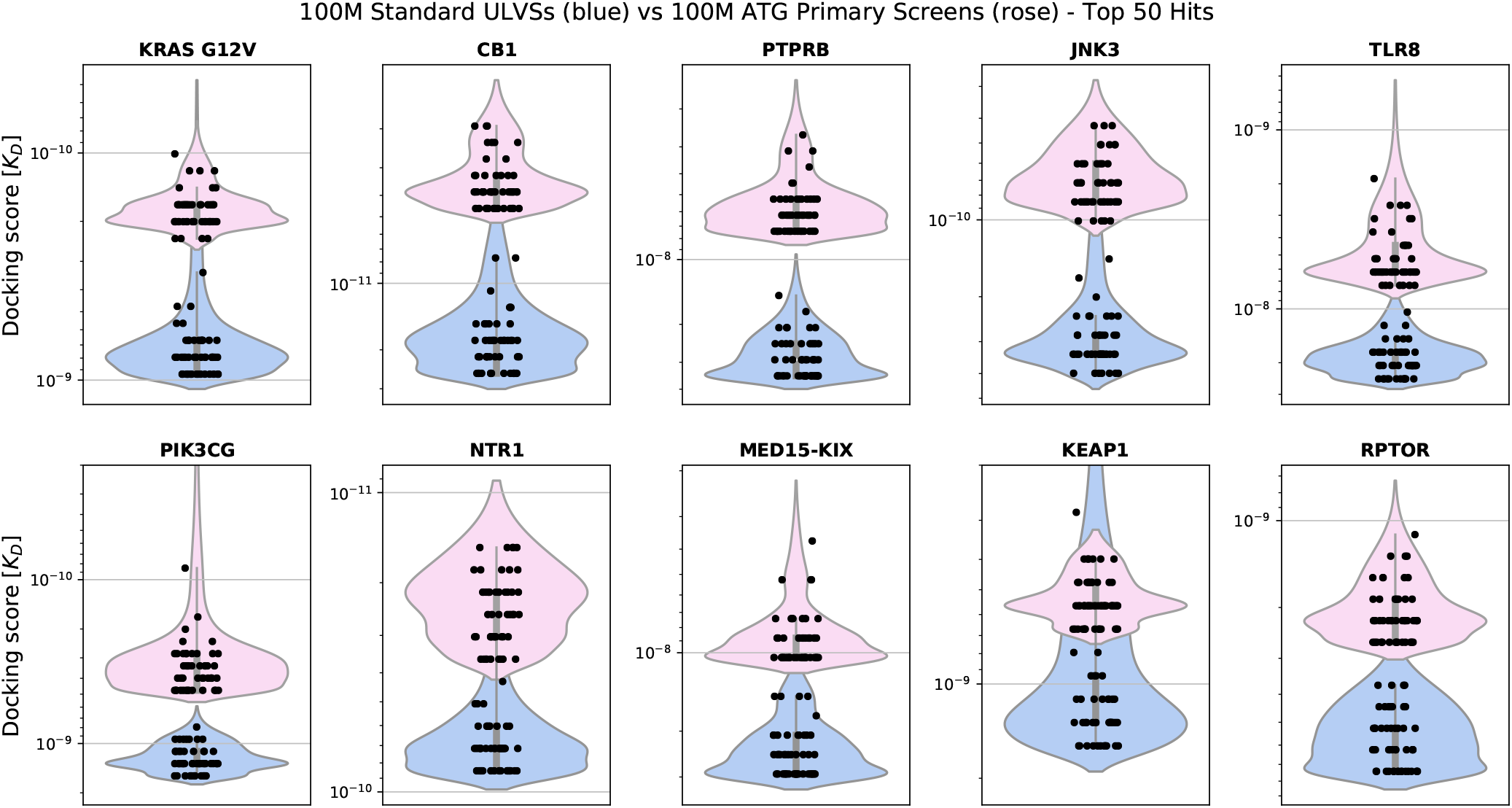
Comparison of Adaptive Target-Guided (ATG) Primary Virtual Screens and standard ULVS. 100 million compounds were screened with standard ULVS of randomly selected compounds, as well as with ATGs where the 100 million compounds to be screened were selected based on the ATG prescreen. Violin plots of the docking scores of the top 50 virtual screening hits obtained by standard ULVSs (blue) and ATG-VS (rose). Stripplots of the data points are overlayed. The docking scores of the top 50 compounds of the adaptive screens are considerably better for all target proteins than for the standard ULVSs.

**Extended Data Fig. 11.**
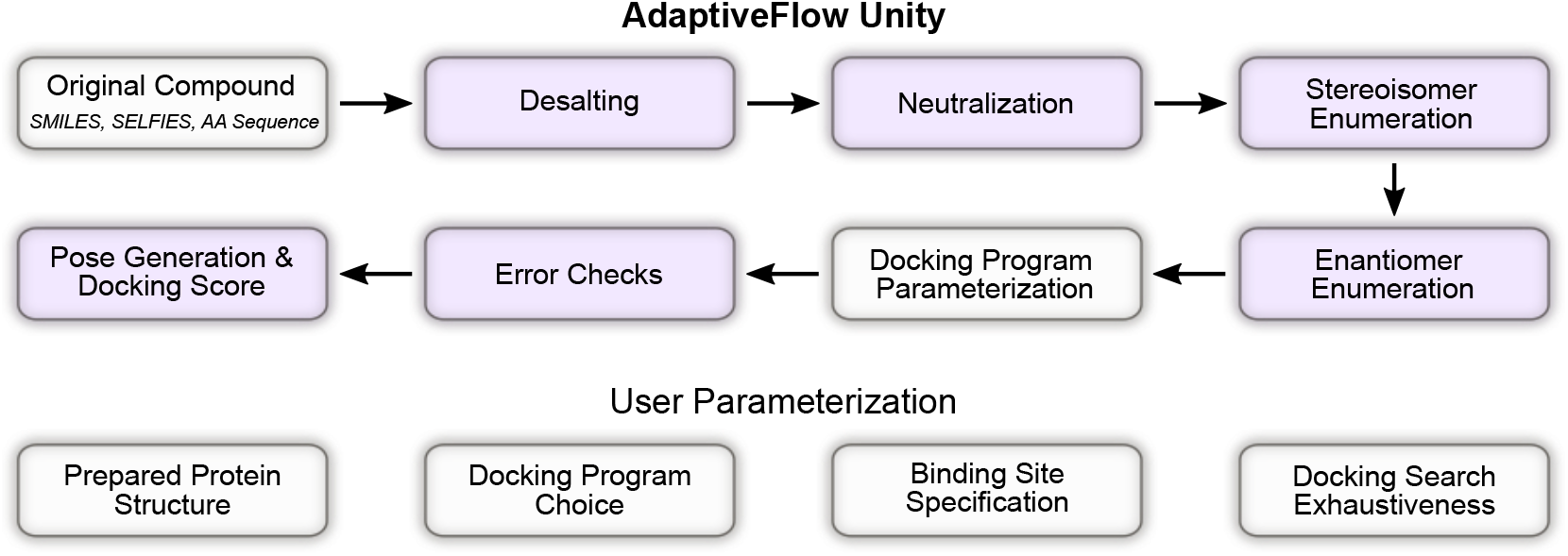
Conceptual Overview of AdaptiveFlow Unity (AFU). AFU takes as input a user-specified string representation of a molecule and processes the provided molecule by desalting, neutralizing, and generating stereoisomer (with RDKit (65)). Subsequently, based on the docking program choice, all ligands are converted into 3D conformers (with OpenBabel (63)) using a default protonation state of 7.4. Finally, built-in error checks are performed depending on a user’s docking program choice, returning the docking pose and scoring function value. Steps in which the parameters of the user are involved are colored in white.

**Extended Data Fig. 12.**
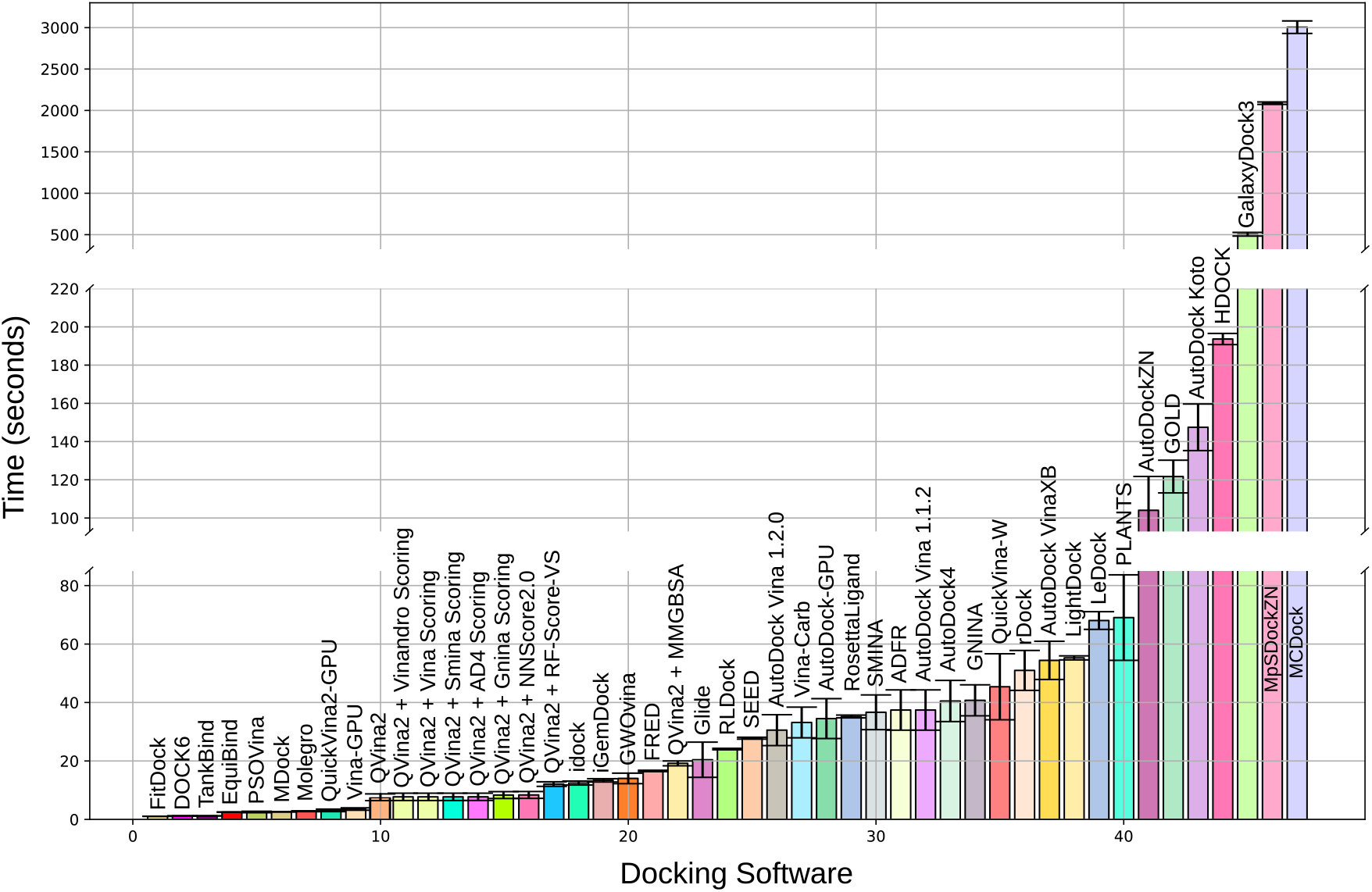
Docking Time Estimates. Timing estimates for running a single calculation averaged over 25 independent runs for all docking programs. The calculations were run using AFU.

**Extended Data Table 1.** Features Supported by AdaptiveFlow. Features grouped by AdaptiveFlow module: General, AFLP, AFVS, and AF Unity. AdaptiveFlow has significantly more features compared to the original version.

| Feature | AdaptiveFlow |
| --- | --- |
| **General Features** |  |
| Codebase | Python |
| Native cloud support | available (AWS) |
| Linear scaling behavior | demonstrated up to 5.6 million CPUs |
| **AdaptiveFlow Ligand Libraries** |  |
| Total size of ligand library | 68.7 billion |
| Additional predicted ligand properties | 28 |
| PDB format | available |
| PDBQT format | available |
| SDF format | available |
| MOL2 format | available |
| SMILES format | available |
| SELFIES format | available |
| Parquet format | available |
| Sparse version (for ATG-VSs) | available |
| Interactive web interface | yes (18 dimensions) |
| Accessibility | AdaptiveFlow website and AWS Open Data |
| **AdaptiveFlow for Ligand Preparation (AFLP) Module** |  |
| Stereoisomer enumeration | supported |
| External open-source tools for ligand preparation | extended support |
| SMILES as input format | supported |
| SELFIES as input format | supported |
| Amino acid sequences as input format | supported |
| SELFIES as output format | supported |
| Prediction of physical molecular properties | supported (27 properties) |
| Automatic multidimensional matrix generation | extended support |
| Quality control | extended support |
| Enhanced speed | extended support |
| **AdaptiveFlow for Virtual Screening (AFVS) Module** |  |
| Adaptive target-guided virtual screens | supported |
| - Machine learning enhancement | supported |
| docking protocols | 1500 |
| Deep learning-based scoring functions | supported |
| Deep learning-based pose prediction methods | supported |
| Molecular dynamics-based methods | supported |
| Protein as target | supported |
| RNA as target | supported |
| DNA as target | supported |
| Covalent docking | extended support |
| Protein ligands (protein-receptor docking) | supported |
| Peptide ligands | supported |
| Parquet output format | supported |
| Docking quality control | supported |
| ARM supported | supported |
| Enhanced x86 performance | available (for several docking protocols) |
| GPU support | supported |
| **AdaptiveFlow Unity (AFU) Module** |  |
| Standalone mode | available |
| API mode | available |

**Extended Data Table 2.** Calculated Properties of Ligands in the Enamine REAL Space. All properties of the ligands that were determined/calculated during the preparation of the REAL Space (version 2022q1-2) with AFVS. A few properties (molecular weight, fsp3, heavy atom count, Enamine compound class) were available in the input library and thus could be directly used without any calculations.

| Index | Property | AdaptiveFlow Parameter | Preparation Method | Used for 18-Dimensional Matrix |
| --- | --- | --- | --- | --- |
| 1 | molecular weight | mw_file | Input library | yes |
| 2 | logP | logp_jchem | JChem | yes |
| 3 | hydrogen bond acceptor count | hba_jchem | JChem | yes |
| 4 | hydrogen bond donor count | hbd_jchem | JChem | yes |
| 5 | rotatable bond count | rotb_jchem | JChem | yes |
| 6 | topological polar surface area | tpsa_jchem | JChem | yes |
| 7 | molecular refractivity | mr_jchem | JChem | yes |
| 8 | solubility [molar] | logs | JChem | yes |
| 9 | number of bonds | bondcount_jchem | JChem | no |
| 10 | aromatic proportion | ap | JChem | yes |
| 11 | ring count | ringcount | JChem | no |
| 12 | aromatic ring count | aromaticringcount | JChem | no |
| 13 | logD | logd | JChem | no |
| 14 | formal charge | formalcharge | AdaptiveFlow | yes |
| 15 | positive charge count | positivechargecount | AdaptiveFlow | yes |
| 16 | negative charge count | negativechargecount | AdaptiveFlow | yes |
| 17 | fraction of sp3 carbon atoms | fsp3_file | Input library | yes |
| 18 | total atom count | atomcount_jchem | JChem | no |
| 19 | heavy atom count | heavyatomcount_file | Input library | no |
| 20 | chiral center count | chiralcentercount | JChem | yes |
| 21 | doublebond stereoisomer count | doublebondstereoisomercount | JChem | yes |
| 22 | halogen atom count | halogencount | JChem | yes |
| 23 | sulfur atom count | sulfurcount | JChem | yes |
| 24 | nitrogen and oxygen atom count | NOcount | JChem | no |
| 25 | electronegative atom count | electronegativeatomcount | JChem | no |
| 26 | Enamine compound class | enamine_compound_class | Input library | yes |
| 27 | quantitative estimate of druglikeness (QED) | qed_rdkit | RDKit | no |
| 28 | Murcko scaffold | scaffold_rdkit | RDKit | no |

**Extended Data Table 3.** Supported Processing Steps of AFLP. AFLP can prepare molecules into a ready-to-dock format, and calculate additional properties of each molecule. For many of these calculations, external tools (JChem from ChemAxon, RDKit, Epik from Schrödinger (98), QupKake (99) and Open Babel) are used, while for a few AFLP calculates them internally. Many properties can also be used directly from the input library if available, avoiding the need to calculate them during the runtime.

| Property | Available Calculation Methods |
| --- | --- |
| **Preparation of Ligands into Ready-To-Dock Format** |  |
| Desalting | AFLP (internal) |
| Neutralization | JChem, Open Babel |
| Stereoisomer Generation | JChem, RDKit |
| Tautomer Generation | JChem, Open Babel, RDKit |
| Protonation State Calculation | JChem, Open Babel, Epik, QupKake |
| 3D Coordinate Conversion | JChem, Open Babel, RDKit |
| Target Format Conversion | Open Babel (for all formats but SELFIES)<br>and SELFIES Python Package (only for SELFIES) |
| **Calculation of Additional Molecular Properties** |  |
| Molecular weight | JChem, Open Babel, Input Library |
| logP | JChem, Open Babel, Input Library |
| HBA count | JChem, Open Babel, Input Library |
| HBD count | JChem, Open Babel, Input Library |
| Rotatable bond count | JChem, Open Babel, Input Library |
| TPSA | JChem, Open Babel, Input Library |
| Molecular refractivity | JChem, Open Babel, Input Library |
| Solubility [molar] | JChem, Open Babel, Input Library |
| Bond count | JChem, Input Library |
| Aromatic proportion | JChem, Input Library |
| Ring count | JChem, Input Library |
| Aromatic ring count | JChem, Input Library |
| logD | JChem, Input Library |
| Formal charge | AFLP (internal) |
| Positive charge count | AFLP (internal) |
| Negative charge count | AFLP (internal) |
| Fraction of sp<sup>3</sup> carbon atoms (fsp<sup>3</sup>) | JChem, Input Library |
| Total atom count | JChem, Open Babel |
| Heavy atom count | Input Library |
| Chiral center count | JChem, Input Library |
| Doublebond stereoisomer count | JChem |
| Halogen atom count | JChem, Input Library |
| Sulfur atom count | JChem, Input Library |
| NO atom count | JChem, Input Library |
| Electronegative atom count | JChem, Input Library |
| Enamine compound class | Input Library |
| Quantitative Estimate of Druglikeness (QED) | RDKit |
| Murcko scaffold | RDKit |

**Extended Data Table 4.** Supported docking programs. The collection of unique underlying docking programs supported within AFVS and AF Unity. These programs serve as the foundational components for the 1,500 supported docking protocols (specific combinations of pose prediction and scoring algorithms) enabled by the platform.

| Docking program | License model | Special features | DL-based | Hardware support | Reference |
| --- | --- | --- | --- | --- | --- |
| AutoDock-GPU | Open Source | GPU-accelerated version of AutoDock4 | ✗ | GPU | (59) |
| AutoDock4 | Open Source | Improved modelling of waters and macrocycles | ✗ | CPU | (100) |
| AutoDockZN | Open Source | Special support for zinc-coordinating ligands | ✗ | CPU | (101) |
| AutoDock Vina 1.2.0 | Open Source | Python support; macromolecule support | ✗ | CPU | (102) |
| AutoDock Vina 1.1.2 | Open Source | Improved optimization, scoring and multithreading | ✗ | CPU | (103) |
| AutoDockFR (ADFR) | Open Source | Improved receptor side-chain flexibility modelling | ✗ | CPU | (104) |
| AutoDock Koto | Open Source | Improved sampling | ✗ | CPU | (105) |
| ATPdock | Open Source | ATP-specific protein-ligand docking | ✗ | CPU | (106) |
| CovDock | Licence Required | Covalent docking | ✗ | CPU | (107) |
| DOCK 6.0 | Free for academics | RNA-ligand, DNA-ligand, protein-protein docking | ✗ | CPU | (108) |
| FlexX | Licence Required | Covalent docking. Ensemble docking | ✗ | CPU | (109) |
| FitDock | Academic Licence | Incorporation of template fitting for docking | ✗ | CPU | (110) |
| GalaxyDock3 | Licence Required | Incorporation of side chain flexibility | ✗ | CPU | (111) |
| Glide (HTVS) | Paid Academic Licencing | Less exhaustive sampling and scoring | ✗ | CPU | (112) |
| Glide (SP) | Paid Academic Licencing | More exhaustive sampling (relative to Glide HTVS) & multiple levels of accuracy | ✗ | CPU | (112) |
| Glide (XP) | Paid Academic Licencing | Most accurate sampling and scoring for Glide & different levels of accuracy | ✗ | CPU | (112) |
| Gnina | Open Source | Deep learning scoring of complex | ✓ | GPU/CPU scoring | (70) |
| GOLD | Licence Required | Genetic algorithm based docking; Covalent docking | ✗ | CPU | (113) |
| GWOVina | Open Source | Grey wolf optimization | ✗ | CPU | (114) |
| HDOCK | Academic Licence | Protein-RNA/DNA docking | ✗ | CPU | (115) |
| HSYMDOCK | Academic Licence | Protein-protein/rna docking with Cn and Dn symmetry | ✗ | CPU | (116) |
| idock | Open Source | AutoDock Vina derivative with improved speed | ✗ | CPU | (117) |
| iGEMDOCK | Open Source | - | ✗ | CPU | (118) |
| LeDock | Freeware for academic use | Excellent Sampling power (comparable to GOLD) | ✗ | CPU | (119) |
| LigandFit | Open Source | Identification of active site regions for docking | ✗ | CPU | (120) |
| LightDock | Open Source | Protein-protein docking | ✗ | CPU | (121) |
| MDock | Licence Required | Ensemble docking for multiple protein conformations | ✗ | CPU | (122) |
| MCDock | Open Source | Use of Monte Carlo simulation technique | ✗ | CPU | (123) |
| Molegro | Licence Required | - | ✗ | CPU | (124) |
| MpsDockzn | Licence Required | Zinc metalloprotein-specific scoring function | ✗ | CPU | (125) |
| PSOVina | Open Source | Improved pose sampling and speed | ✗ | CPU | (126) |
| PLANTS | Open Source | Use of ant colony optimization | ✗ | CPU | (127) |
| PI-LZerD | Open Source | Protein-protein docking | ✗ | CPU | (128) |
| PIPER | Open Source | Use of pairwise interaction potentials for protein-protein docking | ✗ | CPU | (129) |
| QuickVina 2 | Open Source | Extremely fast | ✗ | CPU | (92) |
| QuickVina-W | Open Source | Improved speed for blind docking | ✗ | CPU | (130) |
| QVina2-GPU | Open Source | GPU-accelerated version of QVina2 | ✗ | GPU | (60) |
| QVina2-W-GPU | Open Source | GPU-accelerated version of QVina2-W | ✗ | GPU | (60) |
| rDock | Open Source | Support for RNA targets | ✗ | CPU | (131) |
| Rosetta Ligand | Licence Required | - | ✗ | CPU | (132) |
| RLDOCK | Open Source | RNA-ligand docking | ✗ | CPU | (133) |
| SEED | Open Source | Fragment-based docking | ✗ | CPU | (134) |
| SMINA | Open Source | User specific scoring functions | ✗ | CPU | (135) |
| TANKBind | Open Source | Fast neural network based complex prediction | ✓ | CPU/GPU | (12) |
| VinaCarb | Open Source | Improved glycosidic modelling | ✗ | CPU | (136) |
| VinaXB16 | Open Source | Improved halogen bonding scoring function | ✗ | CPU | (137) |
| Vina-GPU | Open Source | GPU-accelerated version of Vina | ✗ | GPU | (60) |
| Vina-GPU-2.0 | Open Source | GPU-accelerated version of Vina-2.0 | ✗ | GPU | (60) |

**Extended Data Table 5.**
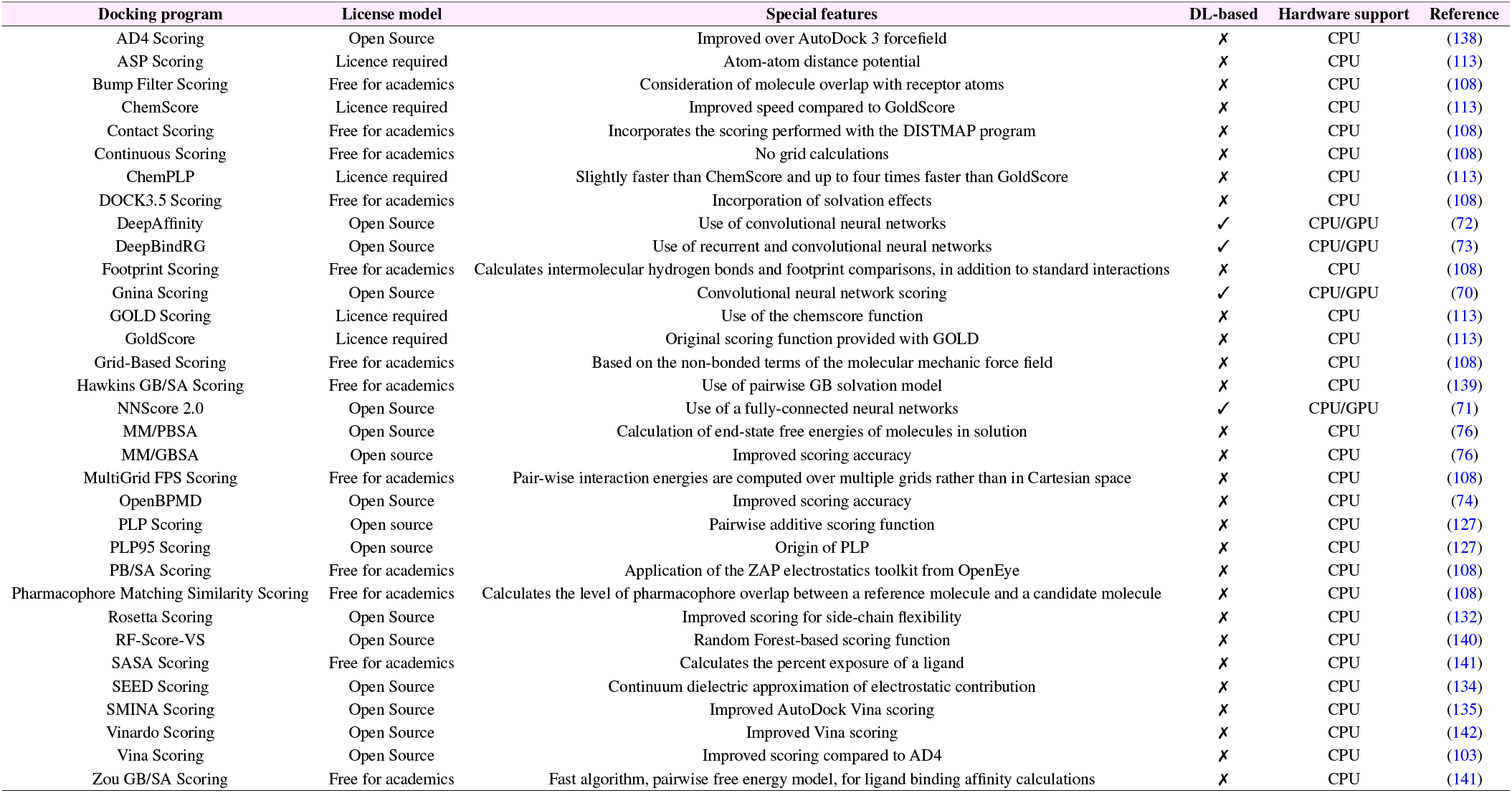
Supported Protein-Ligand Scoring Functions. The collection of docking programs supported within AFVS and AF Unity that perform both pose prediction and scoring.

**Extended Data Table 6.** Supported pose prediction methods. The collection of docking programs supported within AFVS and AF Unity that perform both pose prediction and scoring.

| Docking program | License model | Special features | DL-based | Hardware support | Reference |
| --- | --- | --- | --- | --- | --- |
| DiffDock | Open Source | Blind docking pose prediction | ✓ | GPU/CPU | (68) |
| EquiBind | Open Source | Blind docking pose prediction | ✓ | GPU/CPU | (69) |
| EnzyDock | Open Source | Identifies probable binding modes of multiple ligand states in an enzyme active site | ✗ | CPU | (143) |
| FRED | Academic Licence | Improved pose sampling | ✗ | CPU | (144) |

**Extended Data Table 7.** ATG-VS Benchmark Systems. The target proteins and binding sites used in the benchmark studies comparing ATG-VSs with standard ULVSs.

| Target Protein | Structure ID | Structure Source | Target Site | Target Site Type |
| --- | --- | --- | --- | --- |
| KRAS<sup>G12V</sup> | 4TQ9 | Protein Data Bank (RCSB) | Active site | Enzymatic site |
| CB1 | 6KPG | Protein Data Bank (RCSB) | Primary ligand pocket | Orthosteric site |
| PTPRB | 2H04 | Protein Data Bank (RCSB) | Active site | Enzymatic site |
| JNK3 | 7S1N | Protein Data Bank (RCSB) | Active site | Enzymatic site |
| TLR8 | 6ZJZ | Protein Data Bank (RCSB) | Dimerization site | Protein-protein interaction |
| PIK3CG | 2CHZ | Protein Data Bank (RCSB) | Active site | Enzymatic site |
| NTR1 | 4GRV | Protein Data Bank (RCSB) | Primary ligand pocket | Orthosteric site |
| MED15-KIX | 2GUT | Protein Data Bank (RCSB) | SREBP interaction site | Protein-protein interaction |
| KEAP1 | 5FNQ | Protein Data Bank (RCSB) | NRF2 interaction site | Protein-protein interaction |
| Raptor | AFQ8N122_F1 | AlphaFold Structure Database | TOS motif | Protein-protein interaction |

