## Supplementary Information for "AI-Enhanced Adaptive Virtual Screening Platform Enabling Exploration of 69 Billion Molecules Discovers Structurally Validated FSP1 Inhibitors"

#### A Available docking protocols

AFVS and AFU support the same docking protocols. docking protocols consist of two key components, a pose prediction component (or sampling algorithm) and a scoring function. Supported docking protocols in AdaptiveFlow are either standalone docking programs that contain both a scoring function and a sampling algorithm or composite docking programs that are assembled by AdaptiveFlow from individual scoring functions and pose prediction methods. A list of all docking protocols currently available in AdaptiveFlow is contained in Supplementary Table 1.

| Index | Pose Prediction/Sampling Method | Scoring Function | DL-Based Sampling | DL-Based Scoring |
|---|---|---|---|---|
| 0 | AutoDock-GPU | ASP Scoring | ✗ | ✗ |
| 1 | AutoDock-GPU | AutoDock-GPU | ✗ | ✗ |
| 2 | AutoDock-GPU | Bump Filter Scoring | ✗ | ✗ |
| 3 | AutoDock-GPU | ChemPLP | ✗ | ✗ |
| 4 | AutoDock-GPU | ChemScore | ✗ | ✗ |
| 5 | AutoDock-GPU | Contact Scoring | ✗ | ✗ |
| 6 | AutoDock-GPU | Continuous Scoring | ✗ | ✗ |
| 7 | AutoDock-GPU | DOCK 3.5 Scoring | ✗ | ✗ |
| 8 | AutoDock-GPU | DeepAffinity | ✗ | ✓ |
| 9 | AutoDock-GPU | DeepBindRG | ✗ | ✓ |
| 10 | AutoDock-GPU | Footprint Scoring | ✗ | ✗ |
| 11 | AutoDock-GPU | GOLD Scoring | ✗ | ✗ |
| 12 | AutoDock-GPU | Gnina Scoring | ✗ | ✓ |
| 13 | AutoDock-GPU | GoldScore | ✗ | ✗ |
| 14 | AutoDock-GPU | Grid-Based Scoring | ✗ | ✗ |
| 15 | AutoDock-GPU | Hawkins GB/SA Scoring | ✗ | ✗ |
| 16 | AutoDock-GPU | MM/GBSA Scoring | ✗ | ✗ |
| 17 | AutoDock-GPU | MM/PBSA Scoring | ✗ | ✗ |
| 18 | AutoDock-GPU | MultiGrid FPS Scoring | ✗ | ✗ |
| 19 | AutoDock-GPU | NNScore 2.0 | ✗ | ✓ |
| 20 | AutoDock-GPU | OpenBPMD | ✗ | ✗ |
| 21 | AutoDock-GPU | PB/SA Scoring | ✗ | ✗ |
| 22 | AutoDock-GPU | PLANTS Scoring | ✗ | ✗ |
| 23 | AutoDock-GPU | PLP | ✗ | ✗ |
| 24 | AutoDock-GPU | PLP95 | ✗ | ✗ |
| 25 | AutoDock-GPU | Pharmacophore Matching Similarity Scoring | ✗ | ✗ |
| 26 | AutoDock-GPU | RF-Score-VS | ✗ | ✗ |
| 27 | AutoDock-GPU | Rosetta Scoring | ✗ | ✗ |
| 28 | AutoDock-GPU | SASA Scoring | ✗ | ✗ |
| 29 | AutoDock-GPU | SEED Scoring | ✗ | ✗ |
| 30 | AutoDock-GPU | SMINA Scoring | ✗ | ✗ |
| 31 | AutoDock-GPU | Vina scoring | ✗ | ✗ |
| 32 | AutoDock-GPU | Vinardo Scoring | ✗ | ✗ |
| 33 | AutoDock-GPU | Zou GB/SA Scoring | ✗ | ✗ |
| 34 | AutoDock4 | ASP Scoring | ✗ | ✗ |
| 35 | AutoDock4 | AutoDock4 | ✗ | ✗ |
| 36 | AutoDock4 | Bump Filter Scoring | ✗ | ✗ |
| 37 | AutoDock4 | ChemPLP | ✗ | ✗ |
| 38 | AutoDock4 | ChemScore | ✗ | ✗ |
| 39 | AutoDock4 | Contact Scoring | ✗ | ✗ |
| 40 | AutoDock4 | Continuous Scoring | ✗ | ✗ |
| 41 | AutoDock4 | DOCK 3.5 Scoring | ✗ | ✗ |
| 42 | AutoDock4 | DeepAffinity | ✗ | ✓ |
| 43 | AutoDock4 | DeepBindRG | ✗ | ✓ |
| 44 | AutoDock4 | Footprint Scoring | ✗ | ✗ |
| 45 | AutoDock4 | GOLD Scoring | ✗ | ✗ |
| 46 | AutoDock4 | Gnina Scoring | ✗ | ✓ |
| 47 | AutoDock4 | GoldScore | ✗ | ✗ |
| 48 | AutoDock4 | Grid-Based Scoring | ✗ | ✗ |
| 49 | AutoDock4 | Hawkins GB/SA Scoring | ✗ | ✗ |
| 50 | AutoDock4 | MM/GBSA Scoring | ✗ | ✗ |
| 51 | AutoDock4 | MM/PBSA Scoring | ✗ | ✗ |
| 52 | AutoDock4 | MultiGrid FPS Scoring | ✗ | ✗ |
| 53 | AutoDock4 | NNScore 2.0 | ✗ | ✓ |
| 54 | AutoDock4 | OpenBPMD | ✗ | ✗ |

**Supplementary Table 1. docking protocols supported by the AFVS and AFU Modules.** List of Supported docking protocols. Both the pose prediction/Sampling Method, as well as the scoring function, are listed. For both the sampling method and the scoring function, it is indicated whether they are based on deep learning (DL) or not (continued).

| Index | Pose Prediction/Sampling Method | Scoring Function | DL-Based Sampling | DL-Based Scoring |
|---|---|---|---|---|
| 55 | AutoDock4 | PB/SA Scoring | ✗ | ✗ |
| 56 | AutoDock4 | PLANTS Scoring | ✗ | ✗ |
| 57 | AutoDock4 | PLP | ✗ | ✗ |
| 58 | AutoDock4 | PLP95 | ✗ | ✗ |
| 59 | AutoDock4 | Pharmacophore Matching Similarity Scoring | ✗ | ✗ |
| 60 | AutoDock4 | RF-Score-VS | ✗ | ✗ |
| 61 | AutoDock4 | Rosetta Scoring | ✗ | ✗ |
| 62 | AutoDock4 | SASA Scoring | ✗ | ✗ |
| 63 | AutoDock4 | SEED Scoring | ✗ | ✗ |
| 64 | AutoDock4 | SMINA Scoring | ✗ | ✗ |
| 65 | AutoDock4 | Vina scoring | ✗ | ✗ |
| 66 | AutoDock4 | Vinardo Scoring | ✗ | ✗ |
| 67 | AutoDock4 | Zou GB/SA Scoring | ✗ | ✗ |
| 68 | AutoDockZN | AD4 Scoring | ✗ | ✗ |
| 69 | AutoDockZN | ASP Scoring | ✗ | ✗ |
| 70 | AutoDockZN | AutoDockZN | ✗ | ✗ |
| 71 | AutoDockZN | Bump Filter Scoring | ✗ | ✗ |
| 72 | AutoDockZN | ChemPLP | ✗ | ✗ |
| 73 | AutoDockZN | ChemScore | ✗ | ✗ |
| 74 | AutoDockZN | Contact Scoring | ✗ | ✗ |
| 75 | AutoDockZN | Continuous Scoring | ✗ | ✗ |
| 76 | AutoDockZN | DOCK 3.5 Scoring | ✗ | ✗ |
| 77 | AutoDockZN | DeepAffinity | ✗ | ✓ |
| 78 | AutoDockZN | DeepBindRG | ✗ | ✓ |
| 79 | AutoDockZN | Footprint Scoring | ✗ | ✗ |
| 80 | AutoDockZN | GOLD Scoring | ✗ | ✗ |
| 81 | AutoDockZN | Gnina Scoring | ✗ | ✓ |
| 82 | AutoDockZN | GoldScore | ✗ | ✗ |
| 83 | AutoDockZN | Grid-Based Scoring | ✗ | ✗ |
| 84 | AutoDockZN | Hawkins GB/SA Scoring | ✗ | ✗ |
| 85 | AutoDockZN | MM/GBSA Scoring | ✗ | ✗ |
| 86 | AutoDockZN | MM/PBSA Scoring | ✗ | ✗ |
| 87 | AutoDockZN | MultiGrid FPS Scoring | ✗ | ✗ |
| 88 | AutoDockZN | NNScore 2.0 | ✗ | ✓ |
| 89 | AutoDockZN | OpenBPMD | ✗ | ✗ |
| 90 | AutoDockZN | PB/SA Scoring | ✗ | ✗ |
| 91 | AutoDockZN | PLANTS Scoring | ✗ | ✗ |
| 92 | AutoDockZN | PLP | ✗ | ✗ |
| 93 | AutoDockZN | PLP95 | ✗ | ✗ |
| 94 | AutoDockZN | Pharmacophore Matching Similarity Scoring | ✗ | ✗ |
| 95 | AutoDockZN | RF-Score-VS | ✗ | ✗ |
| 96 | AutoDockZN | Rosetta Scoring | ✗ | ✗ |
| 97 | AutoDockZN | SASA Scoring | ✗ | ✗ |
| 98 | AutoDockZN | SEED Scoring | ✗ | ✗ |
| 99 | AutoDockZN | SMINA Scoring | ✗ | ✗ |
| 100 | AutoDockZN | Vina scoring | ✗ | ✗ |
| 101 | AutoDockZN | Vinardo Scoring | ✗ | ✗ |
| 102 | AutoDockZN | Zou GB/SA Scoring | ✗ | ✗ |
| 103 | AutoDock Vina 1.2.0 | AD4 Scoring | ✗ | ✗ |
| 104 | AutoDock Vina 1.2.0 | ASP Scoring | ✗ | ✗ |
| 105 | AutoDock Vina 1.2.0 | AutoDock Vina 1.2.0 | ✗ | ✗ |
| 106 | AutoDock Vina 1.2.0 | Bump Filter Scoring | ✗ | ✗ |
| 107 | AutoDock Vina 1.2.0 | ChemPLP | ✗ | ✗ |
| 108 | AutoDock Vina 1.2.0 | ChemScore | ✗ | ✗ |
| 109 | AutoDock Vina 1.2.0 | Contact Scoring | ✗ | ✗ |

**Supplementary Table 1 (Continued). docking protocols Supported by the AFVS and AFU Modules.** Each docking protocol is determined by the combination of a pose prediction method and a scoring function. For both of these components, it is indicated whether they are based on deep learning (DL) or not. (Continued on next page.)

| Index | Pose Prediction/Sampling Method | Scoring Function | DL-Based Sampling | DL-Based Scoring |
|---|---|---|---|---|
| 110 | AutoDock Vina 1.2.0 | Continuous Scoring | ✗ | ✗ |
| 111 | AutoDock Vina 1.2.0 | DOCK 3.5 Scoring | ✗ | ✗ |
| 112 | AutoDock Vina 1.2.0 | DeepAffinity | ✗ | ✓ |
| 113 | AutoDock Vina 1.2.0 | DeepBindRG | ✗ | ✓ |
| 114 | AutoDock Vina 1.2.0 | Footprint Scoring | ✗ | ✗ |
| 115 | AutoDock Vina 1.2.0 | GOLD Scoring | ✗ | ✗ |
| 116 | AutoDock Vina 1.2.0 | Gnina Scoring | ✗ | ✓ |
| 117 | AutoDock Vina 1.2.0 | GoldScore | ✗ | ✗ |
| 118 | AutoDock Vina 1.2.0 | Grid-Based Scoring | ✗ | ✗ |
| 119 | AutoDock Vina 1.2.0 | Hawkins GB/SA Scoring | ✗ | ✗ |
| 120 | AutoDock Vina 1.2.0 | MM/GBSA Scoring | ✗ | ✗ |
| 121 | AutoDock Vina 1.2.0 | MM/PBSA Scoring | ✗ | ✗ |
| 122 | AutoDock Vina 1.2.0 | MultiGrid FPS Scoring | ✗ | ✗ |
| 123 | AutoDock Vina 1.2.0 | NNScore 2.0 | ✗ | ✓ |
| 124 | AutoDock Vina 1.2.0 | OpenBPMD | ✗ | ✗ |
| 125 | AutoDock Vina 1.2.0 | PB/SA Scoring | ✗ | ✗ |
| 126 | AutoDock Vina 1.2.0 | PLANTS Scoring | ✗ | ✗ |
| 127 | AutoDock Vina 1.2.0 | PLP | ✗ | ✗ |
| 128 | AutoDock Vina 1.2.0 | PLP95 | ✗ | ✗ |
| 129 | AutoDock Vina 1.2.0 | Pharmacophore Matching Similarity Scoring | ✗ | ✗ |
| 130 | AutoDock Vina 1.2.0 | RF-Score-VS | ✗ | ✗ |
| 131 | AutoDock Vina 1.2.0 | Rosetta Scoring | ✗ | ✗ |
| 132 | AutoDock Vina 1.2.0 | SASA Scoring | ✗ | ✗ |
| 133 | AutoDock Vina 1.2.0 | SEED Scoring | ✗ | ✗ |
| 134 | AutoDock Vina 1.2.0 | SMINA Scoring | ✗ | ✗ |
| 135 | AutoDock Vina 1.2.0 | Vinardo Scoring | ✗ | ✗ |
| 136 | AutoDock Vina 1.2.0 | Zou GB/SA Scoring | ✗ | ✗ |
| 137 | AutoDock Vina 1.1.2 | AD4 Scoring | ✗ | ✗ |
| 138 | AutoDock Vina 1.1.2 | ASP Scoring | ✗ | ✗ |
| 139 | AutoDock Vina 1.1.2 | AutoDock Vina 1.1.2 | ✗ | ✗ |
| 140 | AutoDock Vina 1.1.2 | Bump Filter Scoring | ✗ | ✗ |
| 141 | AutoDock Vina 1.1.2 | ChemPLP | ✗ | ✗ |
| 142 | AutoDock Vina 1.1.2 | ChemScore | ✗ | ✗ |
| 143 | AutoDock Vina 1.1.2 | Contact Scoring | ✗ | ✗ |
| 144 | AutoDock Vina 1.1.2 | Continuous Scoring | ✗ | ✗ |
| 145 | AutoDock Vina 1.1.2 | DOCK 3.5 Scoring | ✗ | ✗ |
| 146 | AutoDock Vina 1.1.2 | DeepAffinity | ✗ | ✓ |
| 147 | AutoDock Vina 1.1.2 | DeepBindRG | ✗ | ✓ |
| 148 | AutoDock Vina 1.1.2 | Footprint Scoring | ✗ | ✗ |
| 149 | AutoDock Vina 1.1.2 | GOLD Scoring | ✗ | ✗ |
| 150 | AutoDock Vina 1.1.2 | Gnina Scoring | ✗ | ✓ |
| 151 | AutoDock Vina 1.1.2 | GoldScore | ✗ | ✗ |
| 152 | AutoDock Vina 1.1.2 | Grid-Based Scoring | ✗ | ✗ |
| 153 | AutoDock Vina 1.1.2 | Hawkins GB/SA Scoring | ✗ | ✗ |
| 154 | AutoDock Vina 1.1.2 | MM/GBSA Scoring | ✗ | ✗ |
| 155 | AutoDock Vina 1.1.2 | MM/PBSA Scoring | ✗ | ✗ |
| 156 | AutoDock Vina 1.1.2 | MultiGrid FPS Scoring | ✗ | ✗ |
| 157 | AutoDock Vina 1.1.2 | NNScore 2.0 | ✗ | ✓ |
| 158 | AutoDock Vina 1.1.2 | OpenBPMD | ✗ | ✗ |
| 159 | AutoDock Vina 1.1.2 | PB/SA Scoring | ✗ | ✗ |
| 160 | AutoDock Vina 1.1.2 | PLANTS Scoring | ✗ | ✗ |
| 161 | AutoDock Vina 1.1.2 | PLP | ✗ | ✗ |
| 162 | AutoDock Vina 1.1.2 | PLP95 | ✗ | ✗ |
| 163 | AutoDock Vina 1.1.2 | Pharmacophore Matching Similarity Scoring | ✗ | ✗ |
| 164 | AutoDock Vina 1.1.2 | RF-Score-VS | ✗ | ✗ |

**Supplementary Table 1 (Continued). docking protocols Supported by the AFVS and AFU Modules.** Each docking protocol is determined by the combination of a pose prediction method and a scoring function. For both of these components, it is indicated whether they are based on deep learning (DL) or not. (Continued on next page.)

| Index | Pose Prediction/Sampling Method | Scoring Function | DL-Based Sampling | DL-Based Scoring |
|---|---|---|---|---|
| 165 | AutoDock Vina 1.1.2 | Rosetta Scoring | ✗ | ✗ |
| 166 | AutoDock Vina 1.1.2 | SASA Scoring | ✗ | ✗ |
| 167 | AutoDock Vina 1.1.2 | SEED Scoring | ✗ | ✗ |
| 168 | AutoDock Vina 1.1.2 | SMINA Scoring | ✗ | ✗ |
| 169 | AutoDock Vina 1.1.2 | Vinardo Scoring | ✗ | ✗ |
| 170 | AutoDock Vina 1.1.2 | Zou GB/SA Scoring | ✗ | ✗ |
| 171 | ADFR | AD4 Scoring | ✗ | ✗ |
| 172 | ADFR | ADFR | ✗ | ✗ |
| 173 | ADFR | ASP Scoring | ✗ | ✗ |
| 174 | ADFR | Bump Filter Scoring | ✗ | ✗ |
| 175 | ADFR | ChemPLP | ✗ | ✗ |
| 176 | ADFR | ChemScore | ✗ | ✗ |
| 177 | ADFR | Contact Scoring | ✗ | ✗ |
| 178 | ADFR | Continuous Scoring | ✗ | ✗ |
| 179 | ADFR | DOCK 3.5 Scoring | ✗ | ✗ |
| 180 | ADFR | DeepAffinity | ✗ | ✓ |
| 181 | ADFR | DeepBindRG | ✗ | ✓ |
| 182 | ADFR | Footprint Scoring | ✗ | ✗ |
| 183 | ADFR | GOLD Scoring | ✗ | ✗ |
| 184 | ADFR | Gnina Scoring | ✗ | ✓ |
| 185 | ADFR | GoldScore | ✗ | ✗ |
| 186 | ADFR | Grid-Based Scoring | ✗ | ✗ |
| 187 | ADFR | Hawkins GB/SA Scoring | ✗ | ✗ |
| 188 | ADFR | MM/GBSA Scoring | ✗ | ✗ |
| 189 | ADFR | MM/PBSA Scoring | ✗ | ✗ |
| 190 | ADFR | MultiGrid FPS Scoring | ✗ | ✗ |
| 191 | ADFR | NNScore 2.0 | ✗ | ✓ |
| 192 | ADFR | OpenBPMD | ✗ | ✗ |
| 193 | ADFR | PB/SA Scoring | ✗ | ✗ |
| 194 | ADFR | PLANTS Scoring | ✗ | ✗ |
| 195 | ADFR | PLP | ✗ | ✗ |
| 196 | ADFR | PLP95 | ✗ | ✗ |
| 197 | ADFR | Pharmacophore Matching Similarity Scoring | ✗ | ✗ |
| 198 | ADFR | RF-Score-VS | ✗ | ✗ |
| 199 | ADFR | Rosetta Scoring | ✗ | ✗ |
| 200 | ADFR | SASA Scoring | ✗ | ✗ |
| 201 | ADFR | SEED Scoring | ✗ | ✗ |
| 202 | ADFR | SMINA Scoring | ✗ | ✗ |
| 203 | ADFR | Vina scoring | ✗ | ✗ |
| 204 | ADFR | Vinardo Scoring | ✗ | ✗ |
| 205 | ADFR | Zou GB/SA Scoring | ✗ | ✗ |
| 206 | AutoDock Koto | AD4 Scoring | ✗ | ✗ |
| 207 | AutoDock Koto | ASP Scoring | ✗ | ✗ |
| 208 | AutoDock Koto | AutoDock Koto | ✗ | ✗ |
| 209 | AutoDock Koto | Bump Filter Scoring | ✗ | ✗ |
| 210 | AutoDock Koto | ChemPLP | ✗ | ✗ |
| 211 | AutoDock Koto | ChemScore | ✗ | ✗ |
| 212 | AutoDock Koto | Contact Scoring | ✗ | ✗ |
| 213 | AutoDock Koto | Continuous Scoring | ✗ | ✗ |
| 214 | AutoDock Koto | DOCK 3.5 Scoring | ✗ | ✗ |
| 215 | AutoDock Koto | DeepAffinity | ✗ | ✓ |
| 216 | AutoDock Koto | DeepBindRG | ✗ | ✓ |
| 217 | AutoDock Koto | Footprint Scoring | ✗ | ✗ |
| 218 | AutoDock Koto | GOLD Scoring | ✗ | ✗ |
| 219 | AutoDock Koto | Gnina Scoring | ✗ | ✓ |

**Supplementary Table 1 (Continued). docking protocols Supported by the AFVS and AFU Modules.** Each docking protocol is determined by the combination of a pose prediction method and a scoring function. For both of these components, it is indicated whether they are based on deep learning (DL) or not. (Continued on next page.)

| Index | Pose Prediction/Sampling Method | Scoring Function | DL-Based Sampling | DL-Based Scoring |
|---|---|---|---|---|
| 220 | AutoDock Koto | GoldScore | ✗ | ✗ |
| 221 | AutoDock Koto | Grid-Based Scoring | ✗ | ✗ |
| 222 | AutoDock Koto | Hawkins GB/SA Scoring | ✗ | ✗ |
| 223 | AutoDock Koto | MM/GBSA Scoring | ✗ | ✗ |
| 224 | AutoDock Koto | MM/PBSA Scoring | ✗ | ✗ |
| 225 | AutoDock Koto | MultiGrid FPS Scoring | ✗ | ✗ |
| 226 | AutoDock Koto | NNScore 2.0 | ✗ | ✓ |
| 227 | AutoDock Koto | OpenBPMD | ✗ | ✗ |
| 228 | AutoDock Koto | PB/SA Scoring | ✗ | ✗ |
| 229 | AutoDock Koto | PLANTS Scoring | ✗ | ✗ |
| 230 | AutoDock Koto | PLP | ✗ | ✗ |
| 231 | AutoDock Koto | PLP95 | ✗ | ✗ |
| 232 | AutoDock Koto | Pharmacophore Matching Similarity Scoring | ✗ | ✗ |
| 233 | AutoDock Koto | RF-Score-VS | ✗ | ✗ |
| 234 | AutoDock Koto | Rosetta Scoring | ✗ | ✗ |
| 235 | AutoDock Koto | SASA Scoring | ✗ | ✗ |
| 236 | AutoDock Koto | SEED Scoring | ✗ | ✗ |
| 237 | AutoDock Koto | SMINA Scoring | ✗ | ✗ |
| 238 | AutoDock Koto | Vina scoring | ✗ | ✗ |
| 239 | AutoDock Koto | Vinardo Scoring | ✗ | ✗ |
| 240 | AutoDock Koto | Zou GB/SA Scoring | ✗ | ✗ |
| 241 | ATPdock | AD4 Scoring | ✗ | ✗ |
| 242 | ATPdock | ASP Scoring | ✗ | ✗ |
| 243 | ATPdock | ATPdock | ✗ | ✗ |
| 244 | ATPdock | Bump Filter Scoring | ✗ | ✗ |
| 245 | ATPdock | ChemPLP | ✗ | ✗ |
| 246 | ATPdock | ChemScore | ✗ | ✗ |
| 247 | ATPdock | Contact Scoring | ✗ | ✗ |
| 248 | ATPdock | Continuous Scoring | ✗ | ✗ |
| 249 | ATPdock | DOCK 3.5 Scoring | ✗ | ✗ |
| 250 | ATPdock | DeepAffinity | ✗ | ✓ |
| 251 | ATPdock | DeepBindRG | ✗ | ✓ |
| 252 | ATPdock | Footprint Scoring | ✗ | ✗ |
| 253 | ATPdock | GOLD Scoring | ✗ | ✗ |
| 254 | ATPdock | Gnina Scoring | ✗ | ✓ |
| 255 | ATPdock | GoldScore | ✗ | ✗ |
| 256 | ATPdock | Grid-Based Scoring | ✗ | ✗ |
| 257 | ATPdock | Hawkins GB/SA Scoring | ✗ | ✗ |
| 258 | ATPdock | MM/GBSA Scoring | ✗ | ✗ |
| 259 | ATPdock | MM/PBSA Scoring | ✗ | ✗ |
| 260 | ATPdock | MultiGrid FPS Scoring | ✗ | ✗ |
| 261 | ATPdock | NNScore 2.0 | ✗ | ✓ |
| 262 | ATPdock | OpenBPMD | ✗ | ✗ |
| 263 | ATPdock | PB/SA Scoring | ✗ | ✗ |
| 264 | ATPdock | PLANTS Scoring | ✗ | ✗ |
| 265 | ATPdock | PLP | ✗ | ✗ |
| 266 | ATPdock | PLP95 | ✗ | ✗ |
| 267 | ATPdock | Pharmacophore Matching Similarity Scoring | ✗ | ✗ |
| 268 | ATPdock | RF-Score-VS | ✗ | ✗ |
| 269 | ATPdock | Rosetta Scoring | ✗ | ✗ |
| 270 | ATPdock | SASA Scoring | ✗ | ✗ |
| 271 | ATPdock | SEED Scoring | ✗ | ✗ |
| 272 | ATPdock | SMINA Scoring | ✗ | ✗ |
| 273 | ATPdock | Vina scoring | ✗ | ✗ |
| 274 | ATPdock | Vinardo Scoring | ✗ | ✗ |

**Supplementary Table 1 (Continued). docking protocols Supported by the AFVS and AFU Modules.** Each docking protocol is determined by the combination of a pose prediction method and a scoring function. For both of these components, it is indicated whether they are based on deep learning (DL) or not. (Continued on next page.)

| Index | Pose Prediction/Sampling Method | Scoring Function | DL-Based Sampling | DL-Based Scoring |
|---|---|---|---|---|
| 275 | ATPdock | Zou GB/SA Scoring | ✗ | ✗ |
| 276 | CovDock | CovDock | ✗ | ✗ |
| 277 | DOCK 6.0 | AD4 Scoring | ✗ | ✗ |
| 278 | DOCK 6.0 | ASP Scoring | ✗ | ✗ |
| 279 | DOCK 6.0 | Bump Filter Scoring | ✗ | ✗ |
| 280 | DOCK 6.0 | ChemPLP | ✗ | ✗ |
| 281 | DOCK 6.0 | ChemScore | ✗ | ✗ |
| 282 | DOCK 6.0 | Contact Scoring | ✗ | ✗ |
| 283 | DOCK 6.0 | Continuous Scoring | ✗ | ✗ |
| 284 | DOCK 6.0 | DOCK 3.5 Scoring | ✗ | ✗ |
| 285 | DOCK 6.0 | DOCK 6.0 | ✗ | ✗ |
| 286 | DOCK 6.0 | DeepAffinity | ✗ | ✓ |
| 287 | DOCK 6.0 | DeepBindRG | ✗ | ✓ |
| 288 | DOCK 6.0 | Footprint Scoring | ✗ | ✗ |
| 289 | DOCK 6.0 | GOLD Scoring | ✗ | ✗ |
| 290 | DOCK 6.0 | Gnina Scoring | ✗ | ✓ |
| 291 | DOCK 6.0 | GoldScore | ✗ | ✗ |
| 292 | DOCK 6.0 | Grid-Based Scoring | ✗ | ✗ |
| 293 | DOCK 6.0 | Hawkins GB/SA Scoring | ✗ | ✗ |
| 294 | DOCK 6.0 | MM/GBSA Scoring | ✗ | ✗ |
| 295 | DOCK 6.0 | MM/PBSA Scoring | ✗ | ✗ |
| 296 | DOCK 6.0 | MultiGrid FPS Scoring | ✗ | ✗ |
| 297 | DOCK 6.0 | NNScore 2.0 | ✗ | ✓ |
| 298 | DOCK 6.0 | OpenBPMD | ✗ | ✗ |
| 299 | DOCK 6.0 | PB/SA Scoring | ✗ | ✗ |
| 300 | DOCK 6.0 | PLANTS Scoring | ✗ | ✗ |
| 301 | DOCK 6.0 | PLP | ✗ | ✗ |
| 302 | DOCK 6.0 | PLP95 | ✗ | ✗ |
| 303 | DOCK 6.0 | Pharmacophore Matching Similarity Scoring | ✗ | ✗ |
| 304 | DOCK 6.0 | RF-Score-VS | ✗ | ✗ |
| 305 | DOCK 6.0 | Rosetta Scoring | ✗ | ✗ |
| 306 | DOCK 6.0 | SASA Scoring | ✗ | ✗ |
| 307 | DOCK 6.0 | SEED Scoring | ✗ | ✗ |
| 308 | DOCK 6.0 | SMINA Scoring | ✗ | ✗ |
| 309 | DOCK 6.0 | Vina scoring | ✗ | ✗ |
| 310 | DOCK 6.0 | Vinardo Scoring | ✗ | ✗ |
| 311 | DOCK 6.0 | Zou GB/SA Scoring | ✗ | ✗ |
| 312 | FlexX | AD4 Scoring | ✗ | ✗ |
| 313 | FlexX | ASP Scoring | ✗ | ✗ |
| 314 | FlexX | Bump Filter Scoring | ✗ | ✗ |
| 315 | FlexX | ChemPLP | ✗ | ✗ |
| 316 | FlexX | ChemScore | ✗ | ✗ |
| 317 | FlexX | Contact Scoring | ✗ | ✗ |
| 318 | FlexX | Continuous Scoring | ✗ | ✗ |
| 319 | FlexX | DOCK 3.5 Scoring | ✗ | ✗ |
| 320 | FlexX | DeepAffinity | ✗ | ✓ |
| 321 | FlexX | DeepBindRG | ✗ | ✓ |
| 322 | FlexX | FlexX | ✗ | ✗ |
| 323 | FlexX | Footprint Scoring | ✗ | ✗ |
| 324 | FlexX | GOLD Scoring | ✗ | ✗ |
| 325 | FlexX | Gnina Scoring | ✗ | ✓ |
| 326 | FlexX | GoldScore | ✗ | ✗ |
| 327 | FlexX | Grid-Based Scoring | ✗ | ✗ |
| 328 | FlexX | Hawkins GB/SA Scoring | ✗ | ✗ |
| 329 | FlexX | MM/GBSA Scoring | ✗ | ✗ |

**Supplementary Table 1 (Continued). docking protocols Supported by the AFVS and AFU Modules.** Each docking protocol is determined by the combination of a pose prediction method and a scoring function. For both of these components, it is indicated whether they are based on deep learning (DL) or not. (Continued on next page.)

| Index | Pose Prediction/Sampling Method | Scoring Function | DL-Based Sampling | DL-Based Scoring |
|---|---|---|---|---|
| 330 | FlexX | MM/PBSA Scoring | ✗ | ✗ |
| 331 | FlexX | MultiGrid FPS Scoring | ✗ | ✗ |
| 332 | FlexX | NNScore 2.0 | ✗ | ✓ |
| 333 | FlexX | OpenBPMD | ✗ | ✗ |
| 334 | FlexX | PB/SA Scoring | ✗ | ✗ |
| 335 | FlexX | PLANTS Scoring | ✗ | ✗ |
| 336 | FlexX | PLP | ✗ | ✗ |
| 337 | FlexX | PLP95 | ✗ | ✗ |
| 338 | FlexX | Pharmacophore Matching Similarity Scoring | ✗ | ✗ |
| 339 | FlexX | RF-Score-VS | ✗ | ✗ |
| 340 | FlexX | Rosetta Scoring | ✗ | ✗ |
| 341 | FlexX | SASA Scoring | ✗ | ✗ |
| 342 | FlexX | SEED Scoring | ✗ | ✗ |
| 343 | FlexX | SMINA Scoring | ✗ | ✗ |
| 344 | FlexX | Vina scoring | ✗ | ✗ |
| 345 | FlexX | Vinardo Scoring | ✗ | ✗ |
| 346 | FlexX | Zou GB/SA Scoring | ✗ | ✗ |
| 347 | FitDock | AD4 Scoring | ✗ | ✗ |
| 348 | FitDock | ASP Scoring | ✗ | ✗ |
| 349 | FitDock | Bump Filter Scoring | ✗ | ✗ |
| 350 | FitDock | ChemPLP | ✗ | ✗ |
| 351 | FitDock | ChemScore | ✗ | ✗ |
| 352 | FitDock | Contact Scoring | ✗ | ✗ |
| 353 | FitDock | Continuous Scoring | ✗ | ✗ |
| 354 | FitDock | DOCK 3.5 Scoring | ✗ | ✗ |
| 355 | FitDock | DeepAffinity | ✗ | ✓ |
| 356 | FitDock | DeepBindRG | ✗ | ✓ |
| 357 | FitDock | FitDock | ✗ | ✗ |
| 358 | FitDock | Footprint Scoring | ✗ | ✗ |
| 359 | FitDock | GOLD Scoring | ✗ | ✗ |
| 360 | FitDock | Gnina Scoring | ✗ | ✓ |
| 361 | FitDock | GoldScore | ✗ | ✗ |
| 362 | FitDock | Grid-Based Scoring | ✗ | ✗ |
| 363 | FitDock | Hawkins GB/SA Scoring | ✗ | ✗ |
| 364 | FitDock | MM/GBSA Scoring | ✗ | ✗ |
| 365 | FitDock | MM/PBSA Scoring | ✗ | ✗ |
| 366 | FitDock | MultiGrid FPS Scoring | ✗ | ✗ |
| 367 | FitDock | NNScore 2.0 | ✗ | ✓ |
| 368 | FitDock | OpenBPMD | ✗ | ✗ |
| 369 | FitDock | PB/SA Scoring | ✗ | ✗ |
| 370 | FitDock | PLANTS Scoring | ✗ | ✗ |
| 371 | FitDock | PLP | ✗ | ✗ |
| 372 | FitDock | PLP95 | ✗ | ✗ |
| 373 | FitDock | Pharmacophore Matching Similarity Scoring | ✗ | ✗ |
| 374 | FitDock | RF-Score-VS | ✗ | ✗ |
| 375 | FitDock | Rosetta Scoring | ✗ | ✗ |
| 376 | FitDock | SASA Scoring | ✗ | ✗ |
| 377 | FitDock | SEED Scoring | ✗ | ✗ |
| 378 | FitDock | SMINA Scoring | ✗ | ✗ |
| 379 | FitDock | Vina scoring | ✗ | ✗ |
| 380 | FitDock | Vinardo Scoring | ✗ | ✗ |
| 381 | FitDock | Zou GB/SA Scoring | ✗ | ✗ |
| 382 | GalaxyDock3 | AD4 Scoring | ✗ | ✗ |
| 383 | GalaxyDock3 | ASP Scoring | ✗ | ✗ |
| 384 | GalaxyDock3 | Bump Filter Scoring | ✗ | ✗ |

**Supplementary Table 1 (Continued). docking protocols Supported by the AFVS and AFU Modules.** Each docking protocol is determined by the combination of a pose prediction method and a scoring function. For both of these components, it is indicated whether they are based on deep learning (DL) or not. (Continued on next page.)

| Index | Pose Prediction/Sampling Method | Scoring Function | DL-Based Sampling | DL-Based Scoring |
|---|---|---|---|---|
| 385 | GalaxyDock3 | ChemPLP | ✗ | ✗ |
| 386 | GalaxyDock3 | ChemScore | ✗ | ✗ |
| 387 | GalaxyDock3 | Contact Scoring | ✗ | ✗ |
| 388 | GalaxyDock3 | Continuous Scoring | ✗ | ✗ |
| 389 | GalaxyDock3 | DOCK 3.5 Scoring | ✗ | ✗ |
| 390 | GalaxyDock3 | DeepAffinity | ✗ | ✓ |
| 391 | GalaxyDock3 | DeepBindRG | ✗ | ✓ |
| 392 | GalaxyDock3 | Footprint Scoring | ✗ | ✗ |
| 393 | GalaxyDock3 | GOLD Scoring | ✗ | ✗ |
| 394 | GalaxyDock3 | GalaxyDock3 | ✗ | ✗ |
| 395 | GalaxyDock3 | Gnina Scoring | ✗ | ✓ |
| 396 | GalaxyDock3 | GoldScore | ✗ | ✗ |
| 397 | GalaxyDock3 | Grid-Based Scoring | ✗ | ✗ |
| 398 | GalaxyDock3 | Hawkins GB/SA Scoring | ✗ | ✗ |
| 399 | GalaxyDock3 | MM/GBSA Scoring | ✗ | ✗ |
| 400 | GalaxyDock3 | MM/PBSA Scoring | ✗ | ✗ |
| 401 | GalaxyDock3 | MultiGrid FPS Scoring | ✗ | ✗ |
| 402 | GalaxyDock3 | NNScore 2.0 | ✗ | ✓ |
| 403 | GalaxyDock3 | OpenBPMD | ✗ | ✗ |
| 404 | GalaxyDock3 | PB/SA Scoring | ✗ | ✗ |
| 405 | GalaxyDock3 | PLANTS Scoring | ✗ | ✗ |
| 406 | GalaxyDock3 | PLP | ✗ | ✗ |
| 407 | GalaxyDock3 | PLP95 | ✗ | ✗ |
| 408 | GalaxyDock3 | Pharmacophore Matching Similarity Scoring | ✗ | ✗ |
| 409 | GalaxyDock3 | RF-Score-VS | ✗ | ✗ |
| 410 | GalaxyDock3 | Rosetta Scoring | ✗ | ✗ |
| 411 | GalaxyDock3 | SASA Scoring | ✗ | ✗ |
| 412 | GalaxyDock3 | SEED Scoring | ✗ | ✗ |
| 413 | GalaxyDock3 | SMINA Scoring | ✗ | ✗ |
| 414 | GalaxyDock3 | Vina scoring | ✗ | ✗ |
| 415 | GalaxyDock3 | Vinardo Scoring | ✗ | ✗ |
| 416 | GalaxyDock3 | Zou GB/SA Scoring | ✗ | ✗ |
| 417 | Glide (HTVS) | AD4 Scoring | ✗ | ✗ |
| 418 | Glide (HTVS) | ASP Scoring | ✗ | ✗ |
| 419 | Glide (HTVS) | Bump Filter Scoring | ✗ | ✗ |
| 420 | Glide (HTVS) | ChemPLP | ✗ | ✗ |
| 421 | Glide (HTVS) | ChemScore | ✗ | ✗ |
| 422 | Glide (HTVS) | Contact Scoring | ✗ | ✗ |
| 423 | Glide (HTVS) | Continuous Scoring | ✗ | ✗ |
| 424 | Glide (HTVS) | DOCK 3.5 Scoring | ✗ | ✗ |
| 425 | Glide (HTVS) | DeepAffinity | ✗ | ✓ |
| 426 | Glide (HTVS) | DeepBindRG | ✗ | ✓ |
| 427 | Glide (HTVS) | Footprint Scoring | ✗ | ✗ |
| 428 | Glide (HTVS) | GOLD Scoring | ✗ | ✗ |
| 429 | Glide (HTVS) | Glide (HTVS) | ✗ | ✗ |
| 430 | Glide (HTVS) | Gnina Scoring | ✗ | ✓ |
| 431 | Glide (HTVS) | GoldScore | ✗ | ✗ |
| 432 | Glide (HTVS) | Grid-Based Scoring | ✗ | ✗ |
| 433 | Glide (HTVS) | Hawkins GB/SA Scoring | ✗ | ✗ |
| 434 | Glide (HTVS) | MM/GBSA Scoring | ✗ | ✗ |
| 435 | Glide (HTVS) | MM/PBSA Scoring | ✗ | ✗ |
| 436 | Glide (HTVS) | MultiGrid FPS Scoring | ✗ | ✗ |
| 437 | Glide (HTVS) | NNScore 2.0 | ✗ | ✓ |
| 438 | Glide (HTVS) | OpenBPMD | ✗ | ✗ |
| 439 | Glide (HTVS) | PB/SA Scoring | ✗ | ✗ |

**Supplementary Table 1 (Continued). docking protocols Supported by the AFVS and AFU Modules.** Each docking protocol is determined by the combination of a pose prediction method and a scoring function. For both of these components, it is indicated whether they are based on deep learning (DL) or not. (Continued on next page.)

| Index | Pose Prediction/Sampling Method | Scoring Function | DL-Based Sampling | DL-Based Scoring |
|---|---|---|---|---|
| 440 | Glide (HTVS) | PLANTS Scoring | ✗ | ✗ |
| 441 | Glide (HTVS) | PLP | ✗ | ✗ |
| 442 | Glide (HTVS) | PLP95 | ✗ | ✗ |
| 443 | Glide (HTVS) | Pharmacophore Matching Similarity Scoring | ✗ | ✗ |
| 444 | Glide (HTVS) | RF-Score-VS | ✗ | ✗ |
| 445 | Glide (HTVS) | Rosetta Scoring | ✗ | ✗ |
| 446 | Glide (HTVS) | SASA Scoring | ✗ | ✗ |
| 447 | Glide (HTVS) | SEED Scoring | ✗ | ✗ |
| 448 | Glide (HTVS) | SMINA Scoring | ✗ | ✗ |
| 449 | Glide (HTVS) | Vina scoring | ✗ | ✗ |
| 450 | Glide (HTVS) | Vinardo Scoring | ✗ | ✗ |
| 451 | Glide (HTVS) | Zou GB/SA Scoring | ✗ | ✗ |
| 452 | Glide (SP) | AD4 Scoring | ✗ | ✗ |
| 453 | Glide (SP) | ASP Scoring | ✗ | ✗ |
| 454 | Glide (SP) | Bump Filter Scoring | ✗ | ✗ |
| 455 | Glide (SP) | ChemPLP | ✗ | ✗ |
| 456 | Glide (SP) | ChemScore | ✗ | ✗ |
| 457 | Glide (SP) | Contact Scoring | ✗ | ✗ |
| 458 | Glide (SP) | Continuous Scoring | ✗ | ✗ |
| 459 | Glide (SP) | DOCK 3.5 Scoring | ✗ | ✗ |
| 460 | Glide (SP) | DeepAffinity | ✗ | ✓ |
| 461 | Glide (SP) | DeepBindRG | ✗ | ✓ |
| 462 | Glide (SP) | Footprint Scoring | ✗ | ✗ |
| 463 | Glide (SP) | GOLD Scoring | ✗ | ✗ |
| 464 | Glide (SP) | Glide (SP) | ✗ | ✗ |
| 465 | Glide (SP) | Gnina Scoring | ✗ | ✓ |
| 466 | Glide (SP) | GoldScore | ✗ | ✗ |
| 467 | Glide (SP) | Grid-Based Scoring | ✗ | ✗ |
| 468 | Glide (SP) | Hawkins GB/SA Scoring | ✗ | ✗ |
| 469 | Glide (SP) | MM/GBSA Scoring | ✗ | ✗ |
| 470 | Glide (SP) | MM/PBSA Scoring | ✗ | ✗ |
| 471 | Glide (SP) | MultiGrid FPS Scoring | ✗ | ✗ |
| 472 | Glide (SP) | NNScore 2.0 | ✗ | ✓ |
| 473 | Glide (SP) | OpenBPMD | ✗ | ✗ |
| 474 | Glide (SP) | PB/SA Scoring | ✗ | ✗ |
| 475 | Glide (SP) | PLANTS Scoring | ✗ | ✗ |
| 476 | Glide (SP) | PLP | ✗ | ✗ |
| 477 | Glide (SP) | PLP95 | ✗ | ✗ |
| 478 | Glide (SP) | Pharmacophore Matching Similarity Scoring | ✗ | ✗ |
| 479 | Glide (SP) | RF-Score-VS | ✗ | ✗ |
| 480 | Glide (SP) | Rosetta Scoring | ✗ | ✗ |
| 481 | Glide (SP) | SASA Scoring | ✗ | ✗ |
| 482 | Glide (SP) | SEED Scoring | ✗ | ✗ |
| 483 | Glide (SP) | SMINA Scoring | ✗ | ✗ |
| 484 | Glide (SP) | Vina scoring | ✗ | ✗ |
| 485 | Glide (SP) | Vinardo Scoring | ✗ | ✗ |
| 486 | Glide (SP) | Zou GB/SA Scoring | ✗ | ✗ |
| 487 | Glide (XP) | AD4 Scoring | ✗ | ✗ |
| 488 | Glide (XP) | ASP Scoring | ✗ | ✗ |
| 489 | Glide (XP) | Bump Filter Scoring | ✗ | ✗ |
| 490 | Glide (XP) | ChemPLP | ✗ | ✗ |
| 491 | Glide (XP) | ChemScore | ✗ | ✗ |
| 492 | Glide (XP) | Contact Scoring | ✗ | ✗ |
| 493 | Glide (XP) | Continuous Scoring | ✗ | ✗ |
| 494 | Glide (XP) | DOCK 3.5 Scoring | ✗ | ✗ |

**Supplementary Table 1 (Continued). docking protocols Supported by the AFVS and AFU Modules.** Each docking protocol is determined by the combination of a pose prediction method and a scoring function. For both of these components, it is indicated whether they are based on deep learning (DL) or not. (Continued on next page.)

| Index | Pose Prediction/Sampling Method | Scoring Function | DL-Based Sampling | DL-Based Scoring |
|---|---|---|---|---|
| 495 | Glide (XP) | DeepAffinity | ✗ | ✓ |
| 496 | Glide (XP) | DeepBindRG | ✗ | ✓ |
| 497 | Glide (XP) | Footprint Scoring | ✗ | ✗ |
| 498 | Glide (XP) | GOLD Scoring | ✗ | ✗ |
| 499 | Glide (XP) | Glide (XP) | ✗ | ✗ |
| 500 | Glide (XP) | Gnina Scoring | ✗ | ✓ |
| 501 | Glide (XP) | GoldScore | ✗ | ✗ |
| 502 | Glide (XP) | Grid-Based Scoring | ✗ | ✗ |
| 503 | Glide (XP) | Hawkins GB/SA Scoring | ✗ | ✗ |
| 504 | Glide (XP) | MM/GBSA Scoring | ✗ | ✗ |
| 505 | Glide (XP) | MM/PBSA Scoring | ✗ | ✗ |
| 506 | Glide (XP) | MultiGrid FPS Scoring | ✗ | ✗ |
| 507 | Glide (XP) | NNScore 2.0 | ✗ | ✓ |
| 508 | Glide (XP) | OpenBPMD | ✗ | ✗ |
| 509 | Glide (XP) | PB/SA Scoring | ✗ | ✗ |
| 510 | Glide (XP) | PLANTS Scoring | ✗ | ✗ |
| 511 | Glide (XP) | PLP | ✗ | ✗ |
| 512 | Glide (XP) | PLP95 | ✗ | ✗ |
| 513 | Glide (XP) | Pharmacophore Matching Similarity Scoring | ✗ | ✗ |
| 514 | Glide (XP) | RF-Score-VS | ✗ | ✗ |
| 515 | Glide (XP) | Rosetta Scoring | ✗ | ✗ |
| 516 | Glide (XP) | SASA Scoring | ✗ | ✗ |
| 517 | Glide (XP) | SEED Scoring | ✗ | ✗ |
| 518 | Glide (XP) | SMINA Scoring | ✗ | ✗ |
| 519 | Glide (XP) | Vina scoring | ✗ | ✗ |
| 520 | Glide (XP) | Vinardo Scoring | ✗ | ✗ |
| 521 | Glide (XP) | Zou GB/SA Scoring | ✗ | ✗ |
| 522 | Gnina | AD4 Scoring | ✗ | ✗ |
| 523 | Gnina | ASP Scoring | ✗ | ✗ |
| 524 | Gnina | Bump Filter Scoring | ✗ | ✗ |
| 525 | Gnina | ChemPLP | ✗ | ✗ |
| 526 | Gnina | ChemScore | ✗ | ✗ |
| 527 | Gnina | Contact Scoring | ✗ | ✗ |
| 528 | Gnina | Continuous Scoring | ✗ | ✗ |
| 529 | Gnina | DOCK 3.5 Scoring | ✗ | ✗ |
| 530 | Gnina | DeepAffinity | ✗ | ✓ |
| 531 | Gnina | DeepBindRG | ✗ | ✓ |
| 532 | Gnina | Footprint Scoring | ✗ | ✗ |
| 533 | Gnina | GOLD Scoring | ✗ | ✗ |
| 534 | Gnina | Gnina | ✗ | ✗ |
| 535 | Gnina | GoldScore | ✗ | ✗ |
| 536 | Gnina | Grid-Based Scoring | ✗ | ✗ |
| 537 | Gnina | Hawkins GB/SA Scoring | ✗ | ✗ |
| 538 | Gnina | MM/GBSA Scoring | ✗ | ✗ |
| 539 | Gnina | MM/PBSA Scoring | ✗ | ✗ |
| 540 | Gnina | MultiGrid FPS Scoring | ✗ | ✗ |
| 541 | Gnina | NNScore 2.0 | ✗ | ✓ |
| 542 | Gnina | OpenBPMD | ✗ | ✗ |
| 543 | Gnina | PB/SA Scoring | ✗ | ✗ |
| 544 | Gnina | PLANTS Scoring | ✗ | ✗ |
| 545 | Gnina | PLP | ✗ | ✗ |
| 546 | Gnina | PLP95 | ✗ | ✗ |
| 547 | Gnina | Pharmacophore Matching Similarity Scoring | ✗ | ✗ |
| 548 | Gnina | RF-Score-VS | ✗ | ✗ |
| 549 | Gnina | Rosetta Scoring | ✗ | ✗ |

**Supplementary Table 1 (Continued). docking protocols Supported by the AFVS and AFU Modules.** Each docking protocol is determined by the combination of a pose prediction method and a scoring function. For both of these components, it is indicated whether they are based on deep learning (DL) or not. (Continued on next page.)

| Index | Pose Prediction/Sampling Method | Scoring Function | DL-Based Sampling | DL-Based Scoring |
|---|---|---|---|---|
| 550 | Gnina | SASA Scoring | ✗ | ✗ |
| 551 | Gnina | SEED Scoring | ✗ | ✗ |
| 552 | Gnina | SMINA Scoring | ✗ | ✗ |
| 553 | Gnina | Vina scoring | ✗ | ✗ |
| 554 | Gnina | Vinardo Scoring | ✗ | ✗ |
| 555 | Gnina | Zou GB/SA Scoring | ✗ | ✗ |
| 556 | GOLD | AD4 Scoring | ✗ | ✗ |
| 557 | GOLD | ASP Scoring | ✗ | ✗ |
| 558 | GOLD | Bump Filter Scoring | ✗ | ✗ |
| 559 | GOLD | ChemPLP | ✗ | ✗ |
| 560 | GOLD | ChemScore | ✗ | ✗ |
| 561 | GOLD | Contact Scoring | ✗ | ✗ |
| 562 | GOLD | Continuous Scoring | ✗ | ✗ |
| 563 | GOLD | DOCK 3.5 Scoring | ✗ | ✗ |
| 564 | GOLD | DeepAffinity | ✗ | ✓ |
| 565 | GOLD | DeepBindRG | ✗ | ✓ |
| 566 | GOLD | Footprint Scoring | ✗ | ✗ |
| 567 | GOLD | GOLD | ✗ | ✗ |
| 568 | GOLD | Gnina Scoring | ✗ | ✓ |
| 569 | GOLD | GoldScore | ✗ | ✗ |
| 570 | GOLD | Grid-Based Scoring | ✗ | ✗ |
| 571 | GOLD | Hawkins GB/SA Scoring | ✗ | ✗ |
| 572 | GOLD | MM/GBSA Scoring | ✗ | ✗ |
| 573 | GOLD | MM/PBSA Scoring | ✗ | ✗ |
| 574 | GOLD | MultiGrid FPS Scoring | ✗ | ✗ |
| 575 | GOLD | NNScore 2.0 | ✗ | ✓ |
| 576 | GOLD | OpenBPMD | ✗ | ✗ |
| 577 | GOLD | PB/SA Scoring | ✗ | ✗ |
| 578 | GOLD | PLANTS Scoring | ✗ | ✗ |
| 579 | GOLD | PLP | ✗ | ✗ |
| 580 | GOLD | PLP95 | ✗ | ✗ |
| 581 | GOLD | Pharmacophore Matching Similarity Scoring | ✗ | ✗ |
| 582 | GOLD | RF-Score-VS | ✗ | ✗ |
| 583 | GOLD | Rosetta Scoring | ✗ | ✗ |
| 584 | GOLD | SASA Scoring | ✗ | ✗ |
| 585 | GOLD | SEED Scoring | ✗ | ✗ |
| 586 | GOLD | SMINA Scoring | ✗ | ✗ |
| 587 | GOLD | Vina scoring | ✗ | ✗ |
| 588 | GOLD | Vinardo Scoring | ✗ | ✗ |
| 589 | GOLD | Zou GB/SA Scoring | ✗ | ✗ |
| 590 | GWOVina | AD4 Scoring | ✗ | ✗ |
| 591 | GWOVina | ASP Scoring | ✗ | ✗ |
| 592 | GWOVina | Bump Filter Scoring | ✗ | ✗ |
| 593 | GWOVina | ChemPLP | ✗ | ✗ |
| 594 | GWOVina | ChemScore | ✗ | ✗ |
| 595 | GWOVina | Contact Scoring | ✗ | ✗ |
| 596 | GWOVina | Continuous Scoring | ✗ | ✗ |
| 597 | GWOVina | DOCK 3.5 Scoring | ✗ | ✗ |
| 598 | GWOVina | DeepAffinity | ✗ | ✓ |
| 599 | GWOVina | DeepBindRG | ✗ | ✓ |
| 600 | GWOVina | Footprint Scoring | ✗ | ✗ |
| 601 | GWOVina | GWOVina | ✗ | ✗ |
| 602 | GWOVina | Gnina Scoring | ✗ | ✓ |
| 603 | GWOVina | GoldScore | ✗ | ✗ |
| 604 | GWOVina | Grid-Based Scoring | ✗ | ✗ |

**Supplementary Table 1 (Continued). docking protocols Supported by the AFVS and AFU Modules.** Each docking protocol is determined by the combination of a pose prediction method and a scoring function. For both of these components, it is indicated whether they are based on deep learning (DL) or not. (Continued on next page.)

| Index | Pose Prediction/Sampling Method | Scoring Function | DL-Based Sampling | DL-Based Scoring |
|---|---|---|---|---|
| 605 | GWOVina | Hawkins GB/SA Scoring | ✗ | ✗ |
| 606 | GWOVina | MM/GBSA Scoring | ✗ | ✗ |
| 607 | GWOVina | MM/PBSA Scoring | ✗ | ✗ |
| 608 | GWOVina | MultiGrid FPS Scoring | ✗ | ✗ |
| 609 | GWOVina | NNScore 2.0 | ✗ | ✓ |
| 610 | GWOVina | OpenBPMD | ✗ | ✗ |
| 611 | GWOVina | PB/SA Scoring | ✗ | ✗ |
| 612 | GWOVina | PLANTS Scoring | ✗ | ✗ |
| 613 | GWOVina | PLP | ✗ | ✗ |
| 614 | GWOVina | PLP95 | ✗ | ✗ |
| 615 | GWOVina | Pharmacophore Matching Similarity Scoring | ✗ | ✗ |
| 616 | GWOVina | RF-Score-VS | ✗ | ✗ |
| 617 | GWOVina | Rosetta Scoring | ✗ | ✗ |
| 618 | GWOVina | SASA Scoring | ✗ | ✗ |
| 619 | GWOVina | SEED Scoring | ✗ | ✗ |
| 620 | GWOVina | SMINA Scoring | ✗ | ✗ |
| 621 | GWOVina | Vina scoring | ✗ | ✗ |
| 622 | GWOVina | Vinardo Scoring | ✗ | ✗ |
| 623 | GWOVina | Zou GB/SA Scoring | ✗ | ✗ |
| 624 | HDOCK | HDOCK | ✗ | ✗ |
| 625 | HSYMDOCK | HSYMDOCK | ✗ | ✗ |
| 626 | idock | AD4 Scoring | ✗ | ✗ |
| 627 | idock | ASP Scoring | ✗ | ✗ |
| 628 | idock | Bump Filter Scoring | ✗ | ✗ |
| 629 | idock | ChemPLP | ✗ | ✗ |
| 630 | idock | ChemScore | ✗ | ✗ |
| 631 | idock | Contact Scoring | ✗ | ✗ |
| 632 | idock | Continuous Scoring | ✗ | ✗ |
| 633 | idock | DOCK 3.5 Scoring | ✗ | ✗ |
| 634 | idock | DeepAffinity | ✗ | ✓ |
| 635 | idock | DeepBindRG | ✗ | ✓ |
| 636 | idock | Footprint Scoring | ✗ | ✗ |
| 637 | idock | GOLD Scoring | ✗ | ✗ |
| 638 | idock | Gnina Scoring | ✗ | ✓ |
| 639 | idock | GoldScore | ✗ | ✗ |
| 640 | idock | Grid-Based Scoring | ✗ | ✗ |
| 641 | idock | Hawkins GB/SA Scoring | ✗ | ✗ |
| 642 | idock | MM/GBSA Scoring | ✗ | ✗ |
| 643 | idock | MM/PBSA Scoring | ✗ | ✗ |
| 644 | idock | MultiGrid FPS Scoring | ✗ | ✗ |
| 645 | idock | NNScore 2.0 | ✗ | ✓ |
| 646 | idock | OpenBPMD | ✗ | ✗ |
| 647 | idock | PB/SA Scoring | ✗ | ✗ |
| 648 | idock | PLANTS Scoring | ✗ | ✗ |
| 649 | idock | PLP | ✗ | ✗ |
| 650 | idock | PLP95 | ✗ | ✗ |
| 651 | idock | Pharmacophore Matching Similarity Scoring | ✗ | ✗ |
| 652 | idock | RF-Score-VS | ✗ | ✗ |
| 653 | idock | Rosetta Scoring | ✗ | ✗ |
| 654 | idock | SASA Scoring | ✗ | ✗ |
| 655 | idock | SEED Scoring | ✗ | ✗ |
| 656 | idock | SMINA Scoring | ✗ | ✗ |
| 657 | idock | Vinardo Scoring | ✗ | ✗ |
| 658 | idock | Zou GB/SA Scoring | ✗ | ✗ |
| 659 | idock | idock | ✗ | ✗ |

**Supplementary Table 1 (Continued). docking protocols Supported by the AFVS and AFU Modules.** Each docking protocol is determined by the combination of a pose prediction method and a scoring function. For both of these components, it is indicated whether they are based on deep learning (DL) or not. (Continued on next page.)

| Index | Pose Prediction/Sampling Method | Scoring Function | DL-Based Sampling | DL-Based Scoring |
|---|---|---|---|---|
| 660 | iGEMDOCK | AD4 Scoring | ✗ | ✗ |
| 661 | iGEMDOCK | ASP Scoring | ✗ | ✗ |
| 662 | iGEMDOCK | Bump Filter Scoring | ✗ | ✗ |
| 663 | iGEMDOCK | ChemPLP | ✗ | ✗ |
| 664 | iGEMDOCK | ChemScore | ✗ | ✗ |
| 665 | iGEMDOCK | Contact Scoring | ✗ | ✗ |
| 666 | iGEMDOCK | Continuous Scoring | ✗ | ✗ |
| 667 | iGEMDOCK | DOCK 3.5 Scoring | ✗ | ✗ |
| 668 | iGEMDOCK | DeepAffinity | ✗ | ✓ |
| 669 | iGEMDOCK | DeepBindRG | ✗ | ✓ |
| 670 | iGEMDOCK | Footprint Scoring | ✗ | ✗ |
| 671 | iGEMDOCK | GOLD Scoring | ✗ | ✗ |
| 672 | iGEMDOCK | Gnina Scoring | ✗ | ✓ |
| 673 | iGEMDOCK | GoldScore | ✗ | ✗ |
| 674 | iGEMDOCK | Grid-Based Scoring | ✗ | ✗ |
| 675 | iGEMDOCK | Hawkins GB/SA Scoring | ✗ | ✗ |
| 676 | iGEMDOCK | MM/GBSA Scoring | ✗ | ✗ |
| 677 | iGEMDOCK | MM/PBSA Scoring | ✗ | ✗ |
| 678 | iGEMDOCK | MultiGrid FPS Scoring | ✗ | ✗ |
| 679 | iGEMDOCK | NNScore 2.0 | ✗ | ✓ |
| 680 | iGEMDOCK | OpenBPMD | ✗ | ✗ |
| 681 | iGEMDOCK | PB/SA Scoring | ✗ | ✗ |
| 682 | iGEMDOCK | PLANTS Scoring | ✗ | ✗ |
| 683 | iGEMDOCK | PLP | ✗ | ✗ |
| 684 | iGEMDOCK | PLP95 | ✗ | ✗ |
| 685 | iGEMDOCK | Pharmacophore Matching Similarity Scoring | ✗ | ✗ |
| 686 | iGEMDOCK | RF-Score-VS | ✗ | ✗ |
| 687 | iGEMDOCK | Rosetta Scoring | ✗ | ✗ |
| 688 | iGEMDOCK | SASA Scoring | ✗ | ✗ |
| 689 | iGEMDOCK | SEED Scoring | ✗ | ✗ |
| 690 | iGEMDOCK | SMINA Scoring | ✗ | ✗ |
| 691 | iGEMDOCK | Vina scoring | ✗ | ✗ |
| 692 | iGEMDOCK | Vinardo Scoring | ✗ | ✗ |
| 693 | iGEMDOCK | Zou GB/SA Scoring | ✗ | ✗ |
| 694 | iGEMDOCK | iGEMDOCK | ✗ | ✗ |
| 695 | LeDock | AD4 Scoring | ✗ | ✗ |
| 696 | LeDock | ASP Scoring | ✗ | ✗ |
| 697 | LeDock | Bump Filter Scoring | ✗ | ✗ |
| 698 | LeDock | ChemPLP | ✗ | ✗ |
| 699 | LeDock | ChemScore | ✗ | ✗ |
| 700 | LeDock | Contact Scoring | ✗ | ✗ |
| 701 | LeDock | Continuous Scoring | ✗ | ✗ |
| 702 | LeDock | DOCK 3.5 Scoring | ✗ | ✗ |
| 703 | LeDock | DeepAffinity | ✗ | ✓ |
| 704 | LeDock | DeepBindRG | ✗ | ✓ |
| 705 | LeDock | Footprint Scoring | ✗ | ✗ |
| 706 | LeDock | GOLD Scoring | ✗ | ✗ |
| 707 | LeDock | Gnina Scoring | ✗ | ✓ |
| 708 | LeDock | GoldScore | ✗ | ✗ |
| 709 | LeDock | Grid-Based Scoring | ✗ | ✗ |
| 710 | LeDock | Hawkins GB/SA Scoring | ✗ | ✗ |
| 711 | LeDock | LeDock | ✗ | ✗ |
| 712 | LeDock | MM/GBSA Scoring | ✗ | ✗ |
| 713 | LeDock | MM/PBSA Scoring | ✗ | ✗ |
| 714 | LeDock | MultiGrid FPS Scoring | ✗ | ✗ |

**Supplementary Table 1 (Continued). docking protocols Supported by the AFVS and AFU Modules.** Each docking protocol is determined by the combination of a pose prediction method and a scoring function. For both of these components, it is indicated whether they are based on deep learning (DL) or not. (Continued on next page.)

| Index | Pose Prediction/Sampling Method | Scoring Function | DL-Based Sampling | DL-Based Scoring |
|---|---|---|---|---|
| 715 | LeDock | NNScore 2.0 | ✗ | ✓ |
| 716 | LeDock | OpenBPMD | ✗ | ✗ |
| 717 | LeDock | PB/SA Scoring | ✗ | ✗ |
| 718 | LeDock | PLANTS Scoring | ✗ | ✗ |
| 719 | LeDock | PLP | ✗ | ✗ |
| 720 | LeDock | PLP95 | ✗ | ✗ |
| 721 | LeDock | Pharmacophore Matching Similarity Scoring | ✗ | ✗ |
| 722 | LeDock | RF-Score-VS | ✗ | ✗ |
| 723 | LeDock | Rosetta Scoring | ✗ | ✗ |
| 724 | LeDock | SASA Scoring | ✗ | ✗ |
| 725 | LeDock | SEED Scoring | ✗ | ✗ |
| 726 | LeDock | SMINA Scoring | ✗ | ✗ |
| 727 | LeDock | Vina scoring | ✗ | ✗ |
| 728 | LeDock | Vinardo Scoring | ✗ | ✗ |
| 729 | LeDock | Zou GB/SA Scoring | ✗ | ✗ |
| 730 | LigandFit | AD4 Scoring | ✗ | ✗ |
| 731 | LigandFit | ASP Scoring | ✗ | ✗ |
| 732 | LigandFit | Bump Filter Scoring | ✗ | ✗ |
| 733 | LigandFit | ChemPLP | ✗ | ✗ |
| 734 | LigandFit | ChemScore | ✗ | ✗ |
| 735 | LigandFit | Contact Scoring | ✗ | ✗ |
| 736 | LigandFit | Continuous Scoring | ✗ | ✗ |
| 737 | LigandFit | DOCK 3.5 Scoring | ✗ | ✗ |
| 738 | LigandFit | DeepAffinity | ✗ | ✓ |
| 739 | LigandFit | DeepBindRG | ✗ | ✓ |
| 740 | LigandFit | Footprint Scoring | ✗ | ✗ |
| 741 | LigandFit | GOLD Scoring | ✗ | ✗ |
| 742 | LigandFit | Gnina Scoring | ✗ | ✓ |
| 743 | LigandFit | GoldScore | ✗ | ✗ |
| 744 | LigandFit | Grid-Based Scoring | ✗ | ✗ |
| 745 | LigandFit | Hawkins GB/SA Scoring | ✗ | ✗ |
| 746 | LigandFit | LigandFit | ✗ | ✗ |
| 747 | LigandFit | MM/GBSA Scoring | ✗ | ✗ |
| 748 | LigandFit | MM/PBSA Scoring | ✗ | ✗ |
| 749 | LigandFit | MultiGrid FPS Scoring | ✗ | ✗ |
| 750 | LigandFit | NNScore 2.0 | ✗ | ✓ |
| 751 | LigandFit | OpenBPMD | ✗ | ✗ |
| 752 | LigandFit | PB/SA Scoring | ✗ | ✗ |
| 753 | LigandFit | PLANTS Scoring | ✗ | ✗ |
| 754 | LigandFit | PLP | ✗ | ✗ |
| 755 | LigandFit | PLP95 | ✗ | ✗ |
| 756 | LigandFit | Pharmacophore Matching Similarity Scoring | ✗ | ✗ |
| 757 | LigandFit | RF-Score-VS | ✗ | ✗ |
| 758 | LigandFit | Rosetta Scoring | ✗ | ✗ |
| 759 | LigandFit | SASA Scoring | ✗ | ✗ |
| 760 | LigandFit | SEED Scoring | ✗ | ✗ |
| 761 | LigandFit | SMINA Scoring | ✗ | ✗ |
| 762 | LigandFit | Vina scoring | ✗ | ✗ |
| 763 | LigandFit | Vinardo Scoring | ✗ | ✗ |
| 764 | LigandFit | Zou GB/SA Scoring | ✗ | ✗ |
| 765 | LightDock | LightDock | ✗ | ✗ |
| 766 | MDock | AD4 Scoring | ✗ | ✗ |
| 767 | MDock | ASP Scoring | ✗ | ✗ |
| 768 | MDock | Bump Filter Scoring | ✗ | ✗ |
| 769 | MDock | ChemPLP | ✗ | ✗ |

**Supplementary Table 1 (Continued). docking protocols Supported by the AFVS and AFU Modules.** Each docking protocol is determined by the combination of a pose prediction method and a scoring function. For both of these components, it is indicated whether they are based on deep learning (DL) or not. (Continued on next page.)

| Index | Pose Prediction/Sampling Method | Scoring Function | DL-Based Sampling | DL-Based Scoring |
|---|---|---|---|---|
| 770 | MDock | ChemScore | ✗ | ✗ |
| 771 | MDock | Contact Scoring | ✗ | ✗ |
| 772 | MDock | Continuous Scoring | ✗ | ✗ |
| 773 | MDock | DOCK 3.5 Scoring | ✗ | ✗ |
| 774 | MDock | DeepAffinity | ✗ | ✓ |
| 775 | MDock | DeepBindRG | ✗ | ✓ |
| 776 | MDock | Footprint Scoring | ✗ | ✗ |
| 777 | MDock | GOLD Scoring | ✗ | ✗ |
| 778 | MDock | Gnina Scoring | ✗ | ✓ |
| 779 | MDock | GoldScore | ✗ | ✗ |
| 780 | MDock | Grid-Based Scoring | ✗ | ✗ |
| 781 | MDock | Hawkins GB/SA Scoring | ✗ | ✗ |
| 782 | MDock | MDock | ✗ | ✗ |
| 783 | MDock | MM/GBSA Scoring | ✗ | ✗ |
| 784 | MDock | MM/PBSA Scoring | ✗ | ✗ |
| 785 | MDock | MultiGrid FPS Scoring | ✗ | ✗ |
| 786 | MDock | NNScore 2.0 | ✗ | ✓ |
| 787 | MDock | OpenBPMD | ✗ | ✗ |
| 788 | MDock | PB/SA Scoring | ✗ | ✗ |
| 789 | MDock | PLANTS Scoring | ✗ | ✗ |
| 790 | MDock | PLP | ✗ | ✗ |
| 791 | MDock | PLP95 | ✗ | ✗ |
| 792 | MDock | Pharmacophore Matching Similarity Scoring | ✗ | ✗ |
| 793 | MDock | RF-Score-VS | ✗ | ✗ |
| 794 | MDock | Rosetta Scoring | ✗ | ✗ |
| 795 | MDock | SASA Scoring | ✗ | ✗ |
| 796 | MDock | SEED Scoring | ✗ | ✗ |
| 797 | MDock | SMINA Scoring | ✗ | ✗ |
| 798 | MDock | Vina scoring | ✗ | ✗ |
| 799 | MDock | Vinardo Scoring | ✗ | ✗ |
| 800 | MDock | Zou GB/SA Scoring | ✗ | ✗ |
| 801 | MCDock | AD4 Scoring | ✗ | ✗ |
| 802 | MCDock | ASP Scoring | ✗ | ✗ |
| 803 | MCDock | Bump Filter Scoring | ✗ | ✗ |
| 804 | MCDock | ChemPLP | ✗ | ✗ |
| 805 | MCDock | ChemScore | ✗ | ✗ |
| 806 | MCDock | Contact Scoring | ✗ | ✗ |
| 807 | MCDock | Continuous Scoring | ✗ | ✗ |
| 808 | MCDock | DOCK 3.5 Scoring | ✗ | ✗ |
| 809 | MCDock | DeepAffinity | ✗ | ✓ |
| 810 | MCDock | DeepBindRG | ✗ | ✓ |
| 811 | MCDock | Footprint Scoring | ✗ | ✗ |
| 812 | MCDock | GOLD Scoring | ✗ | ✗ |
| 813 | MCDock | Gnina Scoring | ✗ | ✓ |
| 814 | MCDock | GoldScore | ✗ | ✗ |
| 815 | MCDock | Grid-Based Scoring | ✗ | ✗ |
| 816 | MCDock | Hawkins GB/SA Scoring | ✗ | ✗ |
| 817 | MCDock | MCDock | ✗ | ✗ |
| 818 | MCDock | MM/GBSA Scoring | ✗ | ✗ |
| 819 | MCDock | MM/PBSA Scoring | ✗ | ✗ |
| 820 | MCDock | MultiGrid FPS Scoring | ✗ | ✗ |
| 821 | MCDock | NNScore 2.0 | ✗ | ✓ |
| 822 | MCDock | OpenBPMD | ✗ | ✗ |
| 823 | MCDock | PB/SA Scoring | ✗ | ✗ |
| 824 | MCDock | PLANTS Scoring | ✗ | ✗ |

**Supplementary Table 1 (Continued). docking protocols Supported by the AFVS and AFU Modules.** Each docking protocol is determined by the combination of a pose prediction method and a scoring function. For both of these components, it is indicated whether they are based on deep learning (DL) or not. (Continued on next page.)

| Index | Pose Prediction/Sampling Method | Scoring Function | DL-Based Sampling | DL-Based Scoring |
|---|---|---|---|---|
| 825 | MCDock | PLP | ✗ | ✗ |
| 826 | MCDock | PLP95 | ✗ | ✗ |
| 827 | MCDock | Pharmacophore Matching Similarity Scoring | ✗ | ✗ |
| 828 | MCDock | RF-Score-VS | ✗ | ✗ |
| 829 | MCDock | Rosetta Scoring | ✗ | ✗ |
| 830 | MCDock | SASA Scoring | ✗ | ✗ |
| 831 | MCDock | SEED Scoring | ✗ | ✗ |
| 832 | MCDock | SMINA Scoring | ✗ | ✗ |
| 833 | MCDock | Vina scoring | ✗ | ✗ |
| 834 | MCDock | Vinardo Scoring | ✗ | ✗ |
| 835 | MCDock | Zou GB/SA Scoring | ✗ | ✗ |
| 836 | Molegro | AD4 Scoring | ✗ | ✗ |
| 837 | Molegro | ASP Scoring | ✗ | ✗ |
| 838 | Molegro | Bump Filter Scoring | ✗ | ✗ |
| 839 | Molegro | ChemPLP | ✗ | ✗ |
| 840 | Molegro | ChemScore | ✗ | ✗ |
| 841 | Molegro | Contact Scoring | ✗ | ✗ |
| 842 | Molegro | Continuous Scoring | ✗ | ✗ |
| 843 | Molegro | DOCK 3.5 Scoring | ✗ | ✗ |
| 844 | Molegro | DeepAffinity | ✗ | ✓ |
| 845 | Molegro | DeepBindRG | ✗ | ✓ |
| 846 | Molegro | Footprint Scoring | ✗ | ✗ |
| 847 | Molegro | GOLD Scoring | ✗ | ✗ |
| 848 | Molegro | Gnina Scoring | ✗ | ✓ |
| 849 | Molegro | GoldScore | ✗ | ✗ |
| 850 | Molegro | Grid-Based Scoring | ✗ | ✗ |
| 851 | Molegro | Hawkins GB/SA Scoring | ✗ | ✗ |
| 852 | Molegro | MM/GBSA Scoring | ✗ | ✗ |
| 853 | Molegro | MM/PBSA Scoring | ✗ | ✗ |
| 854 | Molegro | Molegro | ✗ | ✗ |
| 855 | Molegro | MultiGrid FPS Scoring | ✗ | ✗ |
| 856 | Molegro | NNScore 2.0 | ✗ | ✓ |
| 857 | Molegro | OpenBPMD | ✗ | ✗ |
| 858 | Molegro | PB/SA Scoring | ✗ | ✗ |
| 859 | Molegro | PLANTS Scoring | ✗ | ✗ |
| 860 | Molegro | PLP | ✗ | ✗ |
| 861 | Molegro | PLP95 | ✗ | ✗ |
| 862 | Molegro | Pharmacophore Matching Similarity Scoring | ✗ | ✗ |
| 863 | Molegro | RF-Score-VS | ✗ | ✗ |
| 864 | Molegro | Rosetta Scoring | ✗ | ✗ |
| 865 | Molegro | SASA Scoring | ✗ | ✗ |
| 866 | Molegro | SEED Scoring | ✗ | ✗ |
| 867 | Molegro | SMINA Scoring | ✗ | ✗ |
| 868 | Molegro | Vina scoring | ✗ | ✗ |
| 869 | Molegro | Vinardo Scoring | ✗ | ✗ |
| 870 | Molegro | Zou GB/SA Scoring | ✗ | ✗ |
| 871 | MpsDockzn | AD4 Scoring | ✗ | ✗ |
| 872 | MpsDockzn | ASP Scoring | ✗ | ✗ |
| 873 | MpsDockzn | Bump Filter Scoring | ✗ | ✗ |
| 874 | MpsDockzn | ChemPLP | ✗ | ✗ |
| 875 | MpsDockzn | ChemScore | ✗ | ✗ |
| 876 | MpsDockzn | Contact Scoring | ✗ | ✗ |
| 877 | MpsDockzn | Continuous Scoring | ✗ | ✗ |
| 878 | MpsDockzn | DOCK 3.5 Scoring | ✗ | ✗ |
| 879 | MpsDockzn | DeepAffinity | ✗ | ✓ |

**Supplementary Table 1 (Continued). docking protocols Supported by the AFVS and AFU Modules.** Each docking protocol is determined by the combination of a pose prediction method and a scoring function. For both of these components, it is indicated whether they are based on deep learning (DL) or not. (Continued on next page.)

| Index | Pose Prediction/Sampling Method | Scoring Function | DL-Based Sampling | DL-Based Scoring |
|---|---|---|---|---|
| 880 | MpsDockzn | DeepBindRG | ✗ | ✓ |
| 881 | MpsDockzn | Footprint Scoring | ✗ | ✗ |
| 882 | MpsDockzn | GOLD Scoring | ✗ | ✗ |
| 883 | MpsDockzn | Gnina Scoring | ✗ | ✓ |
| 884 | MpsDockzn | GoldScore | ✗ | ✗ |
| 885 | MpsDockzn | Grid-Based Scoring | ✗ | ✗ |
| 886 | MpsDockzn | Hawkins GB/SA Scoring | ✗ | ✗ |
| 887 | MpsDockzn | MM/GBSA Scoring | ✗ | ✗ |
| 888 | MpsDockzn | MM/PBSA Scoring | ✗ | ✗ |
| 889 | MpsDockzn | MpsDockzn | ✗ | ✗ |
| 890 | MpsDockzn | MultiGrid FPS Scoring | ✗ | ✗ |
| 891 | MpsDockzn | NNScore 2.0 | ✗ | ✓ |
| 892 | MpsDockzn | OpenBPMD | ✗ | ✗ |
| 893 | MpsDockzn | PB/SA Scoring | ✗ | ✗ |
| 894 | MpsDockzn | PLANTS Scoring | ✗ | ✗ |
| 895 | MpsDockzn | PLP | ✗ | ✗ |
| 896 | MpsDockzn | PLP95 | ✗ | ✗ |
| 897 | MpsDockzn | Pharmacophore Matching Similarity Scoring | ✗ | ✗ |
| 898 | MpsDockzn | RF-Score-VS | ✗ | ✗ |
| 899 | MpsDockzn | Rosetta Scoring | ✗ | ✗ |
| 900 | MpsDockzn | SASA Scoring | ✗ | ✗ |
| 901 | MpsDockzn | SEED Scoring | ✗ | ✗ |
| 902 | MpsDockzn | SMINA Scoring | ✗ | ✗ |
| 903 | MpsDockzn | Vina scoring | ✗ | ✗ |
| 904 | MpsDockzn | Vinardo Scoring | ✗ | ✗ |
| 905 | MpsDockzn | Zou GB/SA Scoring | ✗ | ✗ |
| 906 | PSOVina | AD4 Scoring | ✗ | ✗ |
| 907 | PSOVina | ASP Scoring | ✗ | ✗ |
| 908 | PSOVina | Bump Filter Scoring | ✗ | ✗ |
| 909 | PSOVina | ChemPLP | ✗ | ✗ |
| 910 | PSOVina | ChemScore | ✗ | ✗ |
| 911 | PSOVina | Contact Scoring | ✗ | ✗ |
| 912 | PSOVina | Continuous Scoring | ✗ | ✗ |
| 913 | PSOVina | DOCK 3.5 Scoring | ✗ | ✗ |
| 914 | PSOVina | DeepAffinity | ✗ | ✓ |
| 915 | PSOVina | DeepBindRG | ✗ | ✓ |
| 916 | PSOVina | Footprint Scoring | ✗ | ✗ |
| 917 | PSOVina | GOLD Scoring | ✗ | ✗ |
| 918 | PSOVina | Gnina Scoring | ✗ | ✓ |
| 919 | PSOVina | GoldScore | ✗ | ✗ |
| 920 | PSOVina | Grid-Based Scoring | ✗ | ✗ |
| 921 | PSOVina | Hawkins GB/SA Scoring | ✗ | ✗ |
| 922 | PSOVina | MM/GBSA Scoring | ✗ | ✗ |
| 923 | PSOVina | MM/PBSA Scoring | ✗ | ✗ |
| 924 | PSOVina | MultiGrid FPS Scoring | ✗ | ✗ |
| 925 | PSOVina | NNScore 2.0 | ✗ | ✓ |
| 926 | PSOVina | OpenBPMD | ✗ | ✗ |
| 927 | PSOVina | PB/SA Scoring | ✗ | ✗ |
| 928 | PSOVina | PLANTS Scoring | ✗ | ✗ |
| 929 | PSOVina | PLP | ✗ | ✗ |
| 930 | PSOVina | PLP95 | ✗ | ✗ |
| 931 | PSOVina | PSOVina | ✗ | ✗ |
| 932 | PSOVina | Pharmacophore Matching Similarity Scoring | ✗ | ✗ |
| 933 | PSOVina | RF-Score-VS | ✗ | ✗ |
| 934 | PSOVina | Rosetta Scoring | ✗ | ✗ |

**Supplementary Table 1 (Continued). docking protocols Supported by the AFVS and AFU Modules.** Each docking protocol is determined by the combination of a pose prediction method and a scoring function. For both of these components, it is indicated whether they are based on deep learning (DL) or not. (Continued on next page.)

| Index | Pose Prediction/Sampling Method | Scoring Function | DL-Based Sampling | DL-Based Scoring |
|---|---|---|---|---|
| 935 | PSOVina | SASA Scoring | ✗ | ✗ |
| 936 | PSOVina | SEED Scoring | ✗ | ✗ |
| 937 | PSOVina | SMINA Scoring | ✗ | ✗ |
| 938 | PSOVina | Vina scoring | ✗ | ✗ |
| 939 | PSOVina | Vinardo Scoring | ✗ | ✗ |
| 940 | PSOVina | Zou GB/SA Scoring | ✗ | ✗ |
| 941 | PLANTS | AD4 Scoring | ✗ | ✗ |
| 942 | PLANTS | ASP Scoring | ✗ | ✗ |
| 943 | PLANTS | Bump Filter Scoring | ✗ | ✗ |
| 944 | PLANTS | ChemPLP | ✗ | ✗ |
| 945 | PLANTS | ChemScore | ✗ | ✗ |
| 946 | PLANTS | Contact Scoring | ✗ | ✗ |
| 947 | PLANTS | Continuous Scoring | ✗ | ✗ |
| 948 | PLANTS | DOCK 3.5 Scoring | ✗ | ✗ |
| 949 | PLANTS | DeepAffinity | ✗ | ✓ |
| 950 | PLANTS | DeepBindRG | ✗ | ✓ |
| 951 | PLANTS | Footprint Scoring | ✗ | ✗ |
| 952 | PLANTS | GOLD Scoring | ✗ | ✗ |
| 953 | PLANTS | Gnina Scoring | ✗ | ✓ |
| 954 | PLANTS | GoldScore | ✗ | ✗ |
| 955 | PLANTS | Grid-Based Scoring | ✗ | ✗ |
| 956 | PLANTS | Hawkins GB/SA Scoring | ✗ | ✗ |
| 957 | PLANTS | MM/GBSA Scoring | ✗ | ✗ |
| 958 | PLANTS | MM/PBSA Scoring | ✗ | ✗ |
| 959 | PLANTS | MultiGrid FPS Scoring | ✗ | ✗ |
| 960 | PLANTS | NNScore 2.0 | ✗ | ✓ |
| 961 | PLANTS | OpenBPMD | ✗ | ✗ |
| 962 | PLANTS | PB/SA Scoring | ✗ | ✗ |
| 963 | PLANTS | PLANTS | ✗ | ✗ |
| 964 | PLANTS | PLANTS Scoring | ✗ | ✗ |
| 965 | PLANTS | PLP | ✗ | ✗ |
| 966 | PLANTS | PLP95 | ✗ | ✗ |
| 967 | PLANTS | Pharmacophore Matching Similarity Scoring | ✗ | ✗ |
| 968 | PLANTS | RF-Score-VS | ✗ | ✗ |
| 969 | PLANTS | Rosetta Scoring | ✗ | ✗ |
| 970 | PLANTS | SASA Scoring | ✗ | ✗ |
| 971 | PLANTS | SEED Scoring | ✗ | ✗ |
| 972 | PLANTS | SMINA Scoring | ✗ | ✗ |
| 973 | PLANTS | Vina scoring | ✗ | ✗ |
| 974 | PLANTS | Vinardo Scoring | ✗ | ✗ |
| 975 | PLANTS | Zou GB/SA Scoring | ✗ | ✗ |
| 976 | PI-LZerD | PI-LZerD | ✗ | ✗ |
| 977 | PIPER | PIPER | ✗ | ✗ |
| 978 | QuickVina 2 | AD4 Scoring | ✗ | ✗ |
| 979 | QuickVina 2 | ASP Scoring | ✗ | ✗ |
| 980 | QuickVina 2 | Bump Filter Scoring | ✗ | ✗ |
| 981 | QuickVina 2 | ChemPLP | ✗ | ✗ |
| 982 | QuickVina 2 | ChemScore | ✗ | ✗ |
| 983 | QuickVina 2 | Contact Scoring | ✗ | ✗ |
| 984 | QuickVina 2 | Continuous Scoring | ✗ | ✗ |
| 985 | QuickVina 2 | DOCK 3.5 Scoring | ✗ | ✗ |
| 986 | QuickVina 2 | DeepAffinity | ✗ | ✓ |
| 987 | QuickVina 2 | DeepBindRG | ✗ | ✓ |
| 988 | QuickVina 2 | Footprint Scoring | ✗ | ✗ |
| 989 | QuickVina 2 | GOLD Scoring | ✗ | ✗ |

**Supplementary Table 1 (Continued). docking protocols Supported by the AFVS and AFU Modules.** Each docking protocol is determined by the combination of a pose prediction method and a scoring function. For both of these components, it is indicated whether they are based on deep learning (DL) or not. (Continued on next page.)

| Index | Pose Prediction/Sampling Method | Scoring Function | DL-Based Sampling | DL-Based Scoring |
|---|---|---|---|---|
| 990 | QuickVina 2 | Gnina Scoring | ✗ | ✓ |
| 991 | QuickVina 2 | GoldScore | ✗ | ✗ |
| 992 | QuickVina 2 | Grid-Based Scoring | ✗ | ✗ |
| 993 | QuickVina 2 | Hawkins GB/SA Scoring | ✗ | ✗ |
| 994 | QuickVina 2 | MM/GBSA Scoring | ✗ | ✗ |
| 995 | QuickVina 2 | MM/PBSA Scoring | ✗ | ✗ |
| 996 | QuickVina 2 | MultiGrid FPS Scoring | ✗ | ✗ |
| 997 | QuickVina 2 | NNScore 2.0 | ✗ | ✓ |
| 998 | QuickVina 2 | OpenBPMD | ✗ | ✗ |
| 999 | QuickVina 2 | PB/SA Scoring | ✗ | ✗ |
| 1000 | QuickVina 2 | PLANTS Scoring | ✗ | ✗ |
| 1001 | QuickVina 2 | PLP | ✗ | ✗ |
| 1002 | QuickVina 2 | PLP95 | ✗ | ✗ |
| 1003 | QuickVina 2 | Pharmacophore Matching Similarity Scoring | ✗ | ✗ |
| 1004 | QuickVina 2 | QuickVina 2 | ✗ | ✗ |
| 1005 | QuickVina 2 | RF-Score-VS | ✗ | ✗ |
| 1006 | QuickVina 2 | Rosetta Scoring | ✗ | ✗ |
| 1007 | QuickVina 2 | SASA Scoring | ✗ | ✗ |
| 1008 | QuickVina 2 | SEED Scoring | ✗ | ✗ |
| 1009 | QuickVina 2 | SMINA Scoring | ✗ | ✗ |
| 1010 | QuickVina 2 | Vina scoring | ✗ | ✗ |
| 1011 | QuickVina 2 | Vinardo Scoring | ✗ | ✗ |
| 1012 | QuickVina 2 | Zou GB/SA Scoring | ✗ | ✗ |
| 1013 | QuickVina-W | AD4 Scoring | ✗ | ✗ |
| 1014 | QuickVina-W | ASP Scoring | ✗ | ✗ |
| 1015 | QuickVina-W | Bump Filter Scoring | ✗ | ✗ |
| 1016 | QuickVina-W | ChemPLP | ✗ | ✗ |
| 1017 | QuickVina-W | ChemScore | ✗ | ✗ |
| 1018 | QuickVina-W | Contact Scoring | ✗ | ✗ |
| 1019 | QuickVina-W | Continuous Scoring | ✗ | ✗ |
| 1020 | QuickVina-W | DOCK 3.5 Scoring | ✗ | ✗ |
| 1021 | QuickVina-W | DeepAffinity | ✗ | ✓ |
| 1022 | QuickVina-W | DeepBindRG | ✗ | ✓ |
| 1023 | QuickVina-W | Footprint Scoring | ✗ | ✗ |
| 1024 | QuickVina-W | GOLD Scoring | ✗ | ✗ |
| 1025 | QuickVina-W | Gnina Scoring | ✗ | ✓ |
| 1026 | QuickVina-W | GoldScore | ✗ | ✗ |
| 1027 | QuickVina-W | Grid-Based Scoring | ✗ | ✗ |
| 1028 | QuickVina-W | Hawkins GB/SA Scoring | ✗ | ✗ |
| 1029 | QuickVina-W | MM/GBSA Scoring | ✗ | ✗ |
| 1030 | QuickVina-W | MM/PBSA Scoring | ✗ | ✗ |
| 1031 | QuickVina-W | MultiGrid FPS Scoring | ✗ | ✗ |
| 1032 | QuickVina-W | NNScore 2.0 | ✗ | ✓ |
| 1033 | QuickVina-W | OpenBPMD | ✗ | ✗ |
| 1034 | QuickVina-W | PB/SA Scoring | ✗ | ✗ |
| 1035 | QuickVina-W | PLANTS Scoring | ✗ | ✗ |
| 1036 | QuickVina-W | PLP | ✗ | ✗ |
| 1037 | QuickVina-W | PLP95 | ✗ | ✗ |
| 1038 | QuickVina-W | Pharmacophore Matching Similarity Scoring | ✗ | ✗ |
| 1039 | QuickVina-W | QuickVina-W | ✗ | ✗ |
| 1040 | QuickVina-W | RF-Score-VS | ✗ | ✗ |
| 1041 | QuickVina-W | Rosetta Scoring | ✗ | ✗ |
| 1042 | QuickVina-W | SASA Scoring | ✗ | ✗ |
| 1043 | QuickVina-W | SEED Scoring | ✗ | ✗ |
| 1044 | QuickVina-W | SMINA Scoring | ✗ | ✗ |

**Supplementary Table 1 (Continued). docking protocols Supported by the AFVS and AFU Modules.** Each docking protocol is determined by the combination of a pose prediction method and a scoring function. For both of these components, it is indicated whether they are based on deep learning (DL) or not. (Continued on next page.)

| Index | Pose Prediction/Sampling Method | Scoring Function | DL-Based Sampling | DL-Based Scoring |
|---|---|---|---|---|
| 1045 | QuickVina-W | Vina scoring | ✗ | ✗ |
| 1046 | QuickVina-W | Vinardo Scoring | ✗ | ✗ |
| 1047 | QuickVina-W | Zou GB/SA Scoring | ✗ | ✗ |
| 1048 | QVina2-GPU | AD4 Scoring | ✗ | ✗ |
| 1049 | QVina2-GPU | ASP Scoring | ✗ | ✗ |
| 1050 | QVina2-GPU | Bump Filter Scoring | ✗ | ✗ |
| 1051 | QVina2-GPU | ChemPLP | ✗ | ✗ |
| 1052 | QVina2-GPU | ChemScore | ✗ | ✗ |
| 1053 | QVina2-GPU | Contact Scoring | ✗ | ✗ |
| 1054 | QVina2-GPU | Continuous Scoring | ✗ | ✗ |
| 1055 | QVina2-GPU | DOCK 3.5 Scoring | ✗ | ✗ |
| 1056 | QVina2-GPU | DeepAffinity | ✗ | ✓ |
| 1057 | QVina2-GPU | DeepBindRG | ✗ | ✓ |
| 1058 | QVina2-GPU | Footprint Scoring | ✗ | ✗ |
| 1059 | QVina2-GPU | GOLD Scoring | ✗ | ✗ |
| 1060 | QVina2-GPU | Gnina Scoring | ✗ | ✓ |
| 1061 | QVina2-GPU | GoldScore | ✗ | ✗ |
| 1062 | QVina2-GPU | Grid-Based Scoring | ✗ | ✗ |
| 1063 | QVina2-GPU | Hawkins GB/SA Scoring | ✗ | ✗ |
| 1064 | QVina2-GPU | MM/GBSA Scoring | ✗ | ✗ |
| 1065 | QVina2-GPU | MM/PBSA Scoring | ✗ | ✗ |
| 1066 | QVina2-GPU | MultiGrid FPS Scoring | ✗ | ✗ |
| 1067 | QVina2-GPU | NNScore 2.0 | ✗ | ✓ |
| 1068 | QVina2-GPU | OpenBPMD | ✗ | ✗ |
| 1069 | QVina2-GPU | PB/SA Scoring | ✗ | ✗ |
| 1070 | QVina2-GPU | PLANTS Scoring | ✗ | ✗ |
| 1071 | QVina2-GPU | PLP | ✗ | ✗ |
| 1072 | QVina2-GPU | PLP95 | ✗ | ✗ |
| 1073 | QVina2-GPU | Pharmacophore Matching Similarity Scoring | ✗ | ✗ |
| 1074 | QVina2-GPU | QVina2-GPU | ✗ | ✗ |
| 1075 | QVina2-GPU | RF-Score-VS | ✗ | ✗ |
| 1076 | QVina2-GPU | Rosetta Scoring | ✗ | ✗ |
| 1077 | QVina2-GPU | SASA Scoring | ✗ | ✗ |
| 1078 | QVina2-GPU | SEED Scoring | ✗ | ✗ |
| 1079 | QVina2-GPU | SMINA Scoring | ✗ | ✗ |
| 1080 | QVina2-GPU | Vina scoring | ✗ | ✗ |
| 1081 | QVina2-GPU | Vinardo Scoring | ✗ | ✗ |
| 1082 | QVina2-GPU | Zou GB/SA Scoring | ✗ | ✗ |
| 1083 | QVina2-W-GPU | AD4 Scoring | ✗ | ✗ |
| 1084 | QVina2-W-GPU | ASP Scoring | ✗ | ✗ |
| 1085 | QVina2-W-GPU | Bump Filter Scoring | ✗ | ✗ |
| 1086 | QVina2-W-GPU | ChemPLP | ✗ | ✗ |
| 1087 | QVina2-W-GPU | ChemScore | ✗ | ✗ |
| 1088 | QVina2-W-GPU | Contact Scoring | ✗ | ✗ |
| 1089 | QVina2-W-GPU | Continuous Scoring | ✗ | ✗ |
| 1090 | QVina2-W-GPU | DOCK 3.5 Scoring | ✗ | ✗ |
| 1091 | QVina2-W-GPU | DeepAffinity | ✗ | ✓ |
| 1092 | QVina2-W-GPU | DeepBindRG | ✗ | ✓ |
| 1093 | QVina2-W-GPU | Footprint Scoring | ✗ | ✗ |
| 1094 | QVina2-W-GPU | GOLD Scoring | ✗ | ✗ |
| 1095 | QVina2-W-GPU | Gnina Scoring | ✗ | ✓ |
| 1096 | QVina2-W-GPU | GoldScore | ✗ | ✗ |
| 1097 | QVina2-W-GPU | Grid-Based Scoring | ✗ | ✗ |
| 1098 | QVina2-W-GPU | Hawkins GB/SA Scoring | ✗ | ✗ |
| 1099 | QVina2-W-GPU | MM/GBSA Scoring | ✗ | ✗ |

**Supplementary Table 1 (Continued). docking protocols Supported by the AFVS and AFU Modules.** Each docking protocol is determined by the combination of a pose prediction method and a scoring function. For both of these components, it is indicated whether they are based on deep learning (DL) or not. (Continued on next page.)

| Index | Pose Prediction/Sampling Method | Scoring Function | DL-Based Sampling | DL-Based Scoring |
|---|---|---|---|---|
| 1100 | QVina2-W-GPU | MM/PBSA Scoring | ✗ | ✗ |
| 1101 | QVina2-W-GPU | MultiGrid FPS Scoring | ✗ | ✗ |
| 1102 | QVina2-W-GPU | NNScore 2.0 | ✗ | ✓ |
| 1103 | QVina2-W-GPU | OpenBPMD | ✗ | ✗ |
| 1104 | QVina2-W-GPU | PB/SA Scoring | ✗ | ✗ |
| 1105 | QVina2-W-GPU | PLANTS Scoring | ✗ | ✗ |
| 1106 | QVina2-W-GPU | PLP | ✗ | ✗ |
| 1107 | QVina2-W-GPU | PLP95 | ✗ | ✗ |
| 1108 | QVina2-W-GPU | Pharmacophore Matching Similarity Scoring | ✗ | ✗ |
| 1109 | QVina2-W-GPU | QVina2-W-GPU | ✗ | ✗ |
| 1110 | QVina2-W-GPU | RF-Score-VS | ✗ | ✗ |
| 1111 | QVina2-W-GPU | Rosetta Scoring | ✗ | ✗ |
| 1112 | QVina2-W-GPU | SASA Scoring | ✗ | ✗ |
| 1113 | QVina2-W-GPU | SEED Scoring | ✗ | ✗ |
| 1114 | QVina2-W-GPU | SMINA Scoring | ✗ | ✗ |
| 1115 | QVina2-W-GPU | Vina scoring | ✗ | ✗ |
| 1116 | QVina2-W-GPU | Vinardo Scoring | ✗ | ✗ |
| 1117 | QVina2-W-GPU | Zou GB/SA Scoring | ✗ | ✗ |
| 1118 | rDock | AD4 Scoring | ✗ | ✗ |
| 1119 | rDock | ASP Scoring | ✗ | ✗ |
| 1120 | rDock | Bump Filter Scoring | ✗ | ✗ |
| 1121 | rDock | ChemPLP | ✗ | ✗ |
| 1122 | rDock | ChemScore | ✗ | ✗ |
| 1123 | rDock | Contact Scoring | ✗ | ✗ |
| 1124 | rDock | Continuous Scoring | ✗ | ✗ |
| 1125 | rDock | DOCK 3.5 Scoring | ✗ | ✗ |
| 1126 | rDock | DeepAffinity | ✗ | ✓ |
| 1127 | rDock | DeepBindRG | ✗ | ✓ |
| 1128 | rDock | Footprint Scoring | ✗ | ✗ |
| 1129 | rDock | GOLD Scoring | ✗ | ✗ |
| 1130 | rDock | Gnina Scoring | ✗ | ✓ |
| 1131 | rDock | GoldScore | ✗ | ✗ |
| 1132 | rDock | Grid-Based Scoring | ✗ | ✗ |
| 1133 | rDock | Hawkins GB/SA Scoring | ✗ | ✗ |
| 1134 | rDock | MM/GBSA Scoring | ✗ | ✗ |
| 1135 | rDock | MM/PBSA Scoring | ✗ | ✗ |
| 1136 | rDock | MultiGrid FPS Scoring | ✗ | ✗ |
| 1137 | rDock | NNScore 2.0 | ✗ | ✓ |
| 1138 | rDock | OpenBPMD | ✗ | ✗ |
| 1139 | rDock | PB/SA Scoring | ✗ | ✗ |
| 1140 | rDock | PLANTS Scoring | ✗ | ✗ |
| 1141 | rDock | PLP | ✗ | ✗ |
| 1142 | rDock | PLP95 | ✗ | ✗ |
| 1143 | rDock | Pharmacophore Matching Similarity Scoring | ✗ | ✗ |
| 1144 | rDock | RF-Score-VS | ✗ | ✗ |
| 1145 | rDock | Rosetta Scoring | ✗ | ✗ |
| 1146 | rDock | SASA Scoring | ✗ | ✗ |
| 1147 | rDock | SEED Scoring | ✗ | ✗ |
| 1148 | rDock | SMINA Scoring | ✗ | ✗ |
| 1149 | rDock | Vina scoring | ✗ | ✗ |
| 1150 | rDock | Vinardo Scoring | ✗ | ✗ |
| 1151 | rDock | Zou GB/SA Scoring | ✗ | ✗ |
| 1152 | rDock | rDock | ✗ | ✗ |
| 1153 | Rosetta Ligand | AD4 Scoring | ✗ | ✗ |
| 1154 | Rosetta Ligand | ASP Scoring | ✗ | ✗ |

**Supplementary Table 1 (Continued). docking protocols Supported by the AFVS and AFU Modules.** Each docking protocol is determined by the combination of a pose prediction method and a scoring function. For both of these components, it is indicated whether they are based on deep learning (DL) or not. (Continued on next page.)

| Index | Pose Prediction/Sampling Method | Scoring Function | DL-Based Sampling | DL-Based Scoring |
|---|---|---|---|---|
| 1155 | Rosetta Ligand | Bump Filter Scoring | ✗ | ✗ |
| 1156 | Rosetta Ligand | ChemPLP | ✗ | ✗ |
| 1157 | Rosetta Ligand | ChemScore | ✗ | ✗ |
| 1158 | Rosetta Ligand | Contact Scoring | ✗ | ✗ |
| 1159 | Rosetta Ligand | Continuous Scoring | ✗ | ✗ |
| 1160 | Rosetta Ligand | DOCK 3.5 Scoring | ✗ | ✗ |
| 1161 | Rosetta Ligand | DeepAffinity | ✗ | ✓ |
| 1162 | Rosetta Ligand | DeepBindRG | ✗ | ✓ |
| 1163 | Rosetta Ligand | Footprint Scoring | ✗ | ✗ |
| 1164 | Rosetta Ligand | GOLD Scoring | ✗ | ✗ |
| 1165 | Rosetta Ligand | Gnina Scoring | ✗ | ✓ |
| 1166 | Rosetta Ligand | GoldScore | ✗ | ✗ |
| 1167 | Rosetta Ligand | Grid-Based Scoring | ✗ | ✗ |
| 1168 | Rosetta Ligand | Hawkins GB/SA Scoring | ✗ | ✗ |
| 1169 | Rosetta Ligand | MM/GBSA Scoring | ✗ | ✗ |
| 1170 | Rosetta Ligand | MM/PBSA Scoring | ✗ | ✗ |
| 1171 | Rosetta Ligand | MultiGrid FPS Scoring | ✗ | ✗ |
| 1172 | Rosetta Ligand | NNScore 2.0 | ✗ | ✓ |
| 1173 | Rosetta Ligand | OpenBPMD | ✗ | ✗ |
| 1174 | Rosetta Ligand | PB/SA Scoring | ✗ | ✗ |
| 1175 | Rosetta Ligand | PLANTS Scoring | ✗ | ✗ |
| 1176 | Rosetta Ligand | PLP | ✗ | ✗ |
| 1177 | Rosetta Ligand | PLP95 | ✗ | ✗ |
| 1178 | Rosetta Ligand | Pharmacophore Matching Similarity Scoring | ✗ | ✗ |
| 1179 | Rosetta Ligand | RF-Score-VS | ✗ | ✗ |
| 1180 | Rosetta Ligand | Rosetta Ligand | ✗ | ✗ |
| 1181 | Rosetta Ligand | SASA Scoring | ✗ | ✗ |
| 1182 | Rosetta Ligand | SEED Scoring | ✗ | ✗ |
| 1183 | Rosetta Ligand | SMINA Scoring | ✗ | ✗ |
| 1184 | Rosetta Ligand | Vina scoring | ✗ | ✗ |
| 1185 | Rosetta Ligand | Vinardo Scoring | ✗ | ✗ |
| 1186 | Rosetta Ligand | Zou GB/SA Scoring | ✗ | ✗ |
| 1187 | RLDOCK | AD4 Scoring | ✗ | ✗ |
| 1188 | RLDOCK | ASP Scoring | ✗ | ✗ |
| 1189 | RLDOCK | Bump Filter Scoring | ✗ | ✗ |
| 1190 | RLDOCK | ChemPLP | ✗ | ✗ |
| 1191 | RLDOCK | ChemScore | ✗ | ✗ |
| 1192 | RLDOCK | Contact Scoring | ✗ | ✗ |
| 1193 | RLDOCK | Continuous Scoring | ✗ | ✗ |
| 1194 | RLDOCK | DOCK 3.5 Scoring | ✗ | ✗ |
| 1195 | RLDOCK | DeepAffinity | ✗ | ✓ |
| 1196 | RLDOCK | DeepBindRG | ✗ | ✓ |
| 1197 | RLDOCK | Footprint Scoring | ✗ | ✗ |
| 1198 | RLDOCK | GOLD Scoring | ✗ | ✗ |
| 1199 | RLDOCK | Gnina Scoring | ✗ | ✓ |
| 1200 | RLDOCK | GoldScore | ✗ | ✗ |
| 1201 | RLDOCK | Grid-Based Scoring | ✗ | ✗ |
| 1202 | RLDOCK | Hawkins GB/SA Scoring | ✗ | ✗ |
| 1203 | RLDOCK | MM/GBSA Scoring | ✗ | ✗ |
| 1204 | RLDOCK | MM/PBSA Scoring | ✗ | ✗ |
| 1205 | RLDOCK | MultiGrid FPS Scoring | ✗ | ✗ |
| 1206 | RLDOCK | NNScore 2.0 | ✗ | ✓ |
| 1207 | RLDOCK | OpenBPMD | ✗ | ✗ |
| 1208 | RLDOCK | PB/SA Scoring | ✗ | ✗ |
| 1209 | RLDOCK | PLANTS Scoring | ✗ | ✗ |

**Supplementary Table 1 (Continued). docking protocols Supported by the AFVS and AFU Modules.** Each docking protocol is determined by the combination of a pose prediction method and a scoring function. For both of these components, it is indicated whether they are based on deep learning (DL) or not. (Continued on next page.)

| Index | Pose Prediction/Sampling Method | Scoring Function | DL-Based Sampling | DL-Based Scoring |
|---|---|---|---|---|
| 1210 | RLDOCK | PLP | ✗ | ✗ |
| 1211 | RLDOCK | PLP95 | ✗ | ✗ |
| 1212 | RLDOCK | Pharmacophore Matching Similarity Scoring | ✗ | ✗ |
| 1213 | RLDOCK | RF-Score-VS | ✗ | ✗ |
| 1214 | RLDOCK | RLDOCK | ✗ | ✗ |
| 1215 | RLDOCK | Rosetta Scoring | ✗ | ✗ |
| 1216 | RLDOCK | SASA Scoring | ✗ | ✗ |
| 1217 | RLDOCK | SEED Scoring | ✗ | ✗ |
| 1218 | RLDOCK | SMINA Scoring | ✗ | ✗ |
| 1219 | RLDOCK | Vina scoring | ✗ | ✗ |
| 1220 | RLDOCK | Vinardo Scoring | ✗ | ✗ |
| 1221 | RLDOCK | Zou GB/SA Scoring | ✗ | ✗ |
| 1222 | SEED | AD4 Scoring | ✗ | ✗ |
| 1223 | SEED | ASP Scoring | ✗ | ✗ |
| 1224 | SEED | Bump Filter Scoring | ✗ | ✗ |
| 1225 | SEED | ChemPLP | ✗ | ✗ |
| 1226 | SEED | ChemScore | ✗ | ✗ |
| 1227 | SEED | Contact Scoring | ✗ | ✗ |
| 1228 | SEED | Continuous Scoring | ✗ | ✗ |
| 1229 | SEED | DOCK 3.5 Scoring | ✗ | ✗ |
| 1230 | SEED | DeepAffinity | ✗ | ✓ |
| 1231 | SEED | DeepBindRG | ✗ | ✓ |
| 1232 | SEED | Footprint Scoring | ✗ | ✗ |
| 1233 | SEED | GOLD Scoring | ✗ | ✗ |
| 1234 | SEED | Gnina Scoring | ✗ | ✓ |
| 1235 | SEED | GoldScore | ✗ | ✗ |
| 1236 | SEED | Grid-Based Scoring | ✗ | ✗ |
| 1237 | SEED | Hawkins GB/SA Scoring | ✗ | ✗ |
| 1238 | SEED | MM/GBSA Scoring | ✗ | ✗ |
| 1239 | SEED | MM/PBSA Scoring | ✗ | ✗ |
| 1240 | SEED | MultiGrid FPS Scoring | ✗ | ✗ |
| 1241 | SEED | NNScore 2.0 | ✗ | ✓ |
| 1242 | SEED | OpenBPMD | ✗ | ✗ |
| 1243 | SEED | PB/SA Scoring | ✗ | ✗ |
| 1244 | SEED | PLANTS Scoring | ✗ | ✗ |
| 1245 | SEED | PLP | ✗ | ✗ |
| 1246 | SEED | PLP95 | ✗ | ✗ |
| 1247 | SEED | Pharmacophore Matching Similarity Scoring | ✗ | ✗ |
| 1248 | SEED | RF-Score-VS | ✗ | ✗ |
| 1249 | SEED | Rosetta Scoring | ✗ | ✗ |
| 1250 | SEED | SASA Scoring | ✗ | ✗ |
| 1251 | SEED | SEED | ✗ | ✗ |
| 1252 | SEED | SEED Scoring | ✗ | ✗ |
| 1253 | SEED | SMINA Scoring | ✗ | ✗ |
| 1254 | SEED | Vina scoring | ✗ | ✗ |
| 1255 | SEED | Vinardo Scoring | ✗ | ✗ |
| 1256 | SEED | Zou GB/SA Scoring | ✗ | ✗ |
| 1257 | SMINA | AD4 Scoring | ✗ | ✗ |
| 1258 | SMINA | ASP Scoring | ✗ | ✗ |
| 1259 | SMINA | Bump Filter Scoring | ✗ | ✗ |
| 1260 | SMINA | ChemPLP | ✗ | ✗ |
| 1261 | SMINA | ChemScore | ✗ | ✗ |
| 1262 | SMINA | Contact Scoring | ✗ | ✗ |
| 1263 | SMINA | Continuous Scoring | ✗ | ✗ |
| 1264 | SMINA | DOCK 3.5 Scoring | ✗ | ✗ |

**Supplementary Table 1 (Continued). docking protocols Supported by the AFVS and AFU Modules.** Each docking protocol is determined by the combination of a pose prediction method and a scoring function. For both of these components, it is indicated whether they are based on deep learning (DL) or not. (Continued on next page.)

| Index | Pose Prediction/Sampling Method | Scoring Function | DL-Based Sampling | DL-Based Scoring |
|---|---|---|---|---|
| 1265 | SMINA | DeepAffinity | ✗ | ✓ |
| 1266 | SMINA | DeepBindRG | ✗ | ✓ |
| 1267 | SMINA | Footprint Scoring | ✗ | ✗ |
| 1268 | SMINA | GOLD Scoring | ✗ | ✗ |
| 1269 | SMINA | Gnina Scoring | ✗ | ✓ |
| 1270 | SMINA | GoldScore | ✗ | ✗ |
| 1271 | SMINA | Grid-Based Scoring | ✗ | ✗ |
| 1272 | SMINA | Hawkins GB/SA Scoring | ✗ | ✗ |
| 1273 | SMINA | MM/GBSA Scoring | ✗ | ✗ |
| 1274 | SMINA | MM/PBSA Scoring | ✗ | ✗ |
| 1275 | SMINA | MultiGrid FPS Scoring | ✗ | ✗ |
| 1276 | SMINA | NNScore 2.0 | ✗ | ✓ |
| 1277 | SMINA | OpenBPMD | ✗ | ✗ |
| 1278 | SMINA | PB/SA Scoring | ✗ | ✗ |
| 1279 | SMINA | PLANTS Scoring | ✗ | ✗ |
| 1280 | SMINA | PLP | ✗ | ✗ |
| 1281 | SMINA | PLP95 | ✗ | ✗ |
| 1282 | SMINA | Pharmacophore Matching Similarity Scoring | ✗ | ✗ |
| 1283 | SMINA | RF-Score-VS | ✗ | ✗ |
| 1284 | SMINA | Rosetta Scoring | ✗ | ✗ |
| 1285 | SMINA | SASA Scoring | ✗ | ✗ |
| 1286 | SMINA | SEED Scoring | ✗ | ✗ |
| 1287 | SMINA | SMINA | ✗ | ✗ |
| 1288 | SMINA | SMINA Scoring | ✗ | ✗ |
| 1289 | SMINA | Vina scoring | ✗ | ✗ |
| 1290 | SMINA | Vinardo Scoring | ✗ | ✗ |
| 1291 | SMINA | Zou GB/SA Scoring | ✗ | ✗ |
| 1292 | TANKBind | AD4 Scoring | ✓ | ✗ |
| 1293 | TANKBind | ASP Scoring | ✓ | ✗ |
| 1294 | TANKBind | Bump Filter Scoring | ✓ | ✗ |
| 1295 | TANKBind | ChemPLP | ✓ | ✗ |
| 1296 | TANKBind | ChemScore | ✓ | ✗ |
| 1297 | TANKBind | Contact Scoring | ✓ | ✗ |
| 1298 | TANKBind | Continuous Scoring | ✓ | ✗ |
| 1299 | TANKBind | DOCK 3.5 Scoring | ✓ | ✗ |
| 1300 | TANKBind | DeepAffinity | ✓ | ✓ |
| 1301 | TANKBind | DeepBindRG | ✓ | ✓ |
| 1302 | TANKBind | Footprint Scoring | ✓ | ✗ |
| 1303 | TANKBind | GOLD Scoring | ✓ | ✗ |
| 1304 | TANKBind | Gnina Scoring | ✓ | ✓ |
| 1305 | TANKBind | GoldScore | ✓ | ✗ |
| 1306 | TANKBind | Grid-Based Scoring | ✓ | ✗ |
| 1307 | TANKBind | Hawkins GB/SA Scoring | ✓ | ✗ |
| 1308 | TANKBind | MM/GBSA Scoring | ✓ | ✗ |
| 1309 | TANKBind | MM/PBSA Scoring | ✓ | ✗ |
| 1310 | TANKBind | MultiGrid FPS Scoring | ✓ | ✗ |
| 1311 | TANKBind | NNScore 2.0 | ✓ | ✓ |
| 1312 | TANKBind | OpenBPMD | ✓ | ✗ |
| 1313 | TANKBind | PB/SA Scoring | ✓ | ✗ |
| 1314 | TANKBind | PLANTS Scoring | ✓ | ✗ |
| 1315 | TANKBind | PLP | ✓ | ✗ |
| 1316 | TANKBind | PLP95 | ✓ | ✗ |
| 1317 | TANKBind | Pharmacophore Matching Similarity Scoring | ✓ | ✗ |
| 1318 | TANKBind | RF-Score-VS | ✓ | ✗ |
| 1319 | TANKBind | Rosetta Scoring | ✓ | ✗ |

**Supplementary Table 1 (Continued). docking protocols Supported by the AFVS and AFU Modules.** Each docking protocol is determined by the combination of a pose prediction method and a scoring function. For both of these components, it is indicated whether they are based on deep learning (DL) or not. (Continued on next page.)

| Index | Pose Prediction/Sampling Method | Scoring Function | DL-Based Sampling | DL-Based Scoring |
|---|---|---|---|---|
| 1320 | TANKBind | SASA Scoring | ✓ | ✗ |
| 1321 | TANKBind | SEED Scoring | ✓ | ✗ |
| 1322 | TANKBind | SMINA Scoring | ✓ | ✗ |
| 1323 | TANKBind | TANKBind | ✓ | ✗ |
| 1324 | TANKBind | Vina scoring | ✓ | ✗ |
| 1325 | TANKBind | Vinardo Scoring | ✓ | ✗ |
| 1326 | TANKBind | Zou GB/SA Scoring | ✓ | ✗ |
| 1327 | VinaCarb | AD4 Scoring | ✓ | ✗ |
| 1328 | VinaCarb | ASP Scoring | ✓ | ✗ |
| 1329 | VinaCarb | Bump Filter Scoring | ✓ | ✗ |
| 1330 | VinaCarb | ChemPLP | ✓ | ✗ |
| 1331 | VinaCarb | ChemScore | ✓ | ✗ |
| 1332 | VinaCarb | Contact Scoring | ✓ | ✗ |
| 1333 | VinaCarb | Continuous Scoring | ✓ | ✗ |
| 1334 | VinaCarb | DOCK 3.5 Scoring | ✓ | ✗ |
| 1335 | VinaCarb | DeepAffinity | ✓ | ✓ |
| 1336 | VinaCarb | DeepBindRG | ✓ | ✓ |
| 1337 | VinaCarb | Footprint Scoring | ✓ | ✗ |
| 1338 | VinaCarb | GOLD Scoring | ✓ | ✗ |
| 1339 | VinaCarb | Gnina Scoring | ✓ | ✓ |
| 1340 | VinaCarb | GoldScore | ✓ | ✗ |
| 1341 | VinaCarb | Grid-Based Scoring | ✓ | ✗ |
| 1342 | VinaCarb | Hawkins GB/SA Scoring | ✓ | ✗ |
| 1343 | VinaCarb | MM/GBSA Scoring | ✓ | ✗ |
| 1344 | VinaCarb | MM/PBSA Scoring | ✓ | ✗ |
| 1345 | VinaCarb | MultiGrid FPS Scoring | ✓ | ✗ |
| 1346 | VinaCarb | NNScore 2.0 | ✓ | ✓ |
| 1347 | VinaCarb | OpenBPMD | ✓ | ✗ |
| 1348 | VinaCarb | PB/SA Scoring | ✓ | ✗ |
| 1349 | VinaCarb | PLANTS Scoring | ✓ | ✗ |
| 1350 | VinaCarb | PLP | ✓ | ✗ |
| 1351 | VinaCarb | PLP95 | ✓ | ✗ |
| 1352 | VinaCarb | Pharmacophore Matching Similarity Scoring | ✓ | ✗ |
| 1353 | VinaCarb | RF-Score-VS | ✓ | ✗ |
| 1354 | VinaCarb | Rosetta Scoring | ✓ | ✗ |
| 1355 | VinaCarb | SASA Scoring | ✓ | ✗ |
| 1356 | VinaCarb | SEED Scoring | ✓ | ✗ |
| 1357 | VinaCarb | SMINA Scoring | ✓ | ✗ |
| 1358 | VinaCarb | Vina scoring | ✓ | ✗ |
| 1359 | VinaCarb | VinaCarb | ✓ | ✗ |
| 1360 | VinaCarb | Vinardo Scoring | ✓ | ✗ |
| 1361 | VinaCarb | Zou GB/SA Scoring | ✓ | ✗ |
| 1362 | VinaXB16 | AD4 Scoring | ✓ | ✗ |
| 1363 | VinaXB16 | ASP Scoring | ✓ | ✗ |
| 1364 | VinaXB16 | Bump Filter Scoring | ✓ | ✗ |
| 1365 | VinaXB16 | ChemPLP | ✓ | ✗ |
| 1366 | VinaXB16 | ChemScore | ✓ | ✗ |
| 1367 | VinaXB16 | Contact Scoring | ✓ | ✗ |
| 1368 | VinaXB16 | Continuous Scoring | ✓ | ✗ |
| 1369 | VinaXB16 | DOCK 3.5 Scoring | ✓ | ✗ |
| 1370 | VinaXB16 | DeepAffinity | ✓ | ✓ |
| 1371 | VinaXB16 | DeepBindRG | ✓ | ✓ |
| 1372 | VinaXB16 | Footprint Scoring | ✓ | ✗ |
| 1373 | VinaXB16 | GOLD Scoring | ✓ | ✗ |
| 1374 | VinaXB16 | Gnina Scoring | ✓ | ✓ |

**Supplementary Table 1 (Continued). docking protocols Supported by the AFVS and AFU Modules.** Each docking protocol is determined by the combination of a pose prediction method and a scoring function. For both of these components, it is indicated whether they are based on deep learning (DL) or not. (Continued on next page.)

| Index | Pose Prediction/Sampling Method | Scoring Function | DL-Based Sampling | DL-Based Scoring |
|---|---|---|---|---|
| 1375 | VinaXB16 | GoldScore | ✓ | ✗ |
| 1376 | VinaXB16 | Grid-Based Scoring | ✓ | ✗ |
| 1377 | VinaXB16 | Hawkins GB/SA Scoring | ✓ | ✗ |
| 1378 | VinaXB16 | MM/GBSA Scoring | ✓ | ✗ |
| 1379 | VinaXB16 | MM/PBSA Scoring | ✓ | ✗ |
| 1380 | VinaXB16 | MultiGrid FPS Scoring | ✓ | ✗ |
| 1381 | VinaXB16 | NNScore 2.0 | ✓ | ✓ |
| 1382 | VinaXB16 | OpenBPMD | ✓ | ✗ |
| 1383 | VinaXB16 | PB/SA Scoring | ✓ | ✗ |
| 1384 | VinaXB16 | PLANTS Scoring | ✓ | ✗ |
| 1385 | VinaXB16 | PLP | ✓ | ✗ |
| 1386 | VinaXB16 | PLP95 | ✓ | ✗ |
| 1387 | VinaXB16 | Pharmacophore Matching Similarity Scoring | ✓ | ✗ |
| 1388 | VinaXB16 | RF-Score-VS | ✓ | ✗ |
| 1389 | VinaXB16 | Rosetta Scoring | ✓ | ✗ |
| 1390 | VinaXB16 | SASA Scoring | ✓ | ✗ |
| 1391 | VinaXB16 | SEED Scoring | ✓ | ✗ |
| 1392 | VinaXB16 | SMINA Scoring | ✓ | ✗ |
| 1393 | VinaXB16 | Vina scoring | ✓ | ✗ |
| 1394 | VinaXB16 | VinaXB16 | ✓ | ✗ |
| 1395 | VinaXB16 | Vinardo Scoring | ✓ | ✗ |
| 1396 | VinaXB16 | Zou GB/SA Scoring | ✓ | ✗ |
| 1397 | Vina-GPU | AD4 Scoring | ✓ | ✗ |
| 1398 | Vina-GPU | ASP Scoring | ✓ | ✗ |
| 1399 | Vina-GPU | Bump Filter Scoring | ✓ | ✗ |
| 1400 | Vina-GPU | ChemPLP | ✓ | ✗ |
| 1401 | Vina-GPU | ChemScore | ✓ | ✗ |
| 1402 | Vina-GPU | Contact Scoring | ✓ | ✗ |
| 1403 | Vina-GPU | Continuous Scoring | ✓ | ✗ |
| 1404 | Vina-GPU | DOCK 3.5 Scoring | ✓ | ✗ |
| 1405 | Vina-GPU | DeepAffinity | ✓ | ✓ |
| 1406 | Vina-GPU | DeepBindRG | ✓ | ✓ |
| 1407 | Vina-GPU | Footprint Scoring | ✓ | ✗ |
| 1408 | Vina-GPU | GOLD Scoring | ✓ | ✗ |
| 1409 | Vina-GPU | Gnina Scoring | ✓ | ✓ |
| 1410 | Vina-GPU | GoldScore | ✓ | ✗ |
| 1411 | Vina-GPU | Grid-Based Scoring | ✓ | ✗ |
| 1412 | Vina-GPU | Hawkins GB/SA Scoring | ✓ | ✗ |
| 1413 | Vina-GPU | MM/GBSA Scoring | ✓ | ✗ |
| 1414 | Vina-GPU | MM/PBSA Scoring | ✓ | ✗ |
| 1415 | Vina-GPU | MultiGrid FPS Scoring | ✓ | ✗ |
| 1416 | Vina-GPU | NNScore 2.0 | ✓ | ✓ |
| 1417 | Vina-GPU | OpenBPMD | ✓ | ✗ |
| 1418 | Vina-GPU | PB/SA Scoring | ✓ | ✗ |
| 1419 | Vina-GPU | PLANTS Scoring | ✓ | ✗ |
| 1420 | Vina-GPU | PLP | ✓ | ✗ |
| 1421 | Vina-GPU | PLP95 | ✓ | ✗ |
| 1422 | Vina-GPU | Pharmacophore Matching Similarity Scoring | ✓ | ✗ |
| 1423 | Vina-GPU | RF-Score-VS | ✓ | ✗ |
| 1424 | Vina-GPU | Rosetta Scoring | ✓ | ✗ |
| 1425 | Vina-GPU | SASA Scoring | ✓ | ✗ |
| 1426 | Vina-GPU | SEED Scoring | ✓ | ✗ |
| 1427 | Vina-GPU | SMINA Scoring | ✓ | ✗ |
| 1428 | Vina-GPU | Vina scoring | ✓ | ✗ |
| 1429 | Vina-GPU | Vina-GPU | ✓ | ✗ |

**Supplementary Table 1 (Continued). docking protocols Supported by the AFVS and AFU Modules.** Each docking protocol is determined by the combination of a pose prediction method and a scoring function. For both of these components, it is indicated whether they are based on deep learning (DL) or not. (Continued on next page.)

| Index | Pose Prediction/Sampling Method | Scoring Function | DL-Based Sampling | DL-Based Scoring |
|---|---|---|---|---|
| 1430 | Vina-GPU | Vinardo Scoring | ✓ | ✗ |
| 1431 | Vina-GPU | Zou GB/SA Scoring | ✓ | ✗ |
| 1432 | Vina-GPU-2.0 | AD4 Scoring | ✓ | ✗ |
| 1433 | Vina-GPU-2.0 | ASP Scoring | ✓ | ✗ |
| 1434 | Vina-GPU-2.0 | Bump Filter Scoring | ✓ | ✗ |
| 1435 | Vina-GPU-2.0 | ChemPLP | ✓ | ✗ |
| 1436 | Vina-GPU-2.0 | ChemScore | ✓ | ✗ |
| 1437 | Vina-GPU-2.0 | Contact Scoring | ✓ | ✗ |
| 1438 | Vina-GPU-2.0 | Continuous Scoring | ✓ | ✗ |
| 1439 | Vina-GPU-2.0 | DOCK 3.5 Scoring | ✓ | ✗ |
| 1440 | Vina-GPU-2.0 | DeepAffinity | ✓ | ✓ |
| 1441 | Vina-GPU-2.0 | DeepBindRG | ✓ | ✓ |
| 1442 | Vina-GPU-2.0 | Footprint Scoring | ✓ | ✗ |
| 1443 | Vina-GPU-2.0 | GOLD Scoring | ✓ | ✗ |
| 1444 | Vina-GPU-2.0 | Gnina Scoring | ✓ | ✓ |
| 1445 | Vina-GPU-2.0 | GoldScore | ✓ | ✗ |
| 1446 | Vina-GPU-2.0 | Grid-Based Scoring | ✓ | ✗ |
| 1447 | Vina-GPU-2.0 | Hawkins GB/SA Scoring | ✓ | ✗ |
| 1448 | Vina-GPU-2.0 | MM/GBSA Scoring | ✓ | ✗ |
| 1449 | Vina-GPU-2.0 | MM/PBSA Scoring | ✓ | ✗ |
| 1450 | Vina-GPU-2.0 | MultiGrid FPS Scoring | ✓ | ✗ |
| 1451 | Vina-GPU-2.0 | NNScore 2.0 | ✓ | ✓ |
| 1452 | Vina-GPU-2.0 | OpenBPMD | ✓ | ✗ |
| 1453 | Vina-GPU-2.0 | PB/SA Scoring | ✓ | ✗ |
| 1454 | Vina-GPU-2.0 | PLANTS Scoring | ✓ | ✗ |
| 1455 | Vina-GPU-2.0 | PLP | ✓ | ✗ |
| 1456 | Vina-GPU-2.0 | PLP95 | ✓ | ✗ |
| 1457 | Vina-GPU-2.0 | Pharmacophore Matching Similarity Scoring | ✓ | ✗ |
| 1458 | Vina-GPU-2.0 | RF-Score-VS | ✓ | ✗ |
| 1459 | Vina-GPU-2.0 | Rosetta Scoring | ✓ | ✗ |
| 1460 | Vina-GPU-2.0 | SASA Scoring | ✓ | ✗ |
| 1461 | Vina-GPU-2.0 | SEED Scoring | ✓ | ✗ |
| 1462 | Vina-GPU-2.0 | SMINA Scoring | ✓ | ✗ |
| 1463 | Vina-GPU-2.0 | Vina scoring | ✓ | ✗ |
| 1464 | Vina-GPU-2.0 | Vina-GPU-2.0 | ✓ | ✗ |
| 1465 | Vina-GPU-2.0 | Vinardo Scoring | ✓ | ✗ |
| 1466 | Vina-GPU-2.0 | Zou GB/SA Scoring | ✓ | ✗ |
| 1467 | EnzyDock | AD4 Scoring | ✓ | ✗ |
| 1468 | EnzyDock | ASP Scoring | ✓ | ✗ |
| 1469 | EnzyDock | Bump Filter Scoring | ✓ | ✗ |
| 1470 | EnzyDock | ChemPLP | ✓ | ✗ |
| 1471 | EnzyDock | ChemScore | ✓ | ✗ |
| 1472 | EnzyDock | Contact Scoring | ✓ | ✗ |
| 1473 | EnzyDock | Continuous Scoring | ✓ | ✗ |
| 1474 | EnzyDock | DOCK 3.5 Scoring | ✓ | ✗ |
| 1475 | EnzyDock | DeepAffinity | ✓ | ✓ |
| 1476 | EnzyDock | DeepBindRG | ✓ | ✓ |
| 1477 | EnzyDock | Footprint Scoring | ✓ | ✗ |
| 1478 | EnzyDock | GOLD Scoring | ✓ | ✗ |
| 1479 | EnzyDock | Gnina Scoring | ✓ | ✓ |
| 1480 | EnzyDock | GoldScore | ✓ | ✗ |
| 1481 | EnzyDock | Grid-Based Scoring | ✓ | ✗ |
| 1482 | EnzyDock | Hawkins GB/SA Scoring | ✓ | ✗ |
| 1483 | EnzyDock | MM/GBSA Scoring | ✓ | ✗ |

**Supplementary Table 1 (Continued). docking protocols Supported by the AFVS and AFU Modules.** Each docking protocol is determined by the combination of a pose prediction method and a scoring function. For both of these components, it is indicated whether they are based on deep learning (DL) or not. (Continued on next page.)

| Index | Pose Prediction/Sampling Method | Scoring Function | DL-Based Sampling | DL-Based Scoring |
|---|---|---|---|---|
| 1484 | EnzyDock | MM/PBSA Scoring | ✓ | ✗ |
| 1485 | EnzyDock | MultiGrid FPS Scoring | ✓ | ✗ |
| 1486 | EnzyDock | NNScore 2.0 | ✓ | ✓ |
| 1487 | EnzyDock | OpenBPMD | ✓ | ✗ |
| 1488 | EnzyDock | PB/SA Scoring | ✓ | ✗ |
| 1489 | EnzyDock | PLANTS Scoring | ✓ | ✗ |
| 1490 | EnzyDock | PLP | ✓ | ✗ |
| 1491 | EnzyDock | PLP95 | ✓ | ✗ |
| 1492 | EnzyDock | Pharmacophore Matching Similarity Scoring | ✓ | ✗ |
| 1493 | EnzyDock | RF-Score-VS | ✓ | ✗ |
| 1494 | EnzyDock | Rosetta Scoring | ✓ | ✗ |
| 1495 | EnzyDock | SASA Scoring | ✓ | ✗ |
| 1496 | EnzyDock | SEED Scoring | ✓ | ✗ |
| 1497 | EnzyDock | SMINA Scoring | ✓ | ✗ |
| 1498 | EnzyDock | Vina scoring | ✓ | ✗ |
| 1499 | EnzyDock | Vinardo Scoring | ✓ | ✗ |
| 1500 | EnzyDock | Zou GB/SA Scoring | ✓ | ✗ |

**Supplementary Table 1 (Continued). docking protocols Supported by the AFVS and AFU Modules.** Each docking protocol is determined by the combination of a pose prediction method and a scoring function. For both of these components, it is indicated whether they are based on deep learning (DL) or not.

#### B Supported Output File Formats by AFLP

| Index | File Extension | Proper Name | Preparation Method |
|---|---|---|---|
| 1 | acesin | ACES input format | Open Babel |
| 2 | adf | ADF cartesian input format | Open Babel |
| 3 | alc | Alchemy format | Open Babel |
| 4 | ascii | ASCII format | Open Babel |
| 5 | bgf | MSI BGF format | Open Babel |
| 6 | box | Dock 3.5 Box format | Open Babel |
| 7 | bs | Ball and Stick format | Open Babel |
| 8 | c3d1 | Chem3D Cartesian 1 format | Open Babel |
| 9 | c3d2 | Chem3D Cartesian 2 format | Open Babel |
| 10 | cac | CAChe MolStruct format | Open Babel |
| 11 | caccrt | Cacao Cartesian format | Open Babel |
| 12 | cache | CAChe MolStruct format | Open Babel |
| 13 | cacint | Cacao Internal format | Open Babel |
| 14 | can | Canonical SMILES format | Open Babel |
| 15 | cdjson | ChemDoodle JSON | Open Babel |
| 16 | cdxml | ChemDraw CDXML format | Open Babel |
| 17 | cht | Chemtool format | Open Babel |
| 18 | cif | Crystallographic Information File | Open Babel |
| 19 | cml | Chemical Markup Language | Open Babel |
| 20 | cmlr | CML Reaction format | Open Babel |
| 21 | com | Gaussian 98/03 Input | Open Babel |
| 22 | CONFIG | DL-POLY CONFIG | Open Babel |
| 23 | CONTCAR | VASP format | Open Babel |
| 24 | CONTFF | MDFF format | Open Babel |
| 25 | crk2d | Chemical Resource Kit diagram(2D) | Open Babel |
| 26 | crk3d | Chemical Resource Kit 3D format | Open Babel |
| 27 | csr | Accelrys/MSI Quanta CSR format | Open Babel |
| 28 | cssr | CSD CSSR format | Open Babel |
| 29 | ct | ChemDraw Connection Table format | Open Babel |
| 30 | dalmol | DALTON input format | Open Babel |
| 31 | dmol | DMol3 coordinates format | Open Babel |
| 32 | ent | Protein Data Bank format | Open Babel |
| 33 | exyz | Extended XYZ cartesian coordinates format | Open Babel |
| 34 | fa | FASTA format | Open Babel |
| 35 | fasta | FASTA format | Open Babel |
| 36 | feat | Feature format | Open Babel |
| 37 | fh | Fenske-Hall Z-Matrix format | Open Babel |
| 38 | fhiaims | FHIaims XYZ format | Open Babel |
| 39 | fix | SMILES FIX format | Open Babel |
| 40 | fps | FPS text fingerprint format (Dalke) | Open Babel |
| 41 | fpt | Fingerprint format | Open Babel |
| 42 | fract | Free Form Fractional format | Open Babel |
| 43 | fs | Fastsearch format | Open Babel |
| 44 | fsa | FASTA format | Open Babel |
| 45 | gamin | GAMESS Input | Open Babel |
| 46 | gau | Gaussian 98/03 Input | Open Babel |
| 47 | gjc | Gaussian 98/03 Input | Open Babel |
| 48 | gjf | Gaussian 98/03 Input | Open Babel |
| 49 | gpr | Ghemical format | Open Babel |
| 50 | gr96 | GROMOS96 format | Open Babel |
| 51 | gro | GRO format | Open Babel |
| 52 | gukin | GAMESS-UK Input | Open Babel |
| 53 | gzmat | Gaussian Z-Matrix Input | Open Babel |
| 54 | hin | HyperChem HIN format | Open Babel |
| 55 | inchi | InChI format | Open Babel |

**Supplementary Table 2. Supported Chemical File Formats by AFLP for Prepared Output Ligand Library.** AFLP can store the prepared output ligand library in one or more output formats. The table lists the supported formats as well as the preparation method that AFLP uses.

| Index | File Extension | Proper Name | Preparation Method |
|---|---|---|---|
| 56 | inchikey | InChIKey | Open Babel |
| 57 | inp | GAMESS Input | Open Babel |
| 58 | jin | Jaguar input format | Open Babel |
| 59 | lmpdat | The LAMMPS data format | Open Babel |
| 60 | mcdl | MCDL format | Open Babel |
| 61 | mcif | Macromolecular Crystallographic Info | Open Babel |
| 62 | MDFF | MDFF format | Open Babel |
| 63 | mdl | MDL MOL format | Open Babel |
| 64 | ml2 | Sybyl Mol2 format | Open Babel |
| 65 | mmcif | Macromolecular Crystallographic Info | Open Babel |
| 66 | mmd | MacroModel format | Open Babel |
| 67 | mmod | MacroModel format | Open Babel |
| 68 | mna | Multilevel Neighborhoods of Atoms (MNA) | Open Babel |
| 69 | mol | MDL MOL format | Open Babel |
| 70 | mol2 | Sybyl Mol2 format | Open Babel |
| 71 | mold | Molden format | Open Babel |
| 72 | molden | Molden format | Open Babel |
| 73 | molf | Molden format | Open Babel |
| 74 | molreport | Open Babel molecule report | Open Babel |
| 75 | mop | MOPAC Cartesian format | Open Babel |
| 76 | mopcrt | MOPAC Cartesian format | Open Babel |
| 77 | mopin | MOPAC Internal | Open Babel |
| 78 | mp | Molpro input format | Open Babel |
| 79 | mpc | MOPAC Cartesian format | Open Babel |
| 80 | mpd | MolPrint2D format | Open Babel |
| 81 | mpqcin | MPQC simplified input format | Open Babel |
| 82 | mrv | Chemical Markup Language | Open Babel |
| 83 | msms | M.F. Sanner’s MSMS input format | Open Babel |
| 84 | nw | NWChem input format | Open Babel |
| 85 | orcainp | ORCA input format | Open Babel |
| 86 | outmol | DMol3 coordinates format | Open Babel |
| 87 | paint | Painter format | Open Babel |
| 88 | pcjson | PubChem JSON | Open Babel |
| 89 | pcm | PCModel Format | Open Babel |
| 90 | pdb | Protein Data Bank format | Open Babel |
| 91 | pdbqt | AutoDock PDBQT format | Open Babel |
| 92 | png | PNG 2D depiction | Open Babel |
| 93 | pointcloud | Point cloud on VDW surface | Open Babel |
| 94 | POSCAR | VASP format | Open Babel |
| 95 | POSFF | MDFF format | Open Babel |
| 96 | pov | POV-Ray input format | Open Babel |
| 97 | pqr | PQR format | Open Babel |
| 98 | pqs | Parallel Quantum Solutions format | Open Babel |
| 99 | qcin | Q-Chem input format | Open Babel |
| 100 | report | Open Babel report format | Open Babel |
| 101 | rsmi | Reaction SMILES format | Open Babel |
| 102 | rxn | MDL RXN format | Open Babel |
| 103 | sd | MDL MOL format | Open Babel |
| 104 | sdf | MDL MOL format | Open Babel |
| 105 | selfies | SELFIES format | SELFIES Package |
| 106 | smi | SMILES format | Open Babel |
| 107 | smi | SMILES format | AdaptiveFlow internal |
| 108 | stl | STL 3D-printing format | Open Babel |
| 109 | svg | SVG 2D depiction | Open Babel |
| 110 | sy2 | Sybyl Mol2 format | Open Babel |

**Supplementary Table 2 (Continued). Supported Chemical File Formats by AFLP for Prepared Output Ligand Library.** AFLP can store the prepared output ligand library in one or more output formats. The table lists the supported formats as well as the preparation method that AFLP uses.

| Index | File Extension | Proper Name | Preparation Method |
|---|---|---|---|
| 111 | therm | Thermo format | Open Babel |
| 112 | tmol | TurboMole Coordinate format | Open Babel |
| 113 | txt | Title format | Open Babel |
| 114 | txyz | Tinker XYZ format | Open Babel |
| 115 | unixyz | UniChem XYZ format | Open Babel |
| 116 | vmol | ViewMol format | Open Babel |
| 117 | xed | XED format | Open Babel |
| 118 | xyz | XYZ cartesian coordinates format | Open Babel |
| 119 | yob | YASARA.org YOB format | Open Babel |
| 120 | zin | ZINDO input format | Open Babel |

**Supplementary Table 2 (Continued). Supported Chemical File Formats by AFLP for Prepared Output Ligand Libraries.** AFLP can store the prepared output ligand library in one or more output formats. The table lists the supported formats as well as the preparation method that AFLP uses.

### C AdaptiveFlow File and Folder Structures

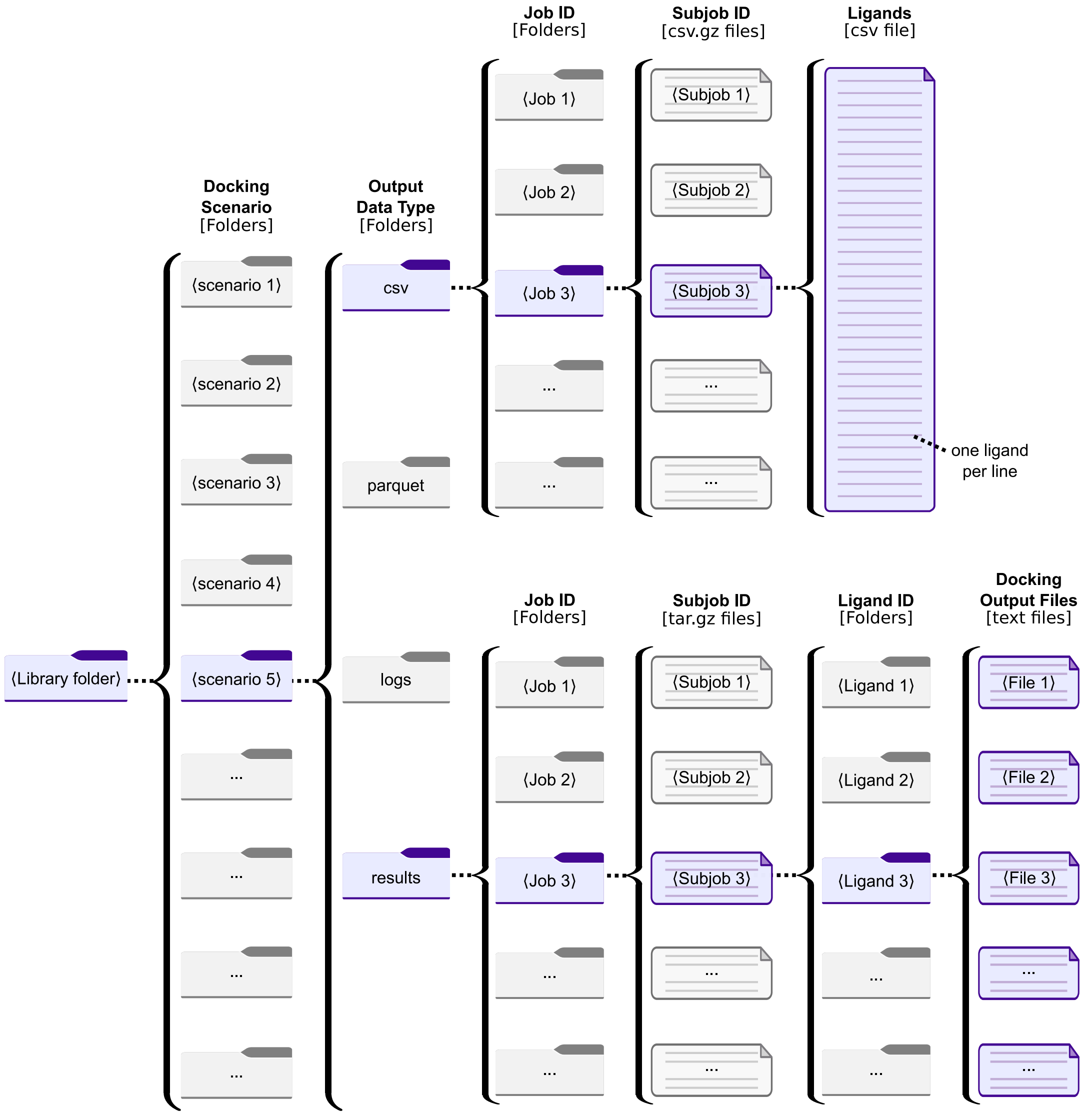

**Supplementary Fig. 1. AFVS Output Library Structure (Standard Format 1).** One of the two output data formats of AFVS is the *standard format 1*. Within the library root folder, each docking scenario has its own folder, containing subfolders for the different types of output files. The results subfolder contains the docking poses and other output files generated by the docking program. The csv and parquet subfolders contain the summary information on the dockings, such as the docking scores, in the csv and parquet formats respectively. The logs subfolder (if enabled) contains log files. Each of these folders contains subfolders for each batchsystem job, and each of them subsequently contains a file for each subjob. In the case of the parquet and the csv folders for the output data type, these files are parquet and gzipped csv files respectively. In the case of the log and the results files, these files are tar.gz files that contain folders for each ligand processed by the subjob.

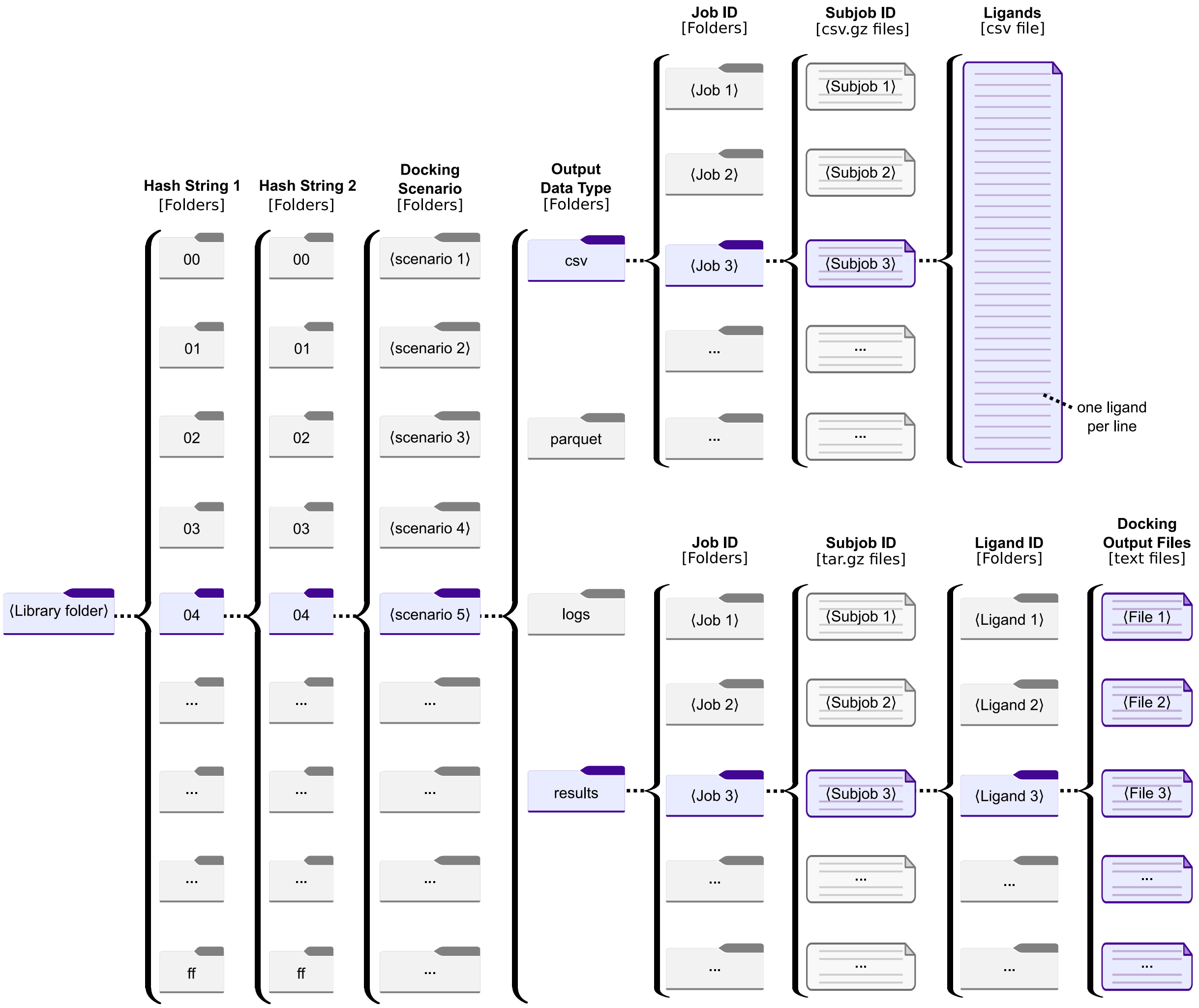

**Supplementary Fig. 2. AFVS Output Library Structure (Hash Format 2).** One of the two output data formats of AFVS is the *hash format 2*. The hash format 2 is principally the same format as the standard format 1, except that two hash subfolders are used directly within the library root folder. The hash folders each consist of two alphanumeric characters, and allow for a balanced distribution of the I/O load during highly parallel workflows, as well as allow to reduce the number of collections files per tranche folder. The hash prefixes are based on the collection and tranche IDs.

#### D AFLP Workflow

A single subjob for AFVS is defined by a set of *collections* to process and the input parameters, which describe the processing steps that should be followed for each ligand that is part of the input collections. A collection contains generally approximately 1,000 ligands and a single subjob will contain one or more of these collections. Each ligand proceeds through several steps based on the configuration that was provided for the job. Supplementary Fig. 7 shows the primary steps, with additional descriptions of each step provided below.

- **Desalting.**
- **Neutralization.**
- **Stereoisomer enumeration.** This process can generate multiple chemical species, so each of these species will process the next steps individually. In the output ligand names, a `_S<index>` will be appended to the original ligand name.
- **Tautomer enumeration.** This process can generate multiple chemical species, so each of these species will process the next steps individually. In the output ligand names, a `_T<index>` will be appended to the ligand name.
- **Attribute generation.** If requested as part of the configuration, various attributes of a ligand can be calculated within AFLP. These attributes can be generated from Open Babel, CxCalc, and RDKit.
- **Protonation state prediction.**
- **3D coordinate generation.** Up until this step, the SMILES format of the ligand has been used for processing. This step will generate a PDB file that will be used for the subsequent steps and target format generation.
- **Energy Check.** The potential energy of the ligand is calculated using Open Babel, and any ligands that exceed the configurable max energy value are discarded.
- **Target format generation.** Each target format is generated by Open Babel from the PDB that was generated as part of the previous step. Any attributes that were calculated as part of the process will be provided as comments in the PDB, PDBQT, SDF, and MOL2 formats.

After all of the ligands within a collection have been processed, AFLP will generate a summary JSON file, and archives for each of the target formats. The summary file provides information on computational timing, logging of decisions made by AFLP, and a summary of the attributes that were calculated for each of the output ligands. It also has the ability to capture intermediate logs for additional analysis. A workflow diagram of AFVS is shown in Supplementary Data Fig. 3.

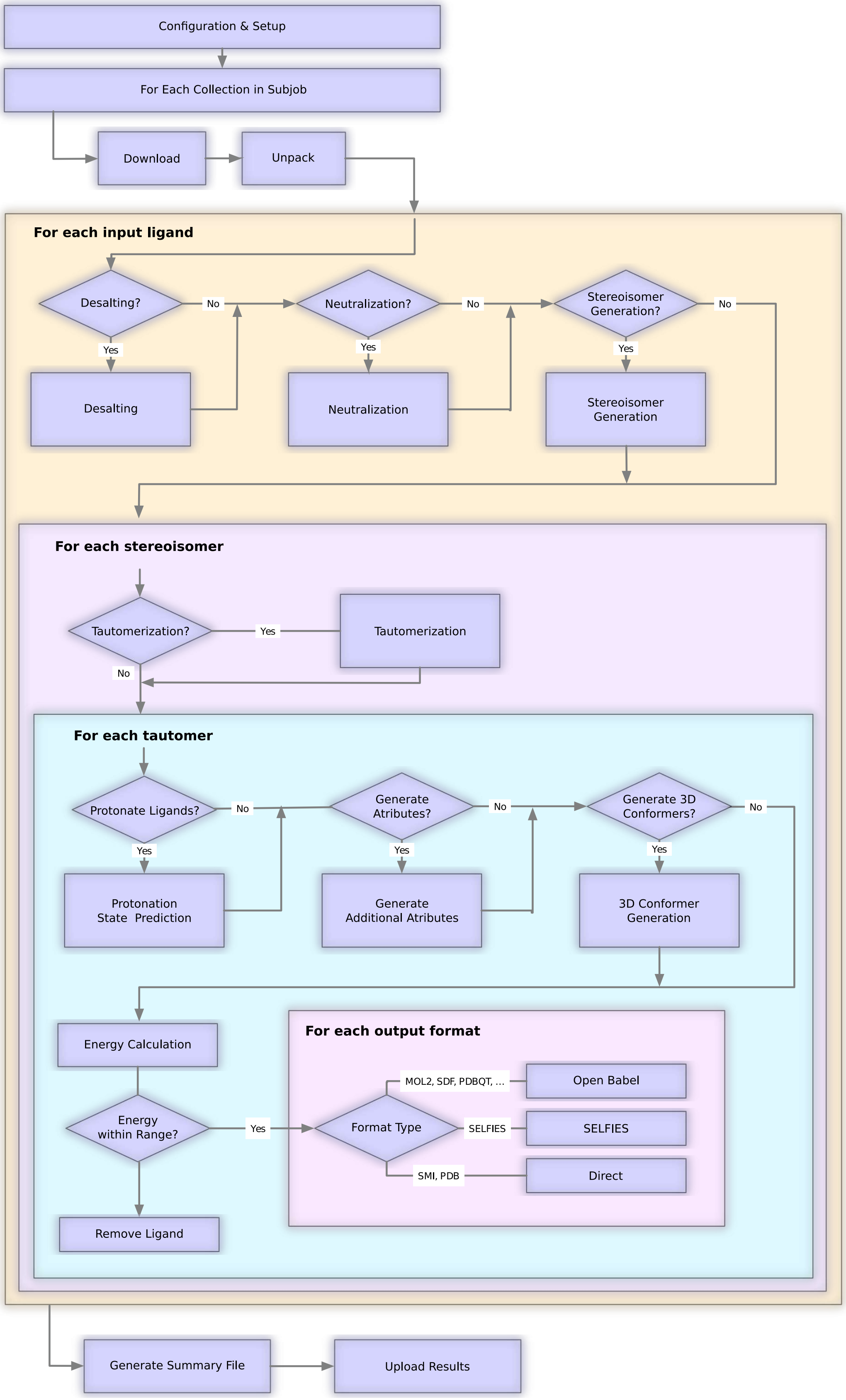

**Supplementary Fig. 3. Overview of the AFLP Subjob Procedure.** AdaptiveFlow uses a multi-process model for each subjob where work is processed via a set of submodules (implemented as processes) through Python queues. This model allows for parallelism within Python, while avoiding the Python Global Interpreter Lock (GIL) that can otherwise prevent efficient parallelism.

#### E AFVS Workflow

A single subjob for AFVS is defined by a set of *collections* to process and the input parameters, which describe the docking steps that should be applied for each ligand that is part of the input collections. A collection is generally approximately 1,000 ligands and a single subjob will contain one or more of these collections. Supplementary Fig. 4 shows the subjob modules.

- **Download.** This module downloads the candidate ligand collections from either a shared filesystem or from object storage.
- **Unpack.** This module expands the compressed archive to split out the individual candidate ligands within the collection
- **Collection Processing.** This module will determine which ligands of a collection should be processed. In the case of a ATG-VS, it will remove ligands that are not needed. Additionally, this module will optionally remove ligands that may be invalid for the scenario that is being processed (e.g. the presence of specific elements). This module will then submit each candidate ligand and docking scenario to the 'Docking' module.
- **Docking.** Processing of each individual docking (candidate ligand and receptor). Based on the docking program being used the module will construct the appropriate command line arguments and parse the output and send it along to the summary module. As here many concurrent processes are executing this is where most of the computational time is spent.
- **Summary.** Collects the individual docking scores and data and stores in memory until all dockings for the subjob are complete (single process)
- **Upload.** Once all of the dockings are complete, this module will upload the summary files to the results storage. (few processes)

All of these are distinct pipeline stages, so they can be overlapped. For example, if a subjob includes multiple collections, the downloading of the second collection can occur while the first collection is being docked.

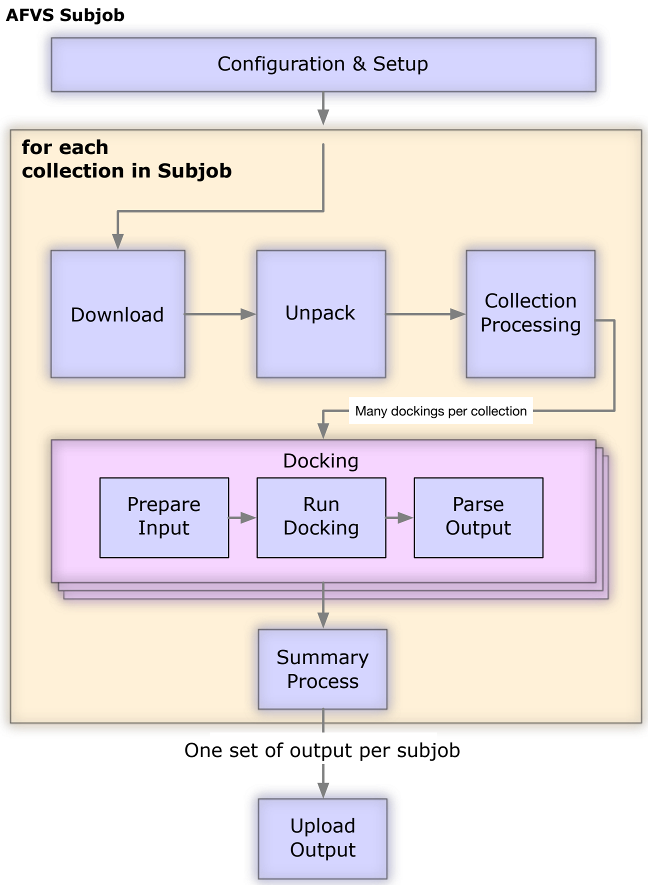

**Supplementary Fig. 4. Overview of the AFVS Subjob Procedure.** AdaptiveFlow uses a multi-process model for each subjob where work is processed via a set of submodules (implemented as processes) through Python queues. This model allows for parallelism within Python, while avoiding the Python Global Interpreter Lock (GIL) that can otherwise prevent efficient parallelism.

#### F AF Unity Workflow

AFU begins by processing the ligand: desalting and neutralizing the molecule, followed by enumerating stereoisomers and enantiomers. Subsequently, OpenBabel (145) is used to convert the molecule into a 3D file format (pdb, pdbt, mol2, sdf, etc.) based on the user's specified docking software. Once the ligands are prepared, the availability of any docking parameterization file (to be provided by the user) is checked, and the calculation is executed. As output, the program returns the docked ligand pose, along with the docking score (if available) of the protein-ligand complex. A step-by-step workflow for the program is described within Fig. 5.

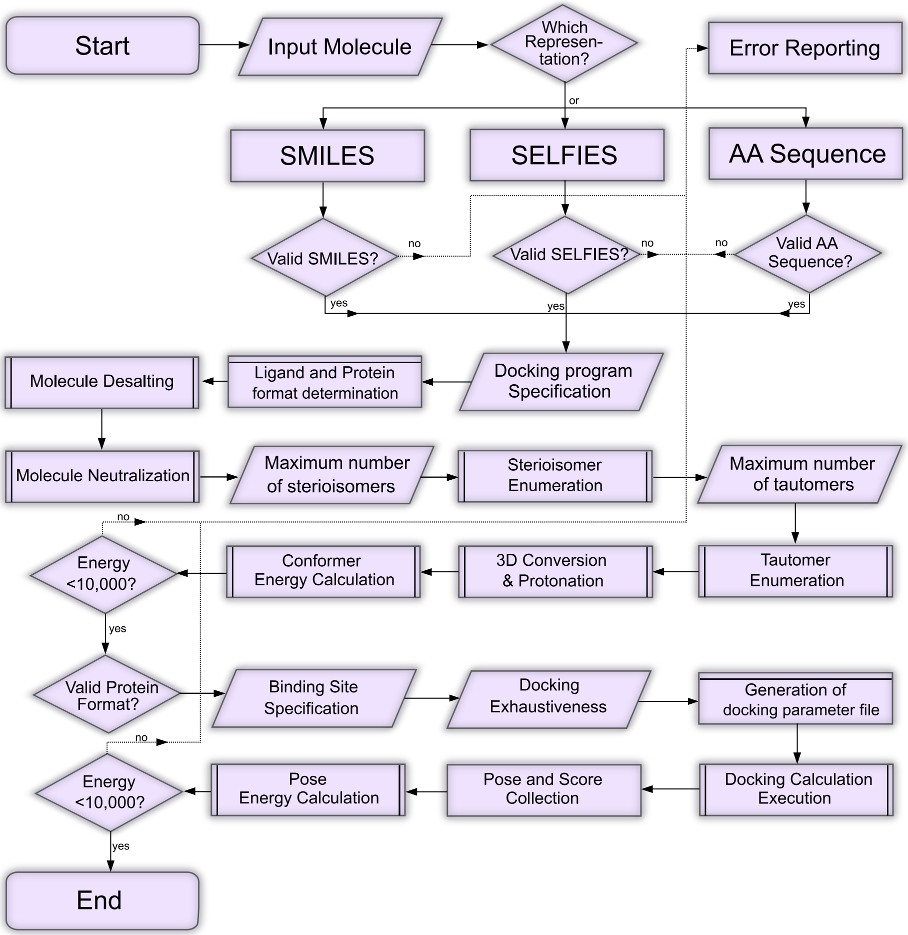

**Supplementary Fig. 5. UML Workflow Diagram of AdaptiveFlow Unity.** Initially, based on a user's specified choice of representation (SMILES, SELFIES, or amino-acid sequence), a molecule is checked for validity (by conversion back to SMILES and conversion to an RDKit mol object). Subsequently, based on the docking program specification, the supported ligand format is determined, allowing for the processing of the molecule by desalting, neutralizing, and enumerating tautomers and enantiomers. The user can control the maximum number of generated tautomers and enantiomers. The molecule is then converted into a 3D format (using OpenBabel) supported by the docking software choice at a specific protonation state (default pH 7.4). To check the successful conversion of the molecule, the energy of each 3D molecule is calculated using the obenergy program (molecules with an energy greater than 10,000 are unstable, resulting in their removal). After the successful processing of the ligand, based on the user's specified docking program, the availability of the required input file is checked. If there is no error, the docking calculation is executed, and the docking energy and the co-complexed ligand pose are returned.

The AFLP and AFVS modules of AdaptiveFlow require input libraries and generate output data/libraries. An overview of the file structures and associated relations is shown in Supplementary Fig. 6.

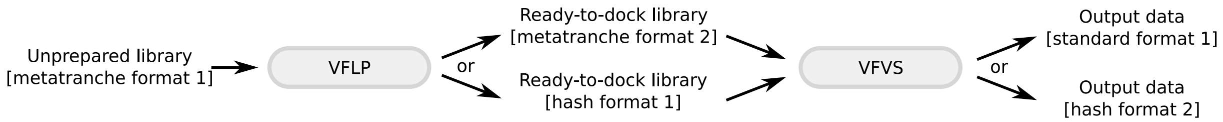

**Supplementary Fig. 6. Overview of Input and Output Library Formats within AdaptiveFlow.** AFLP requires that the processed input ligand library is stored in the *metatranche format 1*. AFLP can then store the output library in one of two types of formats, either the *metatranche format 2* or the *hash format 1*. These two formats for the ligand libraries can be used by AFVS, and the output data of AFVS can be stored in the *standard format 1* or the *hash format 2*.

The input-library format for AFLP is the *metatranche format 1*. In the root library folder are subfolders called metatranches. The metatranche folders contain subfolders that are called tranches. In the tranche folders are gzipped text files, called collections, where each collection file contains any number of ligands (typically up to several thousand ligands) in line notation. Supported line notations are SMILES, SELFIES, and amino acid sequences. The *metatranche format 1* library structure is illustrated in Supplementary Fig. 7. AFLP supports two library output formats, the *metatranche format 2* (Supplementary Fig. 8), as well as the *hash format 1* (Supplementary Fig. 9). These two formats can be used by AFVS for the input ligand libraries. *Hash format 1* includes two hash prefixes/directory levels, which allow a more equal distribution of the I/O to a moderate number of different folders. This can prevent I/O issues when running massively parallel workflows.

The REAL Space library (enumerated, version 2022q1-2) that we provide is stored in the *hash format 1*. Available subversions are the complete REAL Space library, as well as a sparse version that is used by the ATG-VS method. Available ligand file formats are PDB, PDBQT, SDF, MOL2, SMILES, and SELFIES. The library structure is illustrated in Supplementary Fig. 10.

The same two formats that can be generated by AFLP can also be used by AFVS, i.e. the *metatranche format 2* and the *hash format 1*. AFVS itself can produce two types of output formats in which the output data can be stored, the *standard format* (Supplementary Fig. 1) and the *hash format 2* (Supplementary Fig. 2).

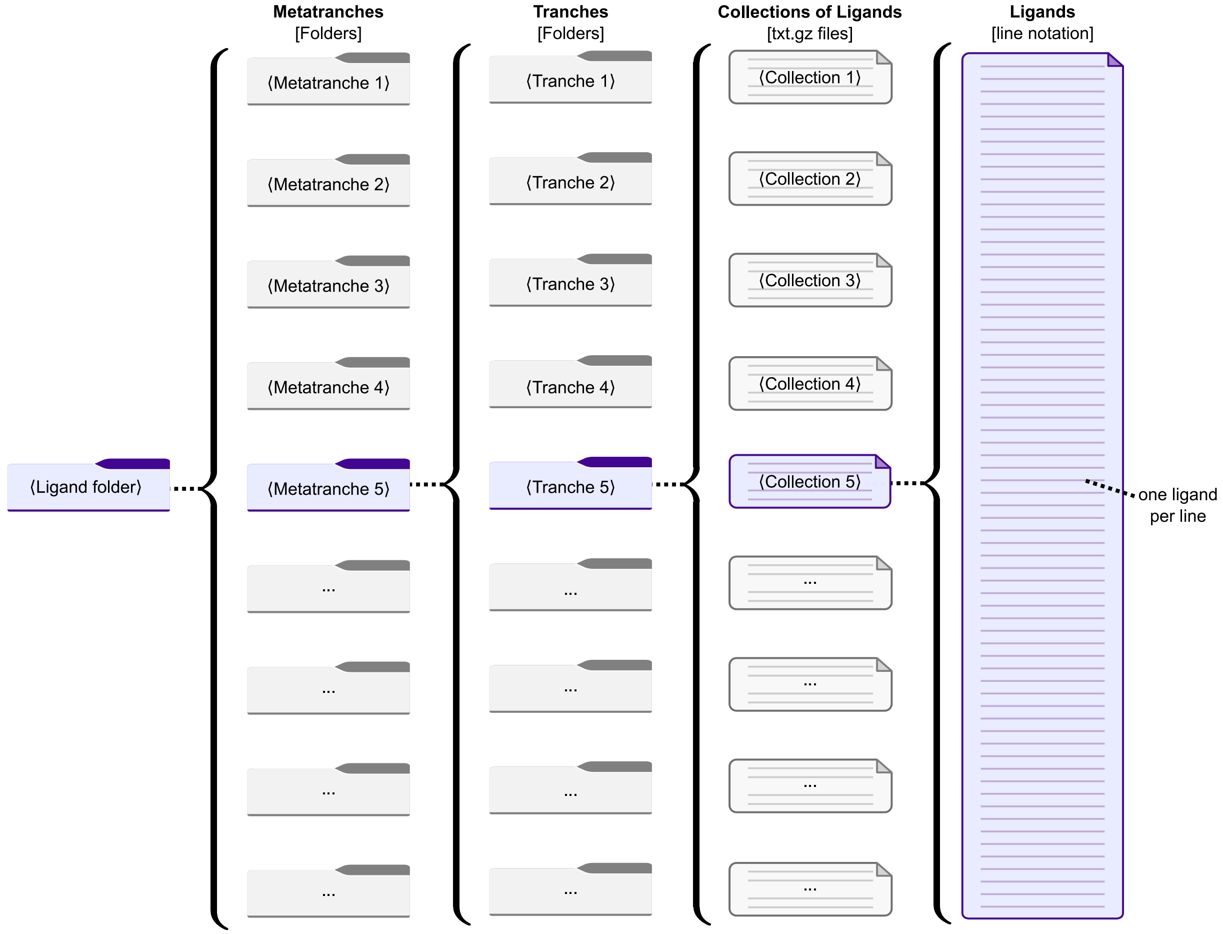

**Supplementary Fig. 7. AFLP Input Library Structure (Metatranche Format 1).** The *metatranche format 1* is used by AFLP as the input library format. In the library root folder are subfolders called metatranches, which contain subfolders called tranches. Each tranche folder contains one or more (ligand) collections, where each collection is a gzipped text file containing ligands in line notation (one ligand per line). Currently supported line notations are SMILES, SELFIES; and amino acid sequences.

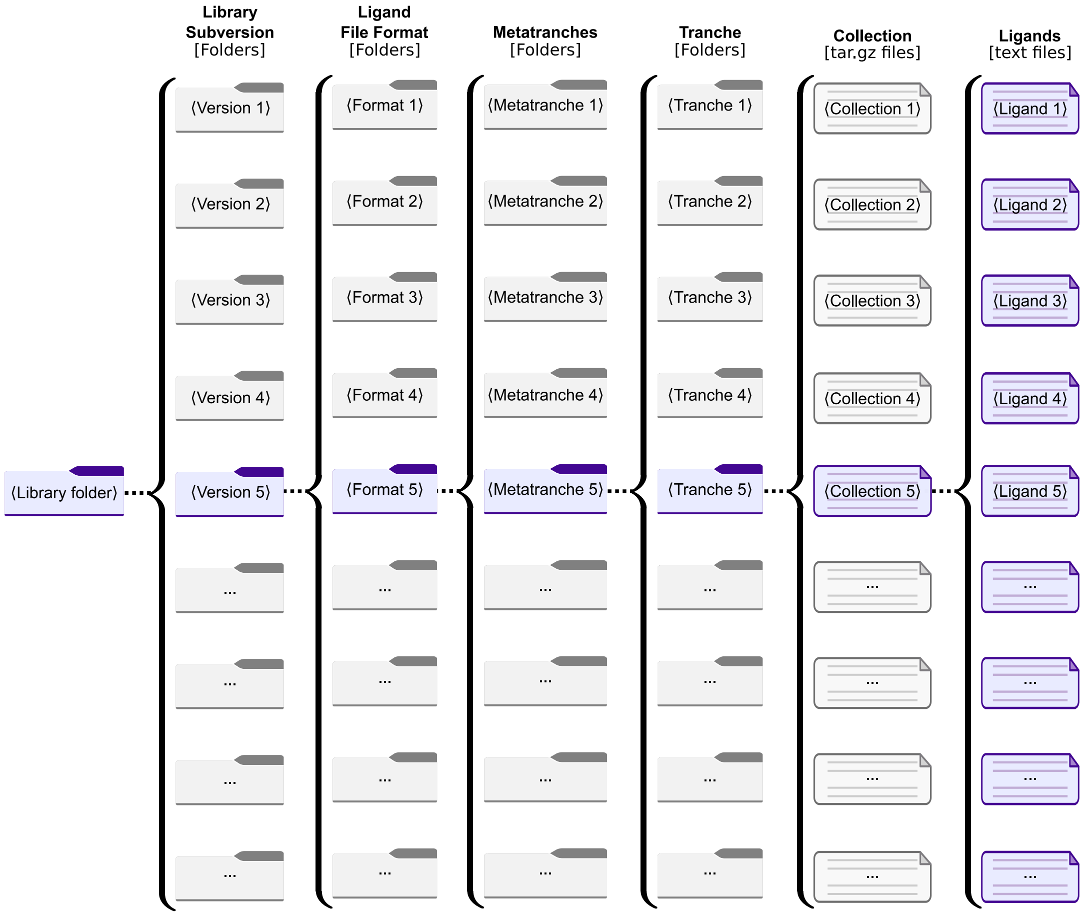

**Supplementary Fig. 8. AFLP Output Library Structure and AFVS Input Library Structure (Metatranche Format 2).** The *metatranche format 2* is one of the two file formats that AFLP can use for the output data ligand library and AFVS for the input ligand library. The library root folder contains subfolders for different library subversions (e.g. the full library, sparse versions, etc.). Each library subversion folder contains subfolders for the ligand file formats in which the ligands are stored. Each file format folder contains subfolders called metatranches, which themselves contain subfolders called tranches. Each tranche folder contains one or more (ligand) collections, where each collection is a gzipped tar archive containing the ligands in individual text files (one ligand per file).

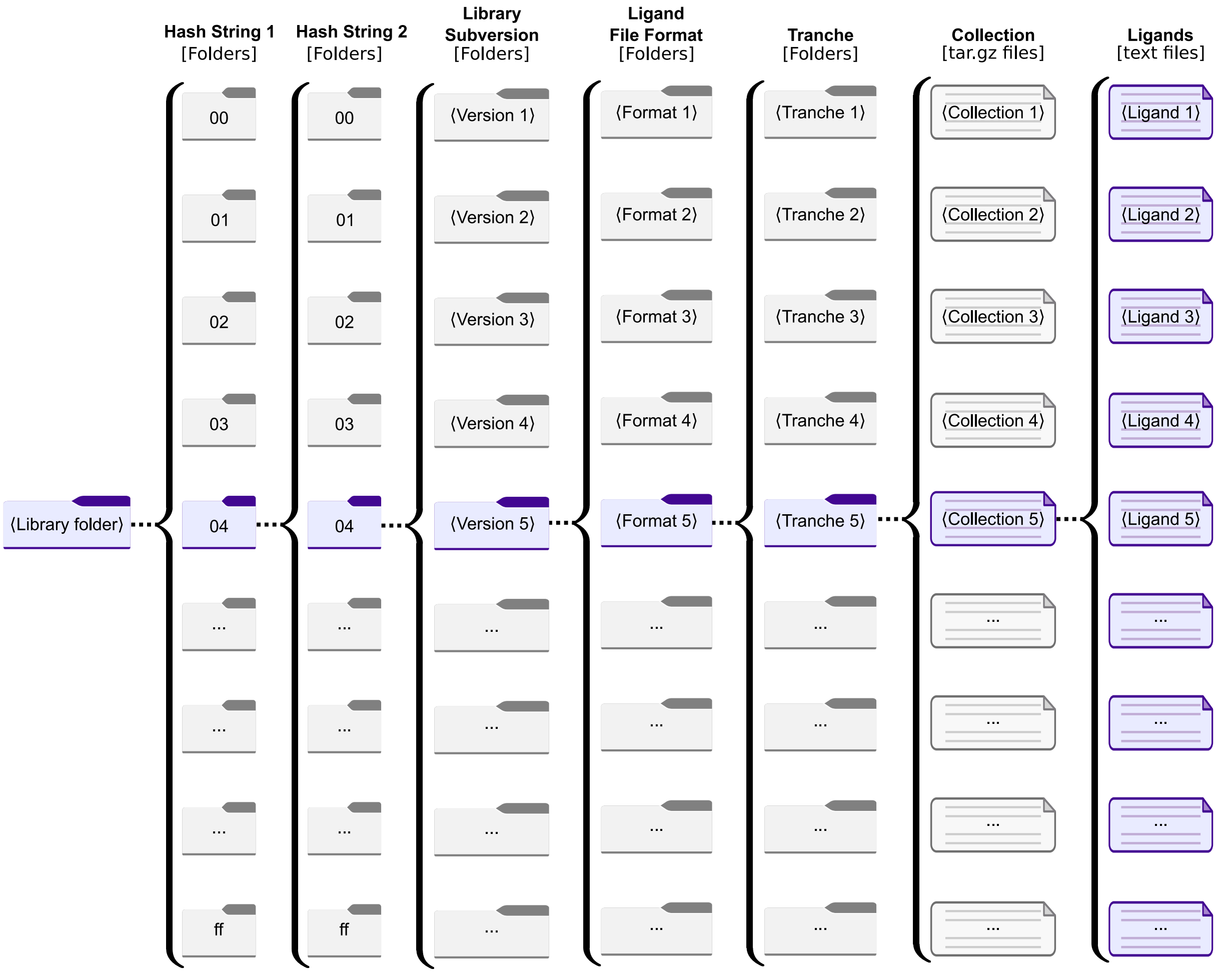

**Supplementary Fig. 9. AFLP Output Library Structure and AFVS Input Library Structure (Hash Format 1).** The hash format 1 is one of the two file formats that AFLP can use for the output data ligand library and AFVS for the input ligand library. The format is the same as the metatranche format 2, except that two hash subfolders are used directly within the library root folder. The hash folders each consist of two alphanumeric characters, and allow for a balanced distribution of the I/O load during highly parallel workflows, as well as allow to reduce the number of collections files per tranche folder. The hash prefixes are based on the collection and tranche IDs.

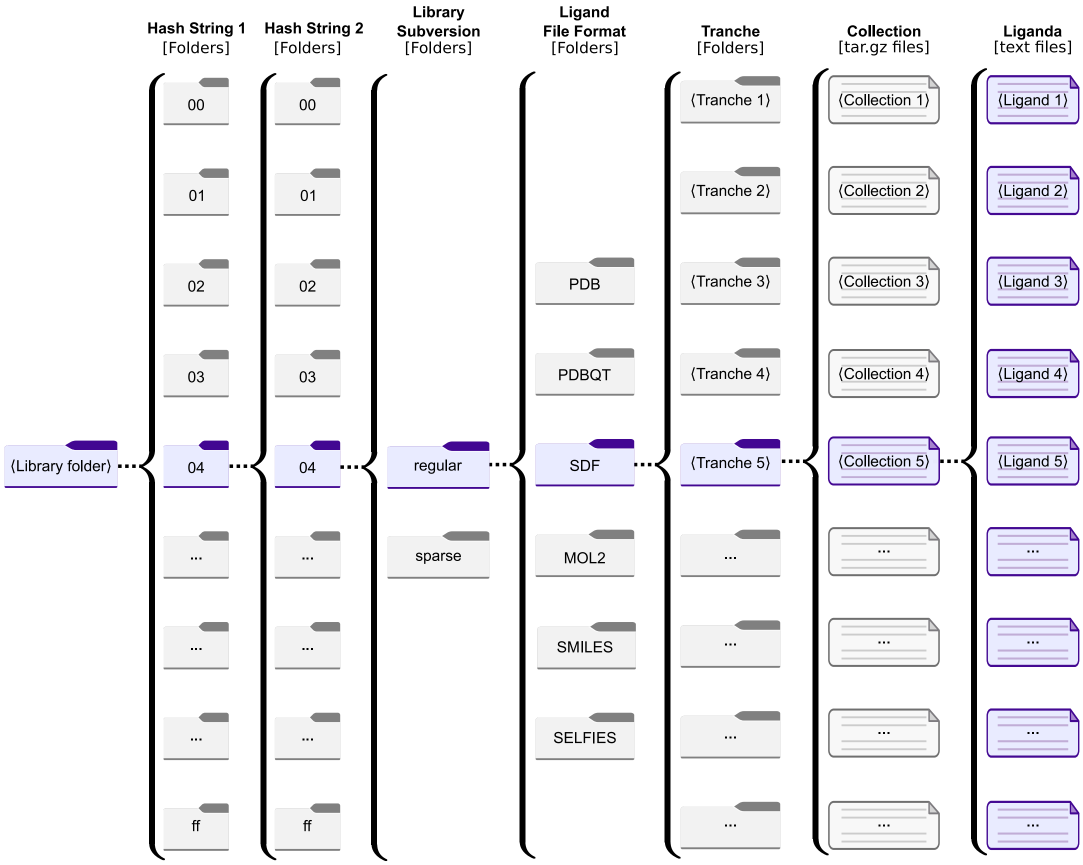

**Supplementary Fig. 10. Structure of the REAL Space in AdaptiveFlow's Hash Format 1.** The REAL Space that was prepared in this work was stored in hash format 1. The general hash format 1 was described in Supplementary Fig. 9. In the case of the REAL Space, there are two subversions of the library available, the regular (full) REAL Space, as well as a sparse version. Regarding the ligand file formats, six file formats are available (PDB, PDBQT, SDF, MOL2, SMILES, and SELFIES).

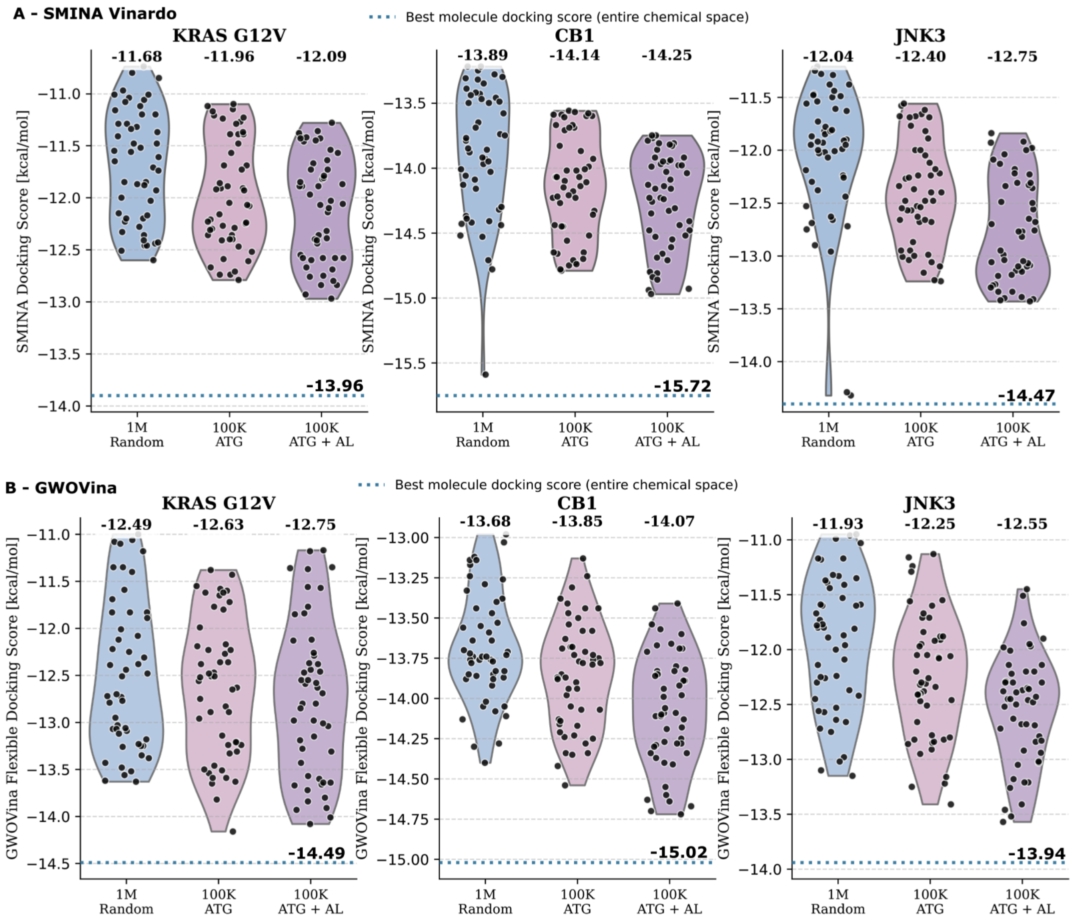

**Supplementary Fig. 11. Robustness of the Adaptive Target-Guided (ATG) strategy across different docking engines.** Performance comparison of the standard ATG and ATG with active learning step (ATG+AL) workflows applied to three distinct protein targets: KRAS G12V, CB1, and JNK3. Violin plots of the docking scores (the more negative, the better) for the top 50 hits obtained using (A) Smina (employing the Vinardo scoring function) and (B) GWOVina (incorporating flexible side-chain docking). The plots compare three approaches: (i) standard ultra-large virtual screening (ULVS) of 1 million random compounds (blue), (ii) adaptive target-guided (ATG) screening with 100,000 molecules (rose), and (iii) ATG screening augmented with active learning (purple). Black dots represent individual data points, and the numbers above the plots indicate the mean docking scores. The horizontal dashed lines indicate the best docking score obtained from a brute-force screen of the entire library, serving as a reference for the absolute best possible hit.

a

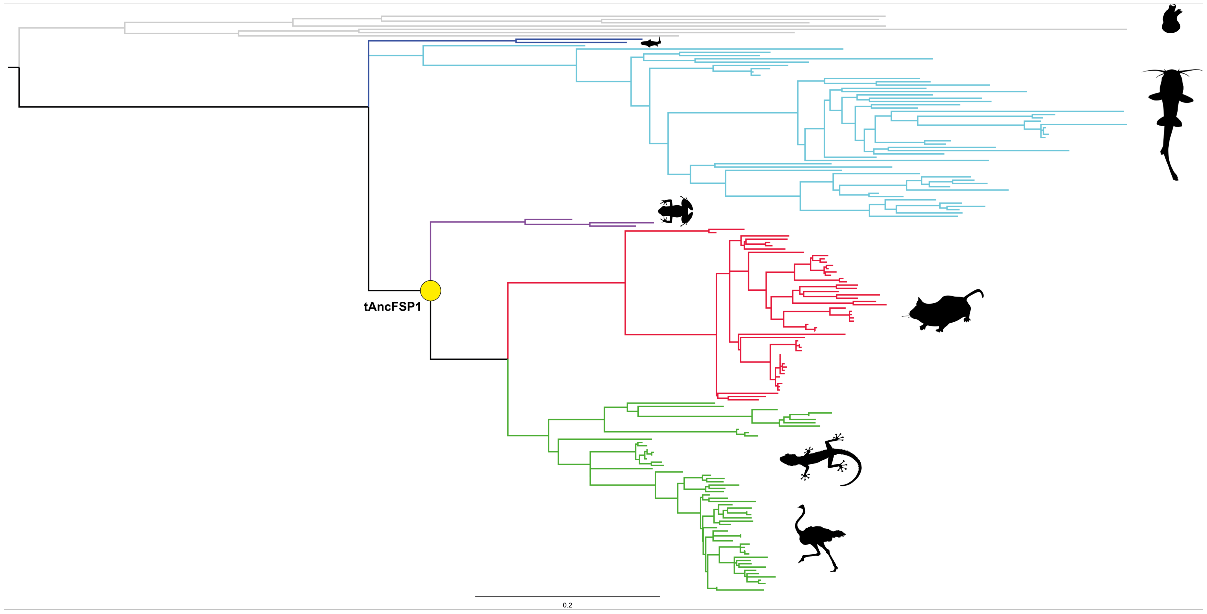

b

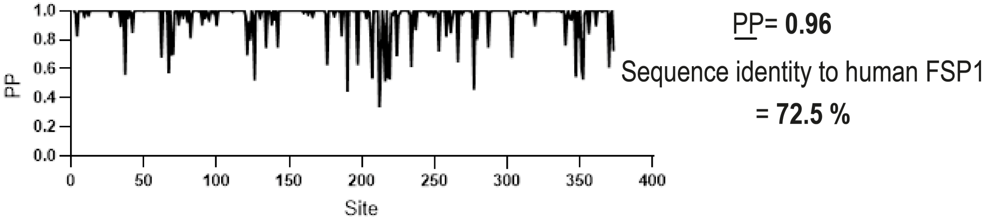

```
>MGSKLSVDDSVHVVIVGGGFGGIAAASQLKSWGVPFVLVDMKDAFHHNVAALRASVESGFAKKTFIPYAATFGDSFKQG
KVVGIDLEKQHVLLEDGEELHFSHLILATGSDGPFPGKFNQPVSMETAIQMYEDMVKEVQKAQRIVVVGGGSAGVEMAAE
IKTDYPDKEVTLIHSKIALADVELLPSVRQGVKEILLQKGVQLLLGQRVSNLQELTLNQVQENMKVKTDKGTEITADLVI
CCTGIKVNSSAYSSAFGDKLAENGALKVNEHLQVEGYDNVYAIGDCADVKEPKMAYHAGLHANVAVTNIINSLTNKPLKS
YKPGSLTMLLSMGRNDGVGQLNGYYVGRFVVRIAKSRDLFVGKSWKEMGQTMPH
```

**Supplementary Fig. 12. Ancestral Sequence Reconstruction of FSP1.** (a) Condensed maximum likelihood phylogenetic tree of Chordata FSP1 proteins. Taxonomic groups are color-coded: Echinodermata, Hemichordata, and Ascidiacea (grey); Chondrichthyes (blue); Actinopteri (cyan); Amphibia (violet); Mammalia (red); and Sauria (dark green). Silhouettes were obtained from `https://www.phylopic.org/`. The yellow circle highlights the reconstructed and experimentally characterized ancestral FSP1 corresponding to the tetrapodal ancestor. (b) Amino acid sequence of the reconstructed FSP1 used in this study, along with the posterior probability distribution per residue. Residues highlighted in red were excluded from the biochemical construct, as they correspond to the N-terminal region involved in myristoylation.

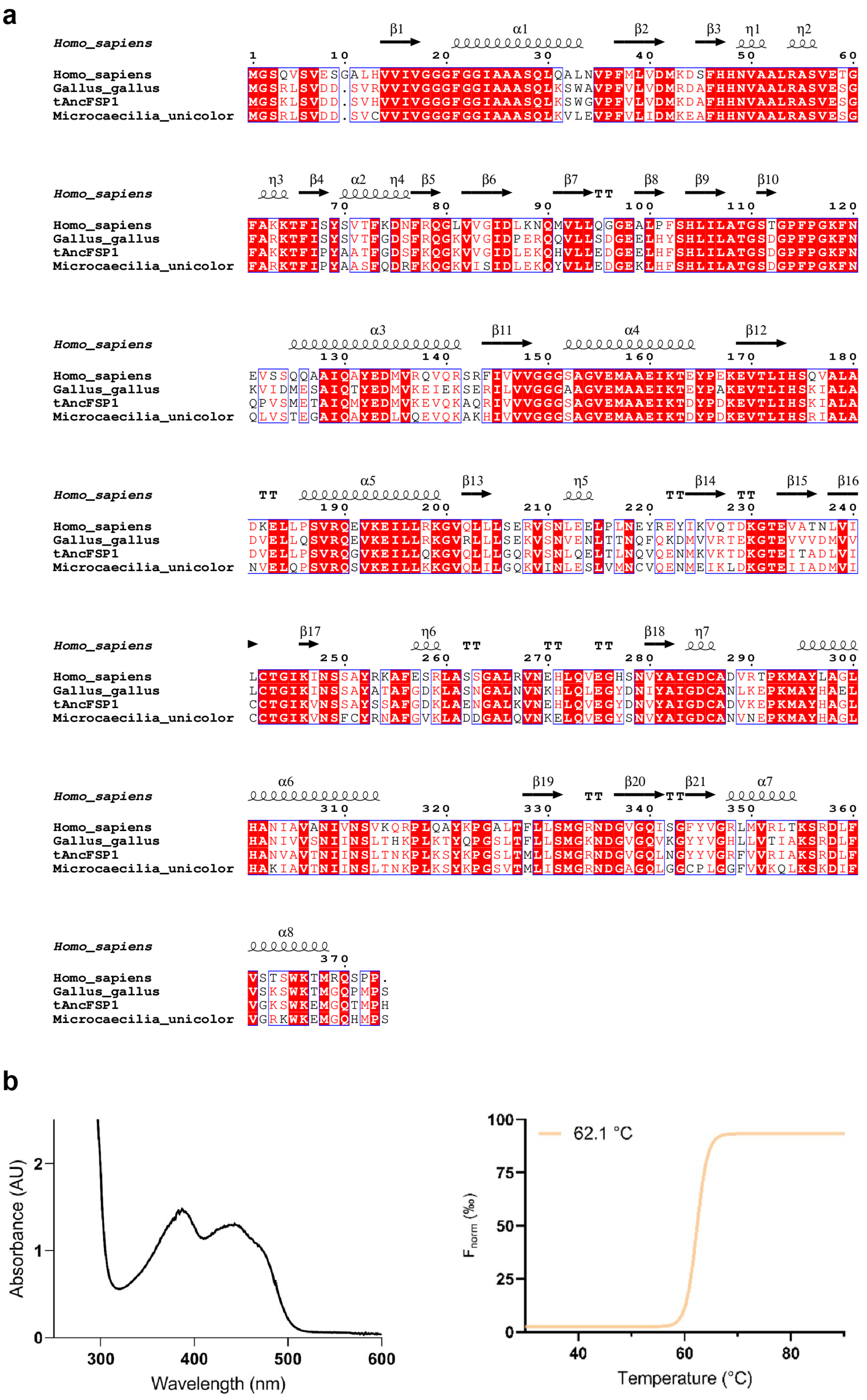

**Supplementary Fig. 13.** (a) Sequence alignment of human FSP1 with representative orthologs from amphibian (*Microcaecilia unicolor*), bird (*Gallus gallus*), and the reconstructed ancestral FSP1. (b) UV–visible absorbance spectrum and ThermoFAD unfolding profile of the ancestral FSP1.

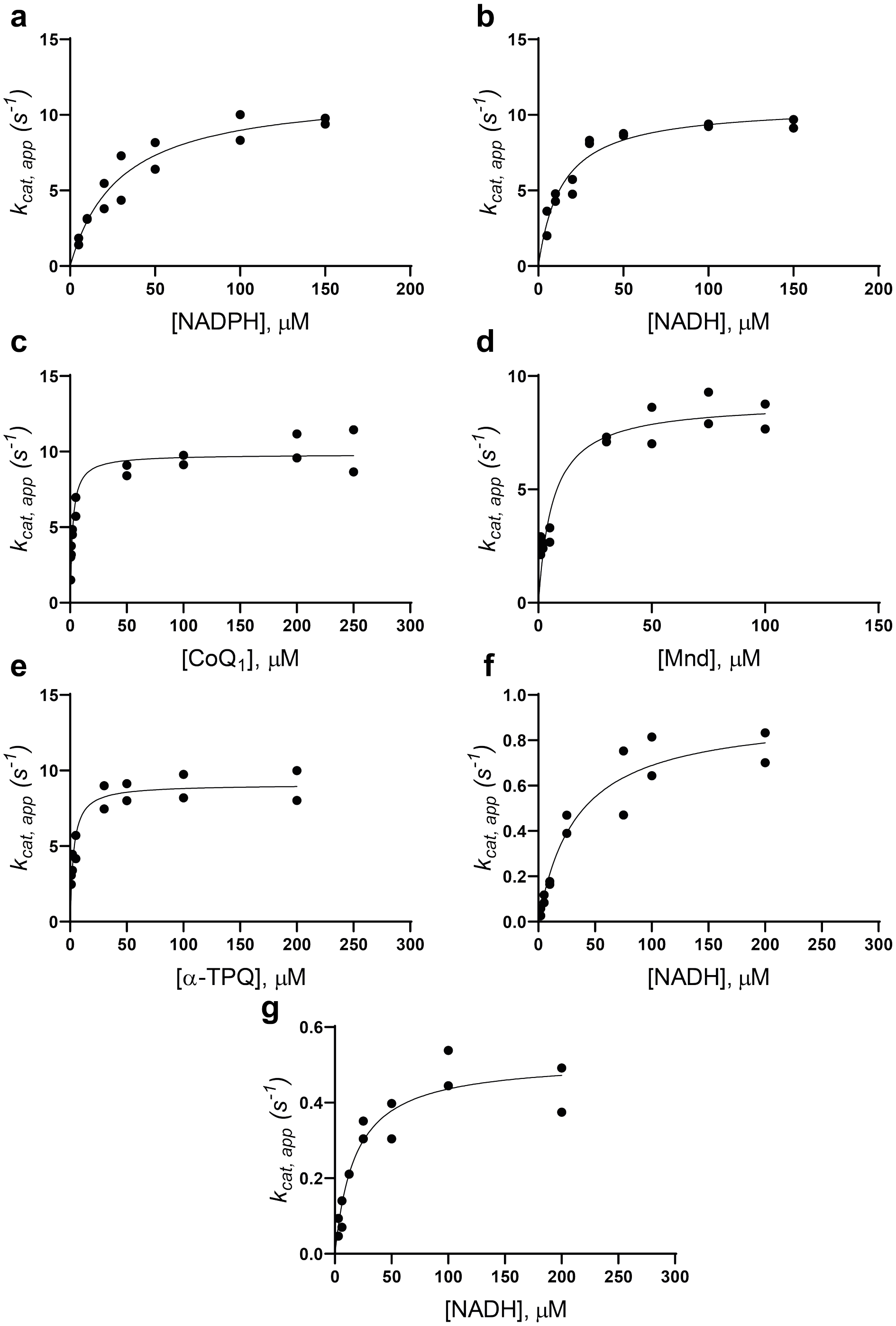

**Supplementary Fig. 14. Steady-state Kinetic Analysis of FSP1 Activity.** (a) NADPH titration at fixed coenzyme $Q_1$ (100 $\mu$M). (b) NADH titration at fixed coenzyme $Q_1$ (100 $\mu$M). (c) Coenzyme $Q_1$ titration at fixed NADH (100 $\mu$M). (d) Menadione titration at fixed NADH (100 $\mu$M). (e) α-tocopherol (quinonic hydrophilic head, α-TPQ) titration at fixed NADH (100 $\mu$M). (f) NADH titration monitored by absorbance at 340 nm. (g) NADH titration monitored by superoxide formation using the nitroblue tetrazolium (NBT) assay. Reactions in (a–e) were monitored via fluorescence-based NAD(P)H consumption, while (f–g) used absorbance-based and superoxide detection methods, respectively. Protein concentrations were 0.2 $\mu$M for panels (a–e) and 1 $\mu$M for panels (f–g). Each curve represents a fit to the Michaelis–Menten model; individual replicates ($n = 2$) are plotted as dots. MND = menadione; α-TPQ = α-tocopherol quinonic hydrophilic head.

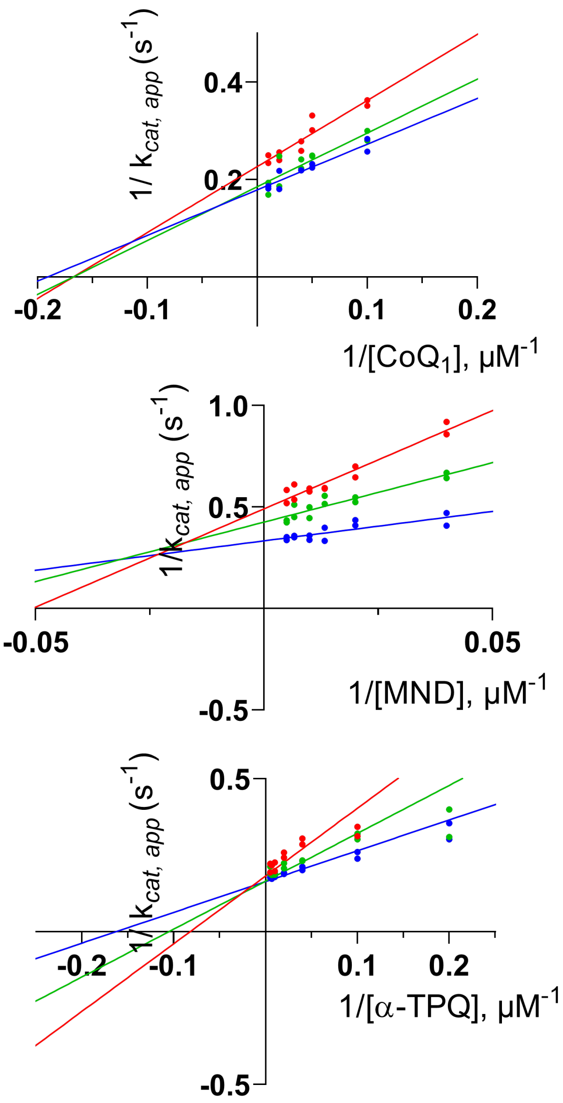

**Supplementary Fig. 15. Lineweaver–Burk Plots for Different Substrate Pairs.** (Top) NADH/coenzyme $Q_1$, (Middle) NADH/menadione (MND), and (Bottom) NADH/α-tocopherol (α-TPQ; quinonic hydrophilic head group). Experiments were performed with varying concentrations of the electron acceptor substrate at fixed NADH concentrations (25, 50, and 100 $\mu$M), shown in red, green, and blue lines, respectively. The convergence of lines to the left of the y-axis indicates a ternary complex mechanism of substrate binding.

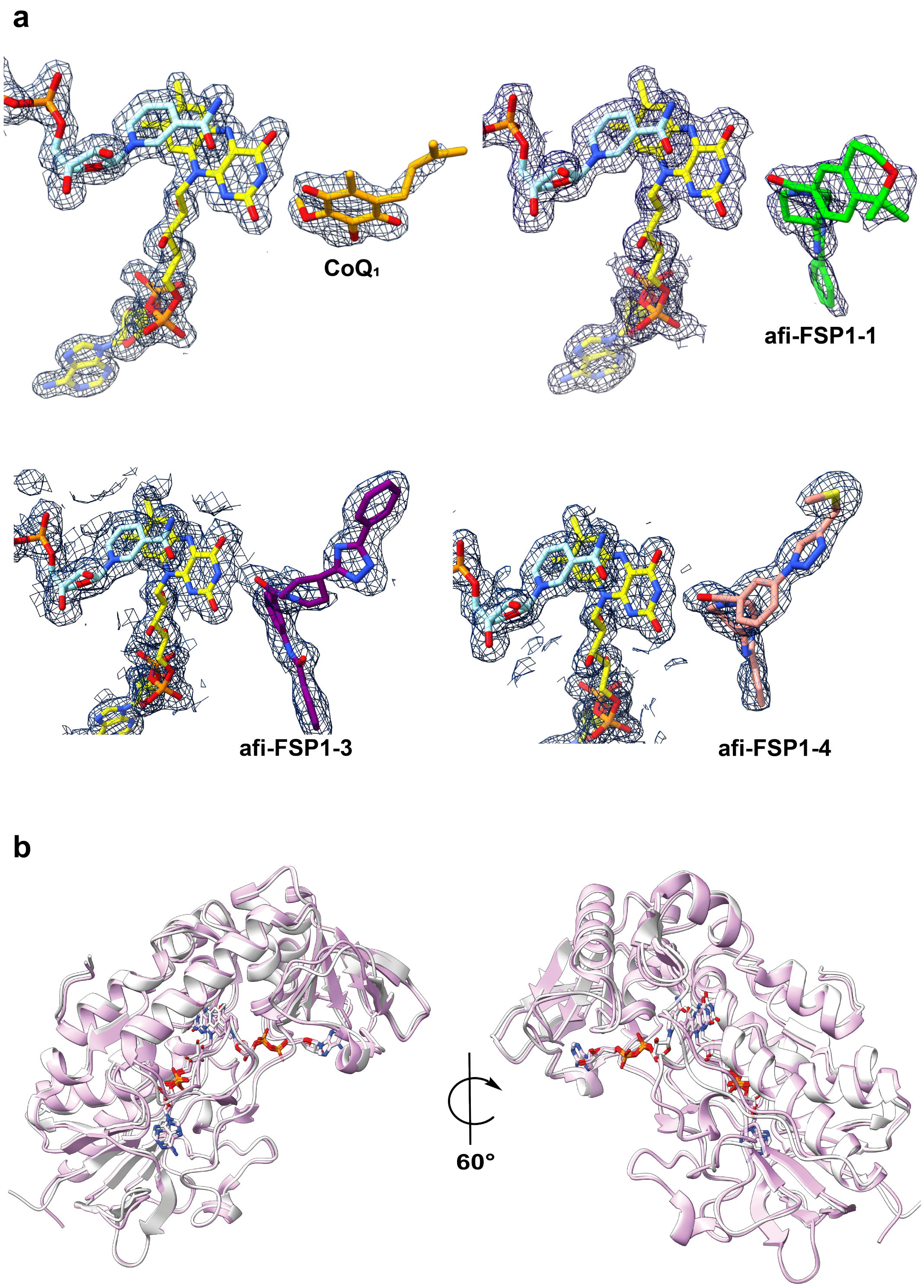

**Supplementary Fig. 16. Quality of Ligand Electron Densities and Structural Similarity of FSP1.** (a) Crystal structures of FSP1 in complex with $NAD^+$ and coenzyme $Q_1$ (orange), $NAD^+$ and compound afi-FSP1-1, $NAD^+$ and compound afi-FSP1-3, and $NAD^+$ and compound afi-FSP1-4 (inhibitors shown in green, purple and pink). FAD carbon atoms are colored yellow, and $NAD^+$ carbons in light cyan. Final $2F_o$–$F_c$ electron density maps are contoured at 1.3–1.4$\sigma$. (b) Structural superposition of ancestral FSP1 (white) and human FSP1 (pink; PDB 1WIK). The root-mean-square deviation (RMSD) is 1.1 Å across 373 C$\alpha$ atoms.

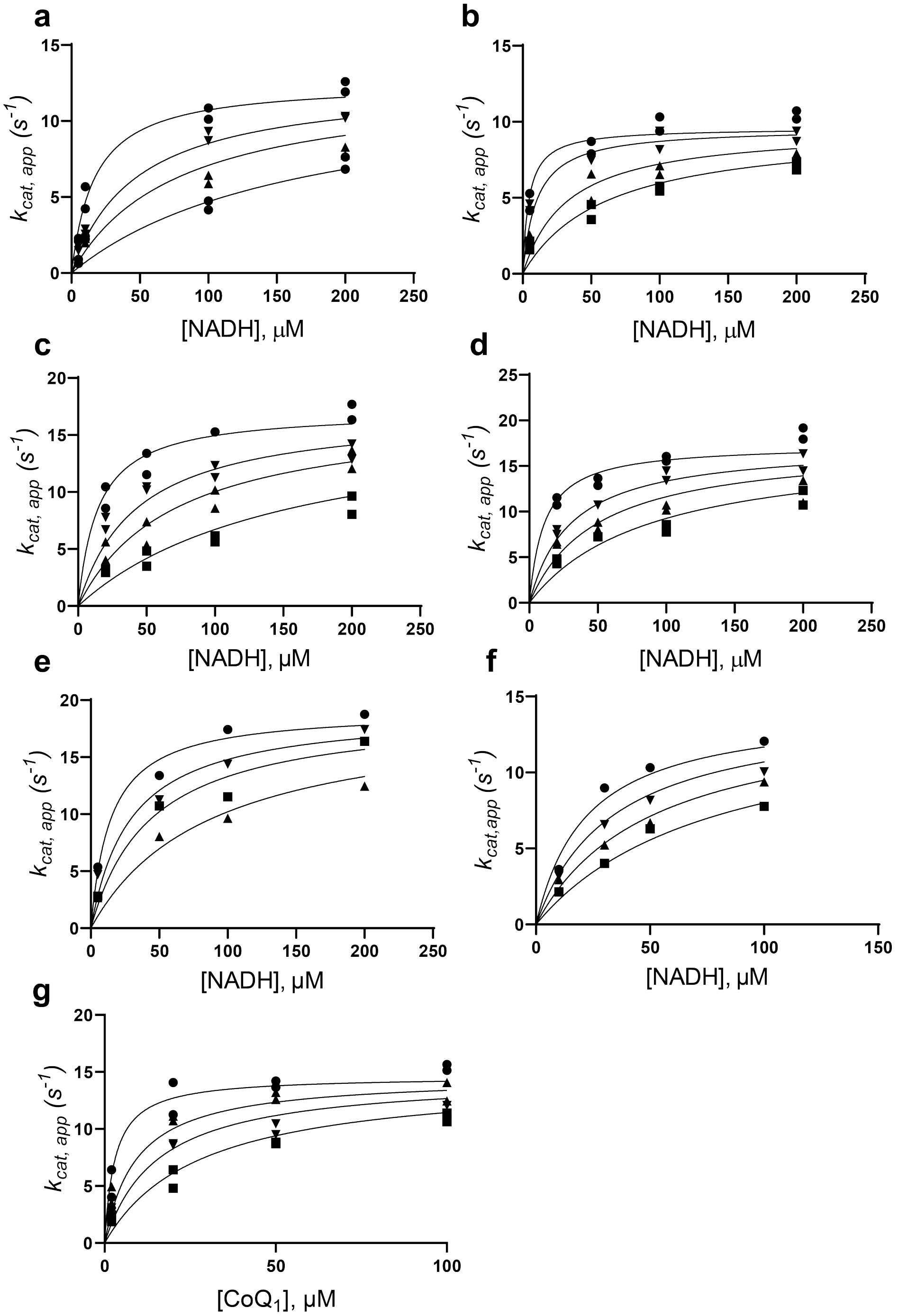

**Supplementary Fig. 17. $K_i$ measurements of FSP1 Inhibitors Under Competitive Inhibition Conditions.** (a) $K_i$ determination for compound afi-FSP1-1 by varying NADH concentrations in the presence of coenzyme $Q_1$ (100 $\mu$M). Data points (circles, downward arrows, upward arrows, squares) correspond to 0, 1.0, 2.0, and 5.0 $\mu$M inhibitor concentrations. (b) $K_i$ for compound afi-FSP1-2, tested at 0, 1.0, 5.0, and 10.0 $\mu$M ($n = 2$). (c) $K_i$ for compound afi-FSP1-3, tested at 0, 1.0, 2.0, and 5.0 $\mu$M. (d) $K_i$ for compound afi-FSP1-4, tested at 0, 0.5, 1.0, and 2.0 $\mu$M ($n = 2$). (e) $K_i$ for compound afi-FSP1-5, tested at 0, 1.0, 2.0, and 5.0 $\mu$M ($n = 1$). (f) $K_i$ for compound afi-FSP1-6, tested at 0, 2.0, 5.0, and 10.0 $\mu$M ($n = 1$). (g) $K_i$ for compound afi-FSP1-1 by varying coenzyme $Q_1$ at fixed NADH (100 $\mu$M), with inhibitor concentrations of 0, 0.5, 1.0, and 2.0 $\mu$M ($n = 2$). All panels show individual replicates plotted as dots, with non-linear regression fits.

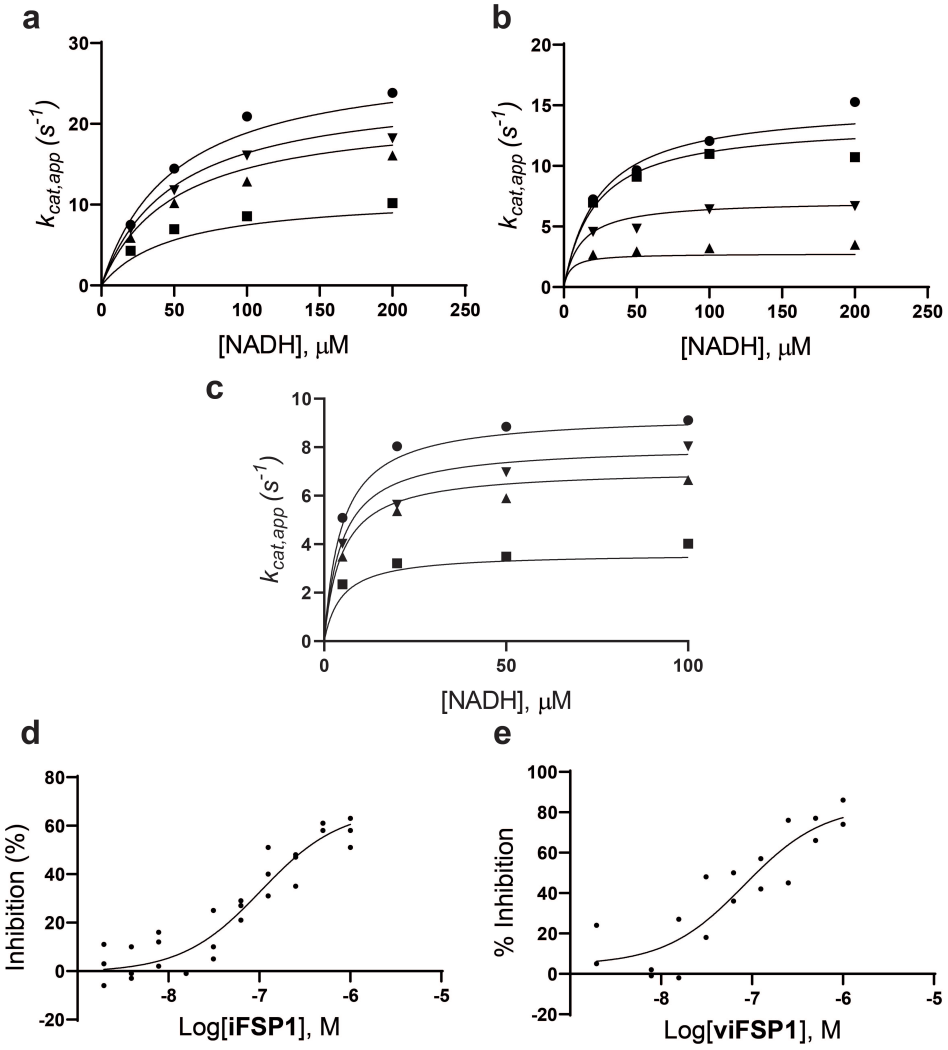

**Supplementary Fig. 18.** $K_i$ **and IC$_{50}$ Measurements for Previously Reported FSP1 Inhibitors.**(a) $K_i$ determination for iFSP1 by varying NADH concentrations in the presence of coenzyme $Q_1$ (100 $\mu$M). Data points (circles, downward arrows, upward arrows, and squares) correspond to 0, 0.5, 1.0, and 5.0 $\mu$M inhibitor concentrations. (b) $K_i$ determination for viFSP1 under the same conditions, using 0, 0.05, 0.5, and 2.0 $\mu$M inhibitor concentrations. (c) $K_i$ determination for FSEN1, tested at 0, 0.05, 0.1, and 0.5 $\mu$M inhibitor concentrations. All $K_i$ measurements were performed in singlicate ($n = 1$). (d–e) IC$_{50}$ values for iFSP1 and viFSP1 determined using the resazurin assay (100 $\mu$M resazurin, 200 $\mu$M NADH) with $n = 3$ and $n = 2$ replicates, respectively.

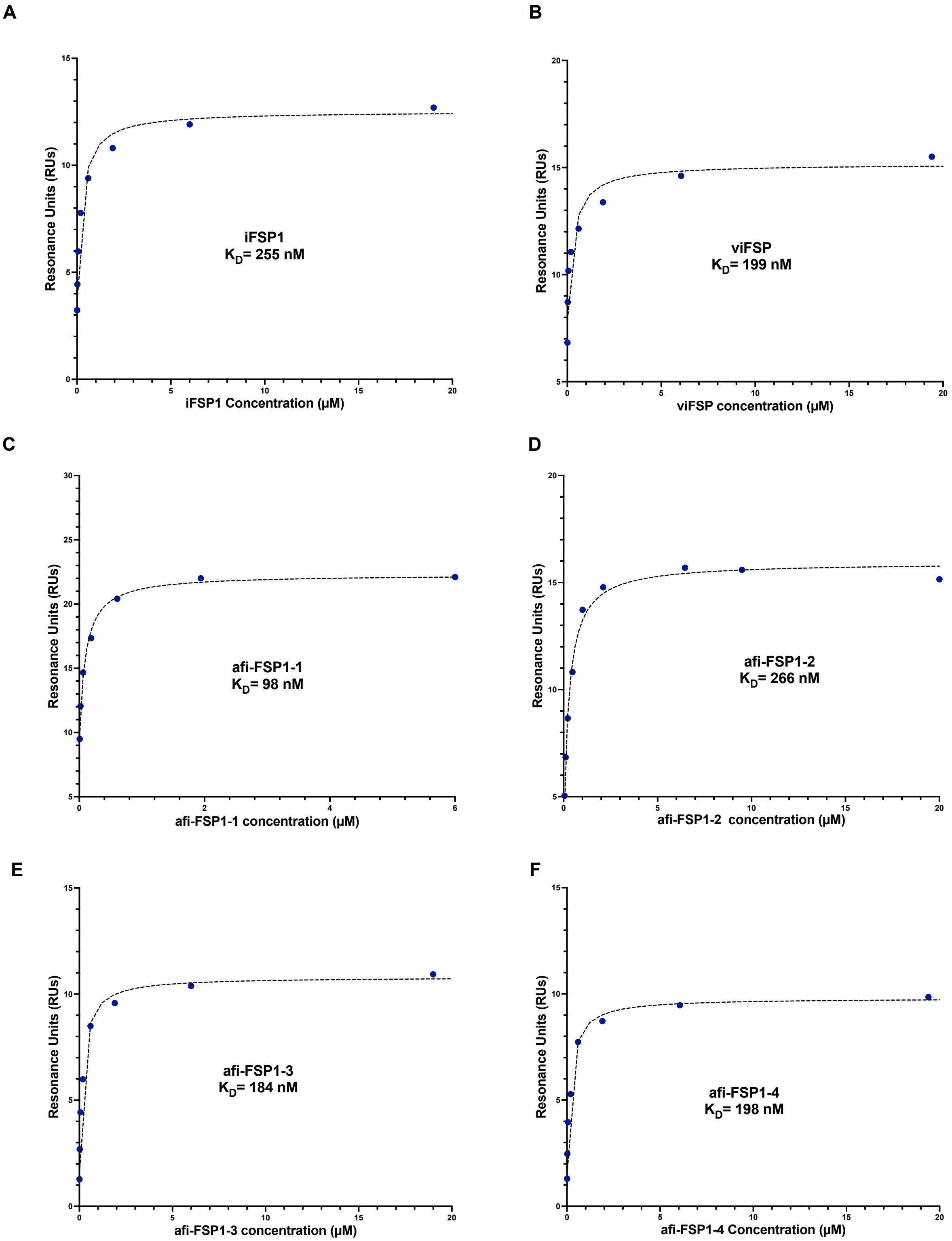

**Supplementary Fig. 19. Steady-State Binding Curves and Corresponding $K_D$ Values.** Steady-state binding curves and corresponding $K_D$ values based on surface plasmon resonance (SPR) for control compounds (A, B) and the top four inhibitors (C-F). Each curve represents binding responses measured across a range of compound concentrations, with data collected from three independent experiments. A single representative dataset is shown for each compound, illustrating the consistency and reproducibility of the observed binding interactions.

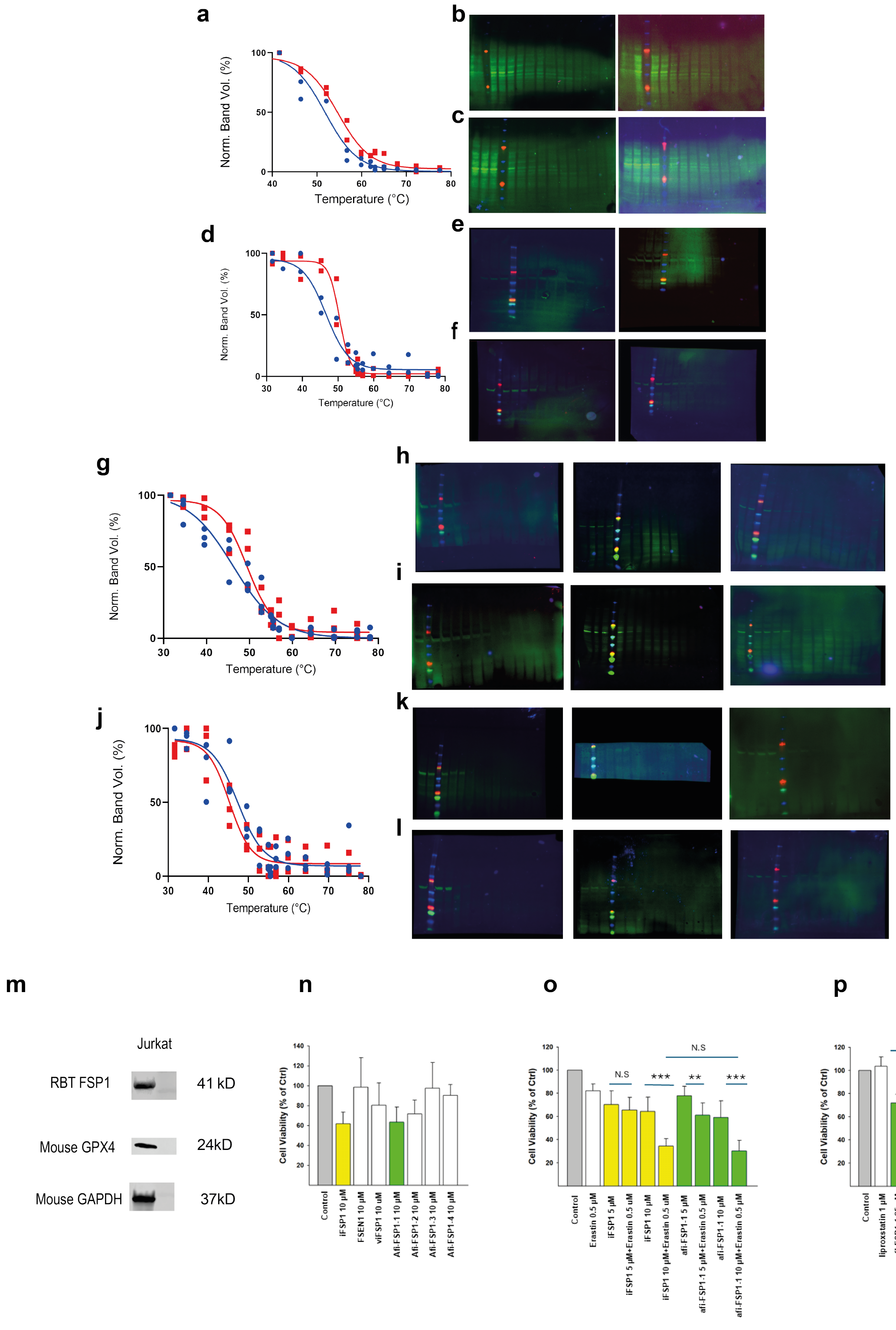

**Supplementary Fig. 20. Cellular Thermal Shift Assay (CETSA) and activity.** (a) 10 µM afi-FSP1-1 (red) engages endogenous human FSP1 (40.53 kDa) in HEK293 cells. The thermal shift is 2.8 ℃. Data are given as $n = 2$ independent experiments. DMSO (blue) was used as a control (b-c) Uncropped western blot gels representing protein quantification after exposure to increasing temperatures without (b) and with afi-FSP1-1 (c). (d) 1 µM afi-FSP1-2 (red) engages recombinant humanFSP1-twinSTREP-eGFP (71.5 kDa). The thermal shift is 3.1 ℃. Data are given as $n = 3$ independent experiments. (e-f) Uncropped western blot gels representing protein quantification after exposure to increasing temperatures without (e) and with afi-FSP1-2 (f). (g) 10 µM afi-FSP1-3 (red) engages human FSP1-twinSTREP-eGFP (71.5 kDa). The thermal shift is 3.2 ℃. Data are given as $n = 3$ independent experiments. (h-i) Uncropped western blot gels representing protein quantification after exposure to increasing temperatures without (h) and with afi-FSP1-3 (i). (j) No positive thermal shift was detected for 10 µM afi-FSP1-4 (red) possibly because of poor cell permeability or cell penetration. Data are given as $n = 3$ independent experiments. (k-l) Uncropped western blot gels representing protein quantification after exposure to increasing temperatures without (k) and with afi-FSP1-4 (l). For all the Western Blots, the molecular weight markers are in red and blue. (m) Jurkat cells express FSP1 and GPX4 (glutathione peroxidase 4) that is implicated in ferroptosis as it protects lipids from oxidation. (n) afi-FSP1 inhibitors (10 µM) are only minimally toxic as single agents. iFSP1, FSEN1, and viFSP1 are known FSP1 inhibitors. (o) Cotreatment of Jurkat cells with the combination of afi-FSP1-1 and erastin, a ferroptosis inducer, decreases cell viability to the same extent as the iFSP1-erastin cotreatment. (p) Liproxstatin, a ferroptosis inhibitor, fully rescues the effect of high 25 µM afi-FSP1-1.

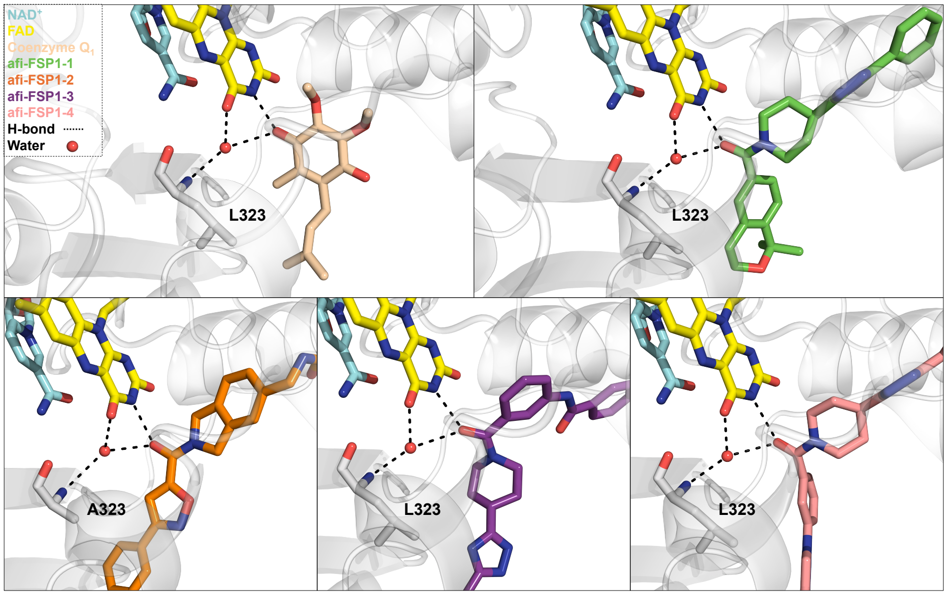

**Supplementary Fig. 21. Water-mediated hydrogen bonds between the flavin, the active-site ligands, and the protein.**

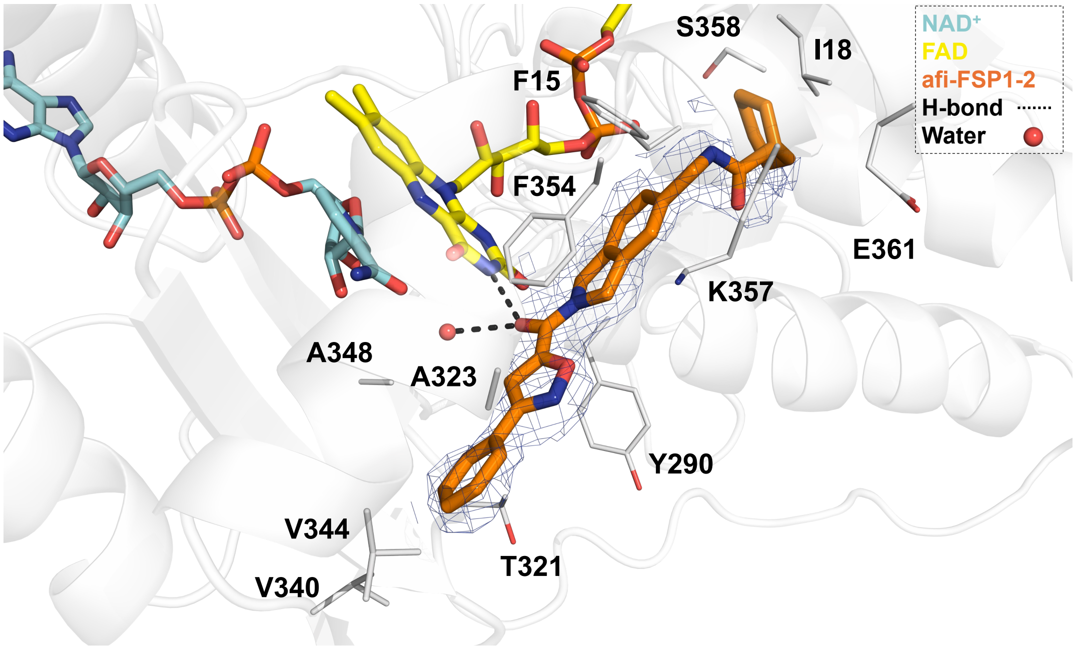

**Supplementary Fig. 22. Co-Crystal Structure of Inhibitor afi-FSP1-2 binding to FSP1 mutant L323A.** The polder omit map for the inhibitor is contoured at the 3.0 σ level. Close-up views of the binding site with afi-FSP1-2 (orange) bound in the region normally occupied by coenzyme $Q_1$, adjacent to FAD (yellow) and $NAD^+$ (cyan). The inhibitor displaces coenzyme $Q_1$ and interacts directly with the catalytic cavity.

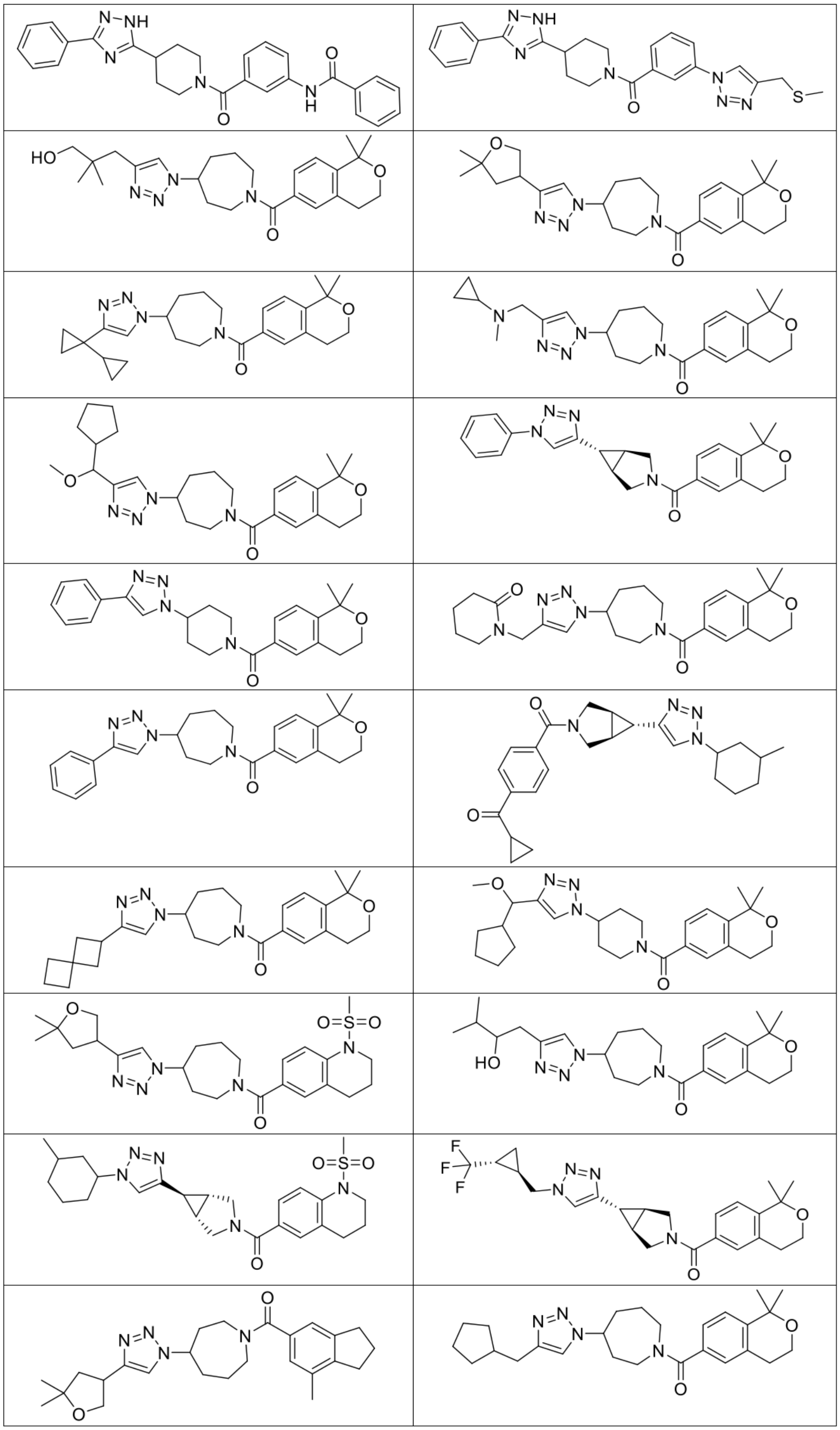

**Supplementary Fig. 23. Additional FSP1 Inhibitors.** Additional FSP1 inhibitors identified from the analog screening of afi-FSP1-1 and validated experimentally as a mixture if the analog has stereoisomers. Compounds showing $>$50% inhibition at 10 $\mu$M are highlighted in the top two rows and are listed in Supplementary Table 4.

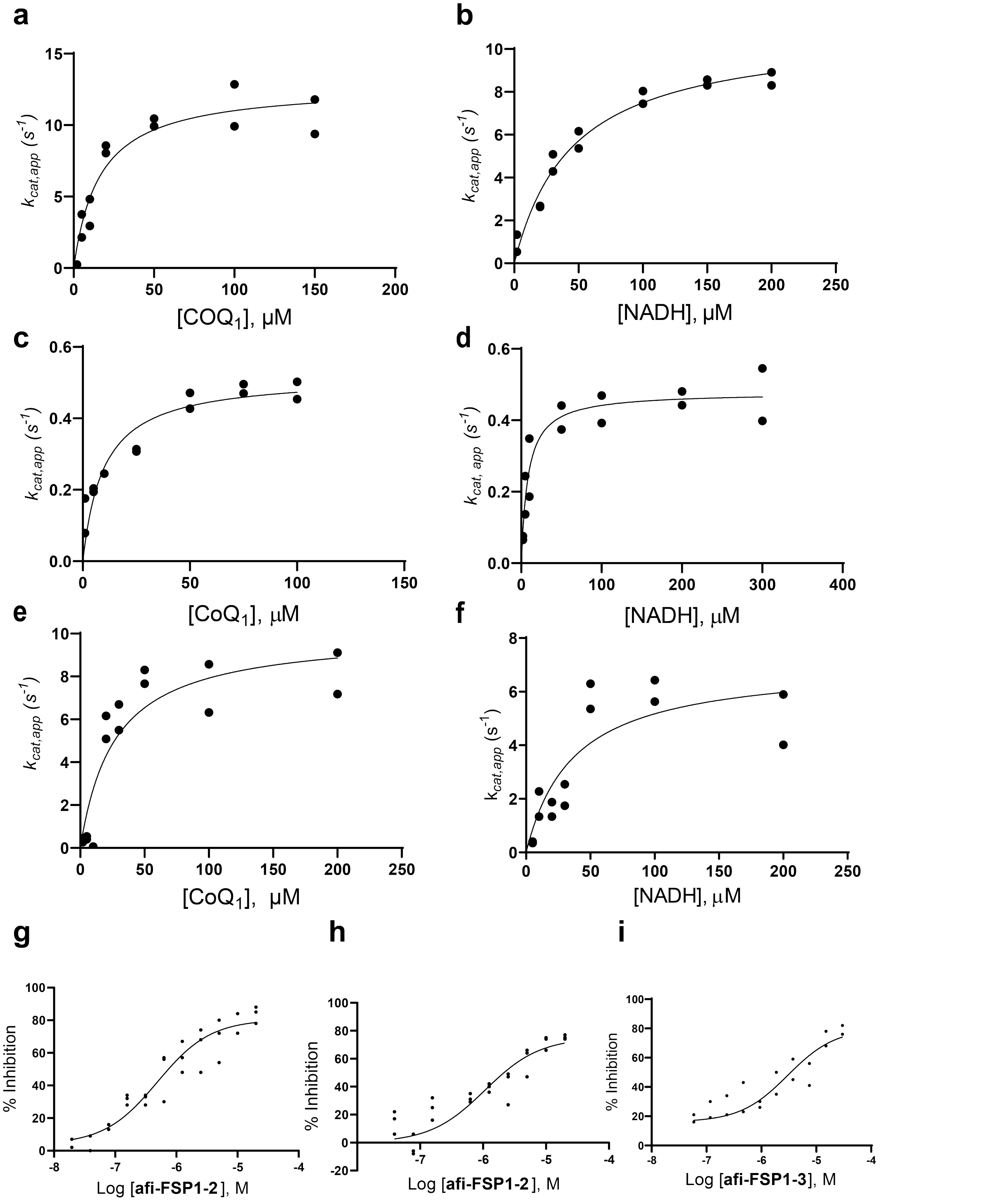

**Supplementary Fig. 24. Steady-State Kinetics and $IC_{50}$ Measurements for F15A and L323A FSP1 Mutants.** (a–b) Michaelis–Menten kinetics of L323A measured by varying coenzyme $Q_1$ at fixed NADH (100 $\mu$M) (a), or varying NADH at fixed coenzyme $Q_1$ (100 $\mu$M) (b). Protein concentration: 0.1 $\mu$M. (c-d) Michaelis–Menten kinetics of F15A under the same conditions. Protein concentration: 1 $\mu$M. Each data point represents an independent replicate ($n = 2$); dots show individual measurements, and curves represent non-linear regression fits. (e-f) Michaelis–Menten kinetics of F354A under the same conditions. Protein concentration: 0.2 $\mu$M. Each data point represents an independent replicate ($n = 2$); dots show individual measurements, and curves represent non-linear regression fits. None of the compounds developed in this study inhibits this mutant. (g) $IC_{50}$ determination for compound afi-FSP1-2 against L323A (0.1 $\mu$M) using the resazurin assay (50 $\mu$M resazurin, 200 $\mu$M NADH, $n = 3$). (h-i) $IC_{50}$ determination for compounds afi-FSP1-2 ($n = 3$) and afi-FSP1-3 ($n = 2$) against F15A (0.5 $\mu$M).

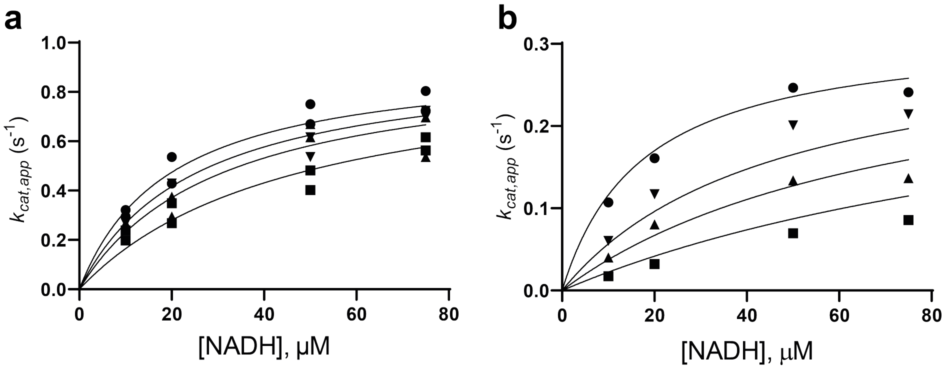

**Supplementary Fig. 25.** $K_i$ **Determination from NADH Oxidase Activity Assays under Competitive Inhibition Conditions.** (a) $K_i$ for compound afi-FSP1-1, measured by varying NADH concentrations in the absence of coenzyme $Q_1$. Data points (circles, downward arrows, upward arrows, and squares) represent inhibitor concentrations of 0, 1.0, 2.0, and 5.0 $\mu$M, respectively ($n = 2$). (b) $K_i$ for compound afi-FSP1-2, measured under the same conditions. Data points represent inhibitor concentrations of 0, 5.0, 10.0, and 20.0 $\mu$M, respectively ($n = 1$).

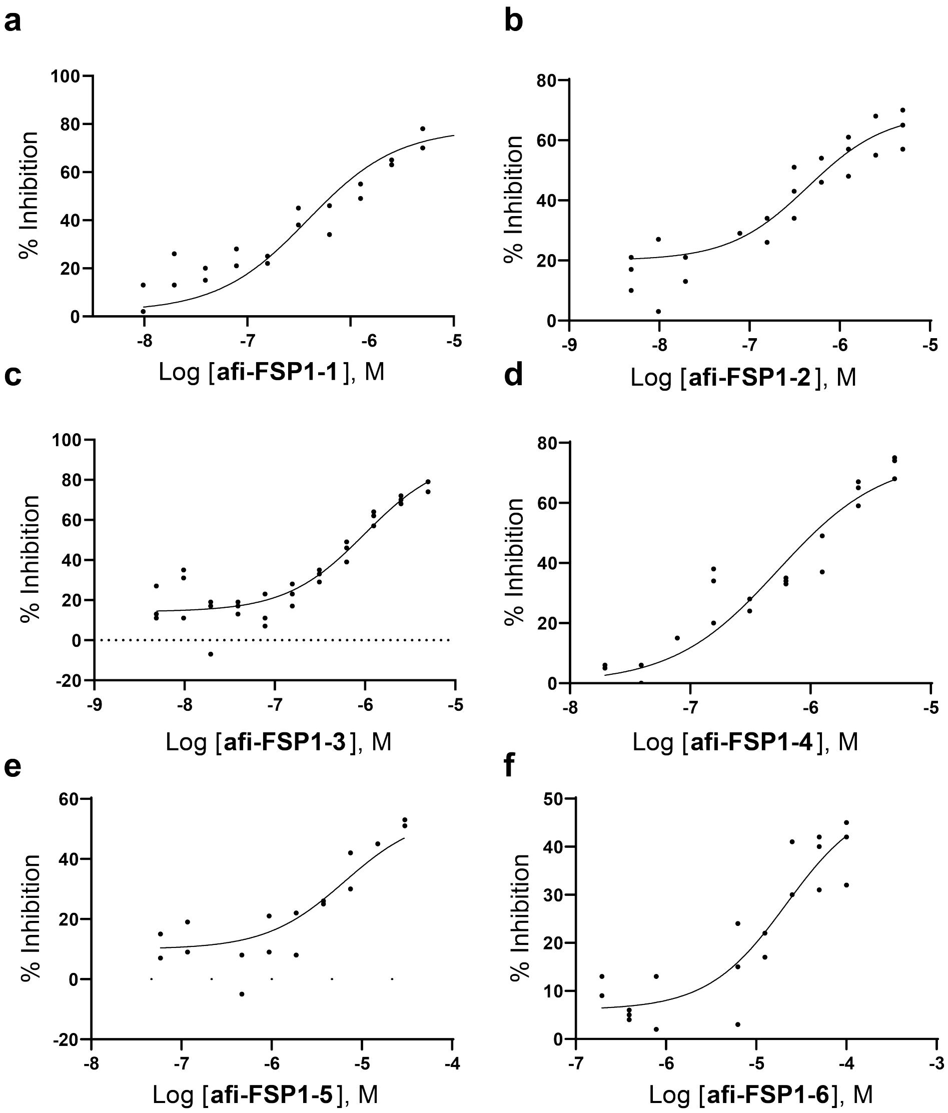

**Supplementary Fig. 26. $IC_{50}$ Values ($\mu$M) of the Most Potent FSP1 Inhibitors.** Panels (a–f) show dose–response curves obtained using the resazurin assay (100 $\mu$M resazurin, 200 $\mu$M NADH). Each curve represents mean values from independent experiments ($n = 2$–3).

| Condition | $k_{cat}$ ($s^{-1}$) | $K_M$ ($\mu$M) | $\frac{k_{cat}}{K_M}$ ($s^{-1}\mu M^{-1}$) |
|---|---|---|---|
| **Oxidoreductase activity**[a] | | | |
| Varying NADPH, fixed coenzyme $Q_1$ | $11.60 \pm 0.88$ | $29.27 \pm 6.26$ | 0.40 |
| Varying NADH, fixed coenzyme $Q_1$ | $10.65 \pm 0.50$ | $13.89 \pm 2.39$ | 0.76 |
| Varying coenzyme $Q_1$, fixed NADH | $9.81 \pm 0.33$ | $2.15 \pm 0.36$ | 4.56 |
| Varying menadione, fixed NADH | $8.86 \pm 0.45$ | $6.33 \pm 0.74$ | 1.40 |
| Varying $\alpha$-tocopherol[b], fixed NADH | $9.07 \pm 0.35$ | $3.00 \pm 0.60$ | 3.02 |
| **NADH oxidase activity at atmospheric $O_2$ concentration** | | | |
| Varying NADH[c] | $0.92 \pm 0.08$ | $33.96 \pm 9.53$ | 0.03 |
| Varying NADH[d] | $0.51 \pm 0.04$ | $18.04 \pm 5.60$ | 0.03 |

**Supplementary Table 3. Steady-state Kinetic Parameters for FSP1 with Different Electron Donors and Acceptors.** $k_{cat}$ and $K_M$ values were determined from Michaelis–Menten fits. [a]Fixed concentrations were 100 $\mu$M. Data shown in Supplementary Fig. 14. [b]Quinone hydrophilic head group. [c]Measured via NADH consumption assay. [d]Measured using nitroblue tetrazolium assay (superoxide detection).

| Parameter | FAD | FAD + $NAD^+$ | FAD + $NAD^+$ + $CoQ_1$ | FAD + $NAD^+$ + afi-FSP1-1 | FAD + $NAD^+$ + afi-FSP1-2 (L323A) | FAD + $NAD^+$ + afi-FSP1-3 | FAD + $NAD^+$ + afi-FSP1-4 |
|---|---|---|---|---|---|---|---|
| Space group | $P2_1$ | $P2_1$ | $P2_1$ | $P2_1$ | $P2_1$ | $P2_1$ | $P2_1$ |
| Unit cell a/b/c (Å) | 43.09<br>81.54<br>57.69 | 43.95<br>80.54<br>60.92 | 43.51<br>79.15<br>112.07 | 43.57<br>79.68<br>113.06 | 44.45<br>78.06<br>115.08 | 43.85<br>79.33<br>113.37 | 43.66<br>79.51<br>112.78 |
| Resolution (Å) | 2.10 | 1.45 | 1.70 | 1.55 | 2.40 | 1.94 | 1.74 |
| PDB code | 9IFZ | 9IFT | 9IFY | 9IFU | 9QRT | 9IG9 | 9IFW |
| $R_{sym}$ [a,b] (%) | 19.4 (98.3) | 8.9 (171.0) | 4.4 (53.6) | 10.6 (39.3) | 12.0 (71.0) | 11.9 (84.6) | 8.0 (74.8) |
| $CC_{1/2}$ (%) | 99.2 (70.2) | 99.9 (45.5) | 99.8 (70.9) | 97.9 (76.5) | 97.3 (36.1) | 95.5 (32.7) | 99.4 (27.7) |
| Completeness[b] (%) | 99.9 (99.8) | 99.0 (99.5) | 95.8 (96.8) | 83.7 (50.6)[e] | 96.1 (98.1) | 93.5 (81.6) | 97.1 (96.5) |
| Unique reflections | 23,156 | 72,942 | 79,096 | 46,979 | 29,339 | 53,297 | 75,055 |
| Redundancy | 6.9 (7.1) | 6.9 (6.7) | 2.7 (2.6) | 2.5 (2.6) | 2.7 (2.6) | 2.0 (2.0) | 2.4 (2.4) |
| $I/\sigma$ [b] | 7.2 (2.1) | 10 (1.0) | 8.9 (1.1) | 5.1 (2.2) | 7.4 (1.6) | 4.7 (0.8) | 5.0 (0.6) |
| Non-H atoms (protein / FAD) | 2770 / 53 | 2804 / 53 | 5594 / 2×53 | 5612 / 2×53 | 5534 / 2×53 | 5548 / 2×53 | 5573 / 2×53 |
| $NAD^+$ | – | 44 | 2×44 | 2×44 | 2×44 | 2×44 | 2×44 |
| Coenzyme $Q_1$ | – | – | 2×17 | – | – | – | – |
| Inhibitor | – | – | – | 2×30 | 33 | 2×34 | 2×32 |
| Water molecules | 118 | 294 | 364 | 556 | 161 | 182 | 273 |
| B-factor (protein / ligands) ($Å^2$) | 27.6 / 28.0 | 22.1 / 18.3 | 30.94 / 31.42 | 21.38 / 24.51 | 29.97 / 29.65 | 29.15 / 30.17 | 25.9 / 29.16 |
| $R_{cryst}$ [b,c] (%) | 19.76 (24.0) | 17.5 (32.0) | 18.81 (34.0) | 17.9 (18.7) | 21.6 (22.9) | 16.3 (32.8) | 20.5 (36.0) |
| $R_{free}$ [b,c] (%) | 22.74 (31.2) | 20.3 (33.0) | 22.58 (37.9) | 24.25 (22.0) | 28.0 (24.0) | 15.9 (26.6) | 20.5 (21.57) |
| RMS bond length (Å) | 0.064 | 0.010 | 0.008 | 0.006 | 0.0070 | 0.0069 | 0.0076 |
| RMS bond angles (°) | 1.37 | 1.63 | 1.47 | 1.34 | 1.47 | 1.46 | 1.53 |

**Supplementary Table 4. Data Collection and Refinement Statistics for FSP1 Crystal Structures.** $R_{\text{sym}} = \sum(|I_i - \langle I \rangle|)/\sum I_i$, where $I_i$ is the intensity of the $i^{\text{th}}$ observation and $\langle I \rangle$ the mean intensity of the reflection. Values in parentheses correspond to the highest resolution shell. $R_{\text{cryst}} = \sum(|F_{\text{obs}} - F_{\text{calc}}|)/\sum|F_{\text{obs}}|$. $R_{\text{free}}$ was calculated using a separate test set. Anisotropic datasets were processed with *STARANISO* (http://staraniso.globalphasing.org/cgi-bin/staraniso.cgi).

| Compound | Structure | $\mathbf{K_i}$ ($\mu$M) | $\mathbf{K_d}$ ($\mu$M) |
|---|---|---|---|
| **iFSP1** | 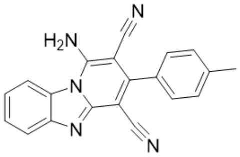 | $3.29 \pm 0.48^b$ | 0.255 |
| **viFSP1** | 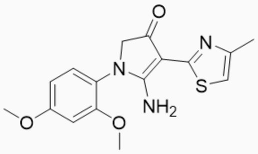 | $0.446 \pm 0.07^c$ | 0.199 |
| **FSEN1** | 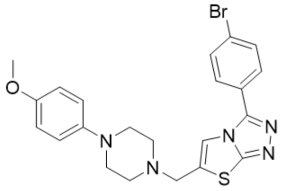 | $0.315 \pm 0.03^d$ | 0.319 |
| **afi-FSP1-1** | 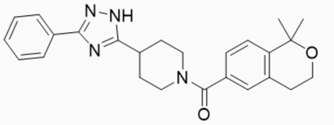 | $0.283 \pm 0.05$ | 0.098 |
| **afi-FSP1-2** | 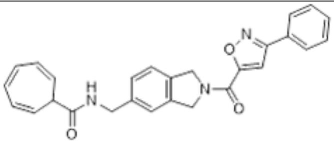 | $0.777 \pm 0.22$ | 0.266 |
| **afi-FSP1-3** | 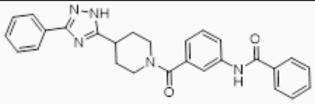 | $0.520 \pm 0.11$ | 0.184 |
| **afi-FSP1-4** | 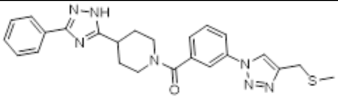 | $0.284 \pm 0.07$ | 0.198 |

**Supplementary Table 5. $K_i$ Values ($\mu$M) and $K_d$ Values ($\mu$M) of FSP1 Inhibitors.** $K_i$ values were measured by monitoring NADH oxidation via absorbance at 340 nm (pH 7.2), using fixed concentrations of 100 $\mu$M coenzyme $Q_1$ and 0.1–0.2 $\mu$M FSP1. Compounds were tested from micromolar to low nanomolar concentrations. Data are shown in Supplementary Fig. 18a–c. The reported $IC_{50}$ for iFSP1 against human FSP1 in the resazurin assay is 0.103 $\mu$M (2). We measured 0.104 $\mu$M for the ancestral FSP1 using the same assay (Supplementary Fig. 18d). For viFSP1, the reported $IC_{50}$ is 0.083 $\mu$M (45); we measured 0.081 $\mu$M for the ancestral protein (Supplementary Fig. 18e). For FSEN1, the reported $IC_{50}$ against human FSP1 is 0.313 $\mu$M (44).

| Mutant | Condition | $\mathbf{k_{cat}}$ (s$^{-1}$) | $\mathbf{K_M}$ ($\mu$M) | $\frac{\mathbf{k_{cat}}}{\mathbf{K_M}}$ (s$^{-1}\mu$M$^{-1}$) |
|---|---|---|---|---|
| **F15A** | Varying NADH, fixed coenzyme $Q_1$ | $0.48 \pm 0.02$ | $8.35 \pm 2.15$ | 0.06 |
| | Varying coenzyme $Q_1$, fixed NADH | $0.52 \pm 0.03$ | $9.48 \pm 2.55$ | 0.05 |
| **L323A** | Varying NADH, fixed coenzyme $Q_1$ | $10.88 \pm 0.57$ | $45.23 \pm 6.90$ | 0.24 |
| | Varying coenzyme $Q_1$, fixed NADH | $12.80 \pm 0.95$ | $16.20 \pm 4.23$ | 0.79 |
| **F354A** | Varying NADH, fixed coenzyme $Q_1$ | $7.06 \pm 1.25$ | $36.59 \pm 17.58$ | 0.19 |
| | Varying coenzyme $Q_1$, fixed NADH | $10.05 \pm 1.34$ | $26.85 \pm 10.62$ | 0.37 |

**Supplementary Table 6. Biochemical Analysis of F15A, L323A and F354A FSP1 Mutants: Steady-State Kinetics.** NADH oxidation was monitored by absorbance at 340 nm (pH 7.2) using 0.2–0.5 $\mu$M FSP1. Fixed concentrations for each assay were 100 $\mu$M NADH or coenzyme $Q_1$, depending on the variable substrate. Both mutants retain enzymatic activity, though F15A exhibits reduced $k_{cat}$ and L323A and F345A show increased $K_M$.

| Compound | Structure | F15A $IC_{50}$ ($\mu$M) | L323A $IC_{50}$ ($\mu$M) |
|---|---|---|---|
| **afi-FSP1-1** | 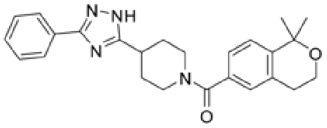 | No inhibition | No inhibition |
| **afi-FSP1-2** | 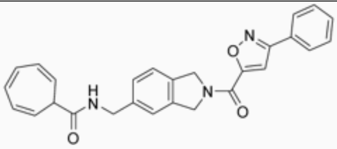 | $1.07 \pm 0.07$ | $0.490 \pm 0.03$ |
| **afi-FSP1-3** | 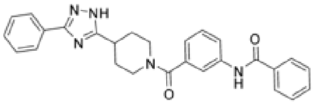 | $3.03 \pm 0.08$ | No inhibition |
| **afi-FSP1-4** | 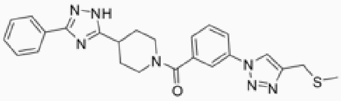 | No inhibition | No inhibition |
| **afi-FSP1-5** | 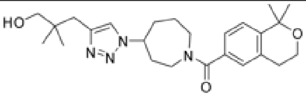 | No inhibition | No inhibition |
| **afi-FSP1-6** | 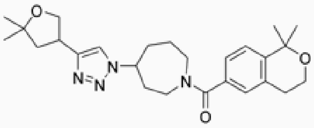 | No inhibition | No inhibition |

**Supplementary Table 7. F15A and L323A FSP1 Mutants $IC_{50}$.** F15A and L323A mutants target residues involved in inhibitor binding, as revealed by crystal structures with compounds afi-FSP1-1, afi-FSP1-3, and afi-FSP1-4. Mutations abolish binding in most cases, except for afi-FSP1-3 (F15A retains binding) and compound afi-FSP1-2, which inhibits both mutants, suggesting a distinct binding mode or reduced reliance on the mutated side chains. Activities were measured by fluorescence-based resorufin formation at pH 7.2, using 100–200 $\mu$M NADH, 50 $\mu$M resazurin, and 0.1–0.5 $\mu$M FSP1. Compounds were tested across micromolar to low nanomolar concentrations. Experiments were performed with the racemic mixture of compound afi-FSP1-5 and the diastereomeric mixture of compound afi-FSP1-6. None of the compounds inhibited F354A.

| Compound | Structure | $K_i$ ($\mu$M) |
|---|---|---|
| **afi-FSP1-1** | 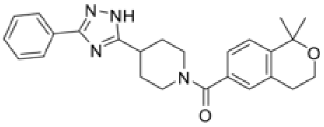 | $3.56 \pm 0.82$ |
| **afi-FSP1-2** | 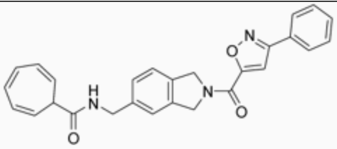 | $3.02 \pm 0.78$ |
| **afi-FSP1-3** | 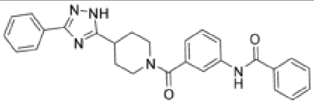 | $\sim 5.41 \pm 2$ |
| **afi-FSP1-4** | 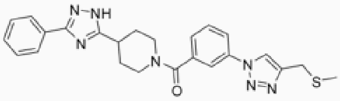 | No inhibition |

**Supplementary Table 8.** $K_i$ values ($\mu$M) of selected FSP1 inhibitors against NADH oxidase activity. Measurements were performed by monitoring NADH oxidation at 340 nm (pH 7.2), using inhibitor concentrations in the micromolar range and 1 $\mu$M FSP1. Data are shown in Supplementary Fig. 25. The $K_i$ value for compound afi-FSP1-3 is an estimate due to poor solubility above 20–50 $\mu$M.

| Compound | Structure | $IC_{50}$ ($\mu$M) |
|---|---|---|
| **afi-FSP1-1** | 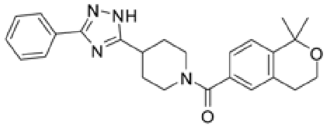 | $0.382 \pm 0.07$ |
| **afi-FSP1-2** | 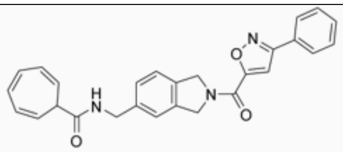 | $0.443 \pm 0.22$ |
| **afi-FSP1-3** | 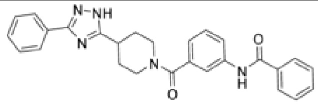 | $0.912 \pm 0.13$ |
| **afi-FSP1-4** | 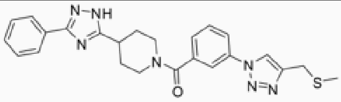 | $0.534 \pm 0.06$ |
| **afi-FSP1-5** | 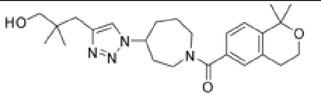 | $6.43 \pm 0.10$ |
| **afi-FSP1-6** | 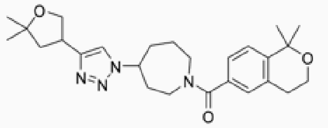 | $21.5 \pm 0.10$ |

**Supplementary Table 9. $IC_{50}$ Values ($\mu$M) of the Most Potent FSP1 Inhibitors.** The experiments were conducted using the racemic mixture of compound afi-FSP1-5 and the diastereomeric mixture of compound afi-FSP1-6. Activities were measured by monitoring fluorescence-based resorufin formation at pH 7.2, with fixed NADH concentrations (100–200 $\mu$M), 50 $\mu$M resazurin, and 0.05–0.1 $\mu$M FSP1. Inhibitors were tested in the low nanomolar to micromolar range. Full data are shown in Supplementary Fig. 26.

#### H ADME analysis of afi-FSP1-1, afi-FSP1-2, afi-FSP1-3, afi-FSP1-4

We performed a standard panel of in vitro Absorption, Distribution, Metabolism, and Excretion (ADME)/developability assays to assess oral bioavailability potential, including aqueous solubility, PAMPA permeability, buffer stability, and metabolic stability in mouse and human liver microsomes. Afi-FSP-1 and afi-FSP-4 were highly soluble (>75%), afi-FSP-3 showed modest solubility (32%), and afi-FSP-2 was poorly soluble. Consistent with these results, afi-FSP-1, -3, and -4 showed moderate PAMPA permeability (40–55% recovery), whereas afi-FSP-2 showed no measurable permeability (Supplementary Table 11). All four compounds were stable in buffer ($t_{1/2}$ >48 h). In mouse microsomes, afi-FSP-2 and afi-FSP-3 showed moderate metabolic stability ($t_{1/2}$ = 0.39 and 0.29 h, respectively), while afi-FSP-1 and afi-FSP-4 cleared rapidly ($t_{1/2}$ = 0.02 h). In human liver microsomes, afi-FSP-3 showed excellent stability ($t_{1/2}$ >2 h; scaled intrinsic clearance = 594.6 mL/min/kg) (Supplementary Table 12). Overall, afi-FSP-3 is the most promising starting point for a medicinal chemistry campaign, combining modest solubility with moderate permeability and excellent human metabolic stability. Afi-FSP-1 is a secondary candidate, offering superior solubility and permeability but requiring substantial improvements in metabolic stability to support preclinical development.

##### ADME Methods

##### Aqueous Solubility Assay

Solubility assays were performed using a Biomek FX lab automation workstation (Beckman Coulter, Inc., Fullerton, CA) and $\mu$SOL Evolution software (pION Inc., Woburn, MA). In a 96-well microplate (Cat. No. 3363, Corning Incorporated, Salt Lake, UT), 10 $\mu$L of a 1 or 10 mM test compound stock in DMSO was added to 190 $\mu$L of 1-propanol (Thermo Scientific, spectroscopy grade, Cat. No. 434360010, Fair Lawn, NJ) to create a reference stock plate. From this reference stock plate, 5 $\mu$L aliquots were mixed with 70 $\mu$L of 1-propanol and 75 $\mu$L of Dulbecco's Phosphate-Buffered Saline (DPBS, 1×; Gibco™, Cat. No. 14190-144, Thermo Fisher Scientific, Waltham, MA) to generate the reference plate.

In a 96-well storage plate (Cat. No. 201276-100, Agilent, Santa Clara, CA), 6 $\mu$L of the test compound stock was added to 600 $\mu$L of DPBS, mixed, sealed, and incubated at room temperature for 18 hours. Solutions were then filtered through a 96-well filter plate (Cat. No. 110037, pION Inc., Woburn, MA). Fractions collected from the filtered sample plate were diluted with 1-propanol at a 1:1 (v/v) ratio to generate the sample plate. Both the reference and sample plates were analyzed by UV spectrometry (SpectraMax Plus 384 Microplate Reader, Molecular Devices, LLC, San Jose, CA). Concentration calculations and solubility values ($\mu$g/mL) were determined using $\mu$SOL Evolution software. All compounds were tested in triplicate wells.

Concentrations in the reference and sample plates were also assessed using UPLC-SQD-MS (Waters, Milford, MA). Solubility ($\mu$M) was determined using the following equation:

$$\text{Solubility} = \frac{\text{Peak Area}_{\text{sample}}}{3} \times \frac{C_0}{\text{Peak Area}_{\text{reference}}}$$

where $C_0$ is the concentration of the DMSO stock solution divided by 100. All compounds were tested in triplicate wells.

##### Parallel Artificial Membrane Permeability Assay (PAMPA)

Six microliters of 1 or 10 mM compound in DMSO were dispensed into each well of a storage plate. The compounds were diluted 200-fold in DPBS to generate 600 µL spiked solution, and 180 µL of that was transferred to the donor plate (Preloaded stirwell PAMPA sandwich, Cat. No. 120551, pION Inc.). An artificial lipid (GIT-0 lipid solution, Cat. No. 110669)–coated acceptor plate containing 200 µL of acceptor sink buffer (Cat. No.110139, pION Inc.) was then placed over the donor plate to form the PAMPA sandwich. Plates were placed on the Gut-Box and stirred at room temperature for 30 min. Fractions were collected from both the donor and acceptor plates. Compound concentrations were quantified by UV spectrometry. All compounds were tested in triplicate wells.

##### Liver Microsomes Stability Assay

NADPH regenerating agent solutions A and B (Corning Gentest$^{TM}$, Cat. No: 451220) were purchased from Discovery Labware (Woburn, MA). Liver microsomes of mouse (CD-1, pooled, GibcoTM, Cat. No: MSMCPL) and human (mixed gender, pool of 50, Cat. No. H0630, Xenotech) were obtained from Thermo Fisher Scientific. Stock solutions of test compounds were prepared at 1 or 10 mM in DMSO. Sample preparation involved transferring 50 nL of stock solutions into six 96-well microplates (0, 0.25, 0.5, 1, and 2 hours) using a Labcyte Echo 650 acoustic dispenser. All compounds were tested in triplicate per time point. The reaction solution contained mouse liver microsomes (20 mg/mL), EDTA, NADPH regenerating agent, and 0.1 M potassium phosphate buffer (pH 7.4). Final concentrations applied: microsome protein 0.5 mg/mL, compound 1 or 5 µM, NADPH A 1.3 mM, NADPH B 0.4 U/mL. Reaction plates were incubated at 37°C on an orbital shaker (100 rpm) and quenched with cold acetonitrile with internal standard. Supernatants were analyzed using UPLC–MS/MS. Metabolic stability was determined via first-order kinetics.

#### PBS Stability Assay

Compound degradation in DPBS was monitored at 0, 1, 3, 6, 24, and 48 hours. Stock solutions were dispensed into triplicate wells of a 96-well microplate using the Labcyte Echo 650 dispenser. DPBS was added to achieve a final compound concentration of 2 µM. Plates were incubated at 37°C on an orbital shaker and quenched with cold acetonitrile with internal standard. Supernatants were analyzed using UPLC-MS/MS. PBS stability was evaluated by first-order kinetics.

#### UPLC/UV/MS System

Chromatographic separation was conducted using a Waters Acquity UPLC system coupled to both UV detector and single quadrupole mass spectrometer. Conditions were set as reported previously.

#### UHPLC-MS/MS System

Separation was performed on an Acquity UPLC BEH C18 1.7 µm, 2.1 x 50 mm column (Waters Corporation, Milford, MA) using a Sciex ExionLC™ with 6500+ Qtrap system. Data were acquired using Analyst v1.7 and analyzed using OS software. Column temperature: 55 °C. Mobile phase A: 0.1% formic acid in MilliQ H2O; Mobile phase B: 0.1% formic acid in acetonitrile. Flow rate: 0.9 mL/min with gradient: 0–0.2 min, B 1–1%; 0.2–0.5 min, B 1–50%; 0.5–1.6 min, B 50–95%; 1.6–1.95 min, B 95–95%; 1.95–1.96 min, B 95–1%; 1.96–2.2 min, B 1–1%. Mass spectrometer operated in positive-ion mode; MRM transitions for liver microsome and PBS stability assays were applied.

| Compound | m/z (Q1 > Q3) | DP (V) | CE (V) | CXP (V) |
|---|---|---|---|---|
| SJ001127367-1 | 411 > 300 | 120 | 32 | 9 |
| SJ001127368-1 | 411 > 300 | 110 | 31 | 9 |
| SJ001127369-1 | 408 > 297 | 130 | 35 | 20 |
| SJ001127371-1 | 408 > 297 | 110 | 33 | 9 |

**Supplementary Table 10.** MRM transitions used for analysis of the 4 FSP1 hits.

| Compound | Avg. Solubility (UV, µM) | % Solubility | Avg. Solubility (LC/MS, µM) | Avg. Permeability (PAMPA, $10^{-6}$ cm/s) | % Recovery |
|---|---|---|---|---|---|
| afi-FSP1-1 | 76.5 ± 2.1 | 77.3 | 44.2 ± 0.7 | 885.7 ± 138.7 | 54.4 ± 1.6 |
| afi-FSP1-2 | 1.02 ± 0.3 | 1.0 | 0.4 ± 0.1 | 0 | 0 |
| afi-FSP1-3 | 31.1 ± 0.9 | 31.4 | 25.6 ± 0.2 | 197 ± 81.6 | 46.7 ± 17.6 |
| afi-FSP1-4 | 75.42 ± 4.7 | 76.2 | 64.2 ± 2.9 | 240.1 ± 115.1 | 42.7 ± 17.4 |
| Albendazole | 2.4 ± 0.2 | 2.5 | 1.9 | NA | NA |
| Carbamazepine | 92.5 ± 6.6 | 93.4 | 64.3 ± 1.5 | 68.9 ± 40.6 | 39.2 ± 5.9 |
| Verapamil | 80.3 ± 6 | 81.1 | 42.1 ± 1.1 | 1251.2 | 64.1 ± 1.1 |

**Supplementary Table 11.** Solubility and PAMPA Permeability of afi-FSP1 compounds: Aqueous kinetic solubility was determined by UV spectrophotometry in Biomek automated 96 well format and by LC–MS–based quantitative analysis; values are reported as mean solubility (µM) ± SD from $n = 3$ independent determinations and as % solubility. PAMPA permeability (Pe, $10^{-6}$ cm/s) was measured across an artificial lipid membrane, and data are shown as mean ± SD ($n = 3$); % recovery is also calculated. Albendazole, carbamazepine, and verapamil were included as reference compounds to benchmark low, moderate, and high permeability/solubility behavior.

| Compound | PBS Stability $t_{1/2}$ (hr) | Mouse Metabolic $t_{1/2}$ (hr) | Mouse Metabolic $CL_{int}$ (mL/min/kg) | Human Metabolic $t_{1/2}$ (hr) | Human Metabolic $CL_{int}$ (mL/min/kg) |
|---|---|---|---|---|---|
| afi-FSP1-1 | >48 | 0.02 | 2320.40 | 0.08 | 267.4 |
| afi-FSP1-2 | >48 | 0.39 | 253.90 | NA | NA |
| afi-FSP1-3 | >48 | 0.29 | 340.36 | >2 | 594.6 |
| afi-FSP1-4 | >48 | 0.02 | 4343.87 | 0.07 | 409.5 |
| Verapamil | NA | 0.01 | 381.65 | 0.17 | 111.8 |

**Supplementary Table 12.** In vitro metabolic stability of afi-FSP1 compounds in Mouse and Human liver microsomes: Apparent in vitro half life ($t_{1/2}$, hr) and intrinsic clearance ($CL_{int}$, mL/min/kg) were determined from the depletion of parent compound in pooled human (HLM) and mouse liver microsomes (MLM) in the presence of NADPH. Data are reported as mean $t_{1/2}$ in hours from $n = 3$ incubations. Reference compound Verapamil is used as a high-clearance control.

#### I Experimental Validation of AdaptiveFlow using PARP1

**I.1 Targeting PARP1.** To evaluate the effectiveness of the AdaptiveFlow platform and the methodology presented, we have conducted a thorough experimental validation using Poly(ADP-ribose) polymerase 1 (PARP1) as our target. PARP1 is a nuclear enzyme critical in the DNA repair process, particularly in the base excision repair pathway. PARP1 is vital for maintaining genomic integrity by identifying and repairing DNA damage, such as single-strand breaks (146). Structurally, PARP1 is composed of distinct domains: a) a DNA-binding domain that enables it to recognize and bind to damaged DNA, a pivotal step for initiating the repair process; b) an auto-modification domain where it adds poly(ADP-ribose) chains to itself, thus regulating its activity and influencing interactions with other cellular components; and c) a catalytic domain that includes an alpha-helical subdomain (HD) and an ADP ribosyl transferase subdomain (CAT), the site of enzymatic activity (Fig. 33(a)) (147, 148). PARP1 uses $NAD^+$ to transfer ADP-ribose units to target proteins, a critical process for DNA repair and various cellular responses (147). In recent years, PARP1 has gained significant attention in drug development and cancer therapy. Pharmacological inhibitors of PARP1 have been developed, mainly for treating cancers with deficiencies in homologous recombination DNA repair, such as those with BRCA mutations (3, 149). Clinical trials have confirmed the efficacy of PARP inhibitors in treating several types of cancer, including ovarian, breast, and prostate cancer. FDA-approved inhibitors, such as olaparib, rucaparib, and niraparib, exemplify this. Current research is extending their potential use in other types of cancer and in combination therapies (150).

**I.2 Applying AdaptiveFlow to PARP1.** To assess the effectiveness of AdaptiveFlow, our analysis was centered on two distinct crystal structures of the PARP1 enzyme in complex with different inhibitors localized within its catalytic domain. Our *in silico* screening approach specifically harnessed the crystal structure of PARP1 conjugated to olaparib (PDB 7AAD) (148) in the first docking scenario. This included both the active site and the regulatory helical domain (HD). Conversely, the second scenario employed the crystal structure of PARP1 complexed with the inhibitor UTT57 (PDB 6NRG) (151), where the HD subdomain was excluded. For docking processes, QuickVina 2 (67) served as the computational tool, alongside the application of the ATG-VS method for both scenarios. Following the *in silico* screening, we established rigorous selection criteria, enabling the identification of 80 promising candidates from each scenario for further experimental verification, as detailed in the Methods section (*Methods: Prioritizing Hits for Experimental Validation*). The preliminary evaluation comprised a PARP1 activity-based assay, which was then complemented by validation of selected hits through protein NMR, protein crystallography, and an examination of cellular activity via a colony formation assay (Fig. 27a).

**I.3 Selection and Characterization of Initial PARP1 Inhibitor Compounds.** The initial phase of our experimental validation involved a dual-concentration enzymatic assay, conducted at $5\mu M$ and $50\mu M$, to measure the inhibitory activity of 160 synthesized compounds (Fig. 27a). Through this assay, we performed duplicate experiments to ensure reproducibility and identified seven compounds that exhibited a notable inhibition of PARP1 activity, with an inhibition percentage exceeding 50% at the lower concentration $5\mu M$. Notably, three compounds were associated with the first docking scenario, while the remaining four were aligned with the second scenario. Importantly, all seven compounds (Fig. 27b) demonstrated compatibility with the assay conditions, showing no signs of interference. To quantify the potency of these inhibitors, we proceeded with determining the half-maximum inhibitory concentration ($IC_{50}$) for each compound. The calculated $IC_{50}$ values are the result of three independent experiments and range from as low as 0.84 nM to approximately 1500 nM, indicating varying degrees of inhibitory efficacy between compounds. To further investigate iParp1 as a strong PARP1 inhibitor against the reference compound olaparib, we performed a complementary intracellular NanoBRET target-engagement assay and derived apparent $K_i$ values by competitive tracer displacement, yielding $K_i = 5.49 \pm 1.68\,\mathrm{nM}$ for iParp1 and $K_i = 6.73 \pm 0.35\,\mathrm{nM}$ for olaparib (Supplementary Fig. 27b, Supplementary Table 13). Representative multi-tracer dose–response curves and the corresponding linear analysis of $IC_{50}$ versus tracer concentration used to extract $K_{i,\mathrm{app}}$ are provided in Supplementary Fig. S27.

| Ligand | Docking Model (PDB) | $IC_{50}$ Exp 1 (nM) | $IC_{50}$ Exp 2 (nM) | $IC_{50}$ Exp 3 (nM) | Mean $IC_{50}$ ± SD (nM) | $K_i$ ± SD (nM) | SMILES |
|---|---|---|---|---|---|---|---|
| iParp1 | 6NRG | 0.544 | 0.530 | 1.44 | 0.84 ± 0.52 | 5.49 ± 1.68 | FC=1C=CC(CC2=NNC(=O)C=3C=CC=CC23)=CC1C(=O)N4CC5CC4C(=O)O5 |
| iParp2 | 6NRG | 75.5 | 42.6 | 67.0 | 61.7 ± 17.1 | — | CC1=C(NC(=O)C=2C=CC=CC12)C(=O)NCC3=NC=4C(=CN=C5C=CC=CC45)N3 |
| iParp3 | 7AAD | 780 | 837 | 700 | 772 ± 68.8 | — | O=C(N1CC2CC1CCN2C(=O)C3=NNC(=O)C=4C=CC=CC34)C=5C=CC=6NC=CC6C5 |
| iParp4 | 7AAD | 1000 | 793 | 992 | 928 ± 117 | — | CC1CC(CN1C(=O)C2=NNC(=O)C=3C=CC=CC23)NC(=O)C4=NNC=5C=CC(C)=CC45 |
| iParp5 | 6NRG | 1790 | 692 | 1360 | 1281 ± 553 | — | O=C(NC1CC2(C1)CN(C2)C(=O)C3=CC=4C=CC=CC4C(=O)N3)C5=CNN=N5 |
| iParp6 | 7AAD | 1430 | 1470 | 1350 | 1417 ± 61.1 | — | O=C(N1CC2CC1CCN2C(=O)C=3C=CC=4C=CN=CC4C3)C5=NNC(=O)C=6C=CC=CC56 |
| iParp7 | 6NRG | 809 | 808 | 766 | 794 ± 24.5 | — | CC=1NN=CC1C(=O)N2CCC(C)(C2)NC(=O)C3=CNC(=O)C=4C=CC=CC34 |
| Olaparib | - | 0.597 | 0.377 | 0.607 | 0.53 ± 0.13 | 6.73 ± 0.35 | O=C(N1CCN(CC1)C(=O)c1cc(ccc1F)Cc1n[nH]c(=O)c2c1cccc2)C1CC1 |

**Supplementary Table 13.** Independent $IC_{50}$ measurements (n=3), mean $IC_{50}$ ± SD, $K_i$ ± SD values for iparp1 and olaparib, corresponding docking model (PDB ID), and SMILES for PARP1 inhibitors.

Subsequently, based on the $IC_{50}$ values determined from triplicate dose–response experiments, we selected four inhibitors with reproducible submicromolar potency ($IC_{50}$ < 1 µM), comprising two representatives for each of the two docking scenarios/PDB structures used, for in-depth biophysical characterization (Supplementary Fig. 27a). Based on this criteria, four compounds (iParp1, iParp2, iParp3, and iParp4) were distinguished for further investigation studies. These inhibitors not only exhibited strong PARP1 inhibition in preliminary assays but also showed promise for significant biological activity, thus warranting additional validation through subsequent NMR spectroscopy, X-ray crystallography, and cell-based assays. Interestingly, hits resulting from computational screening using PDB 6NRG where the HD domain was absent, yielded the highest

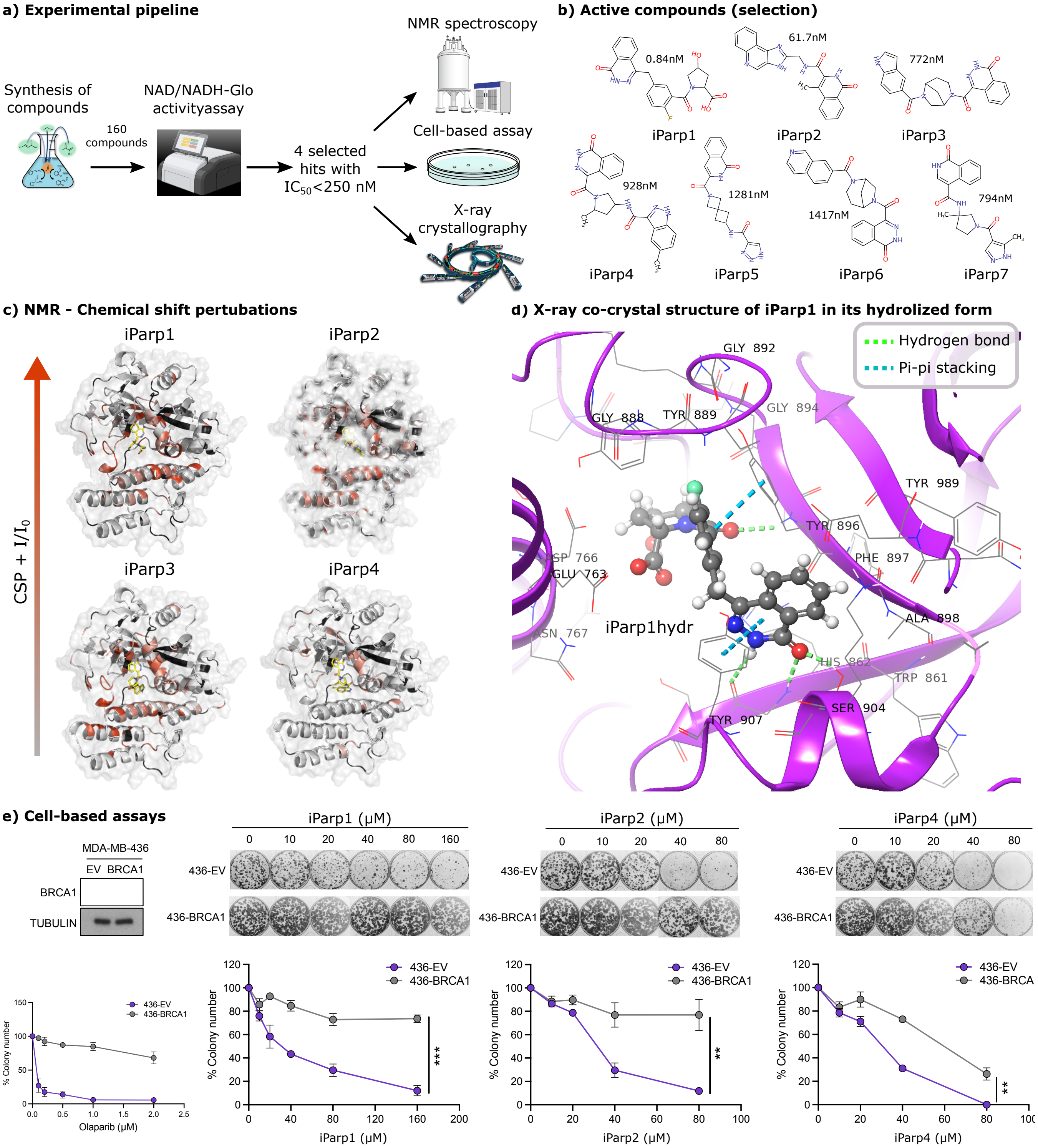

**Supplementary Fig. 27. Experimental Validation of AdaptiveFlow with Human PARP1.** **(a)** Schematic representation of the experimental workflow utilized to validate hits from virtual screening targeting the PARP1 enzyme. The pipeline integrates a suite of biochemical assays, biophysical methods, and cellular-based evaluations. **(b)** A selection of the most potent inhibitors confirmed through experimental assays, with their respective $IC_{50}$ values. **(c)** Docking poses of PARP1 with inhibitors tested by NMR spectroscopy. Residues in red showed higher chemical shift perturbations (CSPs), or stronger broadening, than the mean plus one standard deviation (S1). Shades of red depict the combined size of peak broadening and CSPs. **(d)** Ribbon representation of the co-crystal structure of iParp1 bound to PARP1, detailing the molecular interactions within the catalytic site. Ball representation of iParp1, with carbon, nitrogen, oxygen, and fluorine atoms colored in yellow, blue, red, and cyan, respectively. Hydrogen bonds and pi-pi stacking are represented as dashed blue and dashed green lines. **(e)** Cellular assays conducted on MDA-MB-436, a triple-negative breast cancer cell line, demonstrate the differential cytotoxic effects of the ligands, corroborating the targeted inhibition activity of the compounds. BRCA1 and tubulin protein levels were analyzed by western blot in BRCA1-mutated MDA-MB-436 expressing empty vector (EV) or BRCA1. Clonogenic assay of MDA-MB-436 control (436-EV) and its isogenic BRCA1-reconstituted pair (436-BRCA1) treated with the indicated compounds. Relative quantifications of colonies at the endpoint are shown at the top (n=3, mean ± SEM from three independent experiments) and representative images of the colonies are shown in the line plots.

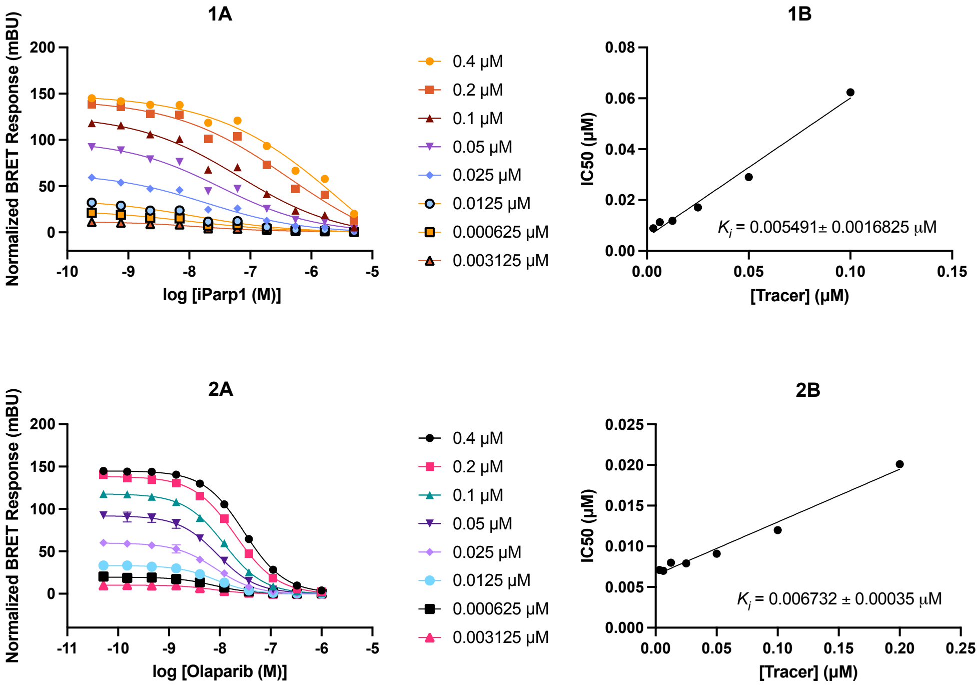

**Supplementary Fig. 28. NanoBRET target engagement for PARP1.** Representative multi-tracer dose–response curves for iParp1 and olaparib, and linear fits of $\mathrm{IC}_{50}$ versus tracer concentration used to obtain $K_{i,\mathrm{app}}$. Reported values: iParp1 $K_{i,\mathrm{app}} = 5.49 \pm 1.68$ nM; olaparib $K_{i,\mathrm{app}} = 6.73 \pm 0.35$ nM.

affinity molecules (iParp1, iParp2).

**I.4 Characterization of PARP1-Inhibitor Interactions via Protein NMR Spectroscopy.** The binding interactions between PARP1 and selected inhibitory compounds were comprehensively elucidated using protein NMR spectroscopy, revealing conclusive evidence of complex formation. We observed significant chemical shift perturbations (CSPs) and peak broadening in the PARP1 NMR spectra upon addition of each inhibitor, indicative of binding events. These CSPs were notable at both 1:1 and 1:2 molar ratios for inhibitors iParp2, iParp4, iParp1, and iParp3, as illustrated in Supplementary Fig. 27c. The perturbations predominantly occurred in regions proximal to PARP1's active site, aligning with the binding locale forecasted by our computational docking simulations (refer to Supplementary Fig. 33a and 27d). Supplementary Fig. 29 and 30 detail the specific residues impacted and the extent of chemical shift perturbations consequent to ligand association.

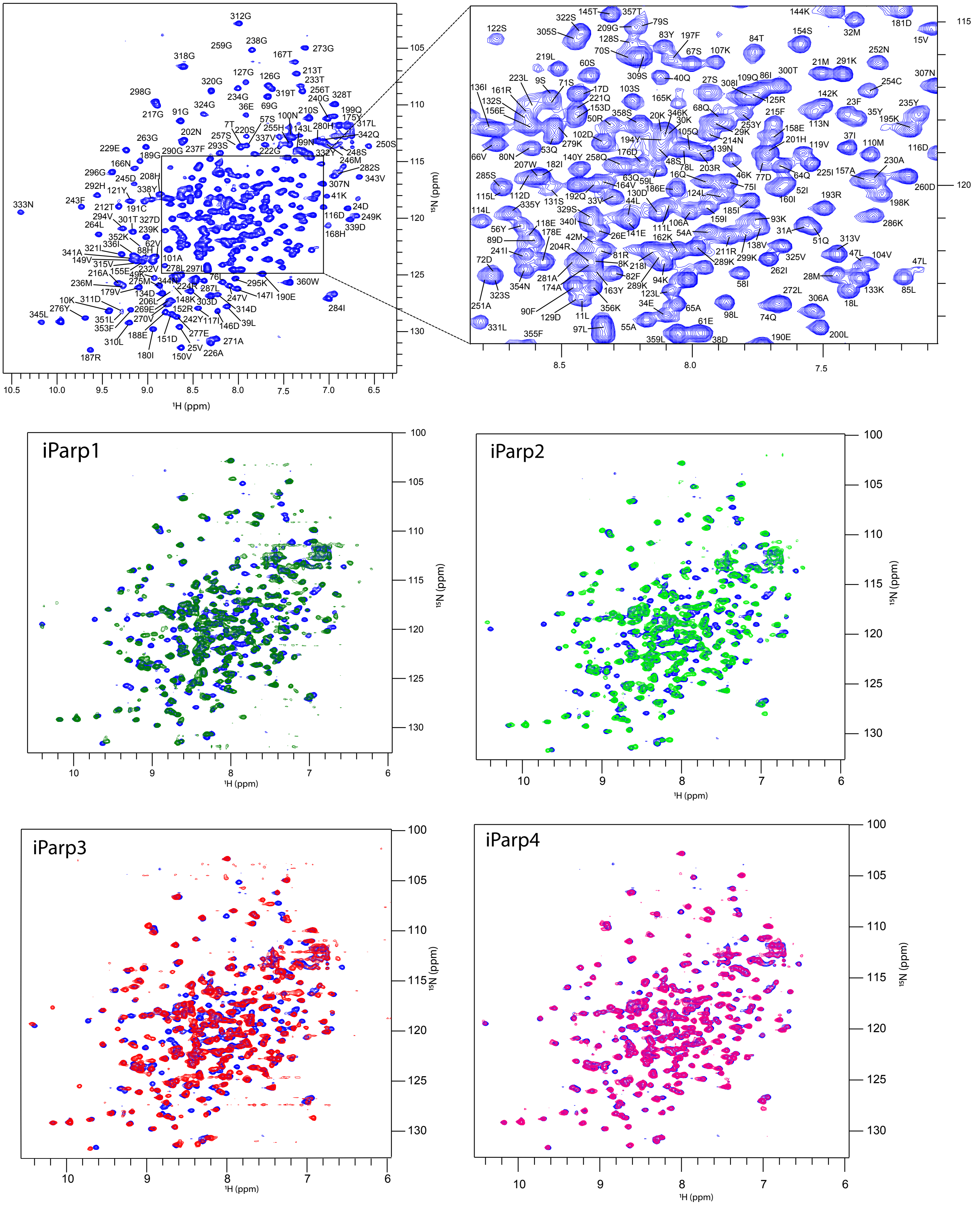

**Supplementary Fig. 29. TROSY NMR spectra illustrating the interaction of PARP1 with various inhibitors**. The assignment of resonances was based on BMRB entry 50454. Spectra are color-coded to indicate different conditions: Blue represents the apoprotein spectrum with 1% DMSO; red indicates the spectrum in the presence of inhibitor iParp3 at a 1:2 protein-to-inhibitor molar ratio; magenta corresponds to inhibitor iParp4 at a 1:1 ratio; dark green shows inhibitor iParp1 at a 1:2 ratio; and light green depicts inhibitor iPARP2 at a 1:1 ratio. The shifts in resonance peaks reflect the binding and perturbation effects of the inhibitors on the PARP1 structure.

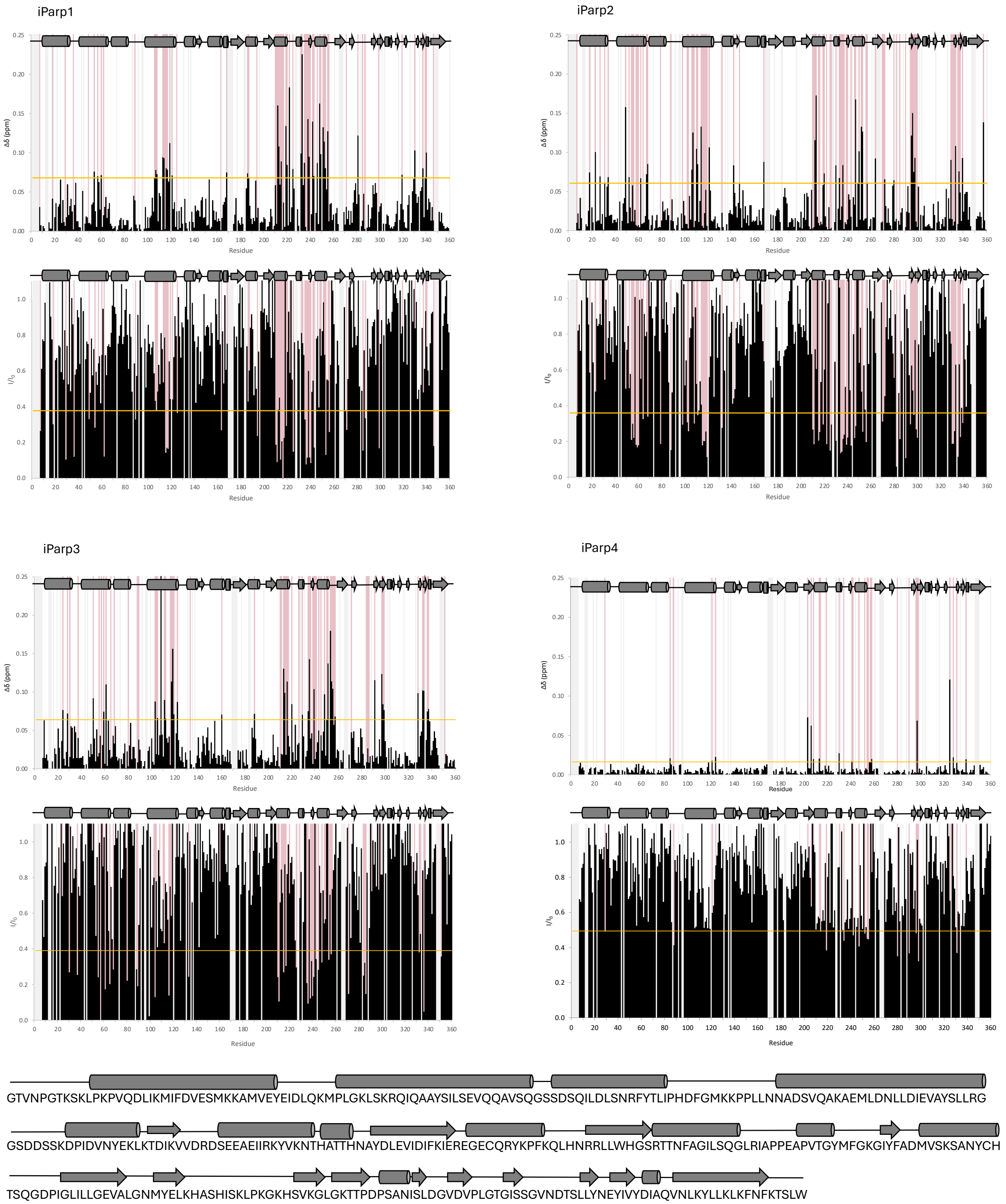

**Supplementary Fig. 30. Nuclear Magnetic Resonance (NMR) Spectral Analysis of PARP1 Protein Interaction with Distinct Inhibitors.** The graphs display the chemical shift perturbation (CSP) profiles and peak broadening profiles for the PARP1 protein upon binding with four different inhibitors: iParp1, iParp2, iParp3, and iParp4. For each inhibitor, the top panel shows the CSPs of backbone resonances in parts per million (ppm) and the bottom panel shows peak broadening ($I/I_0$). The horizontal yellow lines indicate the threshold for significant chemical shift changes, that is mean + 1 STD. Residues that exhibit CSPs above this threshold or signal intensity lower than the threshold are highlighted in red, suggesting regions of PARP1 that are affected by inhibitor binding. Grey background signifies missing assignment or proline residues. The secondary structure of PARP1 is depicted above each graph, with $\alpha$-helices represented by cylinders and $\beta$-strands by arrows. The sequence below the graphs corresponds to the PARP1 regions shown.

**I.5 Co-Crystallization Experiments..** Our crystallization experiments resulted in the determination of the crystal structure of the isolated catalytic domain of human PARP1 (PDB:8VYH) in complex with the hydrolyzed version of iParp1 at a 2.05 Å resolution (Supplementary Fig. 27d). Data collection and refinement statistics can be found in Supplementary Table 14. The unbiased electron density allowed us to model the inhibitor molecule in the enzyme active site and unambiguously show the protein complex with the hydrolyzed form of iParp1 (iParp1hydr, Supplementary Fig. 31). The observed hydrolysis can be explained by the inherent ring strain of the azabicyclo moiety. Fig. Supplementary 32 shows in detail the chemical structures and corresponding 3D structures of iParp1 and its hydrolyzed form iParp1hydr. iParp1hydr binds at the center of the active site of PARP1 performing pi-pi stacking interactions with Tyr896 and Tyr907 (Supplementary Fig. 27d). Furthermore, a hydrogen bond is established between Tyr896 and the oxygen atom close to the hydroxypyrrolidine group. An important hydrogen bond interaction with Glu988 is also observed. To ascertain if AdaptiveFlow could have identified the iParp1hydr had it been present in the library, we conducted docking studies of both iParp1 and iParp1hydr to the PARP1 crystal structure obtained. Remarkably, the docking scores for iParp1 (-12.7 kcal/mol) and iParp1hydr (-12.5 kcal/mol ) were comparable, indicating potential efficacy despite structural alterations induced by hydrolysis.

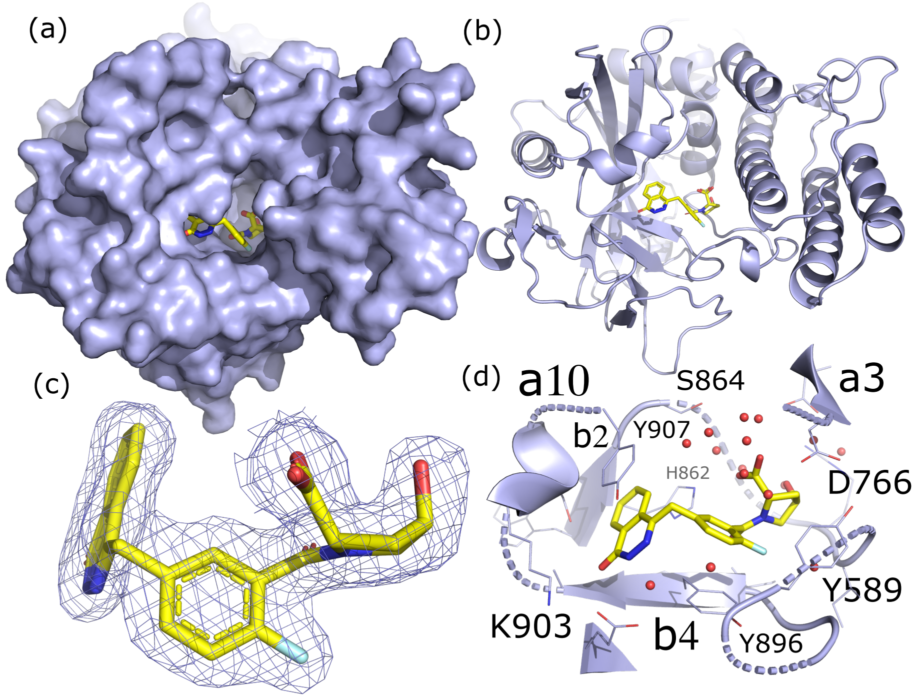

**Supplementary Fig. 31. The Crystal Structure of the PARP1 Catalytic Domain in Complex with iParp1hydr.** a, surface display of human PARP1 and the bound (carbons in yellow) inhibitor, represented in sticks. Oxygen, nitrogen, and fluorine atoms are colored in red, blue, and cyan, respectively. b, ribbon diagram (purple) of PARP1 in complex with iParp1hydr. The compound is in stick representation with carbon, nitrogen, oxygen, and fluorine atoms colored in yellow, blue, red, and cyan, respectively. c, Representation of the electron density of iParp1hydr highlighting the 4-hydroxypyrrolidine-2-carboxylic acid moiety. The compound is in stick representation with carbon, nitrogen, oxygen, and fluorine atoms colored in yellow, blue, red, and cyan, respectively. d, Representation of iParp1hydr and the interactions with its neighboring residues within the catalytic domain active site. The compound is in stick representation with carbon, nitrogen, oxygen, and fluorine atoms colored in yellow, blue, red, and cyan, respectively.

**I.6 Cellular Activity of Identified Hits.** The cellular impact of the identified hits was further delineated through clonogenic assays conducted on MDA-MB-436 cells, a triple-negative breast cancer (TNBC) line with a known BRCA1 5396 + 1G>A mutation. This particular mutation diminishes BRCA1 protein levels, leading to increased sensitivity to PARP1 inhibitors (152, 153) (Supplementary Fig. 27e). To ascertain the specificity of our compounds towards PARP1 in cancer cells, we employed MDA-MB-436 cells harboring an empty vector (control) and their isogenic counterpart expressing wildtype BRCA1. Our findings revealed that the control MDA-MB-436 cells exhibited pronounced sensitivity to iParp1 and iParp2, suggesting these compounds' efficacy in targeting PARP1 within tumor cells. Conversely, cells reintroduced with wildtype BRCA1 exhibited resistance to these inhibitors, underscoring the specificity of iParp1 and iParp2 for PARP1-mediated pathways in the context of BRCA1 deficiency (Supplementary Fig. 27e). iParp4 presented a concentration-dependent inhibitory effect on 436-EV cells, with significant cytotoxicity observed even in 436-BRCA1 cells at higher doses, suggesting a broader range of activity that may transcend PARP1 inhibition. Notably, iParp3 did not exert significant cytotoxicity in either cell line, aligning with the hypothesis of its limited utility in targeting PARP1. Overall, the observed effect of the tested inhibitors on the MDA-MB-436 cells was less significant than the affinity observed with our biochemical assay, probably due to the molecules' limited cell penetration profiles. Such is the case of iParp1, which in its hydrolyzed version bears a carboxylic acid, which

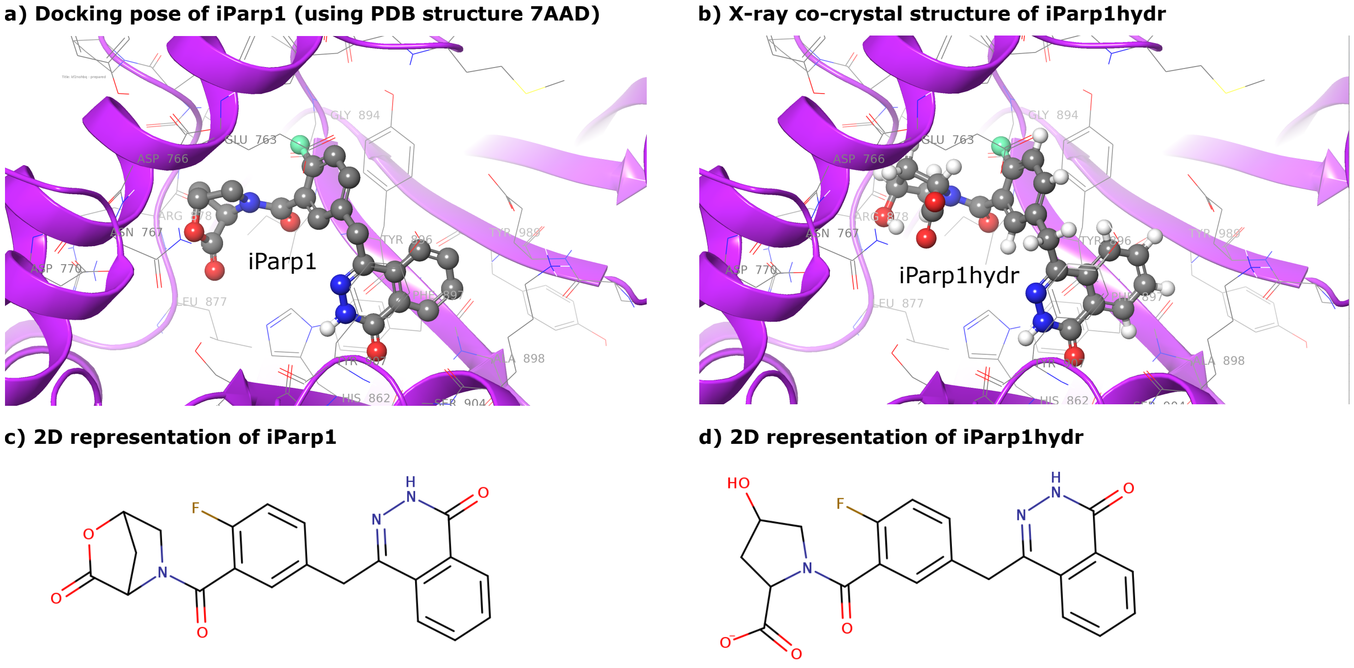

**Supplementary Fig. 32. Chemical and Structural Characterization of iParp1 and its Hydrolyzed Form iParp1hydr.** a, Ribbon representation of the docking pose of the virtual screened iParp1 bound to PARP1. The in-silico screening was performed using the PDB structure 7AAD. Ball representation of iParp1, with carbon, nitrogen, oxygen, and fluorine atoms colored in yellow, blue, red, and cyan, respectively. Hydrogen bonds and pi-pi stacking are represented as dashed blue and dashed green lines. b, Ribbon representation of the co-crystal structure of iParp1hydr with PARP1. Ball representation of iParp1, with carbon, nitrogen, oxygen, and fluorine atoms colored in yellow, blue, red, and cyan, respectively. Hydrogen bonds and pi-pi stacking are represented as dashed blue and dashed green lines. c, Chemical structure of iParp1. d, Chemical structure of iParp1hydr..

consequently leads to poor cell permeability.

***Statistical Analysis.*** Colony formation data are represented as mean ± SEM from three independent experiments, and significance was determined by unpaired, two-tailed Student's t-test.

**I.7 Comparison to Approved Drugs.** A noteworthy outcome of the *in silico* screening conducted using AdaptiveFlow was the identification of a few hits resembling olaparib, an FDA-approved PARP inhibitor. Despite no prior information on known inhibitors being used by AdaptiveFlow during the screens, VirturalFlow A2I was able to discover a few hits (iParp1,iParp3, and iParp6) that are similar to olaparib in structure and also potency. iParp1 has a similar $IC_{50}$ value than olaparib (single digit nanomolar). Olaparib has undergone extensive classical medicinal chemistry during its development, while our compounds come directly from a virtual screen without any optimization, highlighting the efficiency of our approach and increasing the validity of the discovered ligands. All our compounds exhibit distinctiveness in comparison to previously reported inhibitors, and the inhibitors discovered herein through AdaptiveFlow are not covered by prior patents. Utilizing Morgan fingerprints (154) to assess structural similarity, it was observed that among our discovered inhibitors, iParp1, iParp3, and iParp6 exhibited Morgan fingerprint similarities of 0.58, 0.40, and 0.39, respectively. Conversely, iParp2, iParp4, iParp5, and iParp7 are rather different from olaparib, highlighting that our platform can discover novel and diverse sets of molecules with a range of structural features. Olaparib, characterized by its N-acylpiperazine structure with a cyclopropane group and a monofluoro-benzene on an oxo-phthalazine scaffold, shares the oxo-phthalazine core with the first docking scenario series hits, iParp1 and iParp5. This core is altered to a hydroxyisoquinoline in iParp2 and iParp7, while the cyclopropyl N-acylpiperazine feature is replaced with acylated 2,6-diazabicyclo[3.2.1]octane in iParp6 and iParp3. Notable is the modification in iParp1, where the N-acylpiperazine is substituted with a bicyclic lactone prone to hydrolysis into a stable hydroxyproline derivative, a transformation confirmed through NMR and mass spectroscopy and observable in the crystal structure. Remarkably, the docking scores for iParp1 (-12.7 kcal/mol) and iParp1hydr (-12.5 kcal/mol) were comparable, indicating potential efficacy despite structural alterations induced by hydrolysis.

**I.8 Reagents.** Sodium chloride, hepes buffer, phenylmethylsulfonyl fluoride, glycerol, SigmaFast™ Protease Inhibitor Cocktail Tablets EDTA-Free, Ethylenediaminetetraacetic acid (EDTA), tris(2-carboxyethyl) phosphine (TCEP), and dithiothreitol (DTT) were purchased from Sigma-Aldrich. All tested compounds were purchased from the ChemSpace/Enamine supplier. NAD/NADH-Glo assay kit was purchased from Promega (Madison, WI). [$^{15}$N] ammonium chloride, U-$^{13}C_6$D-glucose, alpha-ketobutyric acid-$^{13}$C4,3,3-D2 and alpha-ketoisovaleric acid-U-$^{13}C_5$ were purchased from Cambridge Isotope Laboratories.

alpha/beta-Tubulin Antibody was purchased from Cell Signaling Technology (# 2148S).

**I.9 Receptor Preparation.** The receptor structures for the first and second docking scenarios have PDB ID 7AAD and 6NRG, respectively. The HD domain was present in the first docking scenario, and was absent in the second docking scenario. The structures were prepared with the Protein Preparation Wizard of Maestro by Schrödinger (hydrogens, protonation states, bond orders) (155). MGLTools with AutoDockTools was used to convert the receptor PDB files into PDBQT files, and to set up the docking boxes (156).

**Docking Scenario Setup.** The protein was set up to be rigid during the docking, and QuickVina 2 was used as the docking program for both docking scenarios and both screening stages (ATG prescreen and ATG primary screen). The catalytic domain was targeted in both docking scenarios. In the first docking scenario (with HD domain), the docking box had a size of 20.0 × 16.0 × 16.0 Å, while in the second docking scenario (without HD domain) the docking box had a size of 16.0 × 24.0 × 20.0 Å. In the ATG prescreen, one representative molecule was chosen per tranche, resulting in approximately 12 million compounds in total. During the ATG primary screen, a total of 100 million compounds were screened.

**I.10 Prioritizing Hits for Experimental Validation.** Postprocessing of the results was done with DataWarrior. Basic filters have been applied, such as logP $< 5$, and compounds with problematic functional groups and predicted toxicity (mutagenic compounds, compounds with reproductive effects, tumorigenic compounds) were removed. In addition to the ranking by docking score (compounds displayed docking scores as high as -15.1 kcal/mol), we implemented a filtering and prioritization routine to identify a set of diverse drug-like compounds within the hit list (Supplementary Fig. 27b) (157). The hits were filtered by their aqueous solubility, setting a clogS of at least -5 (10 µM). Many studies have suggested that the aromatic proportion of a molecule has a detrimental impact on solubility (158, 159); for this reason, we filtered out compounds displaying aromaticity higher than 75%. In parallel, the flatness of the library was further assessed by leveraging the importance of an increased number of $sp^3$ hybridized carbons in the increase of positive receptor-ligand interactions, as well as leading to increased solubility (28). As such, the degree of saturation ($F_{sp3}$) was calculated (28), and compounds with values ranging from 0.1–0.4 were given preference (160–162). Finally, clustering was performed using the DataWarrior software to obtain a diverse set of compounds for experimental validation. Supplementary Fig. 33b summarizes the filtering pipeline implemented in this study. All the compounds from the study were provided by Enamine Ltd. (Kyiv, Ukraine).

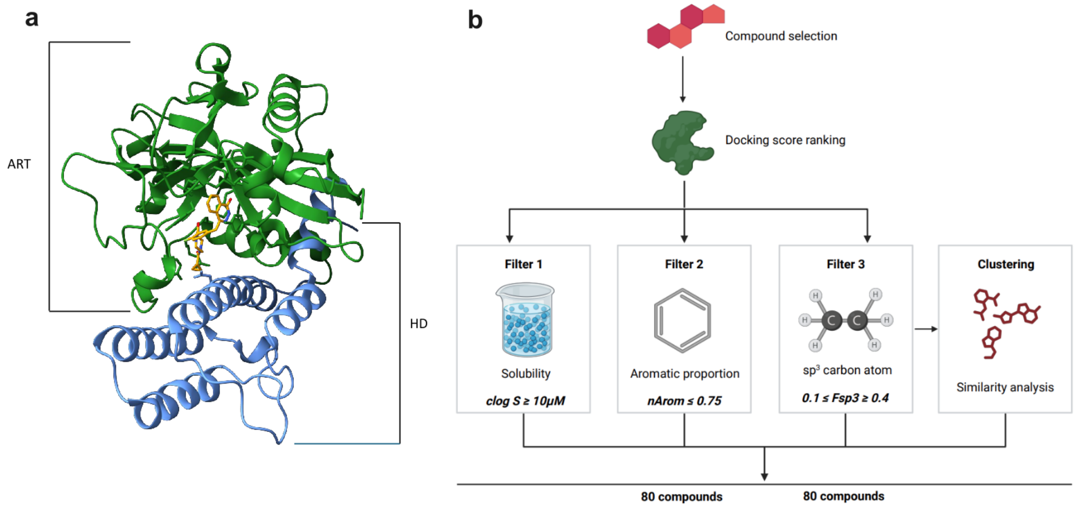

**Supplementary Fig. 33.** a. Model of PARP1 catalytic domain highlighting the alpha-helical subdomain (HD) in blue and the ADP ribosyl transferase subdomain (CAT), in green. The structure is displayed with the inhibitor olaparib (in yellow) bound at the active site. b. Graphic representation of the pipeline used for compound filtering – besides docking score ranking, water solubility, and molecule flatness were considered. The compounds were filtered for nasty functional groups and pan-assay interference compounds (PAINS). Clustering analysis was applied using the DataWarrior software.

**I.11 Protein Expression and Purification.** PARP1 catalytic domain (residues 661–1014, full-length CAT) was expressed as described (163, 164). Unlabeled and $^{15}$N-labled PARP1 was expressed using *E. coli* BL21(DE3) cells in Lysogeny Broth (LB) or M9 medium, respectively, supplemented with 50 $\mu g$/ml kanamycin. Protein expression was induced with 0.2 mM isopropyl $\beta$-D-1-thiogalactopyranoside (IPTG) and performed at 16°C overnight.
The labeled and unlabeled PARP1 catalytic domain was purified by incubation with nickel-nitriloacetic acid (Ni-NTA) agarose resin (Qiagen) for 3h at 4 °C. The resin was washed with 50 ml HEPES, pH 8.0, 500 mM NaCl, and 0.5 mM TCEP. The protein was eluted with 50 ml HEPES, pH 8.0, 100 mM NaCl, 300 mM imidazole, and 0.5 mM TCEP and injected into a Resource Q column for DNA separation. Unlabeled protein was purified by size exclusion chromatography in 50 mM HEPES, pH 8.0, 150

mM NaCl, and 0.5 mM TCEP. PARP1 catalytic domain purified for NMR labeled purposes was purified using 10 mM sodium phosphate, 222 mM sodium chloride, 2.7 mM potassium chloride, and 2 mM DTT.

**I.12 NADH/NAD$^+$ Activity-Based Assay.** NAD$^+$/NADH was measured using the NADH/NAD-Glo Assay from Promega according to the manufacturer's recommendations. Briefly, the enzyme/histones substrate mix prepared in the reaction buffer was added to the reaction plate wells (Corning 3572, Non-Treated). Compounds in 100% DMSO were added into the enzyme mixture by acoustic technology (Echo550; nanoliter range). 100% DMSO was delivered to no compound/DMSO control wells. The reaction plate was centrifuged following a pre-incubation of 20 min at room temperature. The reaction was initiated by adding NAD$^+$ followed by an incubation of 2 hours at room temperature. NAD/NADH-Glo™ prepared as per the kit instructions were added to all assay wells. After incubation of 30 minutes at room temperature in the dark, luminescence with an endpoint read at 30 minutes was recorded. Data were compared with DMSO-treated cells and expressed as % control. Data obtained is the result of three independent experiments (n=3).

##### NanoBRET Target Engagement (Intracellular PARP1)

NanoBRET® Target Engagement assays were performed essentially following the Promega protocol with minor modifications. HEK293 cells were transiently transfected with a PARP1–NanoLuc® fusion construct using FuGENE® HD (Promega). Test compounds were pre-spotted into 384-well assay plates using an Echo 550 acoustic dispenser (Labcyte). Transfected cells were harvested, mixed with the cell-permeable NanoBRET TE PARP Tracer-01, and dispensed into the plates. Plates were incubated for 1 h at 37°C in 5% $CO_2$. NanoBRET® Nano-Glo® Substrate plus Extracellular NanoLuc® Inhibitor solution was then added, followed by a 20 min incubation at room temperature. Donor (460 nm) and acceptor (600 nm) emissions were recorded on an EnVision 2104 multimode plate reader (PerkinElmer). The NanoBRET ratio was computed as:

$$\text{BRET Ratio} = \frac{\text{Acceptor}_{\text{sample}}}{\text{Donor}_{\text{sample}}} - \frac{\text{Acceptor}_{\text{no tracer}}}{\text{Donor}_{\text{no tracer}}}.$$

Concentration–response curves were fitted in GraphPad Prism using a four-parameter logistic model to obtain apparent $IC_{50}$ values. Apparent $K_i$ values were then obtained by competitive tracer-displacement analysis, correcting each $IC_{50}$ for the tracer concentration and its apparent affinity for PARP1 (i.e., $K_i \approx \text{IC}_{50}/[1+[\text{tracer}]/K_{D,\text{app}}]$). Assay conditions were set to support accurate affinity estimation by keeping tracer occupancy modest, controlling expression levels, minimizing ligand depletion and nonspecific binding, and fitting a single-site competition model.

**I.13 NMR Spectroscopy.** The NMR titration samples were prepared by adding the unlabeled inhibitors (iParp1, iParp2, iParp3 and iParp4) into solutions of U-[$^{15}$N, D]-labeled PARP1 to reach a ratio of 1:2 (50 µM protein and 100 µM olaparib, iParp1 or iParp3) or 1:1 for inhibitors with limited water solubility (25 µM protein, and 25 µM iParp2 or iParp4). 500 µL of each sample was transferred into a 5 mm NMR tube and 2D-$^{15}$N-TROSY spectra were acquired at 298 K on a Bruker Avance III HD spectrometer equipped with a TCI cryoprobe and z-shielded gradients, operating at a $^1$H frequency of 800 MHz. Data were processed using NmrPipe (165) and analyzed with CCPNmr (166). To transfer PARP1 CAT-domain assignment, HNCA and HNCACB spectra were acquired and the obtained frequencies were compared to BMRB entry 50454. Chemical shift perturbations and peak broadening were extracted using CCPNmr and their combined normalized value was plotted on the PARP1 docking poses.

**I.14 Crystallography.** PARP1 was co-crystallized with iParp1, the most potent of our inhibitors. The details are described below.

***Protein Expression and Purification.*** The human PARP1 construct (residues 662-1011) contains an N-terminal GST tag and 3C cleavage site, was overexpressed in *E. coli* BL21 (DE3), and purified using affinity chromatography and size-exclusion chromatography. Briefly, cells were grown at 37°C in TB medium in the presence of 100 µg/mL of ampicillin to an OD of 0.8, cooled to 17°C, induced with 400 µM isopropyl-1-thio-D-galactopyranoside (IPTG), incubated overnight for 20 hrs at 17°C, collected by centrifugation, and stored at -80°C. Cell pellets were lysed in buffer A (25 mM HEPES, pH 7.5, 200 mM NaCl, and 1 mM TCEP) using a Microfluidizer (Microfluidics), and the resulting lysate was centrifuged at 30,000g for 40 min. Glutathione beads (Cytiva) were mixed with cleared lysate for 90 min and washed with buffer A. Beads were transferred to an FPLC-compatible column, and the bound protein was washed further with buffer A supplemented with 1 M NaCl for 10 column volumes, followed by buffer A for 10 column volumes. The GST-tag was cleaved on column by adding 3C protease to the washed beads and incubating overnight in the cold room. The cleaved PARP1 was eluted from the column, then concentrated and purified further using a Superdex 75 16/600 column (Cytiva) in buffer A. The fractions containing cleaved PARP1 were concentrated to 6.5 mg/mL and stored at -80°C.

***Crystallization.*** A solution of 6.5 mg/mL human PARP1 was crystallized in 30% PEG-4000, 200 mM Lithium sulfate, and 100 mM Tris, pH 8.5 by sitting-drop vapor diffusion at 20°C. Crystals were transferred briefly into crystallization buffer containing 25% glycerol prior to flash-freezing in liquid nitrogen.

***Structure Determination.*** Diffraction data were collected at beamline 17-ID-2 of the National Synchrotron Light Source II (Brookhaven National Laboratory). Datasets were integrated and scaled using XDS (88). Structures were solved by molecular replacement using the program Phaser (167) and the search model (PDB code 7KK2). Iterative manual model building and refinement using Phenix (168) and Coot (169) led to a model with excellent statistics summarized in Supplementary Table 14.

| Parameter | Value |
|---|---|
| PDB Code | 8VYH |
| Resolution range | 46.28 - 2.05 (2.09 - 2.05) |
| Space group | P 21 21 21 |
| Unit cell | 48.92 93.14 165.55 90 90 90 |
| Total reflections | 189094 (12142) |
| Unique reflections | 45179 (2880) |
| Multiplicity | 4.2 (4.2) |
| Completeness (%) | 94.54 (95.96) |
| Mean I/sigma(I) | 4.12 (70.0) |
| Wilson B-factor | 30.65 |
| R-merge | 0.2088 (1.529) |
| R-pim | 0.1064 (0.773) |
| CC1/2 | 0.987 (0.323) |
| CC* | 0.997 (0.699) |
| Reflections used in refinement | 45134 (2853) |
| Reflections used for R-free | 2275 (147) |
| R-work | 0.1971 (0.3110) |
| R-free | 0.2504 (0.3614) |
| CC(work) | |
| CC(free) | |
| Number of non-hydrogen atoms | 6123 |
| macromolecules | 5506 |
| ligands | 90 |
| solvent | 527 |
| Protein residues | 700 |
| Nucleic acid bases | |
| RMS(bonds) | 0.003 |
| RMS(angles) | 0.55 |
| Ramachandran favored (%) | 99.14 |
| Ramachandran allowed (%) | 0.86 |
| Ramachandran outliers (%) | 0.00 |
| Rotamer outliers (%) | 0.16 |
| Clashscore | 12.26 |
| Average B-factor | |
| macromolecules | 35.77 |
| ligands | 35.49 |
| solvent | 41.46 |
| Number of TLS groups | |

**Supplementary Table 14.** Crystallographic Data and Refinement Statistics for PARP1-iParp1 complex structure.

**I.15 Colony Formation Assays.** MDA-MB-436 cells (1000/well) were seeded in 6-well plates, treated with graded concentrations of the indicated compounds the next day, and grown in the presence of drugs for 15 days. Colonies were fixed with ice-cold methanol for 10 minutes and then stained with 1% crystal violet for 30 minutes. Plates were imaged with GelCount (Oxford Optronix), and quantification of the colony numbers was performed with ImageJ.

#### J Computational Overhead

AdaptiveFlow employs a hierarchical orchestration approach specifically designed to minimize computational overheads at the scale required for large-scale molecular docking operations. At the fundamental level of this system, individual ligand docking operations require between several seconds to several minutes of computation time on a single CPU core, with this duration varying considerably based on the complexity of the molecular structures involved and the specific parameters employed in the docking calculation. When executing distributed docking operations at scale, one will encounter multiple sources of overhead that, while individually small, can compound to create significant inefficiencies. These overhead sources include the workload submission scheduling through systems such as Slurm or AWS Batch, job startup communication protocols, the transfer of input data files containing ligand information, runtime initialization overhead particularly from Java Virtual Machine initialization in Java-based docking programs, the storage and upload of computational results, and finally the job completion overhead associated with scheduler communication. To illustrate the significance of these overheads, consider a conservative scenario where each non-docking operation requires merely 0.1 seconds of computation time, an estimate that underestimates actual overhead in many real-world cases. For a docking operation requiring 10 seconds of computation, these six overhead sources would collectively contribute approximately 0.6 seconds, representing a 6% performance penalty per ligand. When this analysis is extrapolated to operations involving one million ligands, the cumulative overhead becomes substantial and would compound significantly without appropriate mitigation strategies.

**J.1 Hierarchical Strategy.** AdaptiveFlow addresses this scaling challenge through a three-layer hierarchical architecture that systematically amortizes overhead costs across multiple docking operations.

***Data Layer.*** At the foundation of this hierarchy lies the data layer, where ligands are assembled into collections of approximately 1,000 molecules each. These collections are subsequently compressed into archive files, a strategy that serves dual purposes: it substantially reduces the file and object count that must be managed by the storage system, and it amortizes input/output costs across the many docking operations that will be performed on ligands within each collection.

***Compute Layer Architecture.*** The compute layer operates through independent computational tasks termed "subjobs," each of which processes one or more ligand collections. These subjobs are typically dimensioned to execute for 45 to 90 minutes of runtime on computational nodes equipped with 16 virtual CPUs, corresponding to 8 physical processor cores (this sizing is configurable). This particular sizing rationale reflects optimization for cloud-based spot instances and preemptible instances available on high-performance computing platforms, where this configuration typically yields cost savings of 50-60% while maintaining compatibility with HPC scheduler backfill efficiency. Importantly, these parameters remain configurable to accommodate the specific requirements and constraints of different computational environments. Each subjob executes a custom Python program, referred to as the AdaptiveFlow Runner, which leverages standard Python libraries. Notably, the implementation employs Python's multiprocessing package to circumvent the Python Global Interpreter Lock (GIL), thereby enabling true parallel execution. The Runner implements a processing pipeline comprising several connected stages: the initial download or copy of ligand collections from storage, decompression of these archives, execution of docking calculations for each individual ligand, aggregation and summarization of computational results, and finally upload of compressed result files back to persistent storage. For docking programs that use Java, each Runner will share a single JVM through Nailgun, removing the traditional JVM startup time from each docking process. This queue-based architecture incorporating dynamic worker threads provides several critical advantages. First, it enables overlapping of input/output operations with computational work, allowing the next collection to download while docking calculations proceed on the current dataset. Second, the architecture supports configurable oversubscription, wherein more docking threads than physical cores can be instantiated, which has empirically demonstrated throughput improvements as the overhead itself is reduced rather than increased by this approach. Third, the system performs batch compression and storage of results, further amortizing input/output costs across multiple ligands.

***Scheduler Layer.*** At the highest level of the hierarchy, the scheduler layer addresses the rate limits imposed by HPC schedulers on job submission. AdaptiveFlow introduces the concept of "workunits," configurable bundles of subjobs that map to scheduler array jobs. Rather than submitting thousands of individual jobs to the scheduler, which would incur substantial overhead, the system submits a single array job containing thousands of array elements. For instance, instead of submitting 1,000 separate job requests, AdaptiveFlow submits one array job with 1,000 constituent elements, thereby dramatically reducing scheduler overhead and the associated job management costs. AdaptiveFlow can submit to either Slurm or AWS Batch using their respective syntax for array jobs. While generic orchestration frameworks exist within the scientific computing ecosystem, AdaptiveFlow implements a purpose-built model designed specifically for the molecular docking workload pattern. This design decision serves three primary objectives: minimizing software dependencies and deployment complexity, optimizing specifically for the characteristic patterns of docking workflows, and maintaining portability across both traditional HPC environments and cloud computing platforms while optimizing for the specific characteristics of each. The effectiveness of this hierarchical approach is demonstrated through empirical analysis of the system's scaling behavior. Linear scaling observed in deployments utilizing up to 5.6 million virtual CPUs provides compelling evidence that orchestration overhead remains negligible across this range. If Python-based task management imposed significant overhead on the system, two characteristic failure patterns would manifest:

first, sublinear scaling as communication costs begin to dominate computational work, and second, performance plateaus at high core counts where task management overhead saturates available resources. Instead, the sustained linear scaling behavior demonstrates that computational time remains dominated by the actual docking calculations, which require seconds to minutes per ligand, rather than by task management operations, which require only milliseconds. This scaling characteristic provides strong empirical evidence that the hierarchical orchestration approach successfully achieves its design objective of minimizing overhead at scale. To directly validate that orchestration overhead is negligible at the single-run level (in addition to the large-scale scaling results), we benchmarked the same set of 579 ligands using Quick Vina 2 in two modes: (i) a pure serial bash loop that executes one docking at a time, and (ii) execution through the AFVS workunit runner under identical docking parameters ($exhaustiveness = 1$, $num_modes = 1$, $cpu = 1$, $seed = 42$). As summarized in Supplementary Table 15, the per-ligand docking times are on the order of a few seconds in both cases, while the AFVS Python overhead remains at the millisecond level ($\sim$5 ms per docking on average), contributing a negligible fraction of the total per-ligand runtime. Moreover, AFVS reduces the end-to-end elapsed time (wall-clock span) by executing many independent dockings concurrently, without altering docking outcomes, reinforcing that overall runtime is dominated by docking computation rather than workflow management.

**J.2 File Usage Analysis for One Million Molecules.** The hierarchical collection-based architecture described in the previous response was designed not only to minimize computational overhead but also to explicitly address file/inode limitations the reviewer correctly notes that are characteristic of traditional HPC filesystems. We also note that inode constraints are specifically relevant for classical POSIX filesystems; when deploying AdaptiveFlow on cloud infrastructure utilizing AWS S3, these limitations do not apply as object storage architectures do not impose inode quotas. For deployments on traditional HPC filesystems, the collection strategy provides substantial mitigation of inode consumption. Rather than generating individual files for each ligand molecule, an approach that would rapidly exhaust available inodes during large-scale screening campaigns, AdaptiveFlow organizes both input data and output results into aggregated file structures. The input data structure consists of approximately 1,000 compressed collection files, where each collection contains roughly 1,000 ligand molecules. We can also assume one workunit input configuration file (assume 1 collection per subjob, 1000 subjobs to a workunit). On the output side, results are aggregated at the subjob level by the Runner rather than at the individual ligand level, generating approximately 1,000 result files (in Parquet or CSV format containing structured docking results) plus approximately 1,000 log archives for diagnostic purposes. This architecture yields a total file count of approximately 3,001 files per million molecules. The collection and subjob-based strategy serves multiple complementary objectives within the system architecture: it amortizes I/O costs across multiple docking operations as discussed previously, reduces network transfer overhead through compressed archives, and maintains compatibility with HPC inode quotas that would otherwise constrain large-scale screening operations. For cloud-based deployments, AdaptiveFlow provides native support for S3-compatible object storage. While the inode limitations are not there, the same aggregated approach is still used to reduce the overheads and costs that would be associated with per-docking objects.

***Cloud Scalability Benchmark.*** The benchmark was defined as a real-world stress test involving the preparation and docking of the 69 billion compound Enamine REAL Space library. The primary metric was scalability efficiency, defined as the ability to maintain constant processing throughput per core as the total number of concurrent vCPUs increased.

***Characterization.*** The 5.6 million vCPU run was characterized by the simultaneous utilization of Spot Instances (Intel Xeon Scalable processors) across multiple AWS regions, orchestrated via AWS Batch. This scale refers to concurrent virtual CPUs executing the workflow in parallel.

***Overhead Management.*** Overhead was minimized via a hierarchical batching architecture. Ligands were grouped into "collections" ($\sim$1,000 molecules) and further packed into "subjobs" (atomic units of work processing multiple collections). This ensured that the compute duration for each task (approx. 45–90 minutes) significantly exceeded the millisecond-scale scheduling overhead.

***Scaling Behavior.*** The system demonstrated near-linear weak scaling, where throughput (molecules processed per minute) increased linearly with the number of vCPUs up to the 5.6 million limit. This effectively translates to linear strong scaling for the fixed, ultra-large workload (69 billion compounds), enabling the total run time to be compressed from weeks to hours.

We are not aware of any other publicly known examples that show scaling to this level (5.6 million concurrent vCPUs). e.g. other publicly known large scale benchmarks include Clemson public safety video processing at 2.1M vCPUs and 3.1M vCPUs on AWS using Yellowdog.

| Timing metric | QVina02 via AFVS (s) | QVina02 serial (s) |
|---|---|---|
| Records analyzed (count) | 579 | 579 |
| Total docking time | 2311.86 | 2366.44 |
| Average docking time | 3.99 | 4.09 |
| Wall-clock span (min–max) | 145.86 | 2394.00 |
| Total Python overhead | 3.11 | 0.00 |
| Average Python/bash overhead | 0.01 | 0.00 |
| Total time incl. overhead | 2314.96 | 2366.44 |
| Average time incl. overhead | 4.00 | 4.09 |

**Supplementary Table 15. Benchmark timing comparison of Quick Vina 2 docking via AFVS vs. a serial Quick Vina 2 loop.** Per-ligand docking durations were summed to compute total docking time, and mean values report average per-ligand time. The wall-clock span corresponds to the elapsed time between the earliest docking start and the latest docking end. For the serial run, Python/bash overhead is set to zero because AFVS orchestration is not used. Quick Vina 2 settings: $exhaustiveness = 1$, $num_m odes = 1$, $cpu = 1$, $seed = 42$.

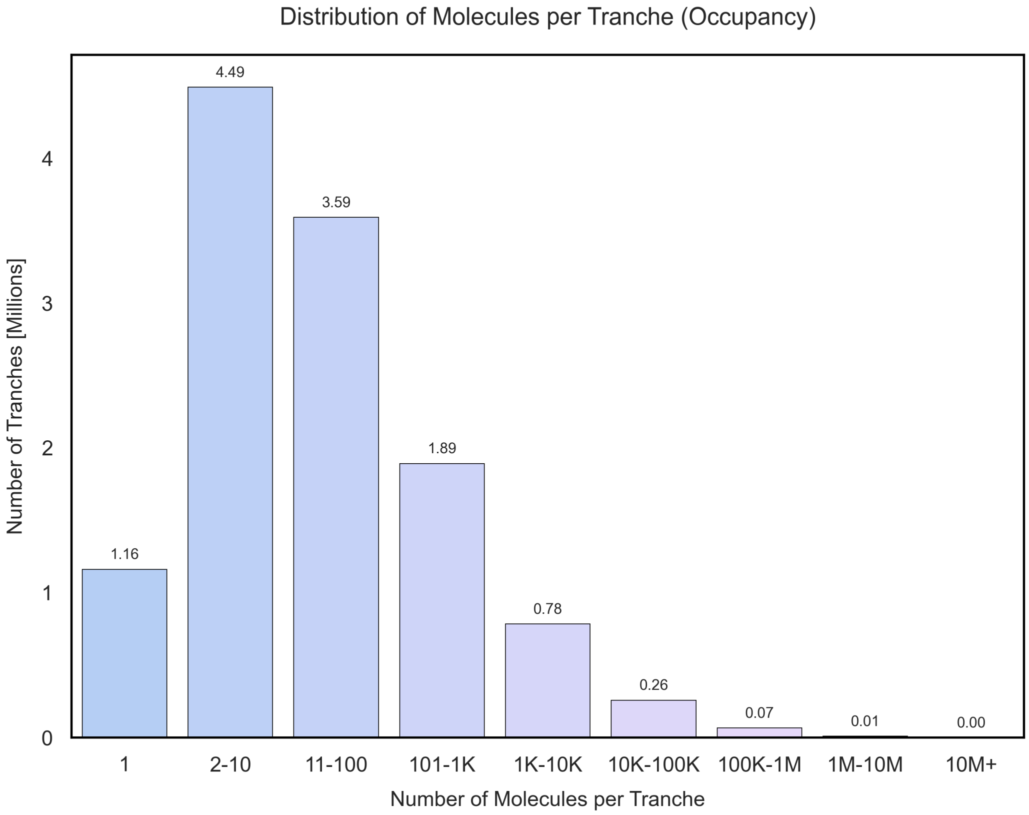

**Supplementary Fig. 34. Distribution of Compound Occupancy Across Tranches in the Enamine REAL Space.** Histogram displaying the number of tranches (y-axis) populated by a given number of molecules (x-axis) within the 18-dimensional property grid. The distribution spans from single-molecule tranches to highly populated tranches containing over 10 million compounds.

#### K Configuration File of AFLP

All options of AFLP are specified in a central configuration file. A sample configuration file is shown in Supplementary Listing 1.

**Supplementary Listing 1.** AFLP Example Configuration File.

```
# Each line that starts with ’#’ is a comment/description of the variable above the comment
# Each line that starts with ’*’ belongs to a section heading

****** Job Resource Configuration

job_name=testing
# alphabetic characters (i.e. letters from a-z or A-Z)
# Used to describe distinct runs (using the same name will
# overwrite data if using S3!)

threads_to_use=384
# This sets how many processes the main execution loop should be using
# to process. This is generally 2x the number of vCPUs or hyperthreads
# available on the system it is being run on

************************************************
** Batch system configuration
************************************************

batchsystem=slurm
# Possible values: awsbatch, slurm

****** AWS Batch Options (if batchsystem=awsbatch)

### To use AWS Batch you must first complete the steps outlined
### in the user guide for AWS Batch

aws_batch_prefix=af
# Prefix for the name of the AWS Batch queues. This is normally ’af’
# if you used the CloudFormation template

aws_batch_number_of_queues=2
# Should be set to the number of queues that are setup for AWS Batch.
# Generally this number is 2 unless you have a large-scale (100K+ vCPUs)
# setup

aws_batch_jobdef=af-jobdef-AFLP
# Generally this is [aws_batch_prefix]-jobdef-AFLP
# (e.g. if aws_batch_prefix=af, then aws_batch_jobdef=af-jobdef-AFLP

aws_batch_array_job_size=2
# Target for the number of jobs that should be in a single array job for AWS Batch.

aws_ecr_repository_name=af-AFLP-ecr
# Set it to the name of the Elastic Container Registry (ECR)
# repository (e.g. af-AFLP-ecr) in your AWS account
# (If you used the template it is generally af-AFLP-ecr)

aws_region=us-east-1
# Set to the AWS location code where you are running AWS Batch
# (e.g. us-east-1 for North America, Northern Virginia)

aws_batch_subjob_vcpus=8
# Set to the number of vCPUs that should be launched per subjob.
# ’threads_to_use’ above should be >= to this value.

aws_batch_subjob_memory=15000
# Memory per subjob to setup for the container in MB

aws_batch_subjob_timeout=10800
# Maximum amount of time (in seconds) that a single AWS Batch job should
# ever run before being terminated.

****** Slurm Options (if batchsystem=slurm)

slurm_template=./templates/template1.slurm.sh
# Template for the slurm job
# Additional slurm attributes can be added directly to this
# template file if they are not available as pass throughs from
# AFVS

slurm_array_job_throttle=1
# Maximum number of jobs running within a single slurm array job

slurm_partition=standard96
# Partition to submit the job
```

```
slurm_cpus=192
# Number of CPUs that are being used

slurm_array_job_size=1
# Maximum number of concurrent jobs from a single array job
# that should be run

************************************************
** Storage configuration
************************************************

job_storage_mode=sharedfs
# This mode determines where data is retrieved and stored from as part of
# AFVS. Valid modes:
#   * s3: Job data is stored on S3 object store, which is the required
#         mode if using AWS Batch. Items under the "S3 Object Store"
#         heading in the configuration are required if this mode is used
#   * sharedfs: This mode requires that all running jobs have access to the
#         same shared filesystem that will allow for both input and output
#         of data. This is required if using Slurm

job_storage_output_addressing=standard
# If input is placed with the hash addressing mode, then use 'hash'.
# otherwise use "standard" for the classic addressing mode
# 'hash' will place data in:
#     [0-f][0-f]/[0-f][0-f]/complete/<output_type>/<metatranche>/<tranche>/<collection>
# 'standard' will place data in:
#     complete/<output_type>/<metatranche>/<tranche>/<collection>

****** Object Store Settings (S3)

object_store_input_ligand_library_bucket=adaptiveflow-data
# Bucket name for the collection data
# Trailing slashes should be used

object_store_input_ligand_library_prefix=AFLP/data/Enamine_2021_test
# Prefix used within the object store to address the collections
# Trailing slashes should be used

object_store_job_output_data_bucket=adaptiveflow-data
# Bucket name for the job data
# Trailing slashes should be used

object_store_job_output_data_prefix=jobs/AFLP
# Where to place job-specific data. This includes where AdaptiveFlow will place
# the input data needed for jobs as well as the output files. Data be be placed
# in object_store_job_output_data_prefix/<job_letter>
# Trailing slashes should be used

****** Shared Filesystem Settings

collection_folder=../input-files/collections
# Path to where the collection file are stored
#  * This is used when job_storage_mode=sharedfs or
#    when the uploader helper script is being used
#
# Slash at the end is not required (optional)
# Either pathname is required w.r.t. the folder tools/
#     or absolute path (e.g. /home/afuser/collections)

# Output of processed ligands will be placed in
# ../workflow/complete (where ../workflow is relative to the tools
# directory of the AFVS installation

****** Workflow Options

ligands_todo_per_queue=1000
# Used as a limit of ligands for the to-do lists.
# A reasonable number for this is generally 1000. The length of time
#     to process will depend on the docking scenarios run

tempdir_default=/dev/shm
# The directory which is used for the temporary workflow files which need a normal performance
# Is normally a local SSD or HDD. A temporary directory will be created underneath this dir

tempdir_fast=/dev/shm
# The directory which is used for the temporary workflow files which need a fast perfomance
# Should be a a local ram filesystem/ramdisk. A temporary directory will be created underneath this dir

store_all_intermediate_logs=false
# This determines if the intermediate files (all intermediate steps) are
# retained for review after a run. This is not recommended unless you
# are debugging as it increases output by over 2x
# Valid values:
```

```
#   * true: Keep the logs. They will be placed in intermediate/
#   * false: Do not retain the logs

***********************************************
** Ligand Preparation Options
***********************************************

****** Desalting

desalting=true
# If true, extracts the largest organic part of the molecule
# Possible values:
#   * false
#   * true

desalting_obligatory=false
# Setting only required if desalting=true
# Possible values:
#   * true: Successful desalting of the ligand is mandatory (unsuccessful desalting leads to the omission of this ligand ).
#   * false: The ligand will continue to be processed even if the desalting step fails.

****** Neutralization

neutralization=true
# Neutralizes molecules using JChem's Standardizer of ChemAxon
# Possible values:
#   * false
#   * true

neutralization_program_1=obabel
neutralization_program_2=none
# Program 1 is used at first for each ligand, and if it fails program 2 is used instead. If the second program also fails, then the ligand is skipped.
# Setting only required if neutralization=true
# Possible values:
#   * obabel
#   * standardizer
#   * none (only for neutralization_program_2)

chemaxon_neutralization_timeout=15
# Time in seconds to wait to timeout for neutralization with chemaxon

obabel_neutralization_timeout=15
# Time in seconds to wait to timeout for neutralization with obabel

neutralization_mode=only_genuine_desalting_and_if_charged
###neutralization_mode=after_desalting_if_charged
# Only relevant if neutralization=true
# Possible values:
#   * always: always neutralize the molecule
#   * only_genuine_desalting: only neutralize the molecule if the input structure
#     contained more than one component
#   * only_genuine_desalting_and_if_charged: only neutralize if the input structure
#     contained more than one component and if the smallest component contained an ion

neutralization_obligatory=false
# Setting only required if neutralization=true
# Possible values:
#   * true: Successful neutralization of the ligand is mandatory (unsuccessful neutralization leads to the omission of this ligand).
#   * false: The ligand will continue to be processed even if the neutralization step fails.
# Settable via range control files: Yes

****** Stereoisomer Generation

stereoisomer_generation=false
# Possible values:
#   * false
#   * true

stereoisomer_generation_program_1=rdkit
stereoisomer_generation_program_2=none
# Program 1 is used at first for each ligand, and if it fails program 2 is used instead. If the second program also fails, then the ligand is skipped.
# Setting only required if stereoisomer_generation=true
# Possible values:
#   * rdkit
#   * cxcalc

stereoisomer_obligatory=false
# Setting only required if stereoisomer_generation=true
# Possible values:
```

```
#   * true: Successful stereoisomer generation of the ligand is mandatory
#           (unsuccessful stereoisomer generation leads to the omission of this ligand).
#   * false: The ligand will continue to be processed even if the stereoisomer generation
#            step fails.

cxcalc_stereoisomer_generation_options=-T -f smiles
# any options which should be passed to the stereoisomer plugin of cxcalc
# This setting has to contain the "-f smiles" option
# The "-T" option for instance specifies that that chiral carbons should be preserved.
# Documentation on the options is available here:
#     https://docs.chemaxon.com/display/docs/cxcalc-calculator-functions.md#src-1806682-cxcalccalculatorfunctions-stereoisomers

rdkit_stereoisomer_generation_unique_correction=false
# Enumeration of stereoisomers by RDkit can result in redundant stereoisomers
# This can be resolved by setting this option to true. However, can be time consuming

cxcalc_stereoisomer_timeout=15
# Time in seconds to wait to timeout for cxcalc

obabel_stereoisomer_timeout=30
# Time in seconds to wait to timeout for obabel

****** Tautomerization

tautomerization=true
# Possible values:
#   * false
#   * true

tautomerization_program_1=obabel
tautomerization_program_2=none
# Program 1 is used at first for each ligand, and if it fails program 2 is used instead. If the second program also fails, then the ligand is skipped.
# Setting only required if tautomerization=true
# Possible values:
#   * obabel
#   * cxcalc
#   * none (only for tautomerization_program_2)

tautomerization_obligatory=false
# Setting only required if tautomerization=true
# Possible values:
#   * true: Successful tautomerization of the ligand is mandatory (unsuccessful tautomerization leads to the omission of this ligand).
#   * false: The ligand will continue to be processed even if the tautomerization step fails.

cxcalc_tautomerization_options=
# any options which should be passed to the tautomerization plugin of cxcalc

cxcalc_tautomerization_timeout=15
# Time in seconds to wait to timeout for tautomerization with cxcalc

obabel_tautomerization_timeout=30
# Time in seconds to wait to timeout for tautomerization with obabel

****** Protonation State Generation

protonation_state_generation=true
# Possible values:
#   * false
#   * true

protonation_program_1=obabel
protonation_program_2=none
# Program 1 is used at first for each ligand, and if it fails program 2 is used instead. If the second program also fails, then the ligand is skipped.
# Setting only required if protonation_state_generation=true
# Possible values:
#   * obabel
#   * cxcalc
#   * none (only for protonation_program_2)

cxcalc_protonation_timeout=15
# Time in seconds to wait to timeout

obabel_protonation_timeout=15
# Time in seconds to wait to timeout

protonation_obligatory=false
# Setting only required if protonation_state_generation=true
# Possible values:
#   * true: Successful protonation of the ligand is mandatory (unsuccessful
#     protonation leads to the omission of this ligand).
#   * false: The ligand will continue to be processed even if the protonation step fails.
#     This might light to protonation states which are unphysiological.
```

```
protonation_pH_value=7.4
# Setting only required if protonation_state_generation=true
# Possible values: floating point number between 0.0 and 14.0

****** Conformation Generation

conformation_generation=true
# Generation of 3D conformation/coordinates of the ligand
# Possible values:
#   * false
#   * true

conformation_program_1=obabel
conformation_program_2=none
# Setting only required if conformation_generation=true
# Program 1 is used at first for each ligand, and if it fails program 2 is used instead.
# If the second program also fails, then the ligand is skipped.
#
# Possible values:
#   * obabel
#   * molconvert
#   * none (only possible for conformation_program_2)

molconvert_3D_options=-3:{fine}
# Setting only required if conformation_generation=true and one of the programs used is molconvert
# 3D conformation generation options for molconvert.
# See also the help text printed by molconvert for additional information
# Possible values:
#   * -3           Defaults to value 3{fast}
#   *  3{fine}     Find low energy conformer Leave failed fragments intact
#   *  3{fast}     Fast clean, if failed, perform fine clean, accept any generated structure (default)
#   *  3{nofaulty} Same as S{fast}, but leave failed fragments intact.

molconvert_conformation_timeout=15
# Time in seconds to wait to timeout

obabel_conformation_timeout=15
# Time in seconds to wait to timeout

conformation_obligatory=true
# Setting only required if conformation_generation=true
# Possible values:
#   * true: Successful 3D conformation generation of the ligand is mandatory
#     (unsuccessful conformation generation leads to the omission of this ligand).
#   * false: The ligand will continue to be processed even if the conformation
#      generation step fails.

****** Target Format Generation

targetformats=pdb:pdbqt:sdf:mol2:smi:selfies
# Possible values:
#     * selfies (https://github.com/aspuru-guzik-group/selfies)
#     * Any format supported by the Open Babel, using the file format identifiers used by Open Babel.
#       A complete list can be obtained by running the command "obabel -L formats"
# Multiple target formats can be specified by separating them with colons, e.g. pdb:sdf:pdbqt

****** Open Babel

obabel_memory_limit=1000000
# In KB
# Recommended value: >= 500000

****** Energy Check

energy_check=true
# Determines whether the potential energy is checked by obenergy (Open Babel Enegy). This can be useuful to filter out
     compounds with unrealistic predicted 3D geometry
# Possible values:
#   * true
#   * false

energy_max=10000
# Maximum allowed energy value. Recommended: 10000
# Possible values: Positive integer

****** Tranche Assignments

tranche_assignments=true
# Should each ligand be assigned a new tranche based based on molecular properties of the ligand?
# This is useful in particular if the tranches of the input ligands are pseudo-tranches (i.e. are arbitrary or have no
```

```
    meaning).

tranche_types=hbd_obabel
# These variables are only needed if tranche_assignments=true
# Multiple values are separated by colons
# Possible values:
#   * mw_jchem: molecular weight
#   * mw_obabel: molecular weight
#   * logp_jchem: octanol water partition coefficient by JChem's cxcalc
#   * logp_obabel: octanol water partition coefficient by Open Babel
#   * hba_jchem: hydrogen bond acceptor count
#   * hba_obabel: hydrogen bond acceptor count
#   * hbd_jchem: hydrogen bond donor count
#   * hbd_obabel: hydrogen bond donor count
#   * rotb_jchem: rotatable bond count
#   * tpsa_jchem: topological polar surface area by JChem's cxcalc
#   * tpsa_obabel: topological polar surface area by Open Babel
#   * logd: octanol water partition coefficient
#   * logs: aqueous solubility in mol/L
#   * atomcount_jchem: number of atoms (including hydrogen)
#   * atomcount_obabel: number of atoms (including hydrogen)
#   * bondcount_jchem: numner of bonds (including hydrogen-heavy atom bonds)
#   * bondcount_obabel: numner of bonds (including hydrogen-heavy atom bonds)
#   * ringcount: number of non-aromatic rings
#   * aromaticringcount: number of aromatic rings
#   * mr_jchem: molecular refractivity by JChem's cxcalc
#   * mr_obabel: molecular refractivity by Open Babel
#   * formalcharge: total (formal) charge of the molecule
#   * positivechargecount: number of positive charged functional groups/atoms
#   * negativechargecount: number of negatively charged functional groups/atoms
#   * fsp3: fraction of sp3 hybridized carbon atoms
#   * chiralcentercount: number of chiral centers
#   * halogencount: number of halogen atoms (F, Br, Cl, I)
#   * sulfurcount: number of sulfur atoms (S)
#   * NOcount: number of oxygen and nitrogen atoms (N, O)
#   * electronegativeatomcount: number of electronegative atoms (N, O, S, P, F, Br, Cl, I)
#   * mw_file: molecular weight
#   * logp_file: octanol water partition coefficient by JChem's cxcalc
#   * hba_file: hydrogen bond acceptor count
#   * hbd_file: hydrogen bond donor count
#   * rotb_file: rotatable bond count
#   * tpsa_file: topological polar surface area by JChem's cxcalc
#   * logd_file: octanol water partition coefficient
#   * logs_file: aqueous solubility in mol/L
#   * heavyatomcount_file: number of atoms (including hydrogen)
#   * bondcount_file: numner of bonds (including hydrogen-heavy atom bonds)
#   * ringcount_file: number of non-aromatic rings
#   * aromaticringcount_file: number of aromatic rings
#   * mr_file: molecular refractivity by Open Babel
#   * formalcharge_file: total (formal) charge of the molecule
#   * positivechargecount_file: number of positive charged functional groups/atoms
#   * negativechargecount_file: number of negatively charged functional groups/atoms
#   * fsp3_file: fraction of sp3 hybridized carbon atoms
#   * chiralcentercount_file: number of chiral centers
#   * halogencount_file: number of halogen atoms (F, Br, Cl, I)
#   * sulfurcount_file: number of sulfur atoms (S)
#   * NOcount_file: number of oxygen and nitrogen atoms (N, O)
#   * electronegativeatomcount_file: number of electronegative atoms (N, O, S, P, F, Br, Cl, I)
#   * enamine_type: Enamine type category
#   * doublebondstereoisomercount_jchem: number of double-bond steroisomers from JChem
#   * aromaticproportion_jchem: aromatic proportion generated from JChem
#   * qed_rdkit: QED generated by RDKit

file_fieldnames=smi:ligand-name:hbd_obabel
# For each tranche type which reads the value from a file, those values must be defined.
# At a minimum 'smi' and 'ligand-name' are REQUIRED (and are generally expected to be the first
# two items per line
#
# e.g. file_fieldnames=smi:ligand-name:-:-:-:-:-:mw:atomcount
#      This means that the first column should be tghe SMILES string, the second
#      column is the 'friendly name', columns 3-7 are values not used,
#      column 8 is the value that should be used for mw_file in tranche assignment,
#      column 9 is the value that should be used for atomcount_file in tranche assignment
#      Other valid options include: enamine (for enamine_type), mw, heavyatomcount,
#         logp, hba, hbd, rotb, fsp3, tpsa

# For each tranche_type X in the variable tranche_types, one additional variable "tranche_<X>_partition" has to be
    specified.
# This variable has to be a set of N >= 1 values (at least one), separated by colons, which partitions the value range
    of the molecular property of tranche <X> into N+1 intervals.
# For example, the partitions '-0:1:2' would result in the 4 intervals: (-∞,0], (0,1], (1,2], (2,∞). The maximum
    number of allowed intervals is 27 (corresponding to 26 values of the alphabet). Each value has to be an integer or
    float, and the values have to be in ascending order.
# These variables are only needed if tranche_assignments=true
```

```
tranche_mw_jchem_partition=200:300:400:450
tranche_mw_obabel_partition=200:300:400:450
tranche_mw_file_partition=200:300:400:450
tranche_logp_jchem_partition=0:2:3:4:5
tranche_logp_obabel_partition=0:2:3:4:5
tranche_logp_file_partition=0:2:3:4:5
tranche_hba_jchem_partition=2:4:7:10
tranche_hba_obabel_partition=2:4:7:10
tranche_hba_file_partition=2:4:7:10
tranche_hbd_jchem_partition=0:2:4:5
tranche_hbd_obabel_partition=0:2:4:5
tranche_hbd_file_partition=0:2:4:5
tranche_rotb_jchem_partition=0:2:4:7:10
tranche_rotb_obabel_partition=0:2:4:7:10
tranche_tpsa_jchem_partition=40:80:110:140
tranche_tpsa_obabel_partition=40:80:110:140
tranche_tpsa_file_partition=40:80:110:140
tranche_logd_partition=0:2:3:4:5
tranche_logd_file_partition=-1:0:1:2:2.5:3:3.5:4:4.5:5
tranche_logs_partition=-6:-5:-4:-3
tranche_logs_file_partition=-6:-5:-4:-3
tranche_atomcount_jchem_partition=10:20:30:40:50
tranche_atomcount_obabel_partition=10:20:30:40:50
tranche_heavyatomcount_file_partition=10:20:30:40:50
tranche_bondcount_jchem_partition=10:20:30:40:50
tranche_bondcount_obabel_partition=10:20:30:40:50
tranche_bondcount_file_partition=10:20:30:40:50
tranche_ringcount_partition=0:1:2:3:4:5
tranche_ringcount_file_partition=0:1:2:3:4:5
tranche_aromaticringcount_partition=0:1:2:3:4:5
tranche_aromaticringcount_file_partition=0:1:2:3:4:5
tranche_formalcharge_partition=-2:-1:0:1
tranche_formalcharge_file_partition=-2:-1:0:1
tranche_mr_jchem_partition=40:80:130
tranche_mr_obabel_partition=40:80:130
tranche_mr_file_partition=40:80:130
tranche_positivechargecount_partition=0:1
tranche_positivechargecount_file_partition=-2:-1:0:1:2
tranche_negativechargecount_partition=0:1
tranche_negativechargecount_file_partition=-2:-1:0:1:2
tranche_fsp3_partition=0.2:0:4:0:6:0.8
tranche_fsp3_file_partition=0.2:0.4:0.6:0.8
tranche_chiralcentercount_partition=0:1
tranche_chiralcentercount_file_partition=0:1:2:3:4:5
tranche_halogencount_partition=0
tranche_halogencount_file_partition=0:1:2:3:4:5
tranche_sulfurcount_partition=0
tranche_sulfurcount_file_partition=0:1:2:3:4:5
tranche_NOcount_partition=0:1:2:3:4:5:6:7:8:9:10
tranche_NOcount_file_partition=0:1:2:3:4:5:6:7:8:9:10
tranche_electronegativeatomcount_partition=0:1:2:3:4:5:6:7:8:9:10
tranche_electronegativeatomcount_file_partition=0:1:2:3:4:5:6:7:8:9:10
tranche_rotb_file_partition=0:2:4:7:10
tranche_aromaticproportion_jchem_partition=0:0.25:0.5:0.75
tranche_doublebondstereoisomercount_jchem_partition=0
tranche_qed_rdkit_partition=0.2:0.4:0.6:0.8:1.0

# For tranche types that are characters or strings (non-numeric)
# they should be in the format tranche_(tranche_type)_mapping=M:A,C:B
#  -- the above sample shows that a string value of "M" should
#     map to tranche of A, "C" to a tranche of B

tranche_enamine_type_mapping=M:A,S:B

****** Attributes

# In addition to the tranche_types above, attributes can be calculated and placed by AFLP
# as remarks in pdb, pdbqt, and mol2 files.

attributes_to_generate=scaffold_rdkit
# Multiple values are separated by colons
# Valid values:
#   * (any of the tranche_types above). e.g. mr_jchem
#   * scaffold_rdkit: Canonical SMI of Murcko scaffold

****** JChem-Related Packages

use_cxcalc_helper=0
# If using AWS Batch, then set this to 1. If not, then set to 0 currently. This option
# provides an optimization to improve performance of the chemaxon properties

jchem_package_filename=none
# Required only if cxcalc or molconvert (both of ChemAxon) are used in the preparation steps, and if tranche_assignments
     =true
```

```
# The filename of the JChem package located in the folder tools/packages/ in the tar.gz format (available on the
    ChemAxon homepage)
# The root folder in the archive has to have the name jchemsuite (normally distributed by ChemAxon in this way)
# Possible values:
#   * <filename>
#   * none

chemaxon_license_filename=none
# Required only if cxcalc or molconvert (both of ChemAxon) are used in the preparation steps.
# The filename of the license file, which has to be located in the folder tools/packages/
# Possible values:
#   * <filename>
#   * none

java_package_filename=none
# Required only if cxcalc or molconvert (both of ChemAxon) are used in the preparation steps.
# Any JRE binary distribution of version of at least version 8.
# If java is provided by the system (e.g. by loading a module), then no Java package needs to be provided.
# This has to be a file in the tar.gz format, which has to be located in the folder tools/packages/
# The root folder in the archive has to have the name "java" (which will be used for the JAVA_HOME variable). This
    normally needs to be manually changed in the Java package after downloading a JRE.
# Possible values:
#   * <filename>
#   * none

ng_package_filename=none
# Required only if cxcalc or molconvert (both of ChemAxon) are used in the preparation steps.
# Nailgun package filename.
# This has to be a file in the tar.gz format, which has to be located in the folder tools/packages/
# The root folder in the archive has to have the name "nailgun"
# Possible values:
#   * <filename>
#   * none

java_max_heap_size=4
# Size in GB
# Recommended: >= 1 GB * queues_per_step
# The required memory depends mainly on how many queues are run per step (and thus per JVM/NG server), since one JVM is
    used per step
```

#### L Configuration File of AFVS

All options of AFVS are specified in a central configuration file. A sample configuration file is shown in Supplementary Listing 2.

**Supplementary Listing 2.** AFVS Example Configuration File.

```
# Each line that starts with '#' is a comment/description of the variable above the comment
# Each line that starts with '*' belongs to a section heading

****** Job Resource Configuration

job_name=testing
# alphabetic characters (i.e. letters from a-z or A-Z)
# Used to describe distinct runs (using the same name will
# overwrite data if using S3!)

threads_per_docking=1
# How many threads should be used for each docking program.

threads_to_use=8
# This sets how many processes the main execution loop should be using
# to process. This is generally 2x the number of vCPUs or hyperthreads
# available on the system it is being run on

program_timeout=90
# How many seconds to wait for each ligand to be processed by a program

************************************************
** Batch system configuration
************************************************

batchsystem=awsbatch
# Possible values: awsbatch, slurm

****** AWS Batch Options (if batchsystem=awsbatch)

### To use AWS Batch you must first complete the steps outlined
### in the user guide for AWS Batch

aws_batch_prefix=af
# Prefix for the name of the AWS Batch queues. This is normally 'af'
# if you used the CloudFormation template

aws_batch_number_of_queues=2
# Should be set to the number of queues that are setup for AWS Batch.
# Generally this number is 2 unless you have a large-scale (100K+ vCPUs)
# setup

aws_batch_jobdef=af-jobdef-AFVS
# Generally this is [aws_batch_prefix]-jobdef-AFVS
# (e.g. if aws_batch_prefix=af, then aws_batch_jobdef=af-jobdef-AFVS

aws_batch_array_job_size=200
# Target for the number of jobs that should be in a single array job for AWS Batch.

aws_ecr_repository_name=af-AFVS-ecr
# Set it to the name of the Elastic Container Registry (ECR)
# repository (e.g. af-AFVS-ecr) in your AWS account
# (If you used the template it is generally af-AFVS-ecr)

aws_region=us-east-1
# Set to the AWS location code where you are running AWS Batch
# (e.g. us-east-1 for North America, Northern Virginia)

aws_batch_subjob_vcpus=8
# Set to the number of vCPUs that should be launched per subjob.
# 'threads_to_use' above should be >= to this value.

aws_batch_subjob_memory=15000
# Memory per subjob to setup for the container in MB

aws_batch_subjob_timeout=10800
# Maximum amount of time (in seconds) that a single AWS Batch job should
# ever run before being terminated.

****** Slurm Options (if batchsystem=slurm)

slurm_template=./templates/template1.slurm.sh
# Template for the slurm job
# Additional slurm attributes can be added directly to this
# template file if they are not available as pass throughs from
# AFVS

slurm_array_job_throttle=100
```

```
# Maximum number of jobs running within a single slurm array job

slurm_partition=partition
# Partition to submit the job

slurm_cpus=18
# Number of CPUs that are being used

slurm_array_job_size=100
# Maximum number of concurrent jobs from a single array job
# that should be run

***********************************************
** Storage configuration
***********************************************

job_storage_mode=s3
# This mode determines where data is retrieved and stored from as part of
# AFVS. Valid modes:
#   * s3: Job data is stored on S3 object store, which is the required
#         mode if using AWS Batch. Items under the "S3 Object Store"
#         heading in the configuration are required if this mode is used
#   * sharedfs: This mode requires that all running jobs have access to the
#         same shared filesystem that will allow for both input and output
#         of data. This is required if using Slurm

data_collection_addressing_mode=metatranche
# If input is placed with the hash addressing mode, then use 'hash'.
# otherwise use "metatranche" for the classic addressing mode

data_collection_identifier=
# This is only used if object_store_data_collection_addressing_mode=hash
# Generally this is the dataset name (e.g. Enamine_REAL_Space_2021q12)

job_addressing_mode=metatranche
# If job output is to be placed with the hash addressing mode, then use 'hash'.
# otherwise use "metatranche" for the classic addressing mode

****** Object Store Settings (S3)

object_store_job_bucket=
# Bucket name for the job data (output and job files)

object_store_job_prefix=jobs
# Where to place job-specific data. This includes where AdaptiveFlow will place
# the input data needed for jobs as well as the output files.
#
# Data be be placed:
# if object_store_job_addressing_mode=hash
#       in object_store_job_prefix/XX/YY/<job_letter>
#       (where 'XX', and 'YY' are hash values that will vary for
#        different files)
# else
#       in object_store_job_prefix/<job_letter>

object_store_data_bucket=
# Bucket name for the input collection data (often the same as the job one)

object_store_data_collection_prefix=
# Prefix used within the object store to address the collections

****** Shared Filesystem Settings

collection_folder=/home/ec2-user/collections
# Path to where the collection file are stored
#  * This is used when job_storage_mode=sharedfs or
#    when the uploader helper script is being used
#
# Slash at the end is not required (optional)
# Either pathname is required w.r.t. the folder tools/
#     or absolute path (e.g. /home/afuser/collections)

***********************************************
** Run configuration
***********************************************

****** Output information

summary_formats=parquet
```

```
# Format for summary files that are generated with the score data.
# Supported values:
#   * txt.gz (space delimited files)
#   * parquet
# Multiple formats can be generated by placing a comma
# (e.g. summary_formats=parquet,txt.gz)

print_smi_in_summary=1
# Whether or not the SMILES string should be printed in
# the summary file (works for pdbqt and mol2)
# Supported values:
#   * 0 : Do not provide
#   * 1 : SMILES string in the summary file

****** Workflow Options

ligands_todo_per_queue=1000
# Used as a limit of ligands for the to-do lists.
# A reasonable number for this is generally 1000. The length of time
#      to process will depend on the docking scenarios run

ligand_library_format=pdbqt
# Supported values:
#   * pdbqt
#   * mol2
# This value is case sensitive
# All AutoDock based docking programs require the library to
#      be in the pdbqt format.
# When the docking program PLANTS is used, both libraries in the
#      pdbqt and the mol2 format are supported.

tempdir_default=/dev/shm
# The directory which is used for the temporary workflow files which need a normal performance
# Is normally a local SSD or HDD. A temporary directory will be created underneath this dir

tempdir_fast=/dev/shm
# The directory which is used for the temporary workflow files which need a fast perfomance
# Should be a a local ram filesystem/ramdisk. A temporary directory will be created underneath this dir

****** Virtual Screening Options

docking_scenario_names=qvina02_rigid_receptor1
# Names for the docking scenarios, separated by colons
# Each docking scenario has one value. Multiple docking scenarios/names have
#     to be separated by colons ":" and without spaces
#
# Example: docking_scenario_names=receptor1_vina_rigid:receptor1_smina_flexible
# The docking scenario names are used for the folder names in which the output files are stored

docking_scenario_programs=qvina02
# For each docking scenario name, a docking protocol has to be specified
# Values have to be separated by colons ":" and without spaces, e.g: docking_scenario_programs=vina:smina
# Possible values:
#      AutoDock-Koto,  AutodockVina_1.2,  AutodockZN,   EquiBind,      FRED
#      FitDock,        GalaxyDock3,       LigandFit,    LightDock,     M-Dock
#      MCDock,         MM-GBSA,           PLANTS,       PSOVina,       RLDock
#      SEED,           adfr,              autodock_cpu, autodock_gpu,  autodock_vina
#      dock6,          flexx,             glide,        gnina,         gnina-scoring
#      gold,           gwovina,           iGemDock,     idock,         ledock
#      molegro,        nnscore2,          qvina,        qvina-w,       rDock
#      rf-score,       rosetta-ligand,    smina,        smina-scoring, vina
#      vina_carb,      vina_xb
# Please note: different pose prediction/docking protocols can be combined with scoring functions.
#              For example: 'qvina+nnscore2'.
# For supported choices/combinations please see the AdaptiveFlow homepage.

docking_scenario_replicas=1
# Series of integers separated by colons ":"
# The number of values has to equal the number of docking programs
#      specified in the variable "docking_programs"
# The values are in the same order as the docking programs specified in the
#      variable "docking_scenario_programs
#      e.g.: docking_scenario_replicas=1:1
# possible range: 1-99999 per field/docking program
# The docking scenario is comprised of all the docking types and their replicas

docking_scenario_basefolder=../input-files
# Relative path to tools directory
# Base directory for where the docking scenarios are held. Nothing other
# than the required files for the docking scenario should be placed here

docking_scenario_inputfolders=qvina02_rigid_receptor1
# folder names inside 'docking_scenario_basefolder'
```

```
# In each input folder must be the file config.txt which is used by the
#     docking program to specify its options
# If other input files are required by the docking type, usually
#     specified in the config.txt file, they have to be in the same folder
```

#### M Configuration File of AdaptiveFlow Unity

All options of AdaptiveFlow Unity are specified in a central configuration file when used in standalone mode. A sample configuration file is shown in Supplementary Listing 3.

**Supplementary Listing 3.** AdaptiveFlow Unity (AFU) Example Configuration File.

```
# The choice of the docking protocol
# Possible choices:
#     AutoDock-Koto,  AutodockVina_1.2,  AutodockZN,   EquiBind,      FRED
#     FitDock,        GalaxyDock3,       LigandFit,    LightDock,     M-Dock
#     MCDock,         MM-GBSA,           PLANTS,       PSOVina,       RLDock
#     SEED,           adfr,              autodock_cpu, autodock_gpu,  autodock_vina
#     dock6,          flexx,             glide,        gnina,         gnina-scoring
#     gold,           gwovina,           iGemDock,     idock,         ledock
#     molegro,        nnscore2,          qvina,        qvina-w,       rDock
#     rf-score,       rosetta-ligand,    smina,        smina-scoring, vina
#     vina_carb,      vina_xb
# Please note: different pose prediction/docking protocols can be combined with scoring functions.
#              For example: 'qvina+nnscore2'.
# For supported choices/combinations please see the AdaptiveFlow homepage.

program_choice=qvina

# The x,y&z coordinates of the center of the docking space. The binding space describes the location where a molecule is
      allowed to bind.
center_x=10
center_y=10
center_z=10

#

# The size (in Angstroms) of the docking space in the x,y&z directions.
size_x=10
size_y=10
size_z=10

# How many poses to search for, providing a limit to the maximum number of iterations that a docking program performs in
      search for good poses
# Large exhaustiveness settings lead to increased computational costs.
exhaustiveness=10

# Molecule (either a string in smiles, selfies or amino-acid sequence)
# If the is_selfies=True, or is_peptide=True, a conversion from selfies->smiles
# and aa-sequence->smiles is performed.
smi=CCCCCCCCCCCCC
is_selfies=False
is_peptide=False

# Location to the prepared receptor file
#  The receptor needs to be in the correct format, supported by the user's selected docking program.
#  Additionally, the file needs to be present in the config directory.
receptor=./config/5wiu_test.pdbqt
```
